# Evidences for a nutritional role of iodine in plants

**DOI:** 10.1101/2020.09.16.300079

**Authors:** Claudia Kiferle, Marco Martinelli, Anna Maria Salzano, Silvia Gonzali, Sara Beltrami, Piero Antonio Salvadori, Katja Hora, Harmen Tjalling Holwerda, Andrea Scaloni, Pierdomenico Perata

## Abstract

Little is known about the role of iodine in plant physiology. We evaluated the impact of low concentrations of iodine on the phenotype, transcriptome and proteome of *Arabidopsis thaliana*. Our experiments showed that removal of iodine from the nutrition solution compromises plant growth, and restoring it in micromolar concentrations is beneficial for biomass accumulation and leads to early flowering. In addition, iodine treatments specifically regulate the expression of several genes, mostly involved in the plant defence response, suggesting that iodine may protect against both biotic and abiotic stress. Finally, we demonstrated iodine organification in proteins. Our bioinformatic analysis of proteomic data revealed that iodinated proteins identified in the shoots are mainly associated with the chloroplast and are functionally involved in photosynthetic processes, whereas those in the roots mostly belong and/or are related to the action of various peroxidases. These results suggest that iodine should be considered as a plant nutrient.

## INTRODUCTION

Plants need macro- and micro-nutrients for their growth and development. Nutrients are chemical elements that are components of biological molecules and/or influence essential metabolic functions. The elements that to date are currently considered as plant nutrients are C, H, O, N, P, K (primary nutrients), Ca, Mg, S (secondary nutrients), and Fe, Zn, Cu, Mn, B, Cl, Mo, Co and Ni (micro-nutrients) (Mengel & Kirkby, 2001). Selenium (Se) is an essential element for animals, while its classification as a micro-nutrient for plants is still controversial (Gupta & Gupta, 2017).

Halogens are the least represented chemical group of plant micro-nutrients, chloride being the only micro-nutrient currently recognised in plant physiology, due to its regulatory action in proton-transfer reactions in the photosystem II (Brahmachari, Guo, Konecny, Obi & Barry, 2018). Studying the effect of different concentration and forms of iodine on the growth of several crops of agricultural importance, Borst Pauwels (1961) referred to iodine as a micro-nutrient for plant. A growing number of recent studies have focused on the benefit of increasing iodine content in plants as a biofortificant in human and animal health (Medrano-Macías, Leija-Martínez, González-Morales, Juárez-Maldonado & Benavides-Mendoza, 2016; Gonzali, Kiferle & Perata, 2017). These studies rarely address the impact of iodine on the physiology of the plant.

At the very beginning of life, many microorganisms such as cyanobacteria started using iodine as an antioxidant compound (Crockford, 2009; Venturi, 2011). Later on, when several organisms (including plants) evolved to live in terrestrial areas, a new set of antioxidants (i.e. ascorbic acid, polyphenols, carotenoids) replaced iodine. Later, novel roles for iodine emerged (Venturi, 2011). In vertebrates, iodine became a key constituent of triiodothyronine (T_3_) and L-thyroxine (T_4_), the two thyroid hormones (THs) that regulate energy metabolism and other essential physiological processes (Zimmermann, Jooste & Pandav, 2008; Crockford, 2009; Mourouzis, Lavecchia & Xinaris, 2020). Iodine can also be incorporated into iodolipids, which have antineoplastic effects (Aceves, Anguiano & Delgado, 2005; Mendieta et al., 2019), and into proteins (Ramachandran, 1956). In proteins, iodination affects various amino acids, depending on the reaction conditions (Ramachandran, 1956), but generally following the reactivity order Tyr >> His > Trp > Cys. In fact, Tyr iodination has been highly characterized in thyroglobulin, a glycoprotein produced from the thyroid gland in vertebrates (Dedieu, Gaillard, Pourcher, Darrouzet & Armengaud, 2011; Xiao, Dorris, Rawitch & Taurog, 1996).

The incorporation of iodine in proteins is therefore one key aspect of the role of iodine as a micro-nutrient for vertebrates.

Plants can absorb iodine from roots or above ground structures (stomata and cuticular waxes) (Medrano-Macías et al., 2016; Gonzali et al., 2017), translocate it mainly through the xylematic route and volatilize it as methyl iodide (CH_3_I) through the action of halide ionmethyltransferase (HMT) and halide/thiol methyltransferase (HTMT) enzymes (Medrano-Macías et al., 2016; Gonzali et al., 2017).

Little is known about the chemical forms of iodine inside plant tissues. Inorganic iodine, iodide (I^−^), however, seems to be predominant (Weng et al., 2008). Plants can also incorporate iodine into organic molecules, such as T_3_ or mono-iodotyrosine (MIT) and di-iodotyrosine (DIT), which are precursors of T_3_ and T_4_ THs in vertebrates (Eales, 1997; Smoleń et al., 2020). The metabolic role of these molecules and their biosynthetic mechanism are still unknown. Protein iodination, which has been verified in several seaweed species (Hou, Yan & Chai, 2000; Romarís-Hortas, Bianga, Moreda-Piñeiro, Bermejo-Barrera & Szpunar, 2014), has not yet been demonstrated in plants.

Iodine is likely involved in several physiological and biochemical processes (Medrano-Macías et al., 2016; Gonzali et al., 2017). Most of the findings regarding the effect of iodine on plant growth define protocols for agronomic biofortification of plants with iodine, in which exogenous iodine is added to the growth medium or spray applied to the crop to increase its availability in the edible part of the plants for human and farm-animal nutrition. In these experiments, however, the control plants are not subjected to iodine enrichment, and are generally grown in iodine-containing soil/solutions, thus preventing the effects related to the presence/absence of this element from being easily observed (Fuge & Johnson, 1986; Ashworth, 2009). The functional role of iodine as a plant nutrient might therefore have been masked.

Plant tissues generally increase their iodine content following its exogenous administration. On the other hand, biomass production, stress resistance or fruit quality can be affected with positive but also negative effects depending on the form and concentration of the iodine applied, the plant species, and the environmental/growing conditions (Medrano-Macías et al., 2016; Gonzali et al., 2017).

The use of low concentrations of iodine is often associated with beneficial effects on plant growth and production, whereas toxic effects are observed when applying iodine at high concentrations, especially in the I^−^ form, which is more phytotoxic than iodate (IO_3_^−^) (Medrano-Macías et al., 2016; Gonzali et al., 2017; Incrocci et al., 2019). Thresholds for beneficial or toxic concentrations have been reported for all micro-nutrients (Welch & Shuman, 1995). Interestingly, the concentrations of iodine added to nutrient solutions that have been associated with positive effects for plants (ranging from approximately 10^2^ to 10^4^ nM) (Medrano-Macìas et al., 2016, Gonzali et al., 2017) are comparable with those generally effective for other elements described as plant micro-nutrients (Sonneveld, 2002), suggesting that iodine plays a similar role in plant nutrition.

We explored the role of iodine as a nutrient for plants using various experimental approaches. Our results showed that iodine, when supplied at a well-defined concentration range, positively affected the phenotype of *Arabidopsis thaliana* plants, and altered the organism’s transcriptome. Most importantly, protein iodination was observed for the first time. These results are strongly suggestive of the role of iodine as a plant nutrient.

## MATERIALS AND METHODS

### Plant material and cultivation system

Plants of *Arabidopsis thaliana*, ecotype Columbia 0, *Solanum lycopersicum* L., cv. Micro-Tom, *Lactuca sativa* L., var. crispa, *Triticum turgidum* L., var. durum, and *Zea mays* L., var. saccharata, were used in the experiments.

Seeds of the different species were sown on rockwool plugs and vernalized for three days. After this period, plants were hydroponically cultivated in a growth chamber (22 °C day/18 °C night with a 12-h photoperiod, a quantum irradiance of 100 μmol photons m^−2^ sec^−1^ and a relative humidity close to 35%), in a floating system. A base nutrient solution, renewed once a week, was prepared minimising iodine contamination by dissolving in MilliQ water, the following amounts of ultrapure salts: 1.25 mM KNO_3_, 1.50 mM Ca(NO_3_)_2_, 0.75 mM MgSO_4_, 0.50 mM KH_2_PO_4_, 50 μM KCl, 50 μM H_3_BO_3_, 10 μM MnSO_4_, 2.0 μM ZnSO_4_, 1.5 μM CuSO_4_, 0.075 μM (NH_4_)_6_Mo_7_O_24_ and 72 μM Fe-EDTA. At preparation, the pH and the electrical conductivity (EC) values were 6.0 and 0.6 dS m^−1^, respectively, whereas the iodine concentration in the nutrient solution was below the detection limit of 8 nM, as determined by ICP-MS analysis.

### Phenotypical determinations

Two separate experiments were performed. In both experiments, Arabidopsis plants were initially fed with the base nutrient solution. After 15 days of growth, plants homogeneous in size and leaf number were selected, grouped, and fed with different concentration and/or type of halogen-containing salts added to the nutrient solution. Plants were distributed in nine separate hydroponic trays (three different trays/treatment), and a total number of 90 plants were cultivated for each experimental condition (30 plants/tray).

In the first experiment (exp. 1), 0.20 or 10 μM KIO_3_ was added to the nutrient solution. One month later, during the formation/elongation of the main inflorescence, half of the plants (15 plants/tray) were harvested and characterized according to the main morphological traits, such as rosette and inflorescence fresh weight (FW), dry weight (DW), dry matter content, rosette diameter, and inflorescence length. The remaining half were allowed to complete the growing cycle and were characterized in terms of flowering and seed production. Flowering, defined as the presence of the first open flower on the stem, was recorded at intervals of three days and expressed on a percentage basis. The percentage of bloomed plants/tray was calculated at each assessment date. Toward the final part of the plants’ life cycle, a periodical harvesting of the produced/matured seeds was carried out until the complete plant desiccation. Seed production was determined in terms of total seed weight/tray (15 plants/tray), number of seeds/silique and number of siliques/plant.

In the second experiment (exp. 2), plants were treated by adding either KI, NaI or KBr (0, 10 or 30 μM) to the nutrient solution. Fifteen days after the salt treatment, half of the plants was characterized in terms of plant FW, DW and dry matter content, while the other half was subsequently characterized in terms of flowering (recorded with intervals of 2 days), as described for Experiment 1.

### Total RNA extraction and processing

Gene expression analysis was performed on 3-week-old Arabidopsis plants hydroponically grown on the base nutrient solution (control plants) or in the same medium to which 10 μM of KBr, NaI or KI was added. Plant material was collected 48 h after the beginning of the treatment. Each sample consisted of a pool of rosettes or roots sampled from three different plants, which were immediately frozen in liquid nitrogen and stored at −80 °C until further analysis.

Total RNA from rosettes was extracted as described by Perata et al. (1997), avoiding the use of aurintricarboxylic acid. RNA from roots was extracted using the Spectrum™ Plant Total RNA Kit (Sigma-Aldrich). RNA was subsequently processed for microarray and qPCR analysis. The TURBO DNA-free kit (Thermo Fisher Scientific) was used to remove DNA contaminations and the iScript TM cDNA synthesis kit (Bio-Rad Laboratories) was used for RNA reverse-transcription.

### Microarray analysis

RNA from rosettes and roots was processed and hybridized to Affymetrix GeneChip Arabidopsis ATH1 Genome Arrays as described by Loreti et al. (2005). Normalization was performed using Microarray Suite 5.0 (MAS5.0). Differentially expressed genes (DEGs) were selected based on the two following criteria: log_2_FC treated/control≥2 and mas5-Detection *p*-value ≤ 0.05. In addition, the absolute expression level ≥100 mas5_Signal was chosen to select only well-expressed genes.

DEGs commonly regulated by KI- and NaI-treated plants and not responding to KBr treatments were considered specifically linked with the iodine treatment and were subjected to gene set enrichment using GOrilla (http://cbl-gorilla.cs.technion.ac.il) and analyzed with Mapman (http://mapman.gabipd.org/mapman), whereas the co-expression analysis was performed using Genevestigator (https://genevestigator.com).

### Gene expression analysis

Quantitative PCR (ABI Prism 7300 Sequence Detection System, Applied Biosystems) was performed using 30 ng cDNA and the iQ SYBR Green Supermix (Bio-Rad Laboratories). *UBIQUITIN10* (At4g05320) and *TIP4* (At2g25810.1) were used as reference genes. Relative expression levels were calculated using GeNorm(http://medgen.ugent.be/~jvdesomp/genorm). The list of the primers and their sequences are reported in Table S1. Four biological replicates were analyzed, each consisting in a pool of rosettes or roots sampled from three different plants.

### Feeding with radioactive iodine

Two separate experiments were performed by feeding radioactive iodine (^125^I - NaI, PerkinElmer) to hydroponically grown *Arabidopsis thaliana* (Experiment 1) or tomato, lettuce, wheat and maize plants (Experiment 2). Treatments were performed on one-month-old plants. The solution of ^125^I was prepared by dissolving 60 μl of the commercial radioactive ^125^I product (2.4 mCi/100 μl – 9,41 μM) in 250 ml of base nutrient solution. Plants were individually transferred into plastic tubes and treated with the hydroponic solution (with or without Na^125^I). Sampling was performed after 48 h of incubation by collecting leaf and root material, which was immediately frozen in liquid nitrogen, and stored at −80 °C until the analysis. Control, non-treated plants (no ^125^I added during their growth) were used in both experiments.

### Protein extraction, electrophoresis, and gel autoradiography

Leaf and root samples from ^125^I-fed and control plants were ground to fine powder in liquid nitrogen. The protein extraction buffer (50 mM TrisHCl, pH 7.0, 1% w/v SDS, P9599 protease inhibitor cocktail, Sigma-Aldrich) was added to the powder. The resulting solution was vortexed vigorously, and then centrifuged (18,407 g, 30 min, 4 °C). Radioactive iodine solution (10 μl; prepared as described above) was added to the control samples during the extraction process to check for the occurrence of false positive signals (technical artifacts), possibly due to unspecific bounds of iodine with the protein extract.

Protein extracts were dissolved in a 5×Laemmli buffer, treated at 95 °C for 10 min, and a volume of 20 μl (corresponding, approximately, to 65 or 20 μg of proteins, in shoot and root samples, respectively) was loaded to Invitrogen NuPAGE gels (10% Bis-Tris Midi Gels, Thermo Fisher Scientific), together with protein markers (Precision Plus Protein™ Dual Color Standards, Bio-Rad). After electrophoresis, the gel was rinsed in MilliQ water, and the proteins were fixed, stained with EZ Blue Gel Staining reagent (Sigma-Aldrich) for 30 min and finally exposed to a multipurpose phosphor storage screen (Cyclone Storage Phosphor System, PerkinElmer) in order to obtain a digital image of the radioactivity distribution. Radioactive signals were quantified after 72-h of gel exposure using a Cyclone Phosphor Imaging System (PerkinElmer). In order to prevent the occurrence of any radioactive emissions from the control samples, after each image acquisition, gels were re-exposed for 15 days, and the absence of ^125^I labelled bands was verified in the newly-acquired images.

### Database search for iodinated peptides in protein datasets from proteomic data repositories

Mass spectrometry data were downloaded from the PRIDE (PRoteomics IDEntification database) archive (https://www.ebi.ac.uk/pride/archive) (Perez-Riverol et al., 2019). The PRIDE archive was searched to select *A. thaliana* datasets based on the analysis of specific plant organs, such as cauline, rosette and roots, and/or subcellular districts, such as chloroplasts and mitochondria. Datasets were excluded if enrichment/immunopurification strategies were used during protein purification. Finally, 21 experimental sets of nano-LC-ESI-MS/MS raw data included in 14 PRIDE repository (March 2020) were obtained and re-analysed by database searching. Only raw files corresponding to the analysis of control/non-treated plants were downloaded and the experimental protocols and the search parameters for each different dataset were annotated. Table S2 lists the experimental sets, with details on the MS instrument, plant organ and/or subcellular compartment, sample preparation, and proteomic strategies adopted.

Raw files were searched separately using Proteome Discoverer 2.4 (Thermo, USA) with the Mascot v. 2.6 search engine (Matrix Science Ltd, UK) against the TAIR10 database (www.arabidopsis.org; 71,567 sequences, accessed May 2017) and a database containing common laboratory contaminants on the MaxQuant website (https://www.maxquant.org/maxquant/). Workflows were built for each experimental dataset, considering the specific mass tolerance values used for the original search and reported on the PRIDE repository, or in the publication associated with the dataset.

For all the workflows, Cys carbamidomethylation was set as a fixed modification, while iodination at Tyr and His (Δm = +125,8966 Da), oxidation at Met, protein N-terminal acetylation, deamidation at Asp, and pyroglutamate formation at N-terminal Gln were selected as variable modifications. Isotopic labeling was also considered in the modification parameters when performed for protein quantification.

Trypsin was selected as the proteolytic enzyme and peptides were allowed to have a maximum of two missed cleavages. The minimum peptide length was set at six amino acids. The site probability threshold for peptide modification was set at 75. Only high confidence peptide identifications were retained by setting the target false discovery rate (FDR) for PSM at 0.01 and further filtered to keep only peptides (P<0.05) with a Mascot Ion score >30. In addition, the results of the identification analysis were processed by putting together the output of iodinated peptides from all the datasets and further applying a filter to keep only those identified with a Mascot Ion score >50, in at least one dataset, to limit the identification to peptides with the best scoring matches and corresponding to high certainty. The presence of the MS/MS spectrum of the unmodified counterpart was verified for each iodinated peptide to further validate the identification.

### Bioinformatics

Proteins containing iodinated peptides were functionally annotated according to MapMan categories by using the Mercator pipeline (http://mapman.gabipd.org/web/guest/app/mercator). Final outputs were integrated with data from the available literature. Protein interaction networks were obtained with STRING v. 11 (http://string-db.org), which was also used to provide information on known gene ontology categories. Venn diagrams were depicted using a web tool at http://bioinformatics.psb.ugent.be/webtools/Venn. The Protein Abundance Database (PAXdb) was also queried to evaluate quantitative levels of modified *A. thaliana* proteins at https://pax-db.org.

### Statistical analysis

Experimental data were analyzed by one-way ANOVA coupled with the LSD post hoc test, when they followed a normal distribution and there was homogeneity of variances. When one of these two prerequisites was violated, a Kruskal-Wallis test for non-parametric statistic was performed and the significance letters were graphically assigned using a box-and-whisker plot with a median notch. Significant differences between the means (P<0.05) are indicated by different letters in the figures and tables.

## RESULTS

### Effects of iodine on the plant phenotype

The effects of low amounts of KIO_3_ (0.20 or 10 μM) on hydroponically grown Arabidopsis plants, compared to plants cultivated on a control nutrient solution were evaluated in terms of plant morphology, biomass, and seed production (Experiment 1). No phytotoxicity symptoms were observed on plants and the most significant phenotypical effect observed was a delay of flowering in the control plants, compared to KIO_3_ (either 0.20 or 10 μM) (Figure 1a,b). Twelve days after the opening of the first flower, plants treated with 0.20 or 10 μM KIO_3_ were close to complete flowering, as 87% and 96% of the plants had bloomed respectively, vs. 69% of the control plants (Figure 1b). Control plants took about 18 days to complete blooming.

**FIGURE 1.**
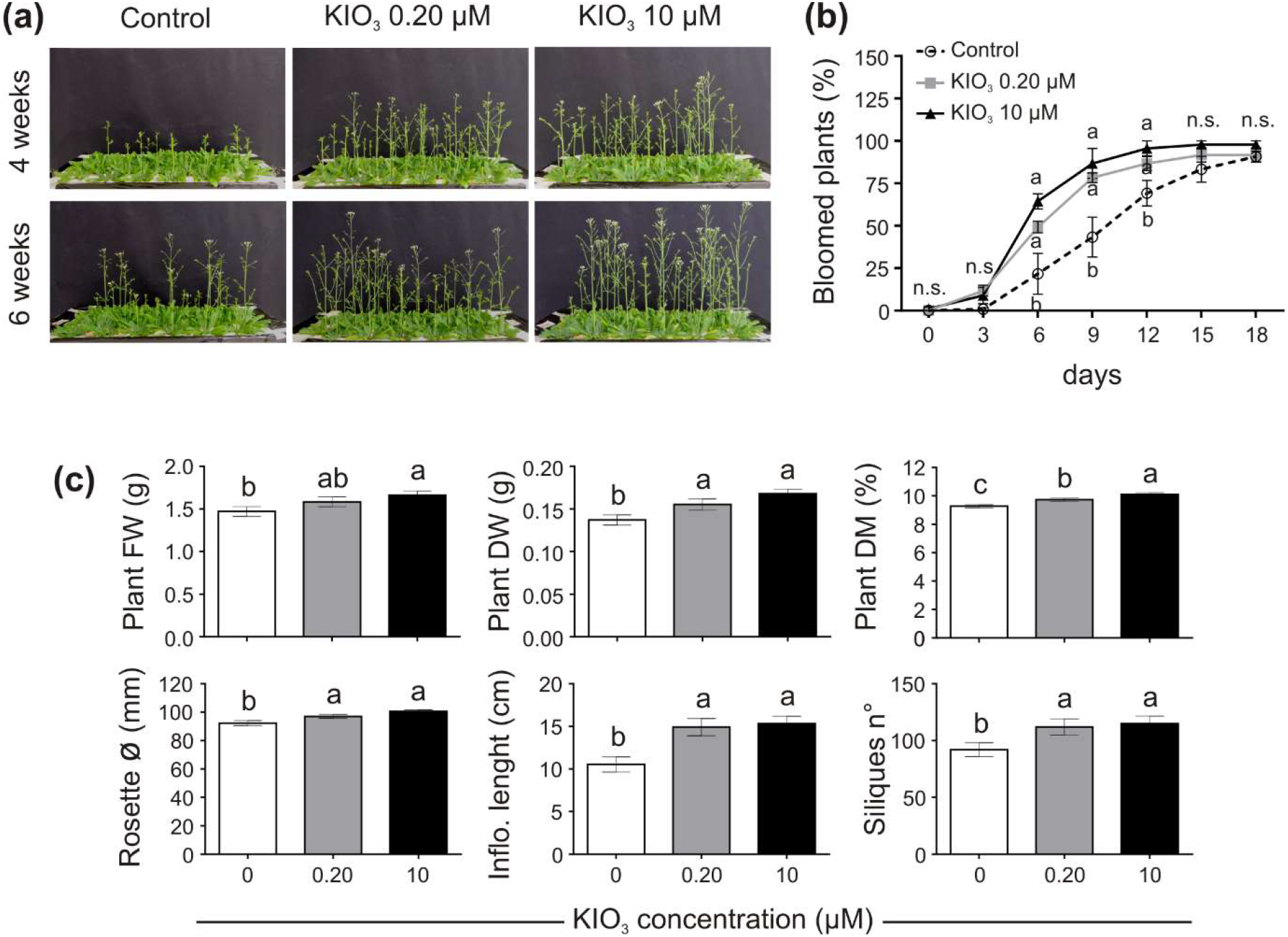
Impact of iodine on plant growth and development (exp. 1). (a), lateral view of plants after 4 or 6 weeks from the onset of KIO_3_ treatment. (b), flowering time curve; the percentage of bloomed plants/tray was calculated every three days after the opening of the first flower (day 0). (c), morphological data on plant FW, DW, dry matter content, rosette diameter, inflorescence length and number of produced siliques/plant, determined one month after the onset of KIO_3_ treatments. When data followed a Normal distribution and there was homogeneity of variances, they were subjected to one-way ANOVA and values indicated by different letters significantly differ from each other (LSD posthoc test, P ≤ 0.05). When one of this two prerequisite was violated, a Kruskal-Wallis test was performed. Statistical analysis of flowering (c) was performed by comparing the percentage of bloomed plants of each tray (considered as biological replicates) within each sampling point. Error bars (±SE) are shown in graphs.

Plant biomass, evaluated one month after the addition of KIO_3_ to the nutrient solution, was significantly lower in control plants, both in FW and DW (Figure 1c). When compared to the control, the plant FW increased by approximately 7.7% and 13% with addition of 0.20 and 10 μM KIO_3_ in the nutrient solution, respectively, whereas the DW increased by 13% and 22%, respectively. The effect on plant FW was mostly ascribable to the inflorescence, as no significant differences were evident in terms of the rosette FW values (Table S3). The concentration of iodine in the nutrient had a marked effect on the inflorescence length, which was approximately 41% and 45% longer compared to the control in 0.20 and 10 μM KIO_3_ treated plants, respectively (Figure 1c) and a comparable effect was seen on the inflorescence’s FW and DW (Table S3). Additionally, the rosette diameter in the control was smaller, and the application of 0.20 or 10 μM KIO_3_ improved rosette diameter by approximately 5% and 9%, respectively (Figure 1c). The plant dry matter content correlated positively with the increased iodate concentrations (Figure 1c).

Seed production was determined in terms of total seed weight, seeds/silique and number of siliques/plant. The number of seeds contained in each silique was not affected by iodate treatments (Table S3), whereas the number of siliques produced by each plant was lower in the control, compared to addition of both 0.20 and 10 μM KIO_3_ (Figure 1c). This influenced the total seed production, which, one month after the addition of KIO_3_ to the nutrient solution, was much higher in plants treated with iodate (more than 50% and 35%, respectively, in 0.20 and 10 μM KIO_3_ treated plants in comparison with the control) (Table S3).

Adding exogenous iodine in the form of KIO_3_, countered the delay in flowering of plants grown with a very low concentration of iodine in the nutrient solution in the control (Figure 1a,b). This was confirmed in experiment 2 (exp.2), when iodine was added in the form of KI or NaI (Figure 2a,b). The possible influences of potassium or bromide, as an additional halogen, were evaluated and then ruled out, as a similar behavior was observed in plants treated with KI or NaI, but not with KBr (Figure 2a,b).

**FIGURE 2.**
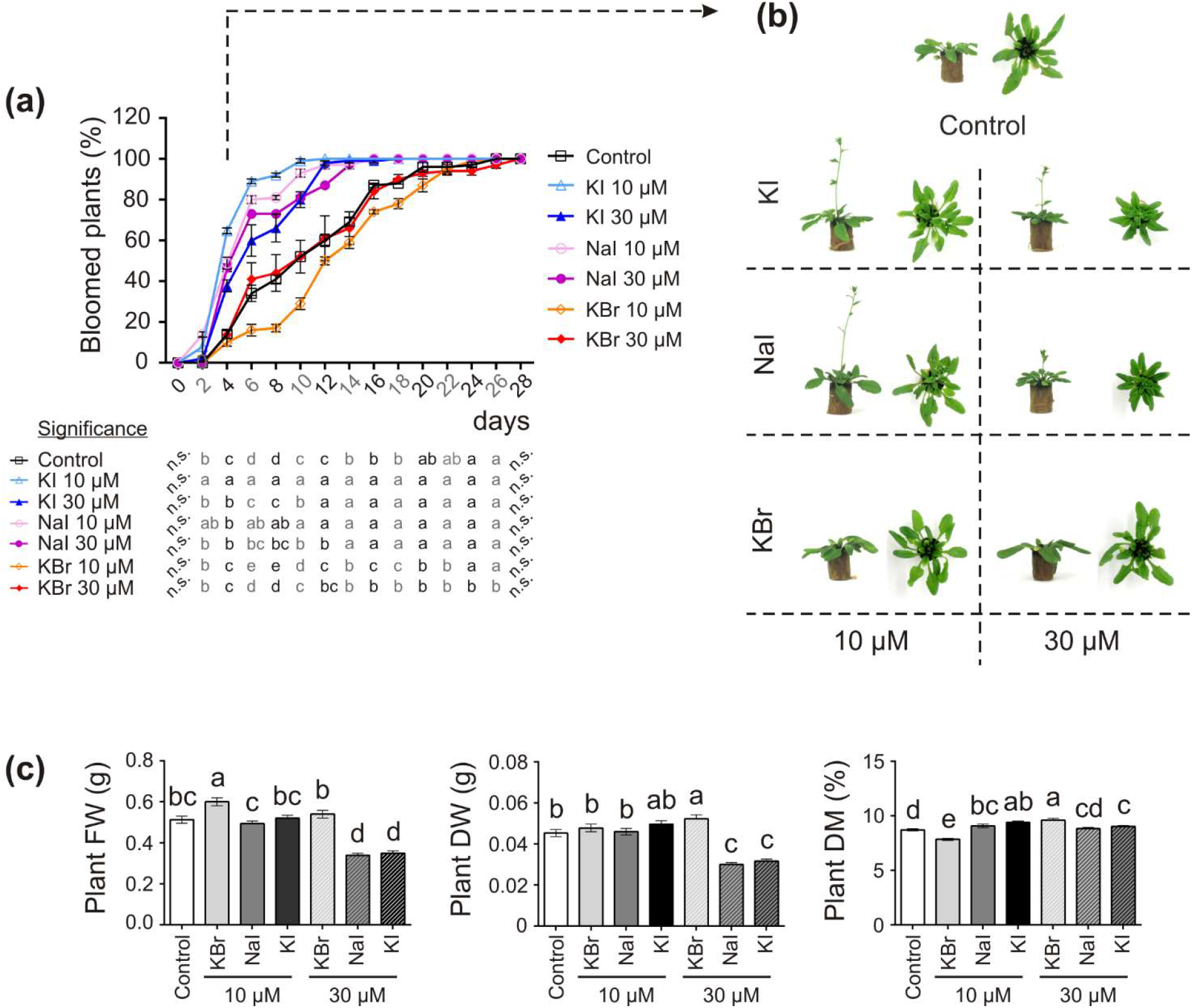
Impact of iodine on plant growth and development (exp.2). (a), flowering curve; the percentage of bloomed plants/tray was calculated every two days after the opening of the first flower (day 0). (b), representative control, and KI-, NaI- or KBr-treated plants (10 and 30 μM) after 15 days from the onset of the treatments. Pictures were taken after 4 days from the opening of the first flower on the main stem. (c), morphological data on plant FW, DW, dry matter content determined 15 days after the onset of the treatments. When data followed a Normal distribution and there was homogeneity of variances, they were subjected to one-way ANOVA and values indicated by different letters significantly differ from each other (LSD posthoc test, P≤0.05). When one of this two prerequisite was violated, a Kruskal-Wallis test was performed. Statistical analysis of flowering (c) was performed by comparing the percentage of bloomed plants of each tray (considered as biological replicates) within each sampling point. Error bars (±SE) are shown in graphs.

The application of 10 μM KI and NaI promoted flowering, without negatively impacting the plant biomass production (Figure 2c), whereas 30 μM KI or NaI reduced plant growth (Figure 2c), although the promoting effect of iodine on flowering was still present (Figure 2a,b). Four days after the opening of the first flower (day 0), more than 50% of KI- and NaI-treated plants had bloomed vs. 14% of the control plants and 10% and 14% of the 10 and 30 μM KBr-treated plants, respectively. Moreover, the floral transition was almost complete in 10 μM KI- and NaI-fed plants in the subsequent six days (10 days after day 0). Two and four more days were required for 30 μM KI- and NaI-fed plants, respectively (12 and 14 days after day 0), whereas the control and KBr-treated plants completed blooming in the subsequent 18 days (22 days after day 0) (Figure 2a).

### Effects of iodine on gene expression

The response of plants to iodine was analysed at the transcriptomic level. To identify genes whose expression was specifically altered by iodine, Arabidopsis plants were treated by adding 10 μM of NaI, KI or KBr to the nutrient solution, compared to the untreated control plants. The resulting RNAs were analysed by hybridization on ATH1 microarrays. To rule out the possible generic effects of halogens, we searched the microarray dataset for genes that responded to KI and NaI, but not to KBr. In addition, a comparison between KBr- and KI-treated plants enabled us to rule out the possible transcriptional regulation of genes exerted by potassium, as K^+^ ion was common to both salts.

Data visualization with a Venn diagram showed that several genes were specifically regulated by iodine, as up- or down-regulated genes in both NaI- and KI-but not in KBr-treated plants were 33 (51% of DEGs) and 15 (33% of DEGs) in the shoot (Figure 3a), and 398 (95% of DEGs) and 133 (79% of DEGs) in the root (Figure 3b), respectively. The similarity and specificity in the expression pattern of KI- and NaI-treated plants were confirmed by the heatmaps generated from the analysis of the shoot (Figure 3c) and root (Figure 3d) expression data.

**FIGURE 3.**
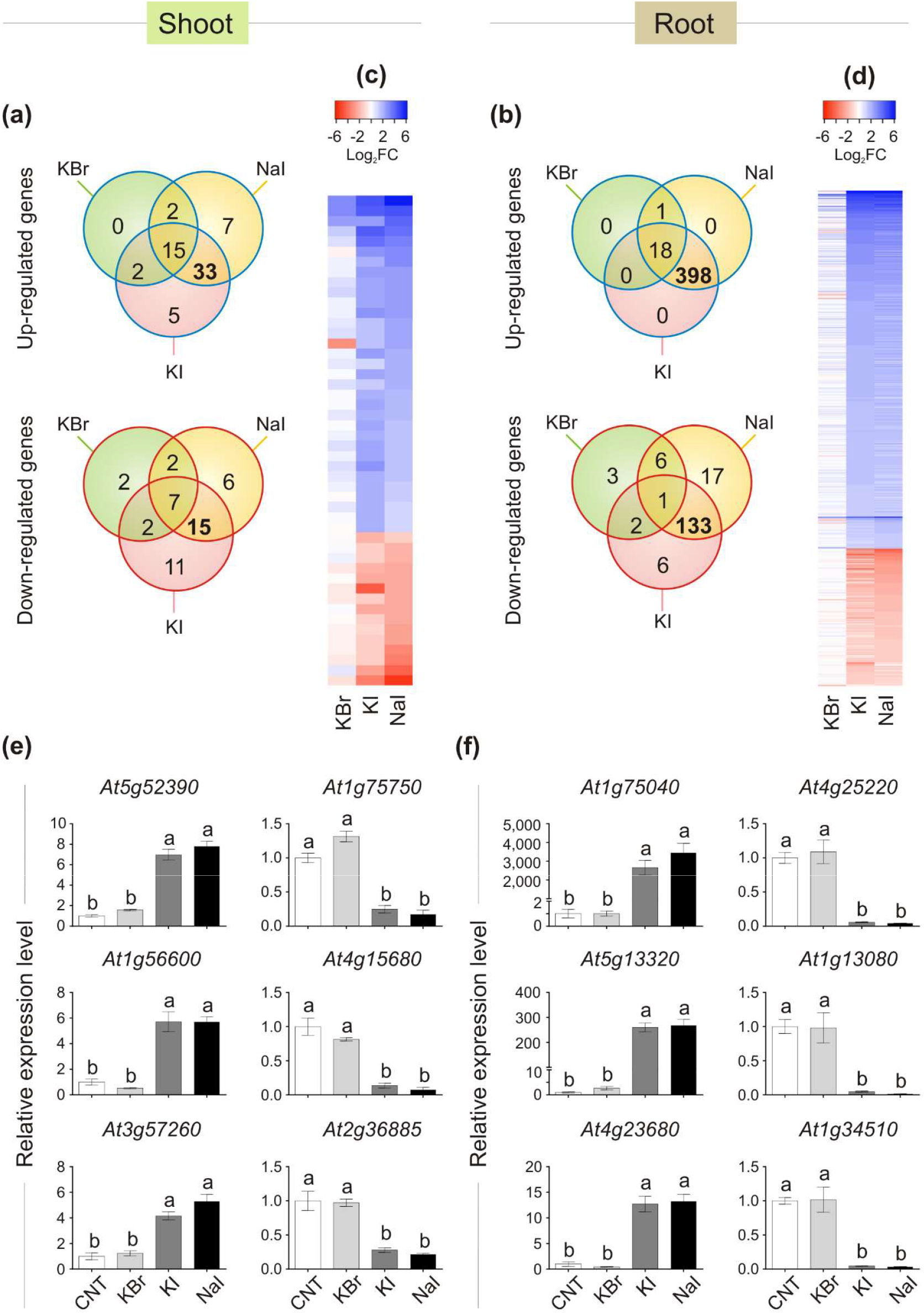
Transcriptional regulation of gene expression induced by iodine. Venn diagram showing the number of genes differentially regulated in shoot (a) or root (b) tissues of KBr-, NaI- and KI-treated plants, when compared with the control. Heatmap showing the pattern of expression of the genes analyzed in the shoot (c) or root (d) tissues that were commonly up- or down-regulated only in NaI- and KI-treated plants, and not in KBr-treated plants, when compared with the control. qPCR validation of selected genes up- or down-regulated by iodine treatments in shoot (e) or root (f) tissues. Data are mean ± SE of four biological replicates, each composed of a pool of three different rosettes. Values indicated by different letters significantly differ from each other (according with one-way ANOVA, LSD posthoc test, P≤0.05).

To validate the microarray analysis, a subset of three I^−^-induced and three I^−^-repressed genes were analyzed by qPCR, corroborating the specific regulation of iodine on their expression in both shoot (Figure 3e) and root (Figure 3f) samples.

The complete list of the KI and NaI commonly up- and down-regulated genes is reported in Tables S4 and S5 (shoot tissue), and Tables S6 and S7 (root tissue), respectively.

The polypeptides codified by the iodine-regulated genes did not show a preferential site of action in the cell, as their predicted localizations include the cytoplasm, chloroplast, cell wall, nucleus, mitochondrion, vacuole and apoplast (Tables S4, S5, S6, S7).

The gene ontology (GO) analysis identified several functional categories regulated by iodine in the roots (Figure 4; Figure S1; Tables S8 and S9), whereas no statistically significant GO terms were identified by analyzing the DEG data on the shoots.

The most representative biological processes affected by iodine in the roots were related to the *response to stimulus* (GO:0050896), followed by *multi-organism process* (GO:0051704), and downstream categories (Figure 4, Table S8). The main molecular functions regulated by iodine in the roots were related to *antioxidant* (GO:0016209) and *oxidoreductase activity* (GO:0016491) and related child terms, in particular *peroxidase activity* (GO:0004601) and *oxidoreductase activity, acting on peroxide as acceptor* (GO:0016684) (Figure S1, Table S9).

**FIGURE 4.**
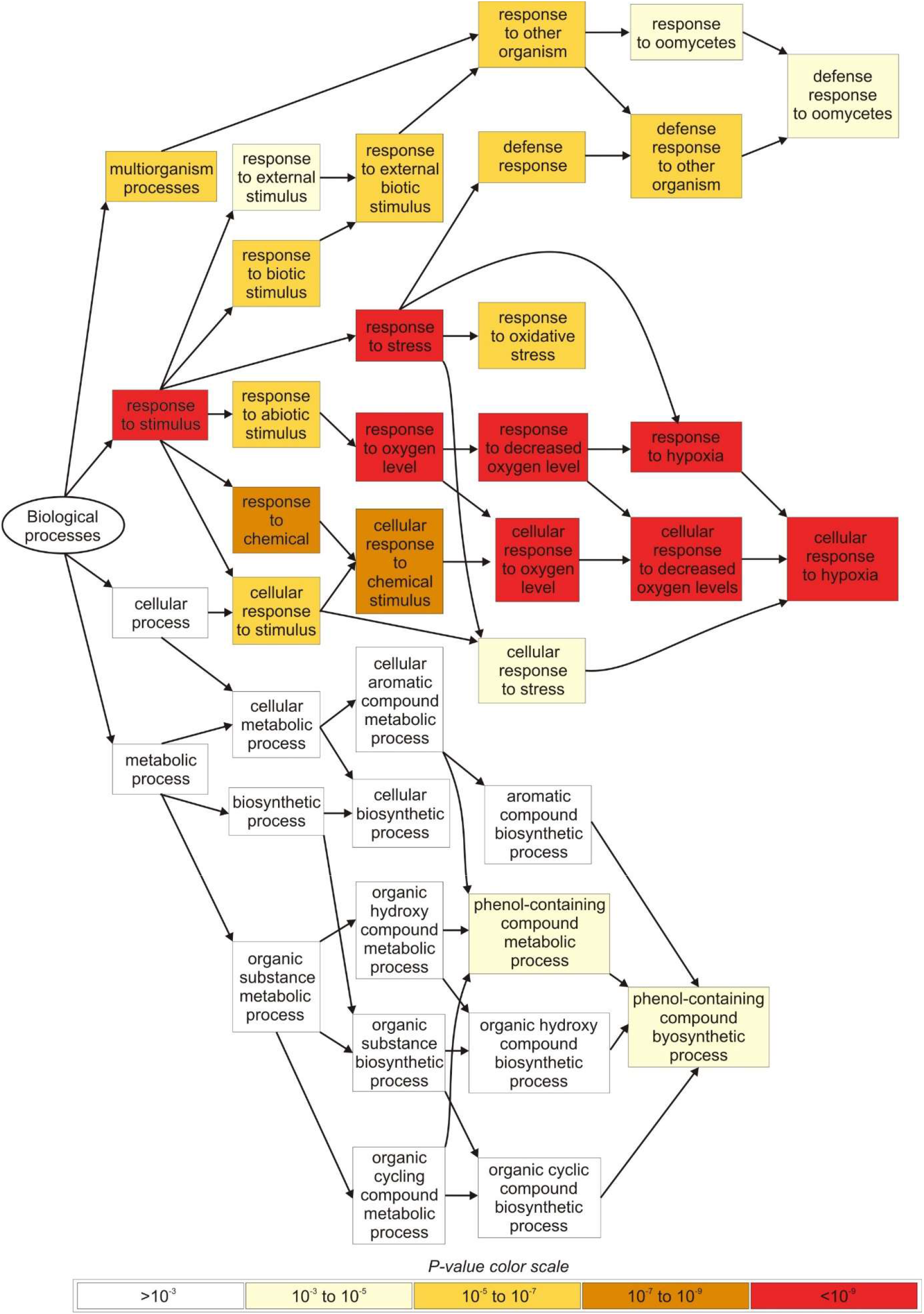
Overview of the main biological processes affected by iodine based on the GO terms enrichment analysis carried out in root tissues. Only genes regulated in NaI- and KI-treated plants, and not in KBr-treated plants, when compared with the control, were analyzed. The figure was extracted from GOrilla (http://cbl-gorilla.cs.technion.ac.il) and reproduced. In this analysis, DEGs with log_2_FC≥2.5 or log_2_FC≤−2.5 were used.

DEGs analysis performed with MapMan highlighted an over-representation of several genes in root samples that were related to calcium regulation and protein modification/degradation (Figure S2a), together with genes encoding for the large enzyme families including peroxidases, oxidases, glutathione S-transferases, and cytochrome P450 (Figure S2b).

The relatively low number of genes regulated by iodine in the shoots prevented a gene ontology analysis from being performed. However, in terms of the most well characterized genes specifically regulated by iodine treatments in the shoot, the main pathways affected were directly or indirectly involved in biotic (approximately 48% or 40% of up- or down-regulates genes, respectively) or abiotic (approximately 45% or 33% of up- or down-regulates genes, respectively) stress response pathways (Tables S4 and S5). Several genes playing a role in the transition to flowering (At4g19191 and At1g75750) and embryo and pollen development (i.e. At1g21310, At3g54150) are also worth mentioning.

The involvement of iodine in the defence response, highlighted by the previous analyses performed on root samples, was also suggested by querying all publicly available microarray datasets (https://genevestigator.com) using the list of iodine-responsive genes of both shoot (Figure S3) and root (Figure S4) tissues. The majority of the up- or down-regulated genes were commonly modulated by the presence of fungal infection, salicylic acid (SA) or synthetic analogues of SA, such as benzothiadiazole (Kouzai et al, 2018).

### Protein iodination in plants

Iodine can be found in plant tissues not only in a mineral form but also in organic compounds (Wang, Zhou, Fredimoses, Liao & Liu, 2014; Smoleń et al., 2020). To verify the possible *in vivo* incorporation of iodine into proteins, we carried out two different experiments by feeding hydroponically grown plants with ^125^I and carrying out the autoradiography of the SDS-PAGE of the relative protein extracts to detect possible radio-labelled proteins. The experiments were performed first with Arabidopsis plants, and then with other species, namely maize, tomato, wheat and lettuce.

The experiment carried out with Arabidopsis plants revealed the presence of at least six radio-labelled bands at different molecular mass values in the protein extracts from shoot tissues (Figure 5a-Exp.1) and eleven radio-labelled bands from root (Figure 5b-Exp.1) tissues, indicating the presence of proteins likely containing iodo-amino acids. Iodinated proteins were preferentially present in root tissues, as the abundance and intensity of ^125^I-labelled bands were higher in the root than in the shoot extracts. No radioactive signals were observed in the shoot and root control samples.

**FIGURE 5.**
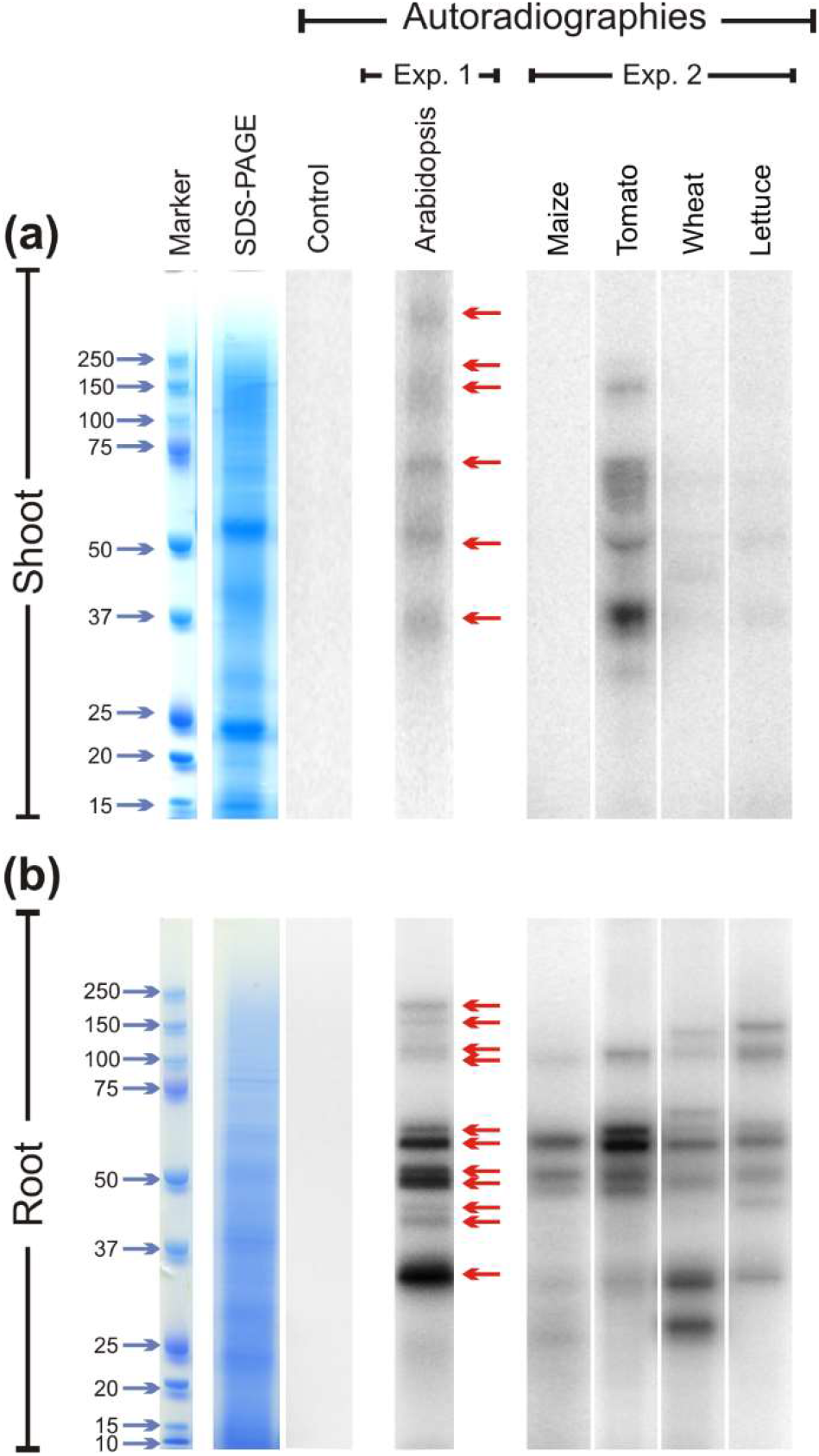
Autoradiographies of the SDS-PAGE gels. Comparison between the position and relative intensities of ^125^I radiolabelled bands of representative shoot (a) and root (b) protein extracts from ^125^I treated Arabidopsis (Exp.1), and maize, tomato, wheat and lettuce plants (Exp.2). Sampling was performed after 48-h of ^125^I incubation. Autoradiographies in both Exp.1 and Exp.2 were acquired after 72-h of gel exposition to the multipurpose phosphor storage screen. Representative pictures of total stained protein extracts (SDS-PAGE) and of the autoradiographies of control samples after 15 days of exposition are also shown (a,b). Controls consisted in protein extracts obtained from plants untreated with ^125^I during their growth, to which the radioactive solution containing ^125^I was added during the extraction process.

Several ^125^I-labelled bands were also observed in the leaf extracts of tomato, wheat and lettuce samples, whereas no ^125^I-containing bands were visible in the leaf protein extracts of maize (Figure 5a-Exp.2). A clear radioactive signal was detected in several root proteins extracted from all the species analyzed, including maize (Figure 5b-Exp.2). Also in this case, the intensity of the radiolabeled bands was higher in root than in shoot extracts. A good degree of conservation of the molecular mass values of the putatively iodinated proteins was observed among the five plant species analyzed (Figure 5).

### Identification of iodinated proteins in *Arabidopsis thaliana*

The identification of the radiolabeled proteins described above was hampered by the presence of a radioactive isotope, which meant that our samples did not meet the safety rules for proteomic facilities. To maximize the probability of success in identifying targets of protein iodination, we then focused on the nano-LC-ESI-MS/MS raw data already acquired within the framework of experimental studies on different organs/subcellular districts of *Arabidopsis thaliana*, and released in the public repository PRoteomics IDEntification Database (PRIDE) Archive (Perez-Riverol et al., 2019).

The datasets considered for our analysis refer to many different experimental conditions in terms of plant growth, treatment, and cultivation regimen, as well as the sample processing and fractionation performed before proteomic analysis. No experiments were explicitly related to iodination studies, the presence of iodine occurred accidentally, as a consequence of its natural presence in the cultivation environment (i.e. soil, air, irrigation water), or because it was conventionally present in the MS growing medium (Murashige & Skoog, 1962), which is widely used in studies based on *in vitro* plant tissue culture.

Following the reactivity order of amino acids described in the Introduction section, mono-iodination at Tyr and His residues were considered in the searching parameters as variable modifications. The output of the database search, in terms of proteins iodinated at Tyr or His residues has been reported in Table S10. The iodinated peptides were identified in 16 out of the 21 datasets analyzed. Iodinated sequences differently modified for deamidation, and/or protein N-terminal acetylation, and/or Met oxidation are reported as a unique iodinated peptide inventory in the Table S11. A total of 106 iodinated peptides, corresponding to 82 protein accessions in the TAIR10 database, were thus identified in the datasets of *A. thaliana* (Table S11).

Most of the modified peptides were found to be iodinated at Tyr residues, while His iodination was identified in only five peptides. Representative MS/MS spectra of Tyr-iodinated peptides are reported in Figure 6a.

**FIGURE 6.**
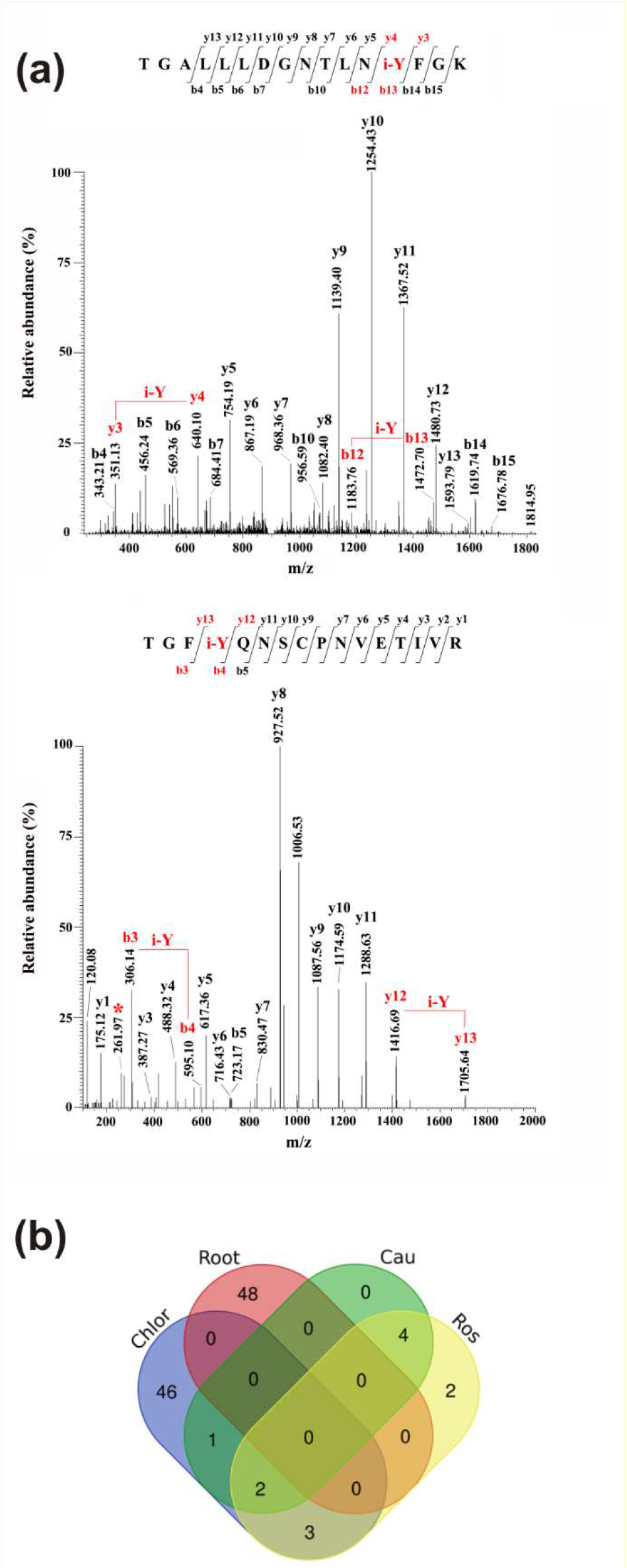
Iodination in *A. thaliana* proteins identified by database searching of nano-LC-ESI-MS/MS raw data from a public repository (PRIDE). (a) Unambiguous assignment of iodination sites by MS/MS analysis in two peptides from chloroplast (light harvesting complex of photosystem II 5, AT4G10340.1), upper panel, and roots (peroxidase superfamily protein, AT4G30170.1), lower panel. The peptides are identified by both y, and b ions. Red labels in the spectra evidence the mass shift corresponding to the iodinated tyrosine (i-Y). (b) Venn Diagram showing the iodinated peptide sequences identified in the datasets of chloroplasts (Chlor), cauline (Cau), rosette (Ros), and roots (Root).

To evaluate the entire output of iodinated peptides identified, iodinated sequences for chloroplasts, caulines, rosettes, and roots were processed and visualized in a Venn diagram (Figure 6b). This showed the presence of the common iodinated peptides for the cauline, rosette, and chloroplast subsets. The root subset was clearly distinct from the other three subsets that were all from the green parts of the plant.

### Iodinated proteins in *A. thaliana* leaves

Iodinated peptides identified in 11 datasets of cauline, rosette, and leaf-isolated chloroplasts were assigned to 42 proteins. Most of the modified species were in the dataset of chloroplastic proteins (Figure 6b). STRING interaction analysis of the modified proteins revealed a single network of 40 iodinated proteins (PPI enrichment p-value:< 1.0e^−16^) (Figure S5a; Table S12). A total of 31 of the 40 proteins in this network were involved in photosynthesis (GO:0015979; false discovery rate 7.82e^−53^) (Table S13), as also attested by their functional analysis, according to MapMan categories (Figure S5b, Table S14). Most of the proteins were constituents of photosystem II (PSII), i.e. proteins of the reaction center (PsbA, PsbB, PsbC and PsbD) and oxygen evolving center (PsbO, PsbP, PsbQ and PsbR), components of the light harvesting complex II (CAB3, LHCB2.1, LHCB1B1, LHCB3, LHCB5), factors involved in the assembly/maintenance of the photosystem (Psb27, Psb29, Psb31 and Psb33), proteins involved in PSII protection from photodamage (MPH1) or in the degradation of the photodamaged D1 reaction center (FtsH plastidial protease and Var1).

Other iodinated proteins were components of the photosystem I (PSI) complex (PsaB, PsaE, PsaF and PsaH), components of the cytochrome b6/f (Cyt b6/f) complex (PetA and PetC), plastocyanin electron carrier (PETE2), ferredoxin-plastoquinone reductase involved in cyclic electron flow around PSI (FNR1), and the component PGRL1-like of cyclic electron flow PGR5/PGRL1 complex (PGR2-like A).

Iodination was also assigned to three subunits of the CF1 subcomplex of ATP synthase (ATPA, ATPB, ATPC1), and several proteins involved in the Calvin Cycle, including some with ribulose-1,5-bisphosphate carboxylase/oxygenase (RuBisCo) (RBCL, ORF110A, RBCS1A, RCA) or fructose-1,6-bisphosphate aldolase (FBA2) activities. Three additional iodination targets were not related to photosynthesis (At2g24940, At1g74470.1 and At3g09260.1).

Some iodinated proteins (RBCL, RBCS1A, ORF110A, PsbB, PsbC, LHB1B1 and DRT112) were identified in at least two datasets apart from that of the chloroplastic species isolated by BN-PAGE. These proteins were thus considered as preferential targets of protein iodination in Arabidopsis leaves. RuBisCo large and small chains (RBCL, RBCS1A) and PSII components PsbB and PsbC were among the most abundant proteins in leaves according to the data reported in the Protein Abundance database (PAXdb).

### Iodinated proteins in *A. thaliana* roots

Iodinated peptides identified in 5 datasets of roots were assigned to 40 proteins. The STRING interaction analysis recognized three networks containing 24 of the 40 iodinated proteins (Figure S6a, Table S12), The GO analysis for these proteins showed a significant over-representation of biological processes related to the *response to stress* (GO:0006950; 3.24e^−07^), *response to oxidative stress* (GO:0006979; 7.87e^−07^), *response to toxic substances* (GO:0009636; 3.30e^−06^), and *response to stimulus* (GO:0050896; 4.53e^−06^). For the molecular functions, the most enriched categories were *copper ion binding* (GO: 0005507; 5.13e^−07^) and *peroxidase activity* (GO:0004601; 2.75e^−06^) (Table S15).

The functional analysis of the iodinated proteins in roots, according to MapMan categories, highlighted a broad range of biological roles (Figure S6b, Table S14). In particular, among the iodinated proteins identified, only 12 were found in two or more datasets. Five proteins belong to the classical plant (class III) peroxidase subfamily (At4g30170, At1g05240, At2g37130, At3g01190, At5g17820). The alignment of the protein sequences of the peroxidases mentioned above showed that iodinated residues in all peptides preferentially corresponded to conserved Tyr residues, while only two iodinated tyrosines were unrelated (Figure S7).

The other iodinated proteins included: i) copper amine oxidase (CUAOy2), a cell-wall oxidase showing primary amine oxidase activity; ii) beta-galactosidase 5 (BGAL5), a glycoside hydrolase involved in the modification of cell wall polysaccharides; iii) glycosyl hydrolases family 32 protein (ATBFRUCT1) acting as a cell wall invertase; iv) Pole Ole1 allergen/extension domain (IPR006041)-containing proline-rich protein-like 1 (PRPL1-MOP10) and root hair specific 13 protein (RHS13), which are cell-wall components; v) D-mannose binding lectin protein (MBL1); and vi) glyceraldehyde-3-phosphate dehydrogenase C sub 1 (GAPC1), a key enzyme in glycolysis. According to PAXdb, some of the above-reported proteins are abundant in the roots.

## DISCUSSION

### Iodine influences plant growth and development and can modulate the plant transcriptome

Establishing whether iodine is important for a plant’s life is complex, as it is always present in variable amounts in the soil, water, and atmosphere. Plants can take up iodine from the soil solution through the root system, but they also assimilate it from the air or absorb it through the leaves if dissolved in salt solutions or in rain. All these processes occur naturally (Fuge & Johnson, 1986; Ashworth, 2009), thus a plant cannot be grown in the complete absence of iodine.

To identify whether iodine can act as a micro-nutrient, we supplied it to plants at very low concentrations (in the micromolar range). These concentrations are typical of many mineral elements that are beneficial or essential when taken up in low doses, and phytotoxic when in excess (Welch & Shuman, 1995). We observed a difference in plant growth between the control and iodine-treated plants. Where these could be perceived as positive effects of addition of a beneficial compound, these can also be interpreted as a negative effect of removal of iodine from the plant’s nutrition. An increase in biomass and seed production, together with a very evident hastening of flowering, was observed by feeding plants with KIO_3_ (Figure 1, Table S1) or KI (Figure 2) at 0.2 or 10 mM. On addition of iodine, flowering was always early and appeared to be specific for iodine, since it was present in the KIO_3_, KI and NaI treatments but completely absent in KBr (Figures 1 and 2). However, the positive effects of iodine on growth were lost at 30 μM. This suggests that a concentration of 30 μM – applied as I^−^ may be above the toxicity threshold.

In the range of 1-10 μM iodine increases biomass in vegetables, e.g. spinach (Zhu, Huang, Hu & Liu, 2003), lettuce (Blasco, Leyva, Romero & Ruiz, 2013), tomato (Lehr, Wybenga & Rosanow M, 1958; Borst Pauwels, 1961), and strawberry (Li et al., 2016), or staple crops, e.g. barley (Borst Pauwels, 1961) and wheat (Cakmak et al., 2017). In tomato, Lehr et al. (1958) demonstrated that treatments with iodine accelerated plant growth, causing early flowering associated with an increase in yield. Similarly, Umaly & Poel (1970) found that the addition of 4 μM KI to the nutrient solution stimulated tomato plants to produce flowers two to three days earlier than the control, whereas the use of iodine at a higher concentration (80 μM KI) delayed flower formation and reduced the number of inflorescences. It must be noted that in most biofortification studies on iodine, its native occurrence in nutrient solution or soil of the control plants is not always reported, and where iodine concentrations in leaf or root tissue in control plants are reported, these always indicate that iodine was available for uptake in the control plants (e.g. Borst Pauwels, 1961, Cakmak et al., 2017).

Flowering is a complex physiological process affected by a multitude of internal and external factors, and its hastening may represent an evolutionary adaptive mechanism to guarantee species survival, by optimizing the seed set in the case of biotic or abiotic stresses (Ionescu, Møller & Sánchez-Pérez, 2017). In Arabidopsis, heat and drought stress are correlated with early flowering, which in turn is generally associated with a reduction in plant growth (Balasubramanian, Sureshkumar, Lempe & Weigel, 2006; Schmalenbach, Zhang, Reymond & Jiménez-Gómez, 2014). In our study, 10 μM iodate or iodide promoted flowering without negatively impacting on biomass production (Figures 1 and 2), which was increased by KIO_3_, thus suggesting the specific flowering-promoting role of iodine in the process. Our transcriptomic analysis of plants treated with 10 μM KI for 48 h showed that several genes were specifically regulated (Figure 3). This was more evident in root than in shoot tissues, probably because iodine was added to the nutrient solution and was used first by roots before the green parts. Interestingly, transcripts regulated by iodine in the roots were mostly involved in the plant response to biotic and/or abiotic stresses (Figure 4; Tables S6,7 and S8) and the selective regulation of iodine on these groups of genes was also observed in the shoots (Tables S4 and S5, Figure S3).

Although no previous data are available on the response of Arabidopsis to iodine at the transcriptomic level, the induction of *HALIDE ION METHYLTRANSFERASE, SALICYLIC ACID CARBOXYL METHYLTRANSFERASE* and *SALICYLIC ACID 3-HYDROXYLASE* genes by aromatic iodine compounds, indicating a possible involvement of iodine in the SA metabolism, has already been described in tomato plants (Halka et al., 2019). SA is a signalling molecule involved in local defence reactions at infection sites and the induction of systemic resistance (Vlot, Dempsey & Klessig, 2009).

Iodine likely has an indirect effect on plant resistance given that iodine treatments induce the biosynthesis of several enzymatic or non-enzymatic compounds involved in the plant response to environmental stresses (Leyva et al., 2011; Blasco et al., 2013; Gupta, Bajpai, Majumdar & Mishra, 2015). The antioxidant response induced by increasing KI and KIO_3_ levels was found to be strongly associated with the synthesis of phenolic compounds (Incrocci et al., 2019; Kiferle et al., 2019), in agreement with our transcriptomic data (Figure 4, Table S8). Iodine treatments have also been associated with the modulation of the essential oil composition in basil plants (Kiferle et al., 2019), which plays a key role in defensive and attraction mechanisms in response to the environment (Lee & Ding, 2016).

In our study, the relationship between iodine and plant resistance to stress was also suggested by the activation of GO terms associated with hypoxia (Figure 4; Table S8) and the over-representation of data points related to calcium (Ca) regulation (Figure S2a). Ca plays a central role in the plant perception of stress by activating a general defence mechanism, which relies on a Ca spiking mechanism and thus on the battery of Ca-dependent proteins that sense Ca and transduce the signal to downstream targets (Dodd, Kudla & Sanders, 2010; Pucciariello, Parlanti, Banti, Novi & Perata, 2012).

Our microarray results did not provide evidence that iodine regulated the expression of anaerobic core genes, such as *PYRUVATE DECARBOXYLASE 1* or *ALCOHOL DEHYDROGENASE* (Mustroph et al., 2010). This suggests that the regulation of hypoxic genes by iodine was not associated with limiting O_2_ levels but with an unspecific plant response to an environmental/biotic stress, as is also highlighted by the activation of enzyme families associated with antioxidant response and xenobiotic detoxification, such as peroxidases, cytochrome P450 or glutathione S-transferases (Figure S2b) (Mittler, Vanderauwera, Gollery & Van Breusegem, 2004; Jouili, Bouazizi & El Ferjani, 2011; Pandian, Sathishraj, Djanaguiraman, Prasad & Jugulam, 2020).

At very low doses, iodine thus activates a general defence response which takes place before any biotic or abiotic danger and thus may prepare the plant for a possible future attack or environmentally unfavourable conditions.

The impact of iodine on the transcriptome related to defence was impressive. For evaluation of iodine as a plant nutrient the possible influence on growth and development is also important, and this appeared to be sustained by more scattered elements, requiring further studies. For instance, we found iodine-driven modulation in the shoots of the expression of specific gene that are known to regulate flowering (Xing et al., 2013; Trapalis, Li & Parish, 2017). This is line with our phenotypical data that indicate that iodine promotes early blooming (Figures 1b and 2a).

### Iodine is incorporated into plant proteins

Using radiolabeled iodine, we observed iodine incorporation into leaf and root proteins in various plant species, with a good level of conservation of the molecular mass values (Figure 5), suggesting that iodination plays a functional role in specific proteins. The absence of radioactive bands in maize leaf extracts may be due to a specific characteristic of the species: maize was the only C4 plant analyzed in our study.

Under alkaline conditions, iodine is known to react *in vitro* with free amino acids, such as Tyr and His, possibly leading to the formation of several I-labelled proteins (Scott, 1954). In our experiments, however, no radioactive signals were present when ^125^ iodine was added to shoot and root control samples after protein extraction, indicating that protein iodination occurred *in vivo*.

To the best of our knowledge, the presence of naturally occurring iodinated proteins in higher plants has never been described before, although it is well known in seaweed. For instance, the fraction of iodine bound to proteins in *Sargassum kjellanianum* accounts for 65.5% of the total element content of this organism (Hou et al., 2000). In addition, 1D and 2D gel-based electrophoresis combined with laser ablation ICP-MS highlighted several iodinated proteins in Nori seaweed, although no further analyses were conducted to identify them (Romarís-Hortas et al., 2014).

In this study, we identified several iodinated peptides from several high-quality proteomic datasets on plants grown without intentional enrichment of the growing media with iodine, but accumulating the iodine naturally present in soil and water or in MS media. Interestingly, these iodinated peptides belong to proteins, which appeared to be involved in well-defined biological contexts within the plant (Tables S10 and S11).

In roots, some of the iodinated protein networks that have been identified belong to various class III peroxidases (EC 1.11.17). These are a large molecular family of isozymes present in higher plants, which catalyze redox processes between H_2_O_2_ and reductants and are involved in growth, cell wall differentiation and the response to various biotic/abiotic stresses (Moerschbacher, 1992; Hiraga, Sasaki, Ito, Ohashi & Matsui, 2001). They are abundant in root tissues, in which high concentrations of H_2_O_2_ occur during root development (Dunand, Crèvecoeur & Penel, 2007). In the presence of low concentrations of iodide, various peroxidases catalytically degrade H_2_O_2_ at neutral pH values, generating hypoiodous acid (HIO) (Davies, Hawkins, Pattison & Rees, 2008; Vlasova, 2018), a strong iodinating agent that can modify proteins in the environment surrounding the site of its generation, also inducing modification at Tyr. Our findings thus suggest that the class III peroxidases involved in the neutralization of H_2_O_2_ present in root tissues, in the presence of iodide and resulting HIO, become themselves the targets of iodination, which in our study occurred at different Tyr residues in their structure. In fact we also found that Tyr iodination occurred in other proteins that directly/indirectly interact with or are functionally linked to class III peroxidases, and/or are present at high levels in root tissues (Figure S6a; Table S12).

Regarding leaves, we identified a number of iodinated peptides from proteins in proteomic datasets from Arabidopsis cauline, rosette, and chloroplast extracts (Figure 6b). Most of these proteins are well-known constitutive subunits of molecular complexes (PSII, PSI, Cytb6f and ATPase) present in the plant photosynthetic machinery (Nevo, Charuvi, Tsabari & Reich, 2012; Dekker & Boekema, 2005). In *in vitro* labeling experiments Machold & Aurich (1981) demonstrated that some of them can be iodinated.

The above-mentioned macromolecular assemblies are involved in the generation and transfer of reactive electrons, from their early formation up to the coupling reactions, where their chemical potential allows the generation of plant ATP, NADP and carbohydrates (Foyer, 2018). In this context, high light intensity is one of the major stress factors in green plant tissues, which leads to the production of highly reactive oxygen species (ROS) (^1^O_2_, O_2_^•-^, H_2_O_2_), and hydroxyl radicals (Schmitt et al., 2014; Foyer, 2018). These reactions also occur during normal photosynthetic conditions and, whenever not controlled by dedicated endogenous antioxidants, can impact on photosynthetic efficiency.

The principal target of light stress is the chloroplast, which is the preferential site of iodine accumulation in leaf (Weng et al., 2013), and PSI and PSII are the main sites of O_2_^•-^ and ^1^O_2_ production, respectively. High light illumination and the corresponding generation of ROS cause photoinhibition of PSII as well as the modification of other photosynthetic complexes, and induce an accelerated turnover of components of these molecular machineries (Li, Aro & Millar, 2018). The latter phenomenon is generally accomplished through the rapid degradation of photo-damaged proteins and concomitant substitution of them with newly synthetized functional copies (Aro et al., 2005; Yamamoto et al., 2014). This oxidative modification is also known to affect redox-regulated enzymes involved in the Calvin cycle. Proteomic studies have already characterized the nature and the sites of various oxidative and nitrosative modifications at Tyr, Trp and His residues in components of PSII, PSI, Cytb6f and ATP synthase complexes in thylakoid membranes from plants exposed to intense illumination (Galetskiy et al., 2011). These modifications were induced by ROS and/or other reactive nitrogen species, e.g. peroxinitrite, which are formed after the reaction of ROS with nitric oxide and other plant nitrogenous species (Bachi, Dalle-Donne & Scaloni, 2013; Lu & Yao, 2018).

Given that such oxidized and nitrated proteins coincide with those found iodinated in our study, ROS likely also react with iodo-containing ions present in the chloroplasts to generate iodinating species that affect Tyr and His residues. We found that iodination processes also affected other proteins functionally related to subunits of PSII, PSI, Cytb6f and ATP synthase complexes, such as those involved in the Calvin Cycle, which are present at high concentrations in the same subcellular district, and have already been reported to directly/indirectly interact with the above-mentioned photosynthetic assemblies (Figure S5a; Table S12).

The addition of ^1^O_2_ to I^−^ forms peroxyiodide, which decomposes into highly reactive iodine and iodo-containing radicals during dye-mediated photodynamic bacterial inactivation in the presence of potassium iodide (Wen et al., 2017). This process is fundamental for antimicrobial photodynamic therapy (Hamblin & Abrahamse, 2018). Thus, in the presence of iodide, the above-mentioned photoactivated reactions or their possible process variants may have contributed to generate highly reactive iodo-containing molecules and/or radicals, thereby leading to the formation of the iodinated proteins observed in this study. The possible functional meaning of these is certainly worth studying.

## CONCLUSIONS

We demonstrated that very low amounts of iodine (between 0.21 and 10 μM, i.e. in the range of the concentrations required by plants for several micro-nutrients) improved plant growth and development thus promoting both biomass production and early flowering, and that this effect could not be achieved by another halogen most resembling iodine (Br). Secondly, we found that iodine was able to modulate gene expression in a specific way, activating multiple pathways, mostly involved in defence responses. Finally, we demonstrated that iodine can be a structural component of several different proteins, and conserved iodinated proteins are synthesized in both the roots and shoots of phylogenetically distant species.

These three lines of evidence highlight that iodine has a nutritional role in plants. This means that the influence of iodine on plants is not merely the result of an indirect priming effect by a potentially phytotoxic compound. Considering that plant nutrients are chemical elements that are components of biological molecules and/or influence essential metabolic functions, iodine matches at least the first part of this definition. Further studies on the importance of organification of iodine in proteins on their catalytic and/or regulatory function will help to complete the picture on the functional role of iodine as a plant nutrient.

## Acknowledgements

This work was supported by Scuola Superiore Sant’Anna and by SQM International N.V..

## Accession numbers

Microarray datasets were deposited in Gene Expression Omnibus (GEO) public repository (accession no. GSE157643; https://www.ncbi.nlm.nih.gov/geo/query/acc.cgi?acc=GSE157643).

## AUTHOR CONTRIBUTIONS

PP, CK, MM, KH and HTH: conceived the project;

CK, MM and SB: experiments on plant phenotype and transcriptomic;

PS, CK and MM: experiments on radioactive feeding;

AMS: bioinformatics analysis of mass spectrometry-based proteomics data sets;

CK and AMS: data analysis and figures preparation;

CK, AMS, AS, SG: writing – original draft preparation;

PP, AS, SG, PS, KH and HTH: general discussion and revision of the article;

all other authors read and contributed to previous versions and approved the final version.

