## Supplementary Figures for "Evidences for a nutritional role of iodine in plants"

**SUPPLEMENTARY INFORMATION_FIGURES**

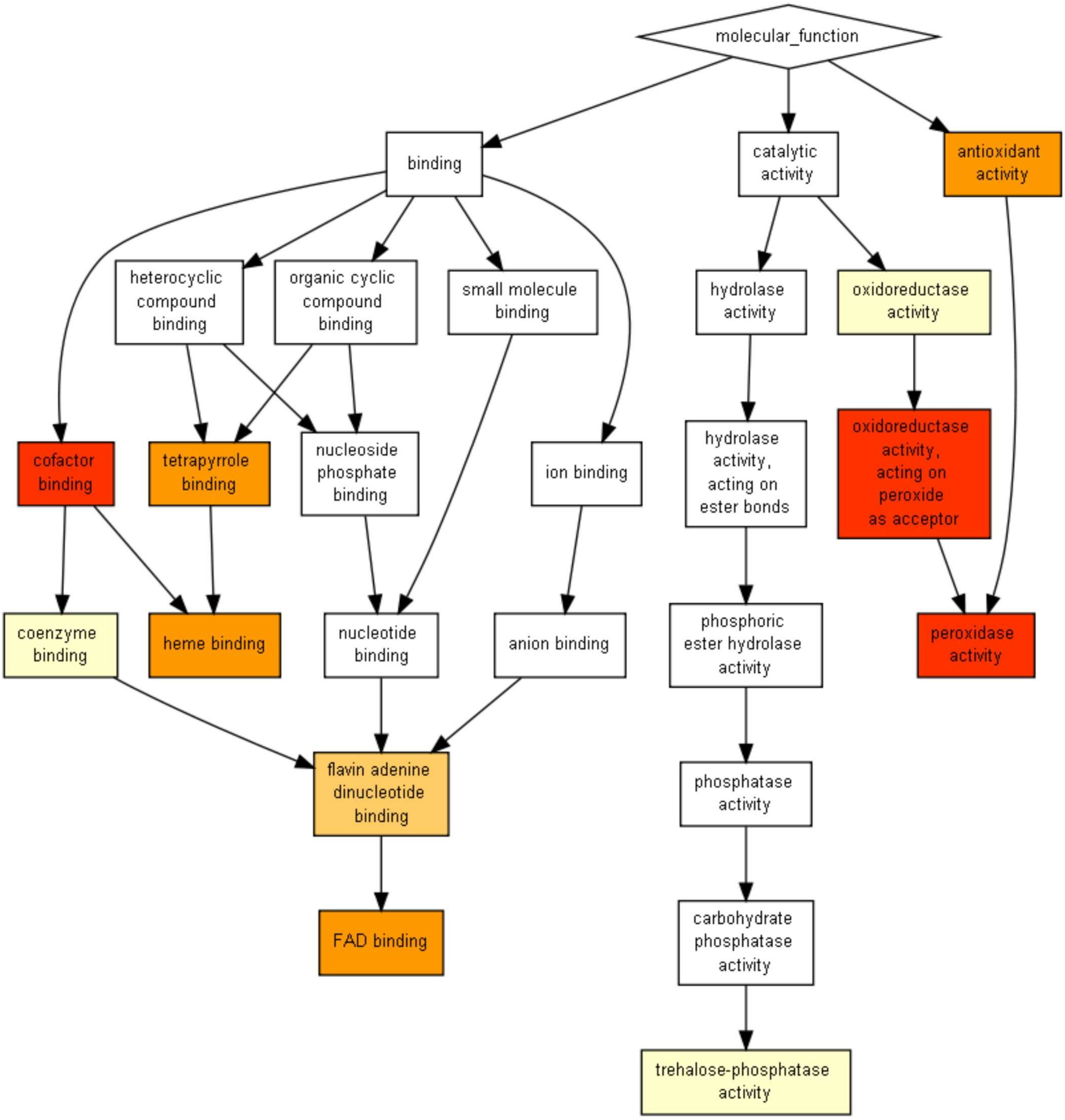


**Figure S1.** Overview of the main molecular functions affected by iodine based on the GO terms enrichment analysis in root tissues (only genes regulated in NaI- and KI-treated plants, and not in KBr-treated plants, when compared with the control were analyzed). The figure was extracted from GOrilla (http://cbl-gorilla.cs.technion.ac.il). In this analysis, DEGs with $\log_2FC \geq 2.5$ or $\log_2FC \leq -2.5$ were used.

(a)

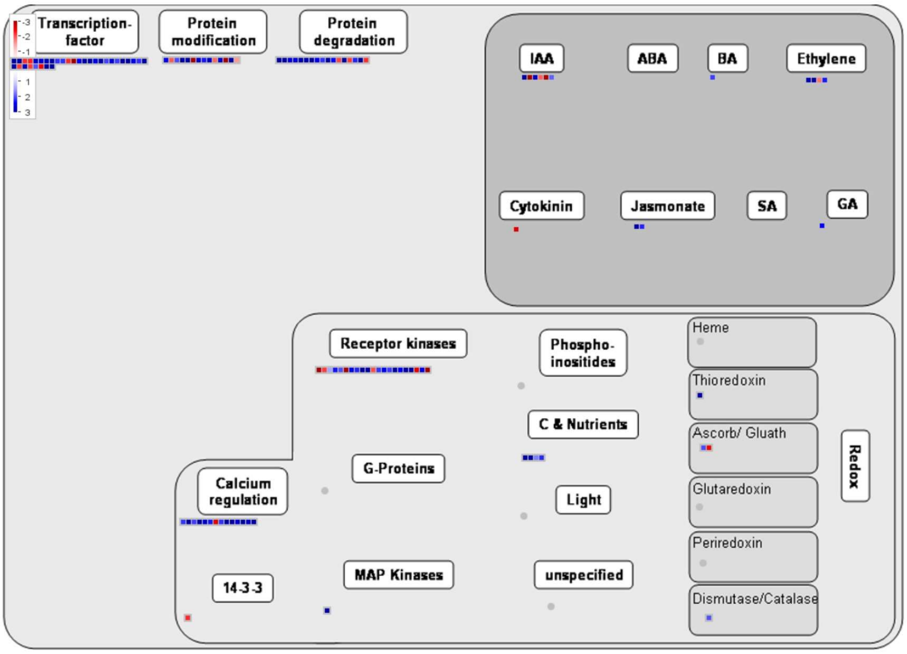


Regulation_overview
mapping: Ath_AGI_TAIR9
mapped:499 of 525 data points
visible: 124 data points

(b)

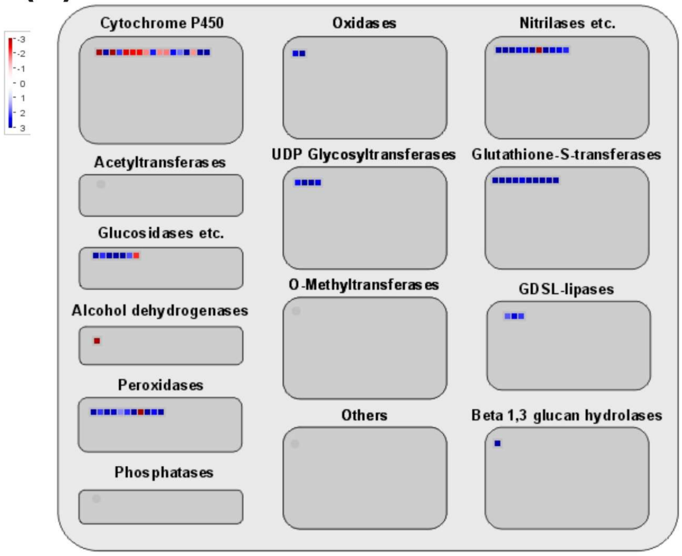


Large_enzyme_families_overview
mapping: Ath_AGI_TAIR9
mapped:499 of 525 data points
visible: 67 data points

**Figure S2.** Regulation overview mapping (a) and large enzyme families (b) overviews of the core list of genes regulated by iodine in root tissue (genes up-regulated only in NaI- and KI-treated plants, and not in KBr-treated plants, when compared with the control). Data and figures were extracted from MapMan (http://mapman.gabipd.org/mapman). In this analysis, DEGs with $log_2FC \geq 2$ or $log_2FC \leq -2$ were used.

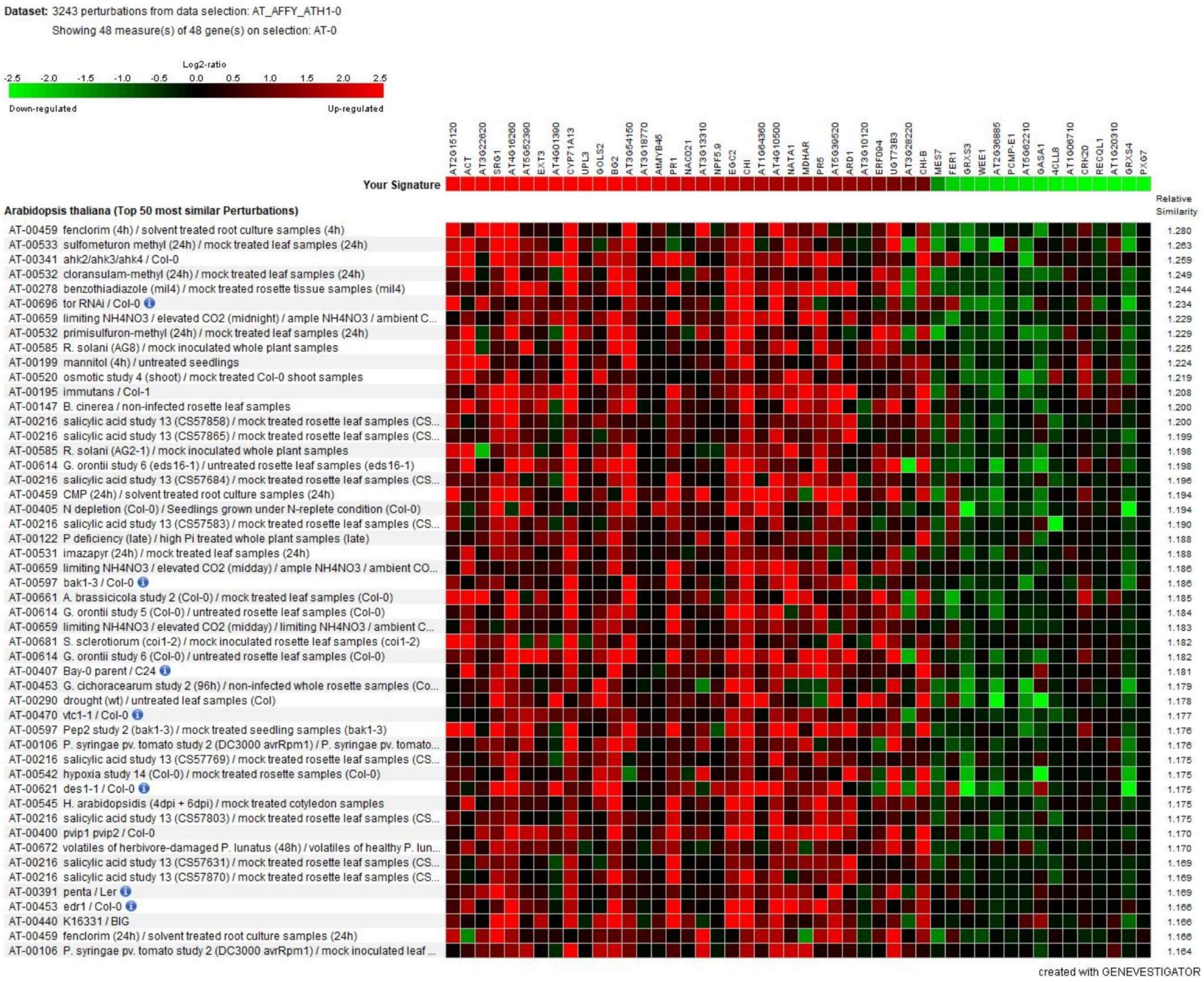


**Figure S3.** Co-regulation of iodine-responding genes in shoot tissues. Heatmap derived by the correlation between the expression level of the genes regulated by iodine with those up- or down-regulated by various other treatments, available on public datasets. Red = induced; green = repressed gene expression. Data and figure were extracted from Genevestigator (https://genevestigator.com). In this analysis, DEGs with log2FC≥2 or log2FC≤-2 were used.

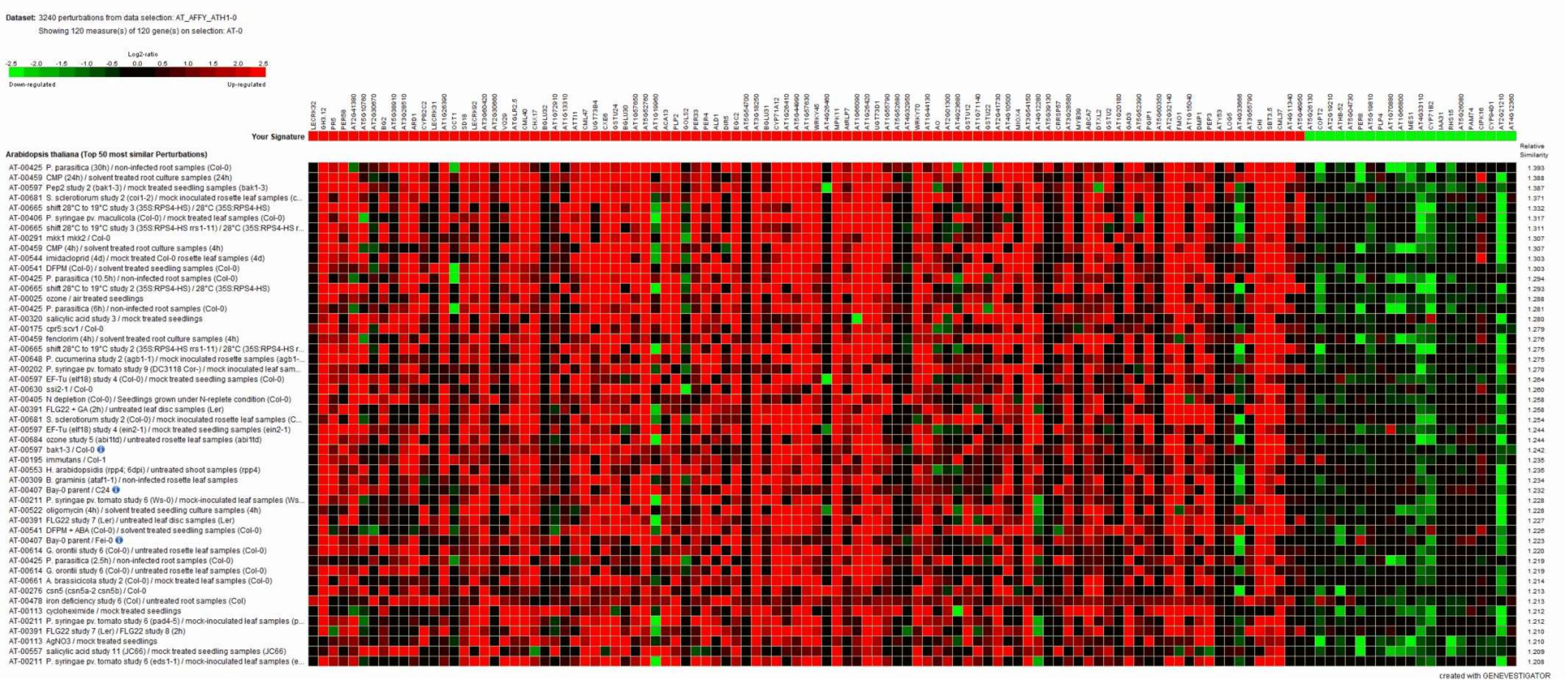

**Figure S4.** Co-regulation of iodine-responding genes in root tissues. Heatmap derived by the correlation between the expression level of the genes regulated by iodine, with those up- or down-regulated by various other treatments, available on public datasets. Red = induced; green = repressed gene expression. Data and figure were extracted from Genevestigator (https://genevestigator.com). In this analysis, DEGs with log2FC≥3.5 or log2FC≤-3.5 were used.

**(a)**

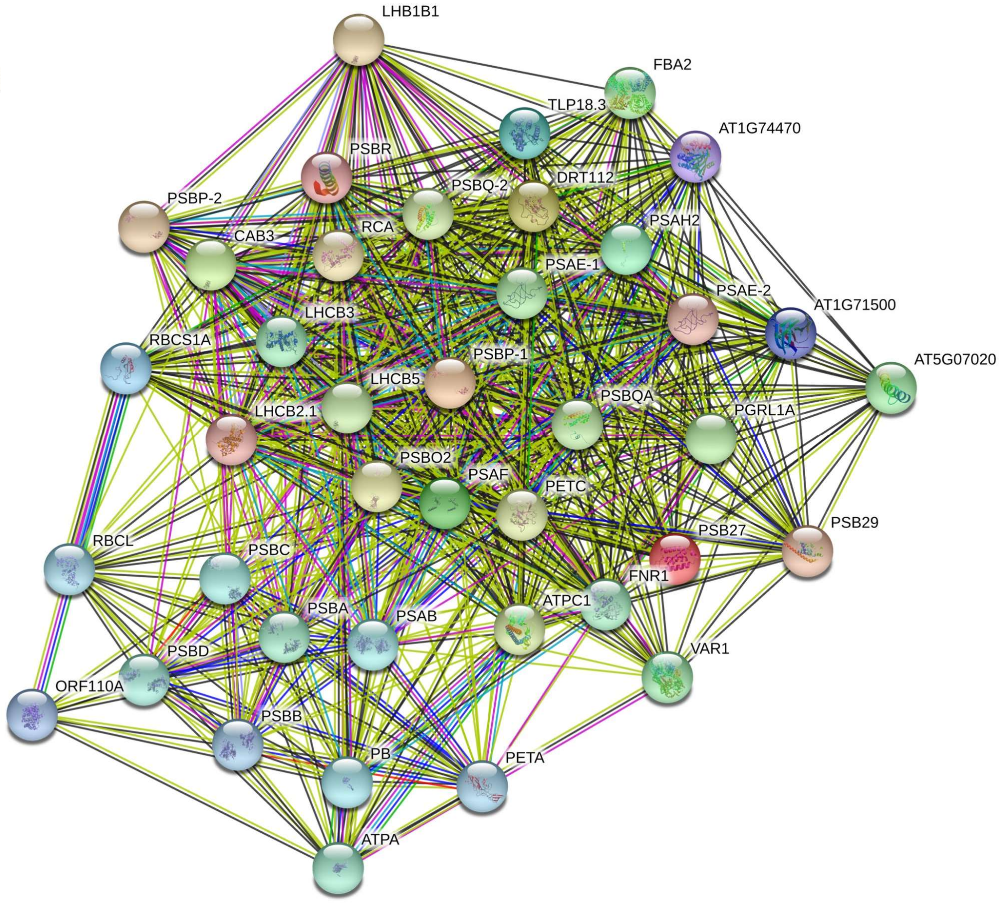


**(b)**

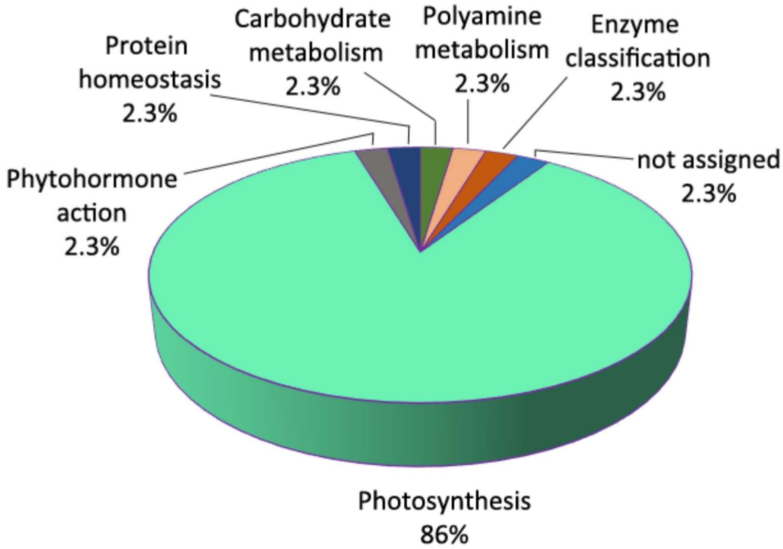


**Figure S5.** String analysis of the iodinated proteins identified in the *A. thaliana* datasets of leaves retrieved from PRIDE repository (a). Medium confidence interactions (score 0.4) are considered. Protein codes are reported in Table S12. Functional distribution of the iodinated proteins identified in the *A.thaliana* datasets of leaves retrieved from PRIDE repository (b). Identified proteins were mapped to one or more plant protein categories (BINs) using Mercator annotation tool (Table S14).

(a)

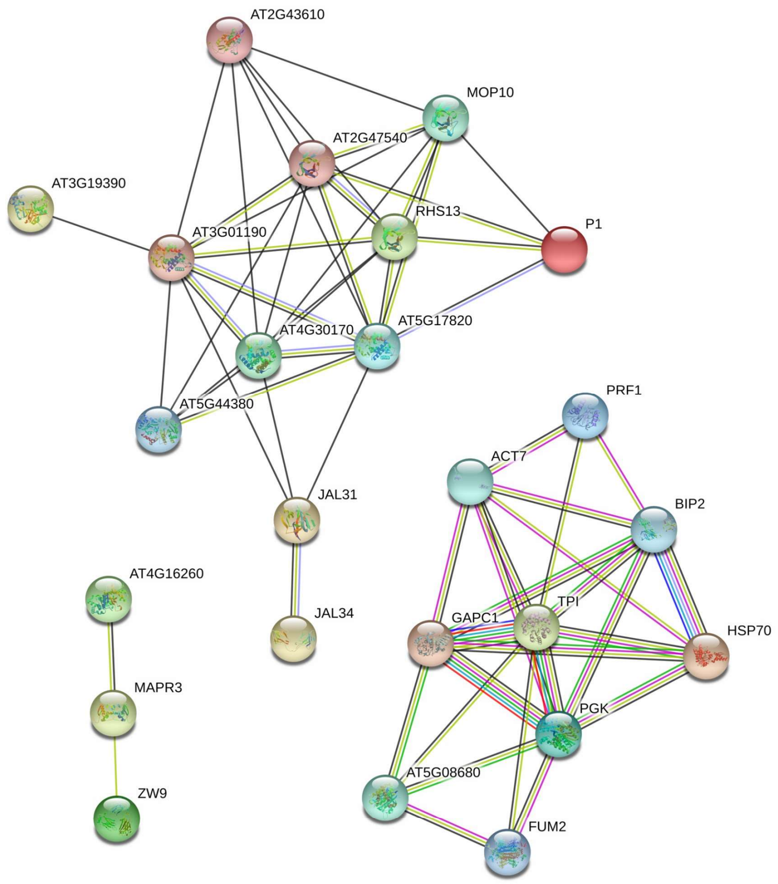


(b)

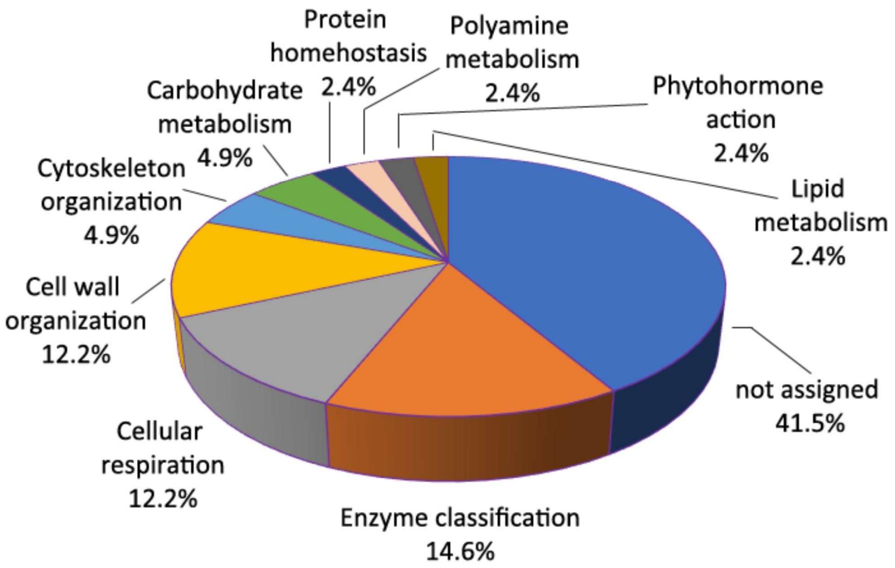


**Figure S6.** String analysis of the iodinated proteins identified in the *A. thaliana* datasets of roots retrieved from PRIDE repository. Medium confidence interactions (score 0.4) are considered. Protein codes are reported in Table S12. Functional distribution of the iodinated proteins identified in the *A.thaliana* datasets of roots retrieved from PRIDE repository (b). Identified proteins were mapped to one or more plant protein categories (BINs) using Mercator annotation tool (Table S14).

```
AT1G05240.1      ----MAIKNILALVVLLSVVGVSVAIPQLLDLDYYRSKCPKAEEIVRGVTVQYVSRQKTL
AT3G01190.1      ----MAASKRLVVSCLFLVLLFAQANSQGLKVGFYSKTCPQLEGIVKKVVFDAMNKAPTL
AT4G30170.1      ---MEKNTSQTIFSNFFLLLLLSSCVSAQLRTGFYQNSCPNVETIVRNAVRQKFQQTFVT
AT5G17820.1      ------MMKGAKFSSLLVLFFIFPIAFAQLRVGFYSQSCPQAETIVRNLVRQRFGVTPTV
AT2G37130.1      MANAKPFCLLGFFCLLLQLFSIFHIGNGELEMNYYKESCPKAEEIIRQQVETLYYKHGNT
                             .  :: :. .       *  .:* ..**: * *::  .

AT1G05240.1      AAKLLRMHFHDCFVRGCDGSVLLKSAKND-AERDAVPNLTLKGYEVVDAAKTALER--KC
AT3G01190.1      GAPLLRMFFHDCFVRGCDGSVLLDKPNNQ-GEKSAVPNLSLRGFGIIDDSKAALEK--VC
AT4G30170.1      APATLRLFFHDCFVRGCDASIMIASPSER-DHPDDMS-LAGDGFDTVVKAKQAVDSNPNC
AT5G17820.1      TAALLRMHFHDCFVKGCDASLLIDSTNS---EKTAGPNGSVREFDLIDRIKAQLEA--AC
AT2G37130.1      AVSWLRNLFHDCVVKSCDASLLLETARGVESEQKSKRSFGMRNFKYVKIIKDALEK--EC
                     **  ****.*:.**.*::: ..     .           :  :   *  ::    *

AT1G05240.1      PNLISCADVLALVARDAVAVIGGP-WWPVPLGRRDGRISKLNDALLNLPSPFADIKTLKK
AT3G01190.1      PGIVSCSDILALVARDAMVALEGP-SWEVETGRRDGRVSNIN--EVNLPSPFDNITKLIS
AT4G30170.1      RNKVSCADILALATREVVVLTGGP-SYPVELGRRDGRISTKASVQSQLPQPEFNLNQLNG
AT5G17820.1      PSTVSCADIVTLATRDSVALAGGP-SYSIPTGRRDGRVSNNLD--VTLPGPTISVSGAVS
AT2G37130.1      PSTVSCADIVALSARDGIVMLKGPKIEMIKTGRRDSRGSYLGDVETLIPNHNDSLSSVIS
                  . :**:*:::* :*: :.   **    :  ****.* *        :*    .:.

AT1G05240.1      NFANKGLNAKDLVVLSGGHTIGISSCALVNSRLYNFTGKGDSDPSMNPSYVRELKRKCPP
AT3G01190.1      DFRSKGLNEKDLVILSGGHTIGMGHCPLLTNRLYNFTGKGDSDPSLDSEYAAKLRKKCKP
AT4G30170.1      MFSRHGLSQTDMIALSGAHTIGFAHCGKMSKRIYNFSPTTRIDPSINRGYVVQLKQMCPI
AT5G17820.1      LFTNKGMNTFDAVALLGAHTVGQGNCGLFSDRITSFQGTGRPDPSMDPALVTSLRNTCRN
AT2G37130.1      TFNSIGIDVEATVALLGAHSVGRVHCVNLVHRLY-----PTIDPTLDPSYALYLKKRCPS
                  *   *:.    : * *.*::*   *  .  *:         **:::   .  *:. *

AT1G05240.1      -TDFRTSLNMDPG---SALTFDTHYFKVVAQKKGLFTSDSTLLDDIETKNYVQTQAILPP
AT3G01190.1      -TDTTTALEMDPG---SFKTFDLSYFTLVAKRRGLFQSDAALLDNSKTRAYVLQQIRT--
AT4G30170.1      GVDVRIAINMDPT---SPRTFDNAYFKNLQQGKGLFTSDQILFTDQRSRSTVNSFANS--
AT5G17820.1      ----SATAALDQS---SPLRFDNQFFKQIRKRRGVLQVDQRLASDPQTRGIVARYANN--
AT2G37130.1      PTPDPNAVLYSRNDRETPMVVDNMYYKNIMAHKGLLVIDDELATDPRTAPFVAKMAADN-
                       :   .     :   .*  ::. :   :*::  *  *  : .:   *

AT1G05240.1      VFSSFNKDFSDSMVKLGFVQILTGKNGEIRKRCAFPN
AT3G01190.1      HGSMFFNDFGVSMVKMGRTGVLTGKAGEIRKTCRSAN
AT4G30170.1      -EGAFRQAFITAITKLGRVGVLTGNAGEIRRDCSRVN
AT5G17820.1      -NAFFKRQFVRAMVKMGAVDVLTGRNGEIRRNCRRFN
AT2G37130.1      --NYFHEQFSRGVRLLSETNPLTGDQGEIRKDCRYVN
                     * . *  .:  :. .  ***  ****: *   *
```

**Figure S7.** Multiple sequence alignment of the class III peroxidases found iodinated in this study. Tyr residues within the sequences are colored in red and iodinated Tyr are highlighted in yellow. The alignment was performed using the Clustal W2 sequence alignment tool.
