## Supplementary Tables_ legends for "Evidences for a nutritional role of iodine in plants"

**SUPPLEMENTARY INFORMATIONS_TABLES**

**Table S1.** Primers used for RT-qPCR gene expression analysis.

**Table S2.** List of *A. thaliana* experimental datasets of nanoLC-ESI-MS/MS raw data retrieved from PRIDE Repository and subjected to database searching for iodinated peptides. The acronym has been indicated in the first column for the datasets in which iodinated peptides have been identified. Informative details on plant organ/subcellular compartment analysed, proteomic experiment performed, mass spectrometer used and related mass tolerances chosen for searching analysis, have been reported.

**Table S3.** Effect of different $KIO_3$ concentrations (0, 0.20 and 10 μM) in the nutrient solution on rosette and inflorescence FW and DW, seed production and number of produced siliques/plant. Each value is the mean (± standard error, SE) of 45 biological replicates, consisting of a single plant, with the exception of data on seed production: the cultivation system did not allow to harvest seeds from single plants, therefore the average of produced seed/tray (3 different trays;15 plants/tray) was compared and statistic analysis was performed accordingly. Values indicated by different superscript letters significantly differ from each other (according with one-way ANOVA, LSD posthoc test, $P \leq 0.05$).

**Table S4.** Up-regulated genes by iodine treatments in shoot tissues. The AGI code, name, and GO-TERM biological processes (BP), cellular component (CC) and molecular function (MF) of each gene are reported, when known.

**Table S5.** Down-regulated genes by iodine treatments in shoot tissues. The AGI code, name, and GO-TERM biological processes (BP), cellular component (CC) and molecular function (MF) of each gene are reported, when known.

**Table S6.** Up-regulated genes by iodine treatments in root tissues. The AGI code, name, and GO-TERM biological processes (BP), cellular component (CC) and molecular function (MF) of each gene are reported, when known.

**Table S7.** Down-regulated genes by iodine treatments in root tissues. The AGI code, name, and GO-TERM biological processes (BP), cellular component (CC) and molecular function (MF) of each gene are reported, when known.

**Table S8.** List of the main biological processes affected by iodine based on the GO terms enrichment analysis in root tissues (only genes regulated in NaI- and KI-treated plants, and not in KBr-treated plants, when compared with the control were analyzed). Data were extracted from Gorilla (http://cbl-gorilla.cs.technion.ac.il). In this analysis, DEGs with log2FC≥2 or log2FC≤-2 were used. 'P-value' is the enrichment p-value computed according to the mHG or HG model. This p-value is not corrected for multiple testing of 4037 GO terms. 'FDR q-value' is the correction of the above p-value for multiple testing using the Benjamini and Hochberg (1995) method (BH procedure). Namely, for the ith term (ranked according to p-value) the FDR q-value is (p-value * number of GO terms) / i. Enrichment (N, B, n, b) is defined as follows: N - is the total number of genes; B - is the total number of genes associated with a specific GO term; n - is the number of genes in the top of the user's input list or in the target set when appropriate; b - is the number of genes in the intersection. Enrichment = (b/n) / (B/N). The genes classified in each GO term are also listed.

**Table S9.** List of the molecular functions affected by iodine based on the GO terms enrichment analysis in root tissues (only genes regulated in NaI- and KI-treated plants, and not in KBr-treated plants, when compared with the control were analyzed). Data were extracted from Gorilla (http://cbl-gorilla.cs.technion.ac.il). In this analysis, DEGs with log2FC≥2 or log2FC≤-2 were used. 'P-value' is the enrichment p-value computed according to the mHG or HG model. This p-value is not corrected for multiple testing of 1737 GO terms. 'FDR q-value' is the correction of the above p-value for multiple testing using the Benjamini and Hochberg (1995) method (BH procedure). Namely, for the ith term (ranked according to p-value) the FDR q-value is (p-value * number of GO terms) / i. Enrichment (N, B, n, b) is defined as follows: N - is the total number of genes; B - is the total number of genes associated with a specific GO term; n - is the number of genes in the top of the user's input list or in the target set when appropriate; b - is the number of genes in the intersection. Enrichment = (b/n) / (B/N). The genes classified in each GO term are also listed.

**Table S10.** Identification details of the *A. thaliana* iodinated proteins identified in this study. The results obtained for the different experimental datasets from PRIDE repository have been reported in different sheets named after the acronyms reported in Table S2. Protein accession and description, sum posterior error probability (PEP) score, amino acid sequence coverage (%), number of identified peptides, peptide spectrum matches (PSMs), number of identified unique peptides, number of aminoacids in the protein (AAs), theoretical molecular mass of the protein (MW), theoretical isoelectric point of the protein (pI) and Mascot identification score have been reported.

**Table S11.** Iodinated peptides identified in *A.thaliana* datasets retrieved from PRIDE repository. Protein Accession and Description, sequence of iodinated peptide and iodination site have been reported. The datasets where the iodinated peptides were identified with Mascot Ion Score > 30 have been indicated.

**Table S12.** Bridged and non-linked nodes of the protein-protein STRING interaction analysis of the iodinated proteins identified in the *A.thaliana* leaf datasets retrieved from PRIDE repository. Results obtained from the analysis of leaves and roots were separately reported.

**Table S13.** List of the biological processes and molecular functions affected by protein iodination in leaves, based on the GO terms enrichment analysis performed through String platform (https://string-db.org). The list of 42 proteins iodinated in leaves was used as input and the enrichment analysis was performed against the whole genome as statistical background. The GO term ID, GO term description, the number of proteins from the input list classified in each GO term and the number of background proteins associated with a specific GO term are reported, with the false discovery rate (FDR) value.

**Table S14.** Iodinated proteins identified in *A.thaliana* datasets retrieved from PRIDE repository. Each protein identifier has been mapped to one or more MapMan "Bins" through Mercator analysis and corresponding names and descriptions have been reported in different sheets. Protein identifiers have been highlighted in pink if they were mapped to more than one Bincode. Protein identifiers were also mapped on the Arabidopsis Information Resource available at https://www.arabidopsis.org and gene model description and primary gene symbol have been also reported. The datasets in which the iodinated proteins were identified have been marked by an "X", in separated sheets for leaves and roots. Gene names for iodinated proteins have been reported in the first column of the table.

**Table S15.** List of the biological processes and molecular functions affected by protein iodination in root tissues, based on the GO terms enrichment analysis performed through String platform (https://string-db.org). The list of 40 proteins iodinated in roots was used as input and the enrichment analysis was performed against the whole genome as statistical background. The GO term ID, GO term description, the number of proteins from the input list classified in each GO term and the number of background proteins associated with a specific GO term are reported, with the false discovery rate (FDR) value.
