## Supplementary material for "Evidences for a nutritional role of iodine in plants": Table S1

| Gene (AGI) | Forward primer | Reverse primer |
|---|---|---|
| At4g05320 (UBQ10) | GGCCTTGTATAATCCCTGATGAATAAG | AAAGAGATAACAGGAACGGAAACATAGT |
| At2g25810.1 (TIP4) | TGATCTTCCCAATGGCTAAGG | TGATCTCCCAATGGCTAAGG |
| At5g52390 | GGTTAAATGCTTAAACCAATGTCC | CTTCTTGACTTCCCTTCTTGTGAT |
| At3g57260 | GCTTCCTTCTTCAACCACACAGC | TGGCAAGGTATCGCCTAGCATC |
| At1g56600 | AAGAAGCAACAGACACTTCAGCAG | TGAAGAGGCGTATGCAGCAAC |
| At1g75750 | TTCTCCAACTCGTCCAGGCTGA | TACACACGCACTCCCACAATCG |
| At4g15680 | GATAGAGCAAGCATTGGCTCAG | GAACCAGAGAGCGATTGAGATG |
| At2g36885 | GGAGCAACTGGTGGTGTTATATCC | CTGTAACAGCGTGCGGACTTTC |
| At5g13320 | GTTGTCACAAATTTCGCTGGCTTG | GCGCGTTGTTGTAGAAACCAGTC |
| At1g75040 | ATCACCCACAGCACAGAGACAC | AGCAATGCCGCTTGTGATGAAC |
| At4g23680 | ACCATGTCTTCCCTGATGCTATCG | TGAACACCTCCTCCTTTCCATCCC |
| At1g34510 | CCGAACGGTTACTGCAGCATTG | GGAAGCATCACAACCTTTGACAAC |
| At4g25220 | TACGTCAGCCACAACATGATCGG | CGACAAGTTTCCTGATGTCTCCTC |
| At1g13080 | TTGATGATCACTTGAAGCCAGAGG | CCCGCGAGAAATACATCCATGAC |
