## Supplementary material for "Evidences for a nutritional role of iodine in plants": Table S2

| Acronym | Pride accession | Organ /subcellular compartment | Experimet type | MS Instrument | ppm | Da |
|---|---|---|---|---|---|---|
| **ChlorBN** | **PXD003162** | choroplast | 1D-BN-PAGE | LTQ Orbitrap Elite | 10 | 0.8 |
| **Chlo_10545** | **PXD010545** | chloroplast | Label free | QExactive Orbitrap | 10 | 0.02 |
| **Chlo3516** | **PXD003516** | chloroplast | TMT labeling | QExactive Orbitrap | 10 | 0.02 |
| **PXD010730** | **PXD010730** | chloroplast | 1D-SDS-PAGE | QExactive Orbitrap | 10 | 0.02 |
| **TMAR** | **PXD000546** | thylakoid microdomains, Margins | 1D-SDS-PAGE | LTQ Orbitrap Velos Pro | 10 | 0.8 |
| CAU | **PXD012708** | Cauline -Multiorgan | MudPIT | Orbitrap Fusion | 10 | 0.02 |
| **CLLF** | **PXD013868** | Cauline leaf -Multiorgan | MudPIT | QExactive Orbitrap | 10 | 0.05 |
| **Ros** | **PXD012708** | Rosette - Multiorgan | MudPIT | Orbitrap Fusion | 10 | 0.02 |
| **LFD** | **PXD013868** | Leaf distal, Rosette - Multiorgan | MudPIT | QExactive Orbitrap | 10 | 0.05 |
| **LFP** | **PXD013868** | Leaf proximal, Rosette - Multiorgan | MudPIT | QExactive Orbitrap | 10 | 0.05 |
| **LFPT** | **PXD013868** | Leaf petiole, Rosette - Multiorgan | MudPIT | QExactive Orbitrap | 10 | 0.05 |

| | | | | | | |
|---|---|---|---|---|---|---|
| **Root** | **PXD012708** | Root - Multiorgan | MudPIT | Orbitrap Fusion | 10 | 0.02 |
| **RT** | **PXD013868** | Root - Multiorgan | MudPIT | QExactive Orbitrap | 10 | 0.05 |
| **RTTP** | **PXD13868** | Root tip - Multiorgan | MudPIT | QExactive Orbitrap | 10 | 0.05 |
| **RTUZ** | **PXD13868** | Root Upper Zone - Multiorgan | MudPIT | QExactive Orbitrap | 10 | 0.05 |
| RTTOF | **PXD0012710** | Root - Multiorgan | MudPIT | Triple TOF | 50 | 0.05 |
| - | **PXD000869** | chloroplast | 1DE_SDS-PAGE, ITRAQ | LTQ Orbitrap Velos | 10 | 0.5 |
| - | **PXD009725** | Rosette | Sthotgun | LTQ Orbitrap | 10 | 0.5 |
| - | **PXD004742** | Rosette | Sthotgun | LTQ Orbitrap Velos | 15 | 0.6 |
| - | **PXD004025** | Rosette | Sthotgun | LTQ Orbitrap Velos Pro | 10 | 0.8 |
| - | **PXD014292** | Mitochondria (cell culture) | glycine-SDS gel | QExactive Orbitrap | 10 | 0.05 |

| Acronym | Pride accession | Publication |
|---|---|---|
| ChlorBN | PXD003162 | Lundquist PK, Mantegazza O, Stefanski A, Stühler K, Weber AP. Surveying the Oligomeric State of Arabidopsis thaliana Chloroplasts. Mol Plant. 2016 Oct 26. pii: S1674-2052(16)30236-2, PubMed: 27794502 |
| Chlo_10545 | PXD010545 | Bouchnak I, Brugière S, Moyet L, Le Gall S, Salvi D, Kuntz M, Tardif M, Rolland N. Unravelling hidden components of the chloroplast envelope proteome: opportunities and limits of better MS sensitivity. Mol Cell Proteomics. 2019, PubMed: 30962257 |
| Chlo3516 | PXD003516 | Wang J, Yu Q, Xiong H, Wang J, Chen S, Yang Z, Dai S. Proteomic Insight into the Response of Arabidopsis Chloroplasts to Darkness. PLoS One. 2016 May 3;11(5):e0154235. eCollection 2016, PubMed: 27137770 |
| PXD010730 | PXD010730 | Wu GZ, Meyer EH, Richter AS, Schuster M, Ling Q, Schöttler MA, Walther D, Zoschke R, Grimm B, Jarvis RP, Bock R. Control of retrograde signalling by protein import and cytosolic folding stress. Nat Plants. 2019 5(5):525-538, PubMed: 31061535 |
| TMAR | PXD000546 | Tomizioli M, Lazar C, Brugière S, Burger T, Salvi D, Gatto L, Moyet L, Breckels LM, Hesse AM, Lilley KS, Seigneurin-Berny D, Finazzi G, Rolland N, Ferro M. Deciphering thylakoid sub-compartments using a mass spectrometry-based approach. Mol Cell Proteomics. 2014 Aug;13(8):2147-67, PubMed: 24872594 |
| CAU | PXD012708 | Zhang H, Liu P, Guo T, Zhao H, Bensaddek D, Aebersold R, Xiong L. Arabidopsis proteome and the mass spectral assay library. Sci Data. 2019 6(1):278, PubMed: 31757973 |
| CLLF | PXD013868 | Mergner J, Frejno M, List M, Papacek M, Chen X, Chaudhary A, Samaras P, Richter S, Shikata H, Messerer M, Lang D, Altmann S, Cyprys P, Zolg DP, Mathieson T, Bantscheff M, Hazarika RR, Schmidt T, Dawid C, Dunkel A, Hofmann T, Sprunck S, Falter-Braun P, Johannes F, Mayer KFX, Jürgens G, Wilhelm M, Baumbach J, Grill E, Schneitz K, Schwechheimer C, Kuster B. Mass-spectrometry-based draft of the Arabidopsis proteome. Nature. 2020 579(7799):409-414, PubMed: 32188942 |
| Ros | PXD012708 | Zhang H, Liu P, Guo T, Zhao H, Bensaddek D, Aebersold R, Xiong L. Arabidopsis proteome and the mass spectral assay library. Sci Data. 2019 6(1):278, PubMed: 31757973 |
| LFD | PXD013868 | Mergner J, Frejno M, List M, Papacek M, Chen X, Chaudhary A, Samaras P, Richter S, Shikata H, Messerer M, Lang D, Altmann S, Cyprys P, Zolg DP, Mathieson T, Bantscheff M, Hazarika RR, Schmidt T, Dawid C, Dunkel A, Hofmann T, Sprunck S, Falter-Braun P, Johannes F, Mayer KFX, Jürgens G, Wilhelm M, Baumbach J, Grill E, Schneitz K, Schwechheimer C, Kuster B. Mass-spectrometry-based draft of the Arabidopsis proteome. Nature. 2020 579(7799):409-414, PubMed: 32188942 |
| LFP | PXD013868 | Mergner J, Frejno M, List M, Papacek M, Chen X, Chaudhary A, Samaras P, Richter S, Shikata H, Messerer M, Lang D, Altmann S, Cyprys P, Zolg DP, Mathieson T, Bantscheff M, Hazarika RR, Schmidt T, Dawid C, Dunkel A, Hofmann T, Sprunck S, Falter-Braun P, Johannes F, Mayer KFX, Jürgens G, Wilhelm M, Baumbach J, Grill E, Schneitz K, Schwechheimer C, Kuster B. Mass-spectrometry-based draft of the Arabidopsis proteome. Nature. 2020 579(7799):409-414, PubMed: 32188942 |
| LFPT | PXD013868 | Mergner J, Frejno M, List M, Papacek M, Chen X, Chaudhary A, Samaras P, Richter S, Shikata H, Messerer M, Lang D, Altmann S, Cyprys P, Zolg DP, Mathieson T, Bantscheff M, Hazarika RR, Schmidt T, Dawid C, Dunkel A, Hofmann T, Sprunck S, Falter-Braun P, Johannes F, Mayer KFX, Jürgens G, Wilhelm M, Baumbach J, Grill E, Schneitz K, Schwechheimer C, Kuster B. Mass-spectrometry-based draft of the Arabidopsis proteome. Nature. 2020 579(7799):409-414, PubMed: 32188942 |
| Root | PXD012708 | Zhang H, Liu P, Guo T, Zhao H, Bensaddek D, Aebersold R, Xiong L. Arabidopsis proteome and the mass spectral assay library. Sci Data. 2019 6(1):278, PubMed: 31757973 |
| RT | PXD013868 | Mergner J, Frejno M, List M, Papacek M, Chen X, Chaudhary A, Samaras P, Richter S, Shikata H, Messerer M, Lang D, Altmann S, Cyprys P, Zolg DP, Mathieson T, Bantscheff M, Hazarika RR, Schmidt T, Dawid C, Dunkel A, Hofmann T, Sprunck S, Falter-Braun P, Johannes F, Mayer KFX, Jürgens G, Wilhelm M, Baumbach J, Grill E, Schneitz K, Schwechheimer C, Kuster B. Mass-spectrometry-based draft of the Arabidopsis proteome. Nature. 2020 579(7799):409-414, PubMed: 32188942 |
| RTTP | PXD13868 | Mergner J, Frejno M, List M, Papacek M, Chen X, Chaudhary A, Samaras P, Richter S, Shikata H, Messerer M, Lang D, Altmann S, Cyprys P, Zolg DP, Mathieson T, Bantscheff M, Hazarika RR, Schmidt T, Dawid C, Dunkel A, Hofmann T, Sprunck S, Falter-Braun P, Johannes F, Mayer KFX, Jürgens G, Wilhelm M, Baumbach J, Grill E, Schneitz K, Schwechheimer C, Kuster B. Mass-spectrometry-based draft of the Arabidopsis proteome. Nature. 2020 579(7799):409-414, PubMed: 32188942 |

| | | |
|---|---|---|
| **RTUZ** | **PXD13868** | Mergner J, Frejno M, List M, Papacek M, Chen X, Chaudhary A, Samaras P, Richter S, Shikata H, Messerer M, Lang D, Altmann S, Cyprys P, Zolg DP, Mathieson T, Bantscheff M, Hazarika RR, Schmidt T, Dawid C, Dunkel A, Hofmann T, Sprunck S, Falter-Braun P, Johannes F, Mayer KFX, Jürgens G, Wilhelm M, Baumbach J, Grill E, Schneitz K, Schwechheimer C, Kuster B. Mass-spectrometry-based draft of the Arabidopsis proteome. Nature. 2020 579(7799):409-414, PubMed: 32188942 |
| RTTOF | **PXD0012710** | Zhang H, Liu P, Guo T, Zhao H, Bensaddek D, Aebersold R, Xiong L. Arabidopsis proteome and the mass spectral assay library. Sci Data. 2019 6(1):278, PubMed: 31757973 |
| - | **PXD000869** | Zhang S, Zhang H, Xia Y, Xiong L. The caseinolytic protease complex component CLPC1 in Arabidopsis maintains proteome and RNA homeostasis in chloroplasts. BMC Plant Biol. 2018 18(1):192, PubMed: 30208840 |
| - | **PXD009725** | Takáč T, Pechan T, Šamajová O, Šamaj J. Proteomic Analysis of Arabidopsis pldα1 Mutants Revealed an Important Role of Phospholipase D Alpha 1 in Chloroplast Biogenesis. Front Plant Sci. 2019 10:89, PubMed: 30833950 |
| - | **PXD004742** | Subramanian S, Souleimanov A, Smith DL. Proteomic Studies on the Effects of Lipo-Chitooligosaccharide and Thuricin 17 under Unstressed and Salt Stressed Conditions in Arabidopsis thaliana. Front Plant Sci. 2016 Aug 30;7:1314. eCollection 2016, PubMed: 27625672 |
| - | **PXD004025** | Al Shweiki MR, Mönchgesang S, Majovsky P, Thieme D, Trutschel D, Hoehenwarter W. Assessment of Label-Free Quantification in Discovery Proteomics and Impact of Technological Factors and Natural Variability of Protein Abundance. J Proteome Res. 2017 Feb 28, PubMed: 28217993 |
| - | **PXD014292** | Fuchs P, Rugen N, Carrie C, Elsässer M, Finkemeier I, Giese J, Hildebrandt TM, Kühn K, Maurino VG, Ruberti C, Schallenberg-Rüdinger M, Steinbeck J, Braun HP, Eubel H, Meyer EH, Müller-Schüssele SJ, Schwarzländer M. Single organelle function and organization as estimated from Arabidopsis mitochondrial proteomics. Plant J. 2019, PubMed: 31520498 |
