## Supplementary material for "Evidences for a nutritional role of iodine in plants": Table S3

| | Control | $KIO_3$ 0.20 µM | $KIO_3$ 10 µM | Significance |
|---|---|---|---|---|
| Rosette FW (g) | 1.22 ± 0,040 | 1.19 ± 0.033 | 1.27 ± 0,033 | n.s. |
| Inflorescence FW (g) | 0.25 ± 0.031 [b] | 0.39 ± 0.038 [a] | 0.39 ± 0.032 [a] | *P-value* ≤0.005 |
| Rosette DW (g) | 0.11 ± 0.004 | 0.11 ± 0.004 | 0.12 ± 0.004 | n.s. |
| Inflorescence DW (g) | 0.03 ± 0.003 [b] | 0.05 ± 0.004 [a] | 0.04 ± 0.004 [a] | *P-value* ≤0.005 |
| Seed production* (g) | 7.00 ± 1.130 | 10.50 ± 1.751 | 9.47 ± 2.062 | n.s. |
| Seed/silique (n°) | 58.59 ± 1.184 | 59.09 ± 1.329 | 59.70 ± 0.740 | n.s. |

* Seed production per hydroponic tray, consisting of 15 plants.
