## Supplementary material for "Evidences for a nutritional role of iodine in plants": Table S4

| AGI code | Name |
|---|---|
| **At4g10500** | **2-oxoglutarate (2OG) and Fe(II)-dependent oxygenase superfamily protein(AT4G10500)** |
| GOTERM_BP_DIRECT | defense response to oomycetes, response to oomycetes, response to bacterium, response to fungus, response to salicylic acid, leaf senescence, secondary metabolic process, salicylic acid catabolic process, oxidation-reduction process, |
| GOTERM_MF_DIRECT | oxidoreductase activity, acting on paired donors, with incorporation or reduction of molecular oxygen, 2-oxoglutarate as one donor, and incorporation of one atom each of oxygen into both donors, metal ion binding, dioxygenase activity, |
| **At2g39030** | **Acyl-CoA N-acyltransferases (NAT) superfamily protein(NATA1)** |
| GOTERM_BP_DIRECT | ornithine metabolic process, defense response, response to jasmonic acid, |
| GOTERM_CC_DIRECT | cytoplasm, chloroplast, |
| GOTERM_MF_DIRECT | N-acetyltransferase activity, |
| **At3g18770** | **Autophagy-related protein 13(AT3G18770)** |
| GOTERM_BP_DIRECT | autophagy, protein transport, |
| GOTERM_CC_DIRECT | nucleus, autophagosome, cytoplasmic vesicle, ATG1/ULK1 kinase complex, |
| **At3g22620** | **Bifunctional inhibitor/lipid-transfer protein/seed storage 2S albumin superfamily protein(AT3G22620)** |
| GOTERM_BP_DIRECT | lipid transport, |
| GOTERM_CC_DIRECT | plasma membrane, chloroplast envelope, |
| GOTERM_MF_DIRECT | lipid binding, |
| **At3g13310** | **Chaperone DnaJ-domain superfamily protein(AT3G13310)** |
| GOTERM_BP_DIRECT | protein folding, |
| GOTERM_CC_DIRECT | chloroplast, |
| **At4g16260** | Glycosyl hydrolase superfamily protein(AT4G16260) |
| GOTERM_BP_DIRECT | defense response to nematode, carbohydrate metabolic process, response to salt stress, defense response to fungus, incompatible interaction, |
| GOTERM_CC_DIRECT | extracellular space, cell wall, vacuolar membrane, anchored component of plasma membrane, apoplast, |
| GOTERM_MF_DIRECT | hydrolase activity, hydrolyzing O-glycosyl compounds, polysaccharide binding, glucan endo-1,3-beta-D-glucosidase activity, |
| **At4g38600** | **HECT ubiquitin protein ligase family protein KAK(KAK)** |
| GOTERM_BP_DIRECT | trichome branching, DNA endoreduplication, protein ubiquitination involved in ubiquitin-dependent protein catabolic process, |
| GOTERM_CC_DIRECT | nucleus, cytoplasm, plasma membrane, |
| GOTERM_MF_DIRECT | ubiquitin-protein transferase activity, ligase activity, |
| **At3g01350** | **Major facilitator superfamily protein(AT3G01350)** |
| GOTERM_BP_DIRECT | transport, oligopeptide transport, |
| GOTERM_CC_DIRECT | plasma membrane, membrane, integral component of membrane, |

| | | |
|---|---|---|
| | GOTERM_MF_DIRECT | transporter activity, |
| | **At1g56010** | **NAC domain containing protein 1(NAC1)** |
| | GOTERM_BP_DIRECT | transcription, DNA-templated, regulation of transcription, DNA-templated, multicellular organism development, auxin-activated signaling pathway, primary shoot apical meristem specification, lateral root development, |
| | GOTERM_CC_DIRECT | nucleus, |
| | GOTERM_MF_DIRECT | DNA binding, transcription factor activity, sequence-specific DNA binding, |
| | **At5g52390** | **PAR1 protein(AT5G52390)** |
| | **At3g54150** | S-adenosyl-L-methionine-dependent methyltransferases superfamily protein(AT3G54150) |
| | GOTERM_BP_DIRECT | pollen germination, pollen exine formation, methylation, |
| | GOTERM_CC_DIRECT | nucleus, endosome, vacuolar membrane, trans-Golgi network, cytosol, |
| | GOTERM_MF_DIRECT | S-adenosylmethionine-dependent methyltransferase activity, |
| | **At3g28220** | **TRAF-like family protein(AT3G28220)** |
| | GOTERM_CC_DIRECT | cytoplasm, vacuole, chloroplast envelope, integral component of membrane, |
| | **At4g01390** | **TRAF-like family protein(AT4G01390)** |
| | GOTERM_CC_DIRECT | extracellular region, nucleus, cytoplasm, |
| | **At4g34131** | UDP-glucosyl transferase 73B3(UGT73B3) |
| | GOTERM_BP_DIRECT | defense response, metabolic process, flavonoid biosynthetic process, response to other organism, flavonoid glucuronidation, |
| | GOTERM_CC_DIRECT | intracellular membrane-bounded organelle, |
| | GOTERM_MF_DIRECT | UDP-glycosyltransferase activity, abscisic acid glucosyltransferase activity, transferase activity, transferring hexosyl groups, UDP-glucosyltransferase activity, flavonol 3-O-glucosyltransferase activity, quercetin 3-O-glucosyltransferase activity, quercetin 7-O-glucosyltransferase activity, daphnetin 3-O-glucosyltransferase activity, |
| | **At2g26400** | **acireductone dioxygenase 3(ARD3**) |
| | GOTERM_BP_DIRECT | methionine metabolic process, L-methionine biosynthetic process from methylthioadenosine, oxidation-reduction process, |
| | GOTERM_CC_DIRECT | nucleus, cytoplasm, plasma membrane, |
| | GOTERM_MF_DIRECT | iron ion binding, heteropolysaccharide binding, acireductone dioxygenase [iron(II)-requiring] activity, metal ion binding, |
| | **At5g61160** | **anthocyanin 5-aromatic acyltransferase 1(AACT1)** |
| | GOTERM_CC_DIRECT | cytoplasm, |
| | GOTERM_MF_DIRECT | transferase activity, agmatine N4-coumaroyltransferase activity, |
| | **At3g12500** | **basic chitinase(HCHIB)** |
| | GOTERM_BP_DIRECT | polysaccharide catabolic process, chitin catabolic process, amino sugar metabolic process, plant-type hypersensitive response, jasmonic acid and ethylene-dependent systemic resistance, ethylene mediated signaling pathway, cell wall macromolecule catabolic process, killing of cells of other organism, response to cadmium ion, defense response to fungus, |
| | GOTERM_CC_DIRECT | extracellular region, intracellular, vacuole, vacuolar membrane, cytosol, |

| | | |
|---|---|---|
| | GOTERM_MF_DIRECT | chitinase activity, chitin binding, |
| | **At3g57260** | **beta-1,3-glucanase 2(BGL2)** |
| | GOTERM_BP_DIRECT | carbohydrate metabolic process, response to cold, systemic acquired resistance, |
| | GOTERM_CC_DIRECT | cell wall, vacuole, endoplasmic reticulum, chloroplast, anchored component of plasma membrane, apoplast, |
| | GOTERM_MF_DIRECT | glucan exo-1,3-beta-glucosidase activity, hydrolase activity, hydrolyzing O-glycosyl compounds, protein binding, cellulase activity, polysaccharide binding, glucan endo-1,3-beta-D-glucosidase activity, |
| | **At2g43570** | **chitinase(CHI)** |
| | GOTERM_BP_DIRECT | polysaccharide catabolic process, carbohydrate metabolic process, chitin catabolic process, amino sugar metabolic process, response to virus, systemic acquired resistance, leaf senescence, response to silver ion, cell wall macromolecule catabolic process, response to amitrole, |
| | GOTERM_CC_DIRECT | extracellular region, intracellular, plant-type cell wall, apoplast, |
| | GOTERM_MF_DIRECT | chitinase activity, chitin binding, |
| | **At2g30770** | **cytochrome P450 family 71 polypeptide(CYP71A13)** |
| | GOTERM_BP_DIRECT | defense response, response to bacterium, induced systemic resistance, camalexin biosynthetic process, defense response to bacterium, defense response to fungus, |
| | GOTERM_CC_DIRECT | membrane, integral component of membrane, |
| | GOTERM_MF_DIRECT | iron ion binding, protein binding, oxidoreductase activity, acting on paired donors, with incorporation or reduction of molecular oxygen, NAD(P)H as one donor, and incorporation of one atom of oxygen, oxygen binding, heme binding, indoleacetaldoxime dehydratase activity, |
| | **At1g21310** | **extensin 3(EXT3)** |
| | GOTERM_BP_DIRECT | plant-type cell wall organization, |
| | GOTERM_CC_DIRECT | extracellular region, primary cell wall, |
| | GOTERM_MF_DIRECT | structural constituent of cell wall, |
| | **At1g56600** | **galactinol synthase 2(GolS2)** |
| | GOTERM_BP_DIRECT | galactose metabolic process, response to oxidative stress, response to cold, response to water deprivation, response to salt stress, response to abscisic acid, carbohydrate biosynthetic process, |
| | GOTERM_CC_DIRECT | nucleus, cytoplasm, |
| | GOTERM_MF_DIRECT | transferase activity, transferring glycosyl groups, transferase activity, transferring hexosyl groups, metal ion binding, inositol 3-alpha-galactosyltransferase activity, |
| | **At5g39520** | **hypothetical protein (DUF1997)(AT5G39520)** |
| | GOTERM_CC_DIRECT | nucleus, |
| | **At1g64360** | **hypothetical protein(AT1G64360)** |
| | GOTERM_BP_DIRECT | response to oxidative stress, leaf senescence, |
| | GOTERM_CC_DIRECT | nucleus, |
| | **At3g10120** | **hypothetical protein(AT3G10120**) |
| | GOTERM_CC_DIRECT | nucleus, |
| | **At3g09940** | monodehydroascorbate reductase(MDHAR) |

| | | |
|---|---|---|
| | GOTERM_BP_DIRECT | response to water deprivation, response to symbiotic fungus, response to salt stress, response to jasmonic acid, regulation of symbiosis, encompassing mutualism through parasitism, oxidation-reduction process, |
| | GOTERM_CC_DIRECT | cytoplasm, cytosol, |
| | GOTERM_MF_DIRECT | oxidoreductase activity, monodehydroascorbate reductase (NADH) activity, flavin adenine dinucleotide binding, |
| | **At3g48920** | **myb domain protein 45(MYB45)** |
| | GOTERM_BP_DIRECT | regulation of transcription, DNA-templated, regulation of transcription from RNA polymerase II promoter, response to salicylic acid, cell differentiation, |
| | GOTERM_CC_DIRECT | nucleus, |
| | GOTERM_MF_DIRECT | RNA polymerase II transcription factor activity, sequence-specific DNA binding, transcription factor activity, RNA polymerase II transcription factor recruiting, DNA binding, transcription factor activity, sequence-specific DNA binding, sequence-specific DNA binding, transcription regulatory region DNA binding, |
| | **At1g06160** | **octadecanoid-responsive AP2/ERF 59(ORA59)** |
| | GOTERM_BP_DIRECT | transcription, DNA-templated, regulation of transcription, DNA-templated, response to ethylene, response to jasmonic acid, jasmonic acid and ethylene-dependent systemic resistance, ethylene-activated signaling pathway, |
| | GOTERM_CC_DIRECT | intracellular, nucleus, |
| | GOTERM_MF_DIRECT | DNA binding, transcription factor activity, sequence-specific DNA binding, |
| | **At2g14610** | **pathogenesis-related protein 1(PR1)** |
| | GOTERM_BP_DIRECT | defense response, response to water deprivation, systemic acquired resistance, response to vitamin B1, |
| | GOTERM_CC_DIRECT | extracellular region, cell wall, apoplast, |
| | **At1g75040** | **pathogenesis-related protein 5(PR5)** |
| | GOTERM_BP_DIRECT | response to virus, systemic acquired resistance, response to UV-B, regulation of anthocyanin biosynthetic process, response to cadmium ion, response to other organism, |
| | GOTERM_CC_DIRECT | extracellular region, cell wall, vacuole, apoplast, |
| | **At2g18660** | **plant natriuretic peptide A(PNP-A)** |
| | GOTERM_BP_DIRECT | amino sugar metabolic process, systemic acquired resistance, alternative respiration, |
| | GOTERM_CC_DIRECT | extracellular region, cell wall, intracellular, apoplast, |
| | GOTERM_MF_DIRECT | chitinase activity, |
| | **At1g17020** | **senescence-related gene 1(SRG1)** |
| | GOTERM_BP_DIRECT | flavonoid biosynthetic process, leaf senescence, oxidation-reduction process, |
| | GOTERM_CC_DIRECT | cytoplasm, |
| | GOTERM_MF_DIRECT | oxidoreductase activity, acting on diphenols and related substances as donors, oxygen as acceptor, oxidoreductase activity, acting on paired donors, with incorporation or reduction of molecular oxygen, 2-oxoglutarate as one donor, and incorporation of one atom each of oxygen into both donors, metal ion binding, dioxygenase activity, |
