## Supplementary material for "Evidences for a nutritional role of iodine in plants": Table S5

| AGI code | Name |
|---|---|
| **At5g38120** | **AMP-dependent synthetase and ligase family protein(4CL8)** |
| GOTERM_BP_DIRECT | metabolic process, |
| GOTERM_CC_DIRECT | peroxisome, chloroplast, |
| GOTERM_MF_DIRECT | ATP binding, 4-coumarate-CoA ligase activity, ligase activity, |
| **At1g23240** | **Caleosin-related family protein(AT1G23240)** |
| GOTERM_BP_DIRECT | oxidation-reduction process, |
| GOTERM_CC_DIRECT | extracellular region, lipid particle, |
| GOTERM_MF_DIRECT | lipase activity, metal ion binding, plant seed peroxidase activity, |
| **At5g62210** | **Embryo-specific protein 3, (ATS3)(AT5G62210)** |
| GOTERM_BP_DIRECT | response to karrikin, |
| GOTERM_CC_DIRECT | anchored component of membrane, |
| **At1g75750** | **GAST1 protein homolog 1(GASA1)** |
| GOTERM_BP_DIRECT | response to abscisic acid, response to gibberellin, gibberellic acid mediated signaling pathway, response to brassinosteroid, unidimensional cell growth, |
| GOTERM_CC_DIRECT | extracellular region, cell wall, plant-type cell wall, |
| **At3g05740** | **RECQ helicase l1(RECQl1)** |
| GOTERM_BP_DIRECT | double-strand break repair via homologous recombination, cellular response to water deprivation, cellular response to cold, |
| GOTERM_CC_DIRECT | nucleus, chromosome, cytoplasm, |
| GOTERM_MF_DIRECT | DNA binding, ATP binding, ATP-dependent helicase activity, four-way junction helicase activity, ATP-dependent 3'-5' DNA helicase activity, metal ion binding, |
| **At1g06710** | **Tetratricopeptide repeat (TPR)-like superfamily protein(AT1G06710)** |
| GOTERM_CC_DIRECT | mitochondrion, |
| **At4g19191** | **Tetratricopeptide repeat (TPR)-like superfamily protein(AT4G19191)** |
| GOTERM_CC_DIRECT | mitochondrion, |
| **At4g15680** | **Thioredoxin superfamily protein(AT4G15680)** |
| GOTERM_BP_DIRECT | cell redox homeostasis, |
| GOTERM_CC_DIRECT | cell, nucleus, cytoplasm, |
| GOTERM_MF_DIRECT | arsenate reductase (glutaredoxin) activity, electron carrier activity, protein disulfide oxidoreductase activity, metal ion binding, 2 iron, 2 sulfur cluster binding, |
| **At4g15700** | **Thioredoxin superfamily protein(AT4G15700)** |
| GOTERM_BP_DIRECT | cell redox homeostasis, |
| GOTERM_CC_DIRECT | cell, nucleus, cytoplasm, |
| GOTERM_MF_DIRECT | arsenate reductase (glutaredoxin) activity, electron carrier activity, protein disulfide oxidoreductase activity, metal ion binding, 2 iron, 2 sulfur cluster binding, |

| | | |
|---|---|---|
| | **At1g02970** | **WEE1-like kinase(WEE1)** |
| | GOTERM_BP_DIRECT | DNA replication checkpoint, cell cycle arrest, mitotic nuclear division, cell division, |
| | GOTERM_CC_DIRECT | nucleus, |
| | GOTERM_MF_DIRECT | protein kinase activity, non-membrane spanning protein tyrosine kinase activity, protein binding, ATP binding, kinase activity, metal ion binding, |
| | **At4g23280** | **cysteine-rich RLK (RECEPTOR-like protein kinase) 20(CRK20)** |
| | GOTERM_BP_DIRECT | protein phosphorylation, defense response, response to salicylic acid, programmed cell death, defense response to bacterium, |
| | GOTERM_CC_DIRECT | extracellular region, plasma membrane, plasmodesma, chloroplast, integral component of membrane, |
| | GOTERM_MF_DIRECT | protein serine/threonine kinase activity, ATP binding, kinase activity, |
| | **At5g01600** | **ferretin 1(FER1)** |
| | GOTERM_BP_DIRECT | response to reactive oxygen species, iron ion transport, cellular iron ion homeostasis, intracellular sequestering of iron ion, response to cold, response to bacterium, response to cytokinin, flower development, response to iron ion, response to zinc ion, photosynthesis, response to hydrogen peroxide, leaf development, iron ion homeostasis, oxidation-reduction process, |
| | GOTERM_CC_DIRECT | cell, mitochondrion, chloroplast, chloroplast thylakoid membrane, chloroplast stroma, thylakoid, membrane, |
| | GOTERM_MF_DIRECT | ferroxidase activity, iron ion binding, ferric iron binding, |
| | **At2g23560** | **methyl esterase 7(MES7)** |
| | GOTERM_BP_DIRECT | systemic acquired resistance, salicylic acid metabolic process, defense response to fungus, incompatible interaction, |
| | GOTERM_CC_DIRECT | cytoplasm, |
| | GOTERM_MF_DIRECT | hydrolase activity, hydrolase activity, acting on ester bonds, methyl indole-3-acetate esterase activity, methyl salicylate esterase activity, |
| | **At1g20310** | **syringolide-induced protein(AT1G20310)** |
| | **At2g36885** | **translation initiation factor(AT2G36885)** |
| | GOTERM_CC_DIRECT | plasma membrane, chloroplast, integral component of membrane, |
