## Supplementary material for "Evidences for a nutritional role of iodine in plants": Table S6

| AGI code | Name |
|---|---|
| **At4g10500** | **2-oxoglutarate (2OG) and Fe(II)-dependent oxygenase superfamily protein(AT4G10500)** |
| GOTERM_BP_DIRECT | defense response to oomycetes, response to oomycetes, response to bacterium, response to fungus, response to salicylic acid, leaf senescence, secondary metabolic process, salicylic acid catabolic process, oxidation-reduction process, |
| GOTERM_MF_DIRECT | oxidoreductase activity, acting on paired donors, with incorporation or reduction of molecular oxygen, 2-oxoglutarate as one donor, and incorporation of one atom each of oxygen into both donors, metal ion binding, dioxygenase activity, |
| **At5g59540** | **2-oxoglutarate (2OG) and Fe(II)-dependent oxygenase superfamily protein(AT5G59540)** |
| GOTERM_BP_DIRECT | oxidation-reduction process, |
| GOTERM_CC_DIRECT | cytoplasm, |
| GOTERM_MF_DIRECT | oxidoreductase activity, metal ion binding, dioxygenase activity, |
| **At3g55090** | **ABC-2 type transporter family protein(ABCG16)** |
| GOTERM_BP_DIRECT | pollen wall assembly, |
| GOTERM_CC_DIRECT | plasma membrane, integral component of membrane, |
| GOTERM_MF_DIRECT | ATP binding, ATPase activity, coupled to transmembrane movement of substances, |
| **At2g37360** | **ABC-2 type transporter family protein(ABCG2)** |
| GOTERM_BP_DIRECT | transport, suberin biosynthetic process, |
| GOTERM_CC_DIRECT | plasma membrane, integral component of membrane, |
| GOTERM_MF_DIRECT | ATP binding, ATPase activity, ATPase activity, coupled to transmembrane movement of substances, |
| **At3g47780** | **ABC2 homolog 6(ABCA7)** |
| GOTERM_BP_DIRECT | lipid transport, |
| GOTERM_CC_DIRECT | plasma membrane, plasmodesma, integral component of membrane, intracellular membrane-bounded organelle, |
| GOTERM_MF_DIRECT | transporter activity, ATP binding, ATPase activity, coupled to transmembrane movement of substances, |
| **At1g74710** | **ADC synthase superfamily protein(EDS16)** |
| GOTERM_BP_DIRECT | defense response, biosynthetic process, response to cold, response to bacterium, systemic acquired resistance, salicylic acid biosynthetic process, stomatal movement, negative regulation of defense response, phylloquinone biosynthetic process, defense response to bacterium, defense response to fungus, |
| GOTERM_CC_DIRECT | chloroplast, plastid, |
| GOTERM_MF_DIRECT | isochorismate synthase activity, |
| **At3g25250** | **AGC (cAMP-dependent, cGMP-dependent and protein kinase C) kinase family protein(AGC2-1)** |
| GOTERM_BP_DIRECT | protein phosphorylation, defense response, response to oxidative stress, response to wounding, intracellular signal transduction, |
| GOTERM_CC_DIRECT | nucleus, cytoplasm, plasma membrane, |
| GOTERM_MF_DIRECT | protein kinase activity, protein serine/threonine kinase activity, protein binding, ATP binding, kinase activity, protein kinase binding, |
| **At2g13810** | **AGD2-like defense response protein 1(ALD1)** |

| | |
|---|---|
| GOTERM_BP_DIRECT | lysine biosynthetic process via diaminopimelate, systemic acquired resistance, salicylic acid mediated signaling pathway, leaf senescence, defense response to bacterium, |
| GOTERM_CC_DIRECT | chloroplast, |
| GOTERM_MF_DIRECT | transaminase activity, L,L-diaminopimelate aminotransferase activity, pyridoxal phosphate binding, |
| **At3g28930** | **AIG2-like (avirulence induced gene) family protein(AIG2)** |
| GOTERM_BP_DIRECT | response to bacterium, |
| GOTERM_CC_DIRECT | nucleus, cytosol, |
| GOTERM_MF_DIRECT | transferase activity, transferring acyl groups, |
| **At1g77240** | **AMP-dependent synthetase and ligase family protein(AAE2)** |
| GOTERM_BP_DIRECT | fatty acid metabolic process, metabolic process, |
| GOTERM_CC_DIRECT | mitochondrion, |
| GOTERM_MF_DIRECT | catalytic activity, ligase activity, |
| **At2g26530** | **AR781, pheromone receptor-like protein (DUF1645)(AR781)** |
| GOTERM_CC_DIRECT | plasma membrane, chloroplast, |
| **At3g02840** | **ARM repeat superfamily protein(AT3G02840)** |
| GOTERM_BP_DIRECT | response to ozone, response to other organism, |
| GOTERM_CC_DIRECT | nucleus, cytoplasm, |
| **At5g67340** | **ARM repeat superfamily protein(AT5G67340)** |
| GOTERM_BP_DIRECT | protein ubiquitination, |
| GOTERM_CC_DIRECT | nucleus, cytoplasm, |
| GOTERM_MF_DIRECT | ubiquitin-protein transferase activity, ligase activity, |
| **At1g57650** | **ATP binding protein(AT1G57650)** |
| GOTERM_BP_DIRECT | defense response, |
| GOTERM_CC_DIRECT | nucleus, |
| **At2g30660** | **ATP-dependent caseinolytic (Clp) protease/crotonase family protein(AT2G30660)** |
| GOTERM_BP_DIRECT | valine catabolic process, fatty acid beta-oxidation, branched-chain amino acid catabolic process, |
| GOTERM_CC_DIRECT | peroxisome, |
| GOTERM_MF_DIRECT | 3-hydroxyisobutyryl-CoA hydrolase activity, |
| **At3g22910** | **ATPase E1-E2 type family protein / haloacid dehalogenase-like hydrolase family protein(AT3G22910)** |
| GOTERM_BP_DIRECT | calcium ion transmembrane transport, |
| GOTERM_CC_DIRECT | integral component of plasma membrane, intracellular membrane-bounded organelle, |
| GOTERM_MF_DIRECT | calcium-transporting ATPase activity, calmodulin binding, ATP binding, metal ion binding, |

| | | |
|---|---|---|
| | **At3g63380** | **ATPase E1-E2 type family protein / haloacid dehalogenase-like hydrolase family protein(AT3G63380)** |
| | GOTERM_BP_DIRECT | calcium ion transmembrane transport, |
| | GOTERM_CC_DIRECT | integral component of plasma membrane, integral component of membrane, intracellular membrane-bounded organelle, |
| | GOTERM_MF_DIRECT | calcium-transporting ATPase activity, calmodulin binding, ATP binding, metal ion binding, |
| | **At2g32020** | **Acyl-CoA N-acyltransferases (NAT) superfamily protein(AT2G32020)** |
| | GOTERM_BP_DIRECT | N-terminal protein amino acid acetylation, response to abscisic acid, |
| | GOTERM_CC_DIRECT | cytoplasm, |
| | GOTERM_MF_DIRECT | N-acetyltransferase activity, |
| | **At5g67430** | **Acyl-CoA N-acyltransferases (NAT) superfamily protein(AT5G67430)** |
| | GOTERM_BP_DIRECT | N-terminal protein amino acid acetylation, |
| | GOTERM_CC_DIRECT | nucleus, protein acetyltransferase complex, |
| | GOTERM_MF_DIRECT | peptide alpha-N-acetyltransferase activity, N-acetyltransferase activity, |
| | **At4g26120** | **Ankyrin repeat family protein / BTB/POZ domain-containing protein(AT4G26120)** |
| | GOTERM_BP_DIRECT | response to chitin, protein ubiquitination, |
| | **At2g24600** | **Ankyrin repeat family protein(AT2G24600)** |
| | GOTERM_BP_DIRECT | signal transduction, |
| | GOTERM_CC_DIRECT | membrane, integral component of membrane, |
| | **At5g50140** | **Ankyrin repeat family protein(AT5G50140)** |
| | GOTERM_BP_DIRECT | signal transduction, |
| | GOTERM_CC_DIRECT | membrane, integral component of membrane, |
| | **At5g54700** | **Ankyrin repeat family protein(AT5G54700)** |
| | GOTERM_BP_DIRECT | signal transduction, |
| | GOTERM_CC_DIRECT | membrane, integral component of membrane, |
| | **At5g54720** | **Ankyrin repeat family protein(AT5G54720)** |
| | **At2g29870** | **Aquaporin-like superfamily protein(AT2G29870)** |
| | GOTERM_BP_DIRECT | transport, cellular water homeostasis, ion transmembrane transport, |
| | GOTERM_CC_DIRECT | integral component of plasma membrane, chloroplast, membrane, |
| | GOTERM_MF_DIRECT | water channel activity, glycerol channel activity, |
| | **At1g76520** | **Auxin efflux carrier family protein(AT1G76520)** |
| | GOTERM_BP_DIRECT | response to auxin, auxin-activated signaling pathway, auxin polar transport, transmembrane transport, |
| | GOTERM_CC_DIRECT | endoplasmic reticulum membrane, integral component of membrane, |

| | | |
|---|---|---|
| | GOTERM_MF_DIRECT | auxin:proton symporter activity, |
| | **At5g13320** | **Auxin-responsive GH3 family protein(PBS3)** |
| | GOTERM_BP_DIRECT | defense response, plant-type hypersensitive response, response to auxin, defense response to bacterium, incompatible interaction, salicylic acid mediated signaling pathway, regulation of systemic acquired resistance, detection of fungus, benzoate metabolic process, positive regulation of plant-type hypersensitive response, defense response to bacterium, interspecies interaction between organisms, cellular response to hypoxia, |
| | GOTERM_CC_DIRECT | cytoplasm, |
| | GOTERM_MF_DIRECT | ligase activity, 4-aminobenzoate amino acid synthetase activity, benzoate amino acid synthetase activity, vanillate amino acid synthetase activity, 4-hydroxybenzoate amino acid synthetase activity, |
| | **At3g29970** | **B12D protein(AT3G29970)** |
| | GOTERM_CC_DIRECT | mitochondrion, integral component of membrane, |
| | **At3g61190** | **BON association protein 1(BAP1)** |
| | GOTERM_BP_DIRECT | defense response, response to temperature stimulus, response to heat, response to cold, response to wounding, response to salicylic acid, cellular homeostasis, negative regulation of defense response, |
| | GOTERM_CC_DIRECT | cell, mitochondrion, membrane, |
| | GOTERM_MF_DIRECT | protein binding, phospholipid binding, |
| | **At2g18370** | **Bifunctional inhibitor/lipid-transfer protein/seed storage 2S albumin superfamily protein(AT2G18370)** |
| | GOTERM_BP_DIRECT | lipid transport, |
| | GOTERM_CC_DIRECT | extracellular region, membrane, |
| | GOTERM_MF_DIRECT | lipid binding, |
| | **At3g58550** | **Bifunctional inhibitor/lipid-transfer protein/seed storage 2S albumin superfamily protein(AT3G58550)** |
| | GOTERM_BP_DIRECT | proteolysis, lipid transport, |
| | GOTERM_CC_DIRECT | plasma membrane, anchored component of membrane, |
| | GOTERM_MF_DIRECT | peptidase activity, lipid binding, |
| | **At2g37430** | **C2H2 and C2HC zinc fingers superfamily protein(ZAT11)** |
| | GOTERM_BP_DIRECT | transcription, DNA-templated, regulation of transcription, DNA-templated, response to chitin, cellular response to nickel ion, regulation of root development, |
| | GOTERM_CC_DIRECT | nucleus, |
| | GOTERM_MF_DIRECT | nucleic acid binding, transcription factor activity, sequence-specific DNA binding, zinc ion binding, sequence-specific DNA binding, transcription regulatory region DNA binding, metal ion binding, |
| | **At3g53600** | **C2H2-type zinc finger family protein(AT3G53600)** |
| | GOTERM_BP_DIRECT | regulation of transcription, DNA-templated, response to chitin, |
| | GOTERM_CC_DIRECT | nucleus, |
| | GOTERM_MF_DIRECT | nucleic acid binding, transcription factor activity, sequence-specific DNA binding, zinc ion binding, sequence-specific DNA binding, transcription regulatory region DNA binding, metal ion binding, |
| | **At5g59820** | **C2H2-type zinc finger family protein(RHL41)** |

| | |
|---|---|
| GOTERM_BP_DIRECT | transcription, DNA-templated, regulation of transcription, DNA-templated, response to oxidative stress, response to heat, response to cold, response to light stimulus, response to wounding, cold acclimation, photosynthetic acclimation, response to chitin, response to UV-B, hyperosmotic salinity response, |
| GOTERM_CC_DIRECT | nucleus, |
| GOTERM_MF_DIRECT | nucleic acid binding, transcription factor activity, sequence-specific DNA binding, zinc ion binding, sequence-specific DNA binding, transcription regulatory region DNA binding, metal ion binding, |
| **At4g33710** | **CAP (Cysteine-rich secretory proteins, Antigen 5, and Pathogenesis-related 1 protein) superfamily protein(AT4G33710)** |
| GOTERM_CC_DIRECT | extracellular region, |
| **At1g48260** | **CBL-interacting protein kinase 17(CIPK17)** |
| GOTERM_BP_DIRECT | protein phosphorylation, signal transduction, intracellular signal transduction, |
| GOTERM_CC_DIRECT | nucleus, cytoplasm, plasma membrane, |
| GOTERM_MF_DIRECT | protein serine/threonine kinase activity, ATP binding, kinase activity, |
| **At1g19020** | **CDP-diacylglycerol-glycerol-3-phosphate 3-phosphatidyltransferase(AT1G19020)** |
| GOTERM_BP_DIRECT | response to oxidative stress, |
| **At2g41100** | **Calcium-binding EF hand family protein(TCH3)** |
| GOTERM_BP_DIRECT | response to temperature stimulus, response to mechanical stimulus, response to absence of light, thigmotropism, |
| GOTERM_CC_DIRECT | mitochondrion, vacuolar membrane, plasmodesma, |
| GOTERM_MF_DIRECT | calcium ion binding, |
| **At2g41090** | **Calcium-binding EF-hand family protein(AT2G41090)** |
| GOTERM_BP_DIRECT | cellular response to oxidative stress, regulation of L-ascorbic acid biosynthetic process, |
| GOTERM_CC_DIRECT | cytoplasm, plasma membrane, |
| GOTERM_MF_DIRECT | calcium ion binding, protein binding, |
| **At3g01830** | **Calcium-binding EF-hand family protein(AT3G01830)** |
| GOTERM_CC_DIRECT | nucleus, |
| GOTERM_MF_DIRECT | calcium ion binding, |
| **At3g47480** | **Calcium-binding EF-hand family protein(AT3G47480)** |
| GOTERM_CC_DIRECT | plasma membrane, |
| GOTERM_MF_DIRECT | calcium ion binding, |
| **At5g39670** | **Calcium-binding EF-hand family protein(AT5G39670)** |
| GOTERM_CC_DIRECT | integral component of membrane, |
| GOTERM_MF_DIRECT | calcium ion binding, |
| **At1g73805** | **Calmodulin binding protein-like protein(SARD1)** |

| | | |
|---|---|---|
| | GOTERM_BP_DIRECT | defense response to oomycetes, transcription, DNA-templated, regulation of transcription, DNA-templated, response to stress, response to bacterium, plant-type hypersensitive response, regulation of systemic acquired resistance, response to UV-B, defense response to bacterium, cellular response to molecule of bacterial origin, regulation of salicylic acid biosynthetic process, positive regulation of defense response to bacterium, |
| | GOTERM_CC_DIRECT | nucleus, |
| | GOTERM_MF_DIRECT | transcription factor activity, sequence-specific DNA binding, calmodulin binding, sequence-specific DNA binding, |
| | **At5g26920** | **Cam-binding protein 60-like G(CBP60G)** |
| | GOTERM_BP_DIRECT | defense response to oomycetes, response to molecule of bacterial origin, transcription, DNA-templated, regulation of transcription, DNA-templated, response to bacterium, response to fungus, plant-type hypersensitive response, salicylic acid biosynthetic process, abscisic acid-activated signaling pathway, positive regulation of abscisic acid-activated signaling pathway, defense response to bacterium, incompatible interaction, regulation of systemic acquired resistance, response to UV-B, defense response to bacterium, cellular response to molecule of bacterial origin, cellular response to hypoxia, regulation of salicylic acid biosynthetic process, positive regulation of response to water deprivation, |
| | GOTERM_CC_DIRECT | nucleus, |
| | GOTERM_MF_DIRECT | transcription factor activity, sequence-specific DNA binding, calmodulin binding, zinc ion binding, sequence-specific DNA binding, |
| | **At3g47540** | **Chitinase family protein(AT3G47540)** |
| | GOTERM_BP_DIRECT | carbohydrate metabolic process, chitin catabolic process, amino sugar metabolic process, cell wall macromolecule catabolic process, |
| | GOTERM_CC_DIRECT | extracellular region, intracellular, |
| | GOTERM_MF_DIRECT | chitinase activity, |
| | **At1g15040** | **Class I glutamine amidotransferase-like superfamily protein(GAT1_2.1)** |
| | GOTERM_BP_DIRECT | glutamine metabolic process, regulation of secondary shoot formation, |
| | GOTERM_CC_DIRECT | cytosol, |
| | GOTERM_MF_DIRECT | transferase activity, hydrolase activity, |
| | **At2g29220** | **Concanavalin A-like lectin protein kinase family protein(AT2G29220)** |
| | GOTERM_BP_DIRECT | protein phosphorylation, |
| | GOTERM_CC_DIRECT | plasma membrane, integral component of membrane, |
| | GOTERM_MF_DIRECT | protein kinase activity, ATP binding, kinase activity, carbohydrate binding, |
| | **At2g29250** | **Concanavalin A-like lectin protein kinase family protein(AT2G29250)** |
| | GOTERM_BP_DIRECT | defense response to oomycetes, protein phosphorylation, |
| | GOTERM_CC_DIRECT | plasma membrane, integral component of membrane, |
| | GOTERM_MF_DIRECT | protein kinase activity, ATP binding, kinase activity, carbohydrate binding, |
| | **At4g28350** | **Concanavalin A-like lectin protein kinase family protein(AT4G28350)** |
| | GOTERM_CC_DIRECT | plasma membrane, integral component of membrane, |
| | GOTERM_MF_DIRECT | protein serine/threonine kinase activity, ATP binding, kinase activity, carbohydrate binding, |
| | **At5g65600** | **Concanavalin A-like lectin protein kinase family protein(AT5G65600)** |
| | GOTERM_BP_DIRECT | defense response to oomycetes, protein phosphorylation, positive regulation of hydrogen peroxide metabolic process, positive regulation of cell death, |

| | | |
|---|---|---|
| | GOTERM_CC_DIRECT | plasma membrane, integral component of membrane, |
| | GOTERM_MF_DIRECT | protein kinase activity, protein serine/threonine kinase activity, protein binding, ATP binding, kinase activity, carbohydrate binding, |
| | **At5g52670** | **Copper transport protein family(AT5G52670)** |
| | GOTERM_BP_DIRECT | metal ion transport, cellular transition metal ion homeostasis, |
| | GOTERM_CC_DIRECT | cytoplasm, |
| | GOTERM_MF_DIRECT | metal ion binding, transition metal ion binding, |
| | **At5g52680** | **Copper transport protein family(AT5G52680)** |
| | GOTERM_BP_DIRECT | metal ion transport, cellular transition metal ion homeostasis, |
| | GOTERM_CC_DIRECT | nucleus, cytoplasm, |
| | GOTERM_MF_DIRECT | metal ion binding, transition metal ion binding, |
| | **At5g52690** | **Copper transport protein family(AT5G52690)** |
| | GOTERM_BP_DIRECT | metal ion transport, cellular transition metal ion homeostasis, |
| | GOTERM_CC_DIRECT | cytoplasm, |
| | GOTERM_MF_DIRECT | metal ion binding, transition metal ion binding, |
| | **At5g52710** | **Copper transport protein family(AT5G52710)** |
| | GOTERM_BP_DIRECT | metal ion transport, cellular transition metal ion homeostasis, |
| | GOTERM_CC_DIRECT | nucleus, cytoplasm, |
| | GOTERM_MF_DIRECT | metal ion binding, transition metal ion binding, |
| | **At5g52760** | **Copper transport protein family(AT5G52760)** |
| | GOTERM_BP_DIRECT | metal ion transport, cellular transition metal ion homeostasis, |
| | GOTERM_CC_DIRECT | nucleus, cytoplasm, |
| | GOTERM_MF_DIRECT | metal ion binding, transition metal ion binding, |
| | **At4g39830** | **Cupredoxin superfamily protein(AT4G39830)** |
| | GOTERM_BP_DIRECT | oxidation-reduction process, |
| | GOTERM_CC_DIRECT | extracellular region, |
| | GOTERM_MF_DIRECT | copper ion binding, oxidoreductase activity, oxidizing metal ions, |
| | **At5g18470** | **Curculin-like (mannose-binding) lectin family protein(AT5G18470)** |
| | GOTERM_BP_DIRECT | response to karrikin, |
| | GOTERM_CC_DIRECT | plasma membrane, plant-type cell wall, |
| | GOTERM_MF_DIRECT | carbohydrate binding, |
| | **At2g44370** | **Cysteine/Histidine-rich C1 domain family protein(AT2G44370)** |

| | | |
|---|---|---|
| | GOTERM_CC_DIRECT | nucleus, cytoplasm, |
| | GOTERM_MF_DIRECT | thioredoxin-disulfide reductase activity, |
| | **At3g26830** | **Cytochrome P450 superfamily protein(PAD3)** |
| | GOTERM_BP_DIRECT | defense response, response to water deprivation, response to bacterium, response to insect, indole phytoalexin biosynthetic process, response to abscisic acid, regulation of systemic acquired resistance, camalexin biosynthetic process, defense response to fungus, oxidation-reduction process, |
| | GOTERM_CC_DIRECT | endoplasmic reticulum, membrane, integral component of membrane, intracellular membrane-bounded organelle, |
| | GOTERM_MF_DIRECT | monooxygenase activity, iron ion binding, dihydrocamalexic acid decarboxylase activity, oxidoreductase activity, acting on the CH-CH group of donors, NAD or NADP as acceptor, oxidoreductase activity, acting on paired donors, with incorporation or reduction of molecular oxygen, oxidoreductase activity, acting on paired donors, with incorporation or reduction of molecular oxygen, NAD(P)H as one donor, and incorporation of one atom of oxygen, oxygen binding, heme binding, |
| | **At2g46750** | **D-arabinono-1,4-lactone oxidase family protein(GulLO2)** |
| | GOTERM_BP_DIRECT | L-ascorbic acid biosynthetic process, oxidation-reduction process, |
| | GOTERM_CC_DIRECT | extracellular region, membrane, |
| | GOTERM_MF_DIRECT | D-arabinono-1,4-lactone oxidase activity, L-gulonolactone oxidase activity, flavin adenine dinucleotide binding, |
| | **At1g66090** | **Disease resistance protein (TIR-NBS class)(AT1G66090)** |
| | GOTERM_BP_DIRECT | defense response, signal transduction, |
| | GOTERM_CC_DIRECT | chloroplast, |
| | GOTERM_MF_DIRECT | ADP binding, |
| | **At1g72890** | **Disease resistance protein (TIR-NBS class)(AT1G72890)** |
| | GOTERM_BP_DIRECT | defense response, signal transduction, |
| | GOTERM_CC_DIRECT | cytoplasm, integral component of membrane, |
| | GOTERM_MF_DIRECT | ADP binding, |
| | **At1g63750** | **Disease resistance protein (TIR-NBS-LRR class) family(AT1G63750)** |
| | GOTERM_BP_DIRECT | defense response, signal transduction, |
| | GOTERM_CC_DIRECT | nucleus, integral component of membrane, |
| | GOTERM_MF_DIRECT | ADP binding, |
| | **At4g11170** | **Disease resistance protein (TIR-NBS-LRR class) family(AT4G11170)** |
| | GOTERM_BP_DIRECT | defense response, signal transduction, response to ozone, |
| | GOTERM_CC_DIRECT | cytoplasm, mitochondrion, |
| | GOTERM_MF_DIRECT | ADP binding, |
| | **At4g11340** | **Disease resistance protein (TIR-NBS-LRR class) family(AT4G11340)** |
| | GOTERM_BP_DIRECT | defense response, signal transduction, |
| | GOTERM_CC_DIRECT | cytoplasm, |
| | **At5g41740** | **Disease resistance protein (TIR-NBS-LRR class) family(AT5G41740)** |

| | | |
|---|---|---|
| | GOTERM_BP_DIRECT | defense response, signal transduction, |
| | GOTERM_CC_DIRECT | nucleus, |
| | GOTERM_MF_DIRECT | ADP binding, |
| | **At1g64160** | **Disease resistance-responsive (dirigent-like protein) family protein(DIR5)** |
| | GOTERM_BP_DIRECT | defense response, lignan biosynthetic process, (-)-pinoresinol biosynthetic process, |
| | GOTERM_CC_DIRECT | extracellular region, apoplast, |
| | GOTERM_MF_DIRECT | guiding stereospecific synthesis activity, |
| | **At1g69570** | **Dof-type zinc finger DNA-binding family protein(AT1G69570)** |
| | GOTERM_BP_DIRECT | transcription, DNA-templated, regulation of transcription, DNA-templated, flower development, |
| | GOTERM_CC_DIRECT | nucleus, |
| | GOTERM_MF_DIRECT | DNA binding, transcription factor activity, sequence-specific DNA binding, metal ion binding, |
| | **At1g13310** | **Endosomal targeting BRO1-like domain-containing protein(AT1G13310)** |
| | **At1g44130** | **Eukaryotic aspartyl protease family protein(AT1G44130)** |
| | GOTERM_BP_DIRECT | proteolysis, protein catabolic process, |
| | GOTERM_CC_DIRECT | extracellular region, Golgi apparatus, plant-type cell wall, |
| | GOTERM_MF_DIRECT | aspartic-type endopeptidase activity, |
| | **At5g10760** | **Eukaryotic aspartyl protease family protein(AT5G10760)** |
| | GOTERM_BP_DIRECT | proteolysis, systemic acquired resistance, protein catabolic process, |
| | GOTERM_CC_DIRECT | extracellular region, apoplast, |
| | GOTERM_MF_DIRECT | aspartic-type endopeptidase activity, |
| | **At5g10770** | **Eukaryotic aspartyl protease family protein(AT5G10770)** |
| | GOTERM_BP_DIRECT | proteolysis, protein catabolic process, |
| | GOTERM_CC_DIRECT | extracellular region, plasma membrane, anchored component of membrane, |
| | GOTERM_MF_DIRECT | DNA binding, aspartic-type endopeptidase activity, |
| | **At3g12700** | **Eukaryotic aspartyl protease family protein(NANA)** |
| | GOTERM_BP_DIRECT | carbohydrate metabolic process, proteolysis, chloroplast-nucleus signaling pathway, protein catabolic process, |
| | GOTERM_CC_DIRECT | extracellular region, nucleus, chloroplast, integral component of membrane, |
| | GOTERM_MF_DIRECT | aspartic-type endopeptidase activity, |
| | **At1g26380** | **FAD-binding Berberine family protein(AT1G26380)** |
| | GOTERM_BP_DIRECT | oxidation-reduction process, cellular response to hypoxia, |
| | GOTERM_CC_DIRECT | cytoplasm, endoplasmic reticulum, plasma membrane, |

| | | |
|---|---|---|
| | GOTERM_MF_DIRECT | electron carrier activity, oxidoreductase activity, acting on CH-OH group of donors, flavin adenine dinucleotide binding, |
| | **At1g26390** | **FAD-binding Berberine family protein(AT1G26390)** |
| | GOTERM_BP_DIRECT | oxidation-reduction process, |
| | GOTERM_CC_DIRECT | cytoplasm, |
| | GOTERM_MF_DIRECT | electron carrier activity, oxidoreductase activity, acting on CH-OH group of donors, flavin adenine dinucleotide binding, |
| | **At1g26410** | **FAD-binding Berberine family protein(AT1G26410)** |
| | GOTERM_BP_DIRECT | oxidation-reduction process, cellular response to hypoxia, |
| | GOTERM_CC_DIRECT | cytoplasm, |
| | GOTERM_MF_DIRECT | electron carrier activity, oxidoreductase activity, acting on CH-OH group of donors, flavin adenine dinucleotide binding, |
| | **At1g26420** | **FAD-binding Berberine family protein(AT1G26420)** |
| | GOTERM_BP_DIRECT | oxidation-reduction process, |
| | GOTERM_CC_DIRECT | cytoplasm, |
| | GOTERM_MF_DIRECT | electron carrier activity, oxidoreductase activity, acting on CH-OH group of donors, flavin adenine dinucleotide binding, |
| | **At1g30700** | **FAD-binding Berberine family protein(AT1G30700)** |
| | GOTERM_BP_DIRECT | oxidation-reduction process, |
| | GOTERM_CC_DIRECT | chloroplast, |
| | GOTERM_MF_DIRECT | electron carrier activity, oxidoreductase activity, acting on CH-OH group of donors, flavin adenine dinucleotide binding, |
| | **At4g20830** | **FAD-binding Berberine family protein(AT4G20830)** |
| | GOTERM_BP_DIRECT | response to oxidative stress, oxidation-reduction process, |
| | GOTERM_CC_DIRECT | mitochondrion, vacuole, cytosol, plasma membrane, plant-type cell wall, plasmodesma, chloroplast, anchored component of membrane, apoplast, |
| | GOTERM_MF_DIRECT | electron carrier activity, oxidoreductase activity, acting on CH-OH group of donors, flavin adenine dinucleotide binding, |
| | **At4g20860** | **FAD-binding Berberine family protein(AT4G20860)** |
| | GOTERM_BP_DIRECT | response to jasmonic acid, oxidation-reduction process, |
| | GOTERM_CC_DIRECT | cytoplasm, cytosol, plant-type cell wall, |
| | GOTERM_MF_DIRECT | electron carrier activity, oxidoreductase activity, acting on CH-OH group of donors, flavin adenine dinucleotide binding, |
| | **At5g44400** | **FAD-binding Berberine family protein(AT5G44400)** |
| | GOTERM_BP_DIRECT | oxidation-reduction process, |
| | GOTERM_CC_DIRECT | cell wall, cytoplasm, plasmodesma, |
| | GOTERM_MF_DIRECT | electron carrier activity, oxidoreductase activity, acting on CH-OH group of donors, flavin adenine dinucleotide binding, |
| | **At5g19260** | **FANTASTIC four-like protein (DUF3049)(FAF3)** |
| | **At3g06020** | **FANTASTIC four-like protein (DUF3049)(FAF4)** |

| | | |
|---|---|---|
| | GOTERM_BP_DIRECT | regulation of meristem growth, |
| | GOTERM_CC_DIRECT | nucleus, |
| | **At4g25100** | **Fe superoxide dismutase 1(FSD1)** |
| | GOTERM_BP_DIRECT | response to oxidative stress, circadian rhythm, response to light intensity, response to ozone, removal of superoxide radicals, response to cadmium ion, response to copper ion, oxidation-reduction process, |
| | GOTERM_CC_DIRECT | cytoplasm, mitochondrion, cytosol, plasma membrane, chloroplast, chloroplast stroma, thylakoid, chloroplast envelope, chloroplast membrane, |
| | GOTERM_MF_DIRECT | superoxide dismutase activity, copper ion binding, metal ion binding, |
| | **At1g12200** | **Flavin-binding monooxygenase family protein(FMO)** |
| | GOTERM_BP_DIRECT | defense response to fungus, oxidation-reduction process, |
| | GOTERM_CC_DIRECT | nucleus, |
| | GOTERM_MF_DIRECT | monooxygenase activity, N,N-dimethylaniline monooxygenase activity, flavin adenine dinucleotide binding, NADP binding, |
| | **At1g28600** | **GDSL-like Lipase/Acylhydrolase superfamily protein(AT1G28600)** |
| | GOTERM_BP_DIRECT | lipid catabolic process, |
| | GOTERM_CC_DIRECT | extracellular region, cell wall, |
| | GOTERM_MF_DIRECT | hydrolase activity, acting on ester bonds, carboxylic ester hydrolase activity, |
| | **At3g50400** | **GDSL-like Lipase/Acylhydrolase superfamily protein(AT3G50400)** |
| | GOTERM_BP_DIRECT | lipid catabolic process, |
| | GOTERM_CC_DIRECT | extracellular region, |
| | GOTERM_MF_DIRECT | hydrolase activity, acting on ester bonds, carboxylic ester hydrolase activity, |
| | **At2g18420** | **Gibberellin-regulated family protein(AT2G18420)** |
| | GOTERM_BP_DIRECT | response to gibberellin, gibberellic acid mediated signaling pathway, |
| | GOTERM_CC_DIRECT | extracellular region, |
| | **At5g48400** | **Glutamate receptor family protein(ATGLR1.2)** |
| | GOTERM_BP_DIRECT | calcium ion transport, cellular calcium ion homeostasis, response to light stimulus, calcium-mediated signaling, cellular response to amino acid stimulus, |
| | GOTERM_CC_DIRECT | extracellular region, intracellular, plasma membrane, integral component of membrane, |
| | GOTERM_MF_DIRECT | ionotropic glutamate receptor activity, intracellular ligand-gated ion channel activity, calcium channel activity, protein binding, glutamate receptor activity, |
| | **At5g44990** | **Glutathione S-transferase family protein(AT5G44990)** |
| | GOTERM_CC_DIRECT | cytoplasm, |
| | GOTERM_MF_DIRECT | transferase activity, |
| | **At4g27850** | **Glycine-rich protein family(AT4G27850)** |
| | GOTERM_CC_DIRECT | nucleus, integral component of membrane, |
| | **At4g39670** | **Glycolipid transfer protein (GLTP) family protein(AT4G39670)** |

| | | |
|---|---|---|
| | GOTERM_BP_DIRECT | glycolipid transport, |
| | GOTERM_CC_DIRECT | nucleus, cytoplasm, |
| | GOTERM_MF_DIRECT | glycolipid transporter activity, glycolipid binding, |
| | **At5g19240** | **Glycoprotein membrane precursor GPI-anchored(AT5G19240)** |
| | GOTERM_CC_DIRECT | plasma membrane, anchored component of membrane, |
| | **At4g19720** | **Glycosyl hydrolase family protein with chitinase insertion domain-containing protein(AT4G19720)** |
| | GOTERM_BP_DIRECT | carbohydrate metabolic process, |
| | GOTERM_CC_DIRECT | nucleus, |
| | GOTERM_MF_DIRECT | hydrolase activity, hydrolyzing O-glycosyl compounds, |
| | **At4g19810** | **Glycosyl hydrolase family protein with chitinase insertion domain-containing protein(ChiC)** |
| | GOTERM_BP_DIRECT | carbohydrate metabolic process, chitin catabolic process, response to salt stress, response to abscisic acid, response to jasmonic acid, |
| | GOTERM_CC_DIRECT | extracellular region, cell wall, |
| | GOTERM_MF_DIRECT | hydrolase activity, hydrolyzing O-glycosyl compounds, chitin binding, endochitinase activity, exochitinase activity, |
| | **At3g60140** | **Glycosyl hydrolase superfamily protein(DIN2)** |
| | GOTERM_BP_DIRECT | carbohydrate metabolic process, aging, response to salt stress, response to hormone, glucosinolate catabolic process, |
| | GOTERM_CC_DIRECT | extracellular region, |
| | GOTERM_MF_DIRECT | hydrolase activity, hydrolyzing O-glycosyl compounds, beta-glucosidase activity, scopolin beta-glucosidase activity, |
| | **At5g42830** | **HXXXD-type acyl-transferase family protein(AT5G42830)** |
| | GOTERM_CC_DIRECT | cytoplasm, cytosol, |
| | GOTERM_MF_DIRECT | transferase activity, transferase activity, transferring acyl groups other than amino-acyl groups, |
| | **At5g63560** | **HXXXD-type acyl-transferase family protein(FACT)** |
| | GOTERM_BP_DIRECT | alkyl caffeate ester biosynthetic process, |
| | GOTERM_CC_DIRECT | cytoplasm, |
| | GOTERM_MF_DIRECT | transferase activity, caffeoyl-CoA: alcohol caffeoyl transferase activity, |
| | **At1g35910** | **Haloacid dehalogenase-like hydrolase (HAD) superfamily protein(TPPD)** |
| | GOTERM_BP_DIRECT | trehalose biosynthetic process, response to osmotic stress, response to oxidative stress, response to salt stress, trehalose metabolism in response to stress, |
| | GOTERM_CC_DIRECT | cytoplasm, chloroplast, |
| | GOTERM_MF_DIRECT | trehalose-phosphatase activity, trehalase activity, |
| | **At1g09080** | **Heat shock protein 70 (Hsp 70) family protein(BIP3)** |
| | GOTERM_BP_DIRECT | transcription, DNA-templated, regulation of transcription, DNA-templated, protein folding, response to heat, pollen tube growth, |
| | GOTERM_CC_DIRECT | endoplasmic reticulum lumen, chloroplast, mediator complex, |

| | |
|---|---|
| GOTERM_MF_DIRECT | ATP binding, |
| **At5g02490** | **Heat shock protein 70 (Hsp 70) family protein(Hsp70-2)** |
| GOTERM_BP_DIRECT | transcription, DNA-templated, regulation of transcription, DNA-templated, protein folding, response to heat, response to virus, response to bacterium, response to cadmium ion, |
| GOTERM_CC_DIRECT | cell wall, nucleus, Golgi apparatus, cytosol, plasma membrane, |
| GOTERM_MF_DIRECT | protein binding, ATP binding, |
| **At5g52750** | **Heavy metal transport/detoxification superfamily protein(AT5G52750)** |
| GOTERM_BP_DIRECT | metal ion transport, cellular transition metal ion homeostasis, |
| GOTERM_CC_DIRECT | nucleus, cytoplasm, |
| GOTERM_MF_DIRECT | metal ion binding, transition metal ion binding, |
| **At2g38340** | **Integrase-type DNA-binding superfamily protein(DREB19)** |
| GOTERM_BP_DIRECT | transcription, DNA-templated, regulation of transcription, DNA-templated, response to water deprivation, response to salt stress, abscisic acid-activated signaling pathway, cellular response to heat, positive regulation of transcription, DNA-templated, |
| GOTERM_CC_DIRECT | nucleus, |
| GOTERM_MF_DIRECT | DNA binding, transcription factor activity, sequence-specific DNA binding, sequence-specific DNA binding, transcription regulatory region DNA binding, |
| **At2g47520** | **Integrase-type DNA-binding superfamily protein(ERF71)** |
| GOTERM_BP_DIRECT | transcription, DNA-templated, regulation of transcription, DNA-templated, ethylene-activated signaling pathway, response to anoxia, |
| GOTERM_CC_DIRECT | nucleus, |
| GOTERM_MF_DIRECT | DNA binding, transcription factor activity, sequence-specific DNA binding, |
| **At5g14760** | **L-aspartate oxidase(AO)** |
| GOTERM_BP_DIRECT | anaerobic respiration, NAD biosynthetic process, |
| GOTERM_CC_DIRECT | mitochondrion, chloroplast, |
| GOTERM_MF_DIRECT | L-aspartate oxidase activity, electron carrier activity, oxidoreductase activity, L-aspartate:fumarate oxidoreductase activity, |
| **At1g67100** | **LOB domain-containing protein 40(LBD40)** |
| GOTERM_BP_DIRECT | response to gibberellin, |
| **At3g54070** | **LOW protein: ankyrin repeat protein(AT3G54070)** |
| GOTERM_CC_DIRECT | integral component of membrane, |
| **At4g23610** | **Late embryogenesis abundant (LEA) hydroxyproline-rich glycoprotein family(AT4G23610)** |
| GOTERM_CC_DIRECT | integral component of membrane, |
| **At2g35980** | **Late embryogenesis abundant (LEA) hydroxyproline-rich glycoprotein family(YLS9)** |
| GOTERM_BP_DIRECT | leaf senescence, defense response to virus, response to other organism, |
| GOTERM_CC_DIRECT | mitochondrion, plasmodesma, chloroplast, anchored component of plasma membrane, |

| | | |
|---|---|---|
| | GOTERM_MF_DIRECT | signal transducer activity, |
| | **At1g53070** | **Legume lectin family protein(AT1G53070)** |
| | GOTERM_BP_DIRECT | response to karrikin, |
| | GOTERM_CC_DIRECT | cell wall, cytosol, plasma membrane, plant-type cell wall, apoplast, |
| | GOTERM_MF_DIRECT | carbohydrate binding, |
| | **At3g16530** | **Legume lectin family protein(AT3G16530)** |
| | GOTERM_BP_DIRECT | response to oomycetes, response to chitin, defense response to fungus, |
| | GOTERM_CC_DIRECT | cell wall, nucleus, plasma membrane, plant-type cell wall, apoplast, |
| | GOTERM_MF_DIRECT | carbohydrate binding, |
| | **At1g33600** | **Leucine-rich repeat (LRR) family protein(AT1G33600)** |
| | GOTERM_BP_DIRECT | signal transduction, |
| | GOTERM_CC_DIRECT | cell wall, plasma membrane, plant-type cell wall, membrane, |
| | **At1g49750** | **Leucine-rich repeat (LRR) family protein(AT1G49750)** |
| | GOTERM_CC_DIRECT | plasma membrane, chloroplast, |
| | **At5g66890** | **Leucine-rich repeat (LRR) family protein(AT5G66890)** |
| | GOTERM_BP_DIRECT | defense response, |
| | **At1g11130** | **Leucine-rich repeat protein kinase family protein(SUB)** |
| | GOTERM_BP_DIRECT | cell morphogenesis, signal transduction, positive regulation of atrichoblast fate specification, positive regulation of trichoblast fate specification, root meristem specification, leaf vascular tissue pattern formation, regulation of cell proliferation, leaf development, shoot system development, floral organ development, plant ovule development, |
| | GOTERM_CC_DIRECT | plasma membrane, plasmodesma, integral component of membrane, |
| | GOTERM_MF_DIRECT | protein kinase activity, receptor signaling protein serine/threonine kinase activity, ATP binding, |
| | **At1g01560** | **MAP kinase 11(MPK11)** |
| | GOTERM_BP_DIRECT | signal transduction, response to abscisic acid, |
| | GOTERM_CC_DIRECT | intracellular, nucleus, cytosol, |
| | GOTERM_MF_DIRECT | MAP kinase activity, ATP binding, kinase activity, |
| | **At1g33110** | **MATE efflux family protein(AT1G33110)** |
| | GOTERM_BP_DIRECT | drug transmembrane transport, |
| | GOTERM_CC_DIRECT | plasma membrane, membrane, integral component of membrane, |
| | GOTERM_MF_DIRECT | transporter activity, drug transmembrane transporter activity, antiporter activity, |
| | **At1g71140** | **MATE efflux family protein(AT1G71140)** |
| | GOTERM_BP_DIRECT | drug transmembrane transport, |

| | |
|---|---|
| GOTERM_CC_DIRECT | vacuolar membrane, plasma membrane, membrane, integral component of membrane, |
| GOTERM_MF_DIRECT | transporter activity, drug transmembrane transporter activity, antiporter activity, |
| **At2g04070** | **MATE efflux family protein(AT2G04070)** |
| GOTERM_BP_DIRECT | transport, drug transmembrane transport, |
| GOTERM_CC_DIRECT | plasma membrane, integral component of membrane, |
| GOTERM_MF_DIRECT | transporter activity, drug transmembrane transporter activity, antiporter activity, |
| **At3g21690** | **MATE efflux family protein(AT3G21690)** |
| GOTERM_BP_DIRECT | drug transmembrane transport, |
| GOTERM_CC_DIRECT | vacuole, vacuolar membrane, plasma membrane, membrane, integral component of membrane, |
| GOTERM_MF_DIRECT | transporter activity, drug transmembrane transporter activity, antiporter activity, |
| **At2g40460** | **Major facilitator superfamily protein(AT2G40460)** |
| GOTERM_BP_DIRECT | transport, oligopeptide transport, response to nematode, |
| GOTERM_CC_DIRECT | plasma membrane, membrane, integral component of membrane, |
| GOTERM_MF_DIRECT | transporter activity, |
| **At3g21670** | **Major facilitator superfamily protein(AT3G21670)** |
| GOTERM_BP_DIRECT | oligopeptide transport, nitrate assimilation, |
| GOTERM_CC_DIRECT | plasma membrane, membrane, integral component of membrane, |
| GOTERM_MF_DIRECT | transporter activity, symporter activity, |
| **At5g26340** | **Major facilitator superfamily protein(MSS1)** |
| GOTERM_BP_DIRECT | response to water deprivation, response to salt stress, response to abscisic acid, monosaccharide transport, glucose import, |
| GOTERM_CC_DIRECT | cytoplasm, plasma membrane, integral component of plasma membrane, plasmodesma, membrane, integral component of membrane, |
| GOTERM_MF_DIRECT | sugar:proton symporter activity, high-affinity hydrogen:glucose symporter activity, hexose:proton symporter activity, carbohydrate transmembrane transporter activity, monosaccharide transmembrane transporter activity, substrate-specific transmembrane transporter activity, |
| **At1g67070** | **Mannose-6-phosphate isomerase, type I(DIN9)** |
| GOTERM_BP_DIRECT | cell wall mannoprotein biosynthetic process, carbohydrate metabolic process, protein glycosylation, GDP-mannose biosynthetic process, response to absence of light, response to sucrose, response to zinc ion, L-ascorbic acid biosynthetic process, response to L-ascorbic acid, response to DDT, response to cadmium ion, |
| GOTERM_CC_DIRECT | cytoplasm, |
| GOTERM_MF_DIRECT | mannose-6-phosphate isomerase activity, zinc ion binding, |
| **At1g24140** | **Matrixin family protein(AT1G24140)** |
| GOTERM_BP_DIRECT | proteolysis, |
| GOTERM_CC_DIRECT | extracellular region, plasma membrane, extracellular matrix, anchored component of membrane, |
| GOTERM_MF_DIRECT | metalloendopeptidase activity, zinc ion binding, |

| | | |
|---|---|---|
| | **At5g38710** | **Methylenetetrahydrofolate reductase family protein(AT5G38710)** |
| | GOTERM_BP_DIRECT | glutamate biosynthetic process, proline catabolic process, response to osmotic stress, response to water deprivation, proline catabolic process to glutamate, oxidation-reduction process, |
| | GOTERM_CC_DIRECT | mitochondrion, |
| | GOTERM_MF_DIRECT | proline dehydrogenase activity, |
| | **At3g16150** | **N-terminal nucleophile aminohydrolases (Ntn hydrolases) superfamily protein(ASPGB1)** |
| | GOTERM_BP_DIRECT | proteolysis, glycoprotein catabolic process, asparagine catabolic process via L-aspartate, protein maturation, |
| | GOTERM_CC_DIRECT | cytoplasm, |
| | GOTERM_MF_DIRECT | asparaginase activity, beta-aspartyl-peptidase activity, hydrolase activity, |
| | **At3g15500** | **NAC domain containing protein 3(NAC3)** |
| | GOTERM_BP_DIRECT | transcription, DNA-templated, regulation of transcription, DNA-templated, multicellular organism development, response to water deprivation, jasmonic acid mediated signaling pathway, |
| | GOTERM_CC_DIRECT | nucleus, |
| | GOTERM_MF_DIRECT | DNA binding, transcription factor activity, sequence-specific DNA binding, protein binding, |
| | **At1g77450** | **NAC domain containing protein 32(NAC032)** |
| | GOTERM_BP_DIRECT | transcription, DNA-templated, regulation of transcription, DNA-templated, multicellular organism development, |
| | GOTERM_CC_DIRECT | nucleus, |
| | GOTERM_MF_DIRECT | DNA binding, transcription factor activity, sequence-specific DNA binding, |
| | **At2g17040** | **NAC domain containing protein 36(NAC036)** |
| | GOTERM_BP_DIRECT | transcription, DNA-templated, regulation of transcription, DNA-templated, multicellular organism development, leaf morphogenesis, response to chitin, negative regulation of cell size, inflorescence morphogenesis, |
| | GOTERM_CC_DIRECT | nucleus, |
| | GOTERM_MF_DIRECT | DNA binding, transcription factor activity, sequence-specific DNA binding, |
| | **At2g24430** | **NAC domain containing protein 38(NAC038)** |
| | GOTERM_BP_DIRECT | transcription, DNA-templated, regulation of transcription, DNA-templated, multicellular organism development, |
| | GOTERM_CC_DIRECT | nucleus, |
| | GOTERM_MF_DIRECT | DNA binding, transcription factor activity, sequence-specific DNA binding, |
| | **At2g43000** | **NAC domain containing protein 42(NAC042)** |
| | GOTERM_BP_DIRECT | trehalose biosynthetic process, transcription, DNA-templated, regulation of transcription, DNA-templated, proline biosynthetic process, multicellular organism development, anthocyanin-containing compound biosynthetic process, camalexin biosynthetic process, leaf senescence, hyperosmotic salinity response, negative regulation of leaf senescence, |
| | GOTERM_CC_DIRECT | nucleus, |
| | GOTERM_MF_DIRECT | DNA binding, transcription factor activity, sequence-specific DNA binding, |
| | **At3g04070** | **NAC domain containing protein 47(NAC047)** |

| | | |
|---|---|---|
| | GOTERM_BP_DIRECT | transcription, DNA-templated, regulation of transcription, DNA-templated, multicellular organism development, |
| | GOTERM_CC_DIRECT | nucleus, |
| | GOTERM_MF_DIRECT | DNA binding, transcription factor activity, sequence-specific DNA binding, sequence-specific DNA binding, |
| | **At5g39610** | **NAC domain containing protein 6(NAC6)** |
| | GOTERM_BP_DIRECT | transcription, DNA-templated, apoptotic process, response to oxidative stress, multicellular organism development, response to salt stress, response to ethylene, response to auxin, response to abscisic acid, regulation of seed germination, leaf senescence, regulation of gene expression, response to hydrogen peroxide, positive regulation of programmed cell death, lateral root development, positive regulation of sequence-specific DNA binding transcription factor activity, stress-induced premature senescence, positive regulation of leaf senescence, response to salt, positive regulation of age-related resistance, |
| | GOTERM_CC_DIRECT | nucleus, |
| | GOTERM_MF_DIRECT | transcription factor activity, sequence-specific DNA binding, protein binding, protein homodimerization activity, sequence-specific DNA binding, protein heterodimerization activity, |
| | **At1g64590** | **NAD(P)-binding Rossmann-fold superfamily protein(AT1G64590)** |
| | GOTERM_MF_DIRECT | oxidoreductase activity, |
| | **At2g29150** | **NAD(P)-binding Rossmann-fold superfamily protein(AT2G29150)** |
| | GOTERM_BP_DIRECT | oxidation-reduction process, |
| | GOTERM_MF_DIRECT | oxidoreductase activity, |
| | **At2g30670** | **NAD(P)-binding Rossmann-fold superfamily protein(AT2G30670)** |
| | GOTERM_BP_DIRECT | oxidation-reduction process, |
| | GOTERM_MF_DIRECT | oxidoreductase activity, |
| | **At1g60730** | **NAD(P)-linked oxidoreductase superfamily protein(AT1G60730)** |
| | GOTERM_BP_DIRECT | oxidation-reduction process, |
| | GOTERM_CC_DIRECT | cytoplasm, cytosol, |
| | GOTERM_MF_DIRECT | aldo-keto reductase (NADP) activity, oxidoreductase activity, |
| | **At1g60750** | **NAD(P)-linked oxidoreductase superfamily protein(AT1G60750)** |
| | GOTERM_BP_DIRECT | oxidation-reduction process, |
| | GOTERM_CC_DIRECT | cytoplasm, |
| | GOTERM_MF_DIRECT | oxidoreductase activity, |
| | **At3g22550** | **NAD(P)H-quinone oxidoreductase subunit, putative (DUF581)(AT3G22550)** |
| | GOTERM_CC_DIRECT | nucleus, |
| | **At1g02450** | **NIM1-interacting 1(NIMIN1)** |
| | GOTERM_BP_DIRECT | regulation of systemic acquired resistance, defense response to bacterium, negative regulation of transcription, DNA-templated, |
| | GOTERM_CC_DIRECT | nucleus, |
| | GOTERM_MF_DIRECT | protein binding, |

| | |
|---|---|
| **At5g45110** | **NPR1-like protein 3(NPR3)** |
| GOTERM_BP_DIRECT | systemic acquired resistance, defense response to bacterium, incompatible interaction, defense response to fungus, incompatible interaction, regulation of proteolysis, protein ubiquitination involved in ubiquitin-dependent protein catabolic process, proteasome-mediated ubiquitin-dependent protein catabolic process, effector dependent induction by symbiont of host immune response, regulation of jasmonic acid mediated signaling pathway, |
| GOTERM_CC_DIRECT | nucleus, cytoplasm, SCF ubiquitin ligase complex, |
| GOTERM_MF_DIRECT | protein binding, ubiquitin protein ligase binding, salicylic acid binding, |
| **At3g14770** | **Nodulin MtN3 family protein(SWEET2)** |
| GOTERM_BP_DIRECT | carbohydrate transport, |
| GOTERM_CC_DIRECT | plasma membrane, integral component of plasma membrane, membrane, integral component of membrane, |
| GOTERM_MF_DIRECT | sugar transmembrane transporter activity, |
| **At5g44820** | **Nucleotide-diphospho-sugar transferase family protein(AT5G44820)** |
| GOTERM_CC_DIRECT | chloroplast, |
| GOTERM_MF_DIRECT | transferase activity, |
| **At1g57980** | **Nucleotide-sugar transporter family protein(AT1G57980)** |
| GOTERM_BP_DIRECT | transport, |
| GOTERM_CC_DIRECT | integral component of membrane, |
| GOTERM_MF_DIRECT | transporter activity, purine nucleobase transmembrane transporter activity, |
| **At1g33030** | **O-methyltransferase family protein(AT1G33030)** |
| GOTERM_BP_DIRECT | lignin biosynthetic process, aromatic compound biosynthetic process, methylation, |
| GOTERM_CC_DIRECT | cytosol, |
| GOTERM_MF_DIRECT | O-methyltransferase activity, S-adenosylmethionine-dependent methyltransferase activity, protein dimerization activity, |
| **At1g43910** | **P-loop containing nucleoside triphosphate hydrolases superfamily protein(AT1G43910)** |
| GOTERM_BP_DIRECT | response to abscisic acid, cellular response to potassium ion starvation, |
| GOTERM_CC_DIRECT | endoplasmic reticulum, Golgi apparatus, plasma membrane, plasmodesma, integral component of membrane, |
| GOTERM_MF_DIRECT | ATP binding, ATPase activity, |
| **At3g28510** | **P-loop containing nucleoside triphosphate hydrolases superfamily protein(AT3G28510)** |
| GOTERM_CC_DIRECT | endoplasmic reticulum, plasma membrane, integral component of membrane, |
| GOTERM_MF_DIRECT | ATP binding, ATPase activity, |
| **At3g28580** | **P-loop containing nucleoside triphosphate hydrolases superfamily protein(AT3G28580)** |
| GOTERM_BP_DIRECT | response to singlet oxygen, response to abscisic acid, |
| GOTERM_CC_DIRECT | endoplasmic reticulum, plasma membrane, integral component of membrane, |
| GOTERM_MF_DIRECT | ATP binding, ATPase activity, |

| | |
|---|---|
| **At5g17760** | **P-loop containing nucleoside triphosphate hydrolases superfamily protein(AT5G17760)** |
| GOTERM_CC_DIRECT | plasma membrane, integral component of membrane, |
| GOTERM_MF_DIRECT | ATP binding, ATPase activity, |
| **At5g52390** | **PAR1 protein(AT5G52390)** |
| **At1g72520** | **PLAT/LH2 domain-containing lipoxygenase family protein(LOX4)** |
| GOTERM_BP_DIRECT | fatty acid biosynthetic process, defense response, pollen development, response to wounding, response to bacterium, jasmonic acid biosynthetic process, anther dehiscence, response to ozone, oxylipin biosynthetic process, lipid oxidation, growth, anther development, stamen filament development, |
| GOTERM_CC_DIRECT | chloroplast, |
| GOTERM_MF_DIRECT | linoleate 13S-lipoxygenase activity, metal ion binding, |
| **At3g29240** | **PPR containing protein (DUF179)(AT3G29240)** |
| GOTERM_CC_DIRECT | chloroplast, |
| **At1g14550** | **Peroxidase superfamily protein(AT1G14550)** |
| GOTERM_BP_DIRECT | response to oxidative stress, hydrogen peroxide catabolic process, oxidation-reduction process, cellular response to hypoxia, |
| GOTERM_CC_DIRECT | extracellular region, |
| GOTERM_MF_DIRECT | peroxidase activity, heme binding, metal ion binding, |
| **At2g18140** | **Peroxidase superfamily protein(AT2G18140)** |
| GOTERM_BP_DIRECT | response to oxidative stress, hydrogen peroxide catabolic process, oxidation-reduction process, |
| GOTERM_CC_DIRECT | extracellular region, cell wall, |
| GOTERM_MF_DIRECT | peroxidase activity, heme binding, metal ion binding, |
| **At2g35380** | **Peroxidase superfamily protein(AT2G35380)** |
| GOTERM_BP_DIRECT | response to oxidative stress, hydrogen peroxide catabolic process, oxidation-reduction process, |
| GOTERM_CC_DIRECT | extracellular region, |
| GOTERM_MF_DIRECT | peroxidase activity, heme binding, metal ion binding, |
| **At4g08780** | **Peroxidase superfamily protein(AT4G08780)** |
| GOTERM_BP_DIRECT | response to oxidative stress, hydrogen peroxide catabolic process, oxidation-reduction process, |
| GOTERM_CC_DIRECT | extracellular region, vacuole, |
| GOTERM_MF_DIRECT | peroxidase activity, heme binding, metal ion binding, |
| **At4g36430** | **Peroxidase superfamily protein(AT4G36430)** |
| GOTERM_BP_DIRECT | response to oxidative stress, hydrogen peroxide catabolic process, response to other organism, oxidation-reduction process, |
| GOTERM_CC_DIRECT | extracellular region, cell wall, |
| GOTERM_MF_DIRECT | peroxidase activity, heme binding, metal ion binding, |
| **At5g06730** | **Peroxidase superfamily protein(AT5G06730)** |

| | | |
|---|---|---|
| | GOTERM_BP_DIRECT | response to oxidative stress, hydrogen peroxide catabolic process, oxidation-reduction process, |
| | GOTERM_CC_DIRECT | extracellular region, vacuolar membrane, |
| | GOTERM_MF_DIRECT | peroxidase activity, heme binding, metal ion binding, |
| | **At5g14130** | **Peroxidase superfamily protein(AT5G14130)** |
| | GOTERM_BP_DIRECT | response to oxidative stress, hydrogen peroxide catabolic process, oxidation-reduction process, |
| | GOTERM_CC_DIRECT | extracellular region, |
| | GOTERM_MF_DIRECT | peroxidase activity, heme binding, metal ion binding, |
| | **At5g19880** | **Peroxidase superfamily protein(AT5G19880)** |
| | GOTERM_BP_DIRECT | response to oxidative stress, response to virus, response to ethylene, hydrogen peroxide catabolic process, oxidation-reduction process, |
| | GOTERM_CC_DIRECT | extracellular region, |
| | GOTERM_MF_DIRECT | peroxidase activity, heme binding, metal ion binding, |
| | **At5g39580** | **Peroxidase superfamily protein(AT5G39580)** |
| | GOTERM_BP_DIRECT | response to oxidative stress, plant-type cell wall organization, hydrogen peroxide catabolic process, defense response to fungus, oxidation-reduction process, |
| | GOTERM_CC_DIRECT | extracellular region, Golgi apparatus, plant-type cell wall, |
| | GOTERM_MF_DIRECT | peroxidase activity, heme binding, metal ion binding, |
| | **At5g64110** | **Peroxidase superfamily protein(AT5G64110)** |
| | GOTERM_BP_DIRECT | response to oxidative stress, plant-type cell wall organization, hydrogen peroxide catabolic process, oxidation-reduction process, |
| | GOTERM_CC_DIRECT | extracellular region, plant-type cell wall, membrane, |
| | GOTERM_MF_DIRECT | peroxidase activity, heme binding, metal ion binding, |
| | **At5g64120** | **Peroxidase superfamily protein(AT5G64120)** |
| | GOTERM_BP_DIRECT | response to oxidative stress, plant-type cell wall organization, lignin metabolic process, hydrogen peroxide catabolic process, respiratory burst, rhythmic process, defense response to fungus, oxidation-reduction process, |
| | GOTERM_CC_DIRECT | extracellular region, cell wall, plant-type cell wall, membrane, apoplast, |
| | GOTERM_MF_DIRECT | peroxidase activity, heme binding, metal ion binding, |
| | **At3g01420** | **Peroxidase superfamily protein(DOX1)** |
| | GOTERM_BP_DIRECT | fatty acid alpha-oxidation, lipid metabolic process, fatty acid biosynthetic process, response to oxidative stress, cell death, plant-type hypersensitive response, systemic acquired resistance, response to abscisic acid, response to salicylic acid, oxylipin biosynthetic process, cellular response to reactive oxygen species, defense response to bacterium, defense response to fungus, response to other organism, cellular response to salicylic acid stimulus, cellular response to nitric oxide, (R)-2-hydroxy-alpha-linolenic acid biosynthetic process, |
| | GOTERM_CC_DIRECT | extracellular region, monolayer-surrounded lipid storage body, |
| | GOTERM_MF_DIRECT | peroxidase activity, oxidoreductase activity, acting on single donors with incorporation of molecular oxygen, incorporation of two atoms of oxygen, heme binding, metal ion binding, |
| | **At1g14540** | **Peroxidase superfamily protein(PER4)** |
| | GOTERM_BP_DIRECT | response to oxidative stress, hydrogen peroxide catabolic process, oxidation-reduction process, cellular response to hypoxia, |

| | |
|---|---|
| GOTERM_CC_DIRECT | extracellular region, |
| GOTERM_MF_DIRECT | peroxidase activity, heme binding, metal ion binding, |
| **At5g05340** | **Peroxidase superfamily protein(PRX52)** |
| GOTERM_BP_DIRECT | response to oxidative stress, lignin biosynthetic process, xylem development, hydrogen peroxide catabolic process, oxidation-reduction process, positive regulation of syringal lignin biosynthetic process, |
| GOTERM_CC_DIRECT | extracellular region, cell wall, Golgi apparatus, cytosol, apoplast, |
| GOTERM_MF_DIRECT | peroxidase activity, protein binding, heme binding, metal ion binding, |
| **At1g35140** | **Phosphate-responsive 1 family protein(PHI-1)** |
| GOTERM_BP_DIRECT | response to hypoxia, growth, |
| GOTERM_CC_DIRECT | extracellular region, extracellular space, Golgi apparatus, plant-type cell wall, apoplast, |
| **At4g28940** | **Phosphorylase superfamily protein(AT4G28940)** |
| GOTERM_BP_DIRECT | nucleoside metabolic process, |
| GOTERM_CC_DIRECT | plasma membrane, |
| GOTERM_MF_DIRECT | catalytic activity, |
| **At5g46950** | **Plant invertase/pectin methylesterase inhibitor superfamily protein(AT5G46950)** |
| GOTERM_BP_DIRECT | negative regulation of catalytic activity, |
| GOTERM_CC_DIRECT | chloroplast, cell periphery, |
| GOTERM_MF_DIRECT | pectinesterase activity, pectinesterase inhibitor activity, |
| **At4g23680** | **Polyketide cyclase/dehydrase and lipid transport superfamily protein(AT4G23680)** |
| GOTERM_BP_DIRECT | defense response, response to biotic stimulus, |
| **At3g26470** | **Powdery mildew resistance protein, RPW8 domain-containing protein(AT3G26470)** |
| GOTERM_CC_DIRECT | mitochondrion, |
| **At1g68690** | **Protein kinase superfamily protein(PERK9)** |
| GOTERM_BP_DIRECT | protein phosphorylation, |
| GOTERM_CC_DIRECT | plasma membrane, integral component of membrane, |
| GOTERM_MF_DIRECT | protein serine/threonine kinase activity, ATP binding, protein kinase binding, |
| **At4g32950** | **Protein phosphatase 2C family protein(AT4G32950)** |
| GOTERM_BP_DIRECT | protein dephosphorylation, |
| GOTERM_MF_DIRECT | protein serine/threonine phosphatase activity, metal ion binding, |
| **At4g35190** | **Putative lysine decarboxylase family protein(LOG5)** |
| GOTERM_BP_DIRECT | cytokinin biosynthetic process, |
| GOTERM_CC_DIRECT | nucleus, cytosol, |

| | | |
|---|---|---|
| | GOTERM_MF_DIRECT | hydrolase activity, hydrolyzing N-glycosyl compounds, |
| | **At3g18250** | **Putative membrane lipoprotein(AT3G18250)** |
| | GOTERM_CC_DIRECT | endoplasmic reticulum, integral component of membrane, |
| | **At1g05880** | **RING/U-box superfamily protein(ARI12)** |
| | GOTERM_BP_DIRECT | protein polyubiquitination, response to UV-B, positive regulation of proteasomal ubiquitin-dependent protein catabolic process, protein ubiquitination involved in ubiquitin-dependent protein catabolic process, cellular response to hypoxia, |
| | GOTERM_CC_DIRECT | ubiquitin ligase complex, nucleus, cytoplasm, |
| | GOTERM_MF_DIRECT | ligase activity, ubiquitin conjugating enzyme binding, metal ion binding, ubiquitin protein ligase activity, |
| | **At2g42360** | **RING/U-box superfamily protein(AT2G42360)** |
| | GOTERM_BP_DIRECT | protein ubiquitination, proteasome-mediated ubiquitin-dependent protein catabolic process, |
| | GOTERM_CC_DIRECT | nucleus, integral component of membrane, |
| | GOTERM_MF_DIRECT | ubiquitin-protein transferase activity, zinc ion binding, ligase activity, ubiquitin protein ligase activity, |
| | **At3g61390** | **RING/U-box superfamily protein(AT3G61390)** |
| | GOTERM_BP_DIRECT | protein ubiquitination, |
| | GOTERM_CC_DIRECT | nucleus, |
| | GOTERM_MF_DIRECT | ubiquitin-protein transferase activity, ligase activity, |
| | **At5g10380** | **RING/U-box superfamily protein(RING1)** |
| | GOTERM_BP_DIRECT | response to molecule of fungal origin, apoptotic process, defense response, response to bacterium, response to chitin, programmed cell death, protein ubiquitination, positive regulation of programmed cell death, protein autoubiquitination, |
| | GOTERM_CC_DIRECT | extracellular region, plasma membrane, integral component of membrane, |
| | GOTERM_MF_DIRECT | ubiquitin-protein transferase activity, zinc ion binding, ligase activity, |
| | **At3g48450** | **RPM1-interacting protein 4 (RIN4) family protein(AT3G48450)** |
| | GOTERM_BP_DIRECT | response to nitrate, |
| | GOTERM_CC_DIRECT | nucleus, |
| | **At5g41280** | **Receptor-like protein kinase-related family protein(AT5G41280)** |
| | GOTERM_CC_DIRECT | plasma membrane, anchored component of membrane, anchored component of plasma membrane, |
| | **At1g13340** | **Regulator of Vps4 activity in the MVB pathway protein(AT1G13340)** |
| | GOTERM_BP_DIRECT | response to oxidative stress, protein transport, |
| | GOTERM_CC_DIRECT | cytoplasm, |
| | **At2g17850** | **Rhodanese/Cell cycle control phosphatase superfamily protein(AT2G17850)** |
| | GOTERM_CC_DIRECT | chloroplast, |
| | **At4g35770** | **Rhodanese/Cell cycle control phosphatase superfamily protein(SEN1)** |

| | | |
|---|---|---|
| | GOTERM_BP_DIRECT | response to oxidative stress, aging, response to wounding, response to jasmonic acid, |
| | GOTERM_CC_DIRECT | chloroplast, thylakoid, |
| | **At5g38910** | **RmlC-like cupins superfamily protein(AT5G38910)** |
| | GOTERM_BP_DIRECT | oxalate metabolic process, |
| | GOTERM_CC_DIRECT | extracellular region, cell wall, apoplast, |
| | GOTERM_MF_DIRECT | manganese ion binding, nutrient reservoir activity, oxalate decarboxylase activity, |
| | **At5g39110** | **RmlC-like cupins superfamily protein(AT5G39110)** |
| | GOTERM_BP_DIRECT | oxalate metabolic process, |
| | GOTERM_CC_DIRECT | extracellular region, cell wall, apoplast, |
| | GOTERM_MF_DIRECT | manganese ion binding, nutrient reservoir activity, oxalate decarboxylase activity, |
| | **At5g39130** | **RmlC-like cupins superfamily protein(AT5G39130)** |
| | GOTERM_BP_DIRECT | oxalate metabolic process, |
| | GOTERM_CC_DIRECT | extracellular region, cell wall, apoplast, |
| | GOTERM_MF_DIRECT | manganese ion binding, nutrient reservoir activity, oxalate decarboxylase activity, |
| | **At2g41380** | **S-adenosyl-L-methionine-dependent methyltransferases superfamily protein(AT2G41380)** |
| | GOTERM_BP_DIRECT | methylation, response to cadmium ion, |
| | GOTERM_CC_DIRECT | mitochondrion, endosome, vacuolar membrane, trans-Golgi network, cytosol, |
| | GOTERM_MF_DIRECT | S-adenosylmethionine-dependent methyltransferase activity, |
| | **At3g54150** | **S-adenosyl-L-methionine-dependent methyltransferases superfamily protein(AT3G54150)** |
| | GOTERM_BP_DIRECT | pollen germination, pollen exine formation, methylation, |
| | GOTERM_CC_DIRECT | nucleus, endosome, vacuolar membrane, trans-Golgi network, cytosol, |
| | GOTERM_MF_DIRECT | S-adenosylmethionine-dependent methyltransferase activity, |
| | **At4g22530** | **S-adenosyl-L-methionine-dependent methyltransferases superfamily protein(AT4G22530)** |
| | GOTERM_BP_DIRECT | methylation, |
| | GOTERM_CC_DIRECT | nucleus, cytoplasm, endosome, vacuolar membrane, trans-Golgi network, cytosol, |
| | GOTERM_MF_DIRECT | S-adenosylmethionine-dependent methyltransferase activity, |
| | **At4g26460** | **S-adenosyl-L-methionine-dependent methyltransferases superfamily protein(AT4G26460)** |
| | GOTERM_BP_DIRECT | methylation, |
| | GOTERM_CC_DIRECT | nucleus, |
| | GOTERM_MF_DIRECT | methyltransferase activity, |
| | **At1g11330** | **S-locus lectin protein kinase family protein(AT1G11330)** |

| | | |
|---|---|---|
| | GOTERM_BP_DIRECT | protein phosphorylation, innate immune response, recognition of pollen, |
| | GOTERM_CC_DIRECT | plasma membrane, plasmodesma, integral component of membrane, |
| | GOTERM_MF_DIRECT | protein serine/threonine kinase activity, calmodulin binding, ATP binding, kinase activity, carbohydrate binding, ubiquitin protein ligase binding, |
| | **At1g61500** | **S-locus lectin protein kinase family protein(AT1G61500)** |
| | GOTERM_BP_DIRECT | protein phosphorylation, innate immune response, recognition of pollen, |
| | GOTERM_CC_DIRECT | plasma membrane, plasmodesma, integral component of membrane, |
| | GOTERM_MF_DIRECT | protein serine/threonine kinase activity, calmodulin binding, ATP binding, kinase activity, carbohydrate binding, ubiquitin protein ligase binding, |
| | **At1g61550** | **S-locus lectin protein kinase family protein(AT1G61550)** |
| | GOTERM_BP_DIRECT | protein phosphorylation, innate immune response, recognition of pollen, |
| | GOTERM_CC_DIRECT | plasma membrane, plasmodesma, integral component of membrane, |
| | GOTERM_MF_DIRECT | protein serine/threonine kinase activity, calmodulin binding, ATP binding, kinase activity, carbohydrate binding, ubiquitin protein ligase binding, |
| | **At5g37690** | **SGNH hydrolase-type esterase superfamily protein(AT5G37690)** |
| | GOTERM_BP_DIRECT | lipid catabolic process, |
| | GOTERM_CC_DIRECT | extracellular region, |
| | GOTERM_MF_DIRECT | lipase activity, hydrolase activity, acting on ester bonds, |
| | **At5g25250** | **SPFH/Band 7/PHB domain-containing membrane-associated protein family(FLOT1)** |
| | GOTERM_BP_DIRECT | endocytosis, membrane invagination, cellular response to hypoxia, |
| | GOTERM_CC_DIRECT | endosome, vacuole, vacuolar membrane, plasma membrane, caveola, plasmodesma, |
| | **At5g20150** | **SPX domain-containing protein 1(SPX1)** |
| | GOTERM_BP_DIRECT | cellular response to phosphate starvation, positive regulation of cellular response to phosphate starvation, |
| | GOTERM_CC_DIRECT | nucleus, endoplasmic reticulum, |
| | **At2g38870** | **Serine protease inhibitor, potato inhibitor I-type family protein(AT2G38870)** |
| | GOTERM_BP_DIRECT | proteolysis, response to wounding, defense response to fungus, |
| | GOTERM_CC_DIRECT | extracellular region, cell wall, |
| | GOTERM_MF_DIRECT | serine-type endopeptidase inhibitor activity, peptidase activity, |
| | **At1g32940** | **Subtilase family protein(SBT3.5)** |
| | GOTERM_BP_DIRECT | proteolysis, metabolic process, regulation of growth, |
| | GOTERM_CC_DIRECT | extracellular region, cell wall, apoplast, |
| | GOTERM_MF_DIRECT | serine-type endopeptidase activity, |
| | **At4g01390** | **TRAF-like family protein(AT4G01390)** |
| | GOTERM_CC_DIRECT | extracellular region, nucleus, cytoplasm, |

| | | |
|---|---|---|
| | **At5g24600** | **TRP-like ion channel protein (Protein of unknown function, DUF599)(AT5G24600)** |
| | GOTERM_CC_DIRECT | mitochondrion, plasma membrane, integral component of membrane, |
| | **At5g38900** | **Thioredoxin superfamily protein(AT5G38900)** |
| | GOTERM_BP_DIRECT | defense response to fungus, incompatible interaction, |
| | GOTERM_CC_DIRECT | cytosol, |
| | GOTERM_MF_DIRECT | protein disulfide oxidoreductase activity, |
| | **At1g57630** | **Toll-Interleukin-Resistance (TIR) domain family protein(AT1G57630)** |
| | GOTERM_BP_DIRECT | defense response, signal transduction, cellular response to hypoxia, |
| | GOTERM_CC_DIRECT | chloroplast, |
| | **At1g72910** | **Toll-Interleukin-Resistance (TIR) domain-containing protein(AT1G72910)** |
| | GOTERM_BP_DIRECT | defense response, signal transduction, |
| | GOTERM_CC_DIRECT | cytoplasm, |
| | GOTERM_MF_DIRECT | ADP binding, |
| | **At1g72940** | **Toll-Interleukin-Resistance (TIR) domain-containing protein(AT1G72940)** |
| | GOTERM_BP_DIRECT | defense response, signal transduction, |
| | GOTERM_CC_DIRECT | cytoplasm, |
| | GOTERM_MF_DIRECT | ADP binding, |
| | **At3g02800** | **Tyrosine phosphatase family protein(PFA-DSP3)** |
| | GOTERM_CC_DIRECT | nucleus, cytoplasm, |
| | GOTERM_MF_DIRECT | phosphoprotein phosphatase activity, protein tyrosine phosphatase activity, phosphatase activity, |
| | **At2g30140** | **UDP-Glycosyltransferase superfamily protein(UGT87A2)** |
| | GOTERM_BP_DIRECT | metabolic process, flavonoid biosynthetic process, regulation of flower development, flavonoid glucuronidation, |
| | GOTERM_CC_DIRECT | nucleus, cytoplasm, cytosol, |
| | GOTERM_MF_DIRECT | UDP-glycosyltransferase activity, transferase activity, transferring glycosyl groups, transferase activity, transferring hexosyl groups, quercetin 3-O-glucosyltransferase activity, quercetin 7-O-glucosyltransferase activity, |
| | **At4g34131** | **UDP-glucosyl transferase 73B3(UGT73B3)** |
| | GOTERM_BP_DIRECT | defense response, metabolic process, flavonoid biosynthetic process, response to other organism, flavonoid glucuronidation, |
| | GOTERM_CC_DIRECT | intracellular membrane-bounded organelle, |
| | GOTERM_MF_DIRECT | UDP-glycosyltransferase activity, abscisic acid glucosyltransferase activity, transferase activity, transferring hexosyl groups, UDP-glucosyltransferase activity, flavonol 3-O-glucosyltransferase activity, quercetin 3-O-glucosyltransferase activity, quercetin 7-O-glucosyltransferase activity, daphnetin 3-O-glucosyltransferase activity, |
| | **At3g53150** | **UDP-glucosyl transferase 73D1(UGT73D1)** |
| | GOTERM_BP_DIRECT | metabolic process, flavonoid biosynthetic process, flavonoid glucuronidation, |

| | |
|---|---|
| GOTERM_CC_DIRECT | intracellular membrane-bounded organelle, |
| GOTERM_MF_DIRECT | UDP-glycosyltransferase activity, transferase activity, transferring hexosyl groups, quercetin 3-O-glucosyltransferase activity, quercetin 7-O-glucosyltransferase activity, |
| **At2g26480** | **UDP-glucosyl transferase 76D1(UGT76D1)** |
| GOTERM_BP_DIRECT | metabolic process, flavonoid biosynthetic process, flavonoid glucuronidation, |
| GOTERM_CC_DIRECT | nucleus, intracellular membrane-bounded organelle, |
| GOTERM_MF_DIRECT | UDP-glycosyltransferase activity, transferase activity, transferring glycosyl groups, transferase activity, transferring hexosyl groups, quercetin 3-O-glucosyltransferase activity, quercetin 7-O-glucosyltransferase activity, |
| **At2g15490** | **UDP-glycosyltransferase 73B4(UGT73B4)** |
| GOTERM_BP_DIRECT | metabolic process, response to toxic substance, flavonoid biosynthetic process, response to other organism, flavonoid glucuronidation, |
| GOTERM_CC_DIRECT | cytosol, intracellular membrane-bounded organelle, |
| GOTERM_MF_DIRECT | UDP-glycosyltransferase activity, transferase activity, transferring glycosyl groups, transferase activity, transferring hexosyl groups, UDP-glucosyltransferase activity, flavonol 3-O-glucosyltransferase activity, quercetin 3-O-glucosyltransferase activity, quercetin 7-O-glucosyltransferase activity, daphnetin 3-O-glucosyltransferase activity, |
| **At3g46110** | **UPSTREAM OF FLC-like protein (DUF966)(AT3G46110)** |
| GOTERM_CC_DIRECT | nucleus, plasma membrane, |
| **At1g58420** | **Uncharacterized conserved protein UCP031279(AT1G58420)** |
| GOTERM_CC_DIRECT | mitochondrion, |
| **At4g15610** | **Uncharacterized protein family (UPF0497)(AT4G15610)** |
| GOTERM_CC_DIRECT | endosome, Golgi apparatus, trans-Golgi network, plasma membrane, integral component of membrane, |
| **At2g22880** | **VQ motif-containing protein(AT2G22880)** |
| GOTERM_BP_DIRECT | response to UV-B, |
| GOTERM_CC_DIRECT | nucleus, |
| **At4g37710** | **VQ motif-containing protein(AT4G37710)** |
| GOTERM_BP_DIRECT | flower development, cellular response to hypoxia, |
| GOTERM_CC_DIRECT | nucleus, |
| **At5g22570** | **WRKY DNA-binding protein 38(WRKY38)** |
| GOTERM_BP_DIRECT | transcription, DNA-templated, regulation of transcription, DNA-templated, salicylic acid mediated signaling pathway, defense response to bacterium, |
| GOTERM_CC_DIRECT | nucleus, |
| GOTERM_MF_DIRECT | transcription factor activity, sequence-specific DNA binding, sequence-specific DNA binding, |
| **At3g01970** | **WRKY DNA-binding protein 45(WRKY45)** |
| GOTERM_BP_DIRECT | transcription, DNA-templated, regulation of transcription, DNA-templated, phosphate ion transport, |
| GOTERM_CC_DIRECT | nucleus, |

| | | |
|---|---|---|
| | GOTERM_MF_DIRECT | transcription factor activity, sequence-specific DNA binding, transcription factor binding, sequence-specific DNA binding, transcription regulatory region DNA binding, |
| | **At2g46400** | **WRKY DNA-binding protein 46(WRKY46)** |
| | GOTERM_BP_DIRECT | transcription, DNA-templated, regulation of transcription, DNA-templated, response to chitin, lateral root development, |
| | GOTERM_CC_DIRECT | nucleus, nucleolus, |
| | GOTERM_MF_DIRECT | core promoter proximal region sequence-specific DNA binding, transcription factor activity, sequence-specific DNA binding, sequence-specific DNA binding, transcription regulatory region DNA binding, |
| | **At2g40740** | **WRKY DNA-binding protein 55(WRKY55)** |
| | GOTERM_BP_DIRECT | transcription, DNA-templated, regulation of transcription, DNA-templated, |
| | GOTERM_CC_DIRECT | nucleus, |
| | GOTERM_MF_DIRECT | transcription factor activity, sequence-specific DNA binding, sequence-specific DNA binding, |
| | **At1g80590** | **WRKY DNA-binding protein 66(WRKY66)** |
| | GOTERM_BP_DIRECT | transcription, DNA-templated, regulation of transcription, DNA-templated, |
| | GOTERM_CC_DIRECT | nucleus, |
| | GOTERM_MF_DIRECT | transcription factor activity, sequence-specific DNA binding, sequence-specific DNA binding, |
| | **At3g56400** | **WRKY DNA-binding protein 70(WRKY70)** |
| | GOTERM_BP_DIRECT | transcription, DNA-templated, regulation of transcription, DNA-templated, response to salicylic acid, response to jasmonic acid, systemic acquired resistance, salicylic acid mediated signaling pathway, induced systemic resistance, jasmonic acid mediated signaling pathway, response to chitin, regulation of defense response, defense response to bacterium, negative regulation of transcription, DNA-templated, defense response to fungus, negative regulation of leaf senescence, |
| | GOTERM_CC_DIRECT | nucleus, |
| | GOTERM_MF_DIRECT | transcription factor activity, sequence-specific DNA binding, protein binding, sequence-specific DNA binding, |
| | **At5g13080** | **WRKY DNA-binding protein 75(WRKY75)** |
| | GOTERM_BP_DIRECT | negative regulation of transcription from RNA polymerase II promoter, transcription, DNA-templated, regulation of transcription, DNA-templated, atrichoblast differentiation, regulation of response to nutrient levels, regulation of DNA-templated transcription in response to stress, lateral root development, |
| | GOTERM_CC_DIRECT | nucleus, |
| | GOTERM_MF_DIRECT | core promoter sequence-specific DNA binding, transcription factor activity, sequence-specific DNA binding, sequence-specific DNA binding, transcription regulatory region DNA binding, |
| | **At2g26400** | **acireductone dioxygenase 3(ARD3)** |
| | GOTERM_BP_DIRECT | methionine metabolic process, L-methionine biosynthetic process from methylthioadenosine, oxidation-reduction process, |
| | GOTERM_CC_DIRECT | nucleus, cytoplasm, plasma membrane, |
| | GOTERM_MF_DIRECT | iron ion binding, heteropolysaccharide binding, acireductone dioxygenase [iron(II)-requiring] activity, metal ion binding, |
| | **At1g68620** | **alpha/beta-Hydrolases superfamily protein(AT1G68620)** |
| | GOTERM_BP_DIRECT | metabolic process, catabolic process, |
| | GOTERM_CC_DIRECT | nucleus, |

| | | |
|---|---|---|
| | GOTERM_MF_DIRECT | hydrolase activity, carboxylic ester hydrolase activity, |
| | **At2g18360** | **alpha/beta-Hydrolases superfamily protein(AT2G18360)** |
| | GOTERM_MF_DIRECT | hydrolase activity, |
| | **At2g39400** | **alpha/beta-Hydrolases superfamily protein(AT2G39400)** |
| | GOTERM_BP_DIRECT | lipid metabolic process, |
| | GOTERM_CC_DIRECT | cytoplasm, endoplasmic reticulum, Golgi apparatus, membrane, |
| | GOTERM_MF_DIRECT | lipase activity, hydrolase activity, acylglycerol lipase activity, |
| | **At5g24210** | **alpha/beta-Hydrolases superfamily protein(AT5G24210)** |
| | GOTERM_BP_DIRECT | lipid metabolic process, |
| | GOTERM_CC_DIRECT | nucleus, |
| | GOTERM_MF_DIRECT | triglyceride lipase activity, hydrolase activity, |
| | **At1g06800** | **alpha/beta-Hydrolases superfamily protein(PLA-I{gamma}1)** |
| | GOTERM_BP_DIRECT | lipid metabolic process, lipid catabolic process, |
| | GOTERM_CC_DIRECT | chloroplast, |
| | GOTERM_MF_DIRECT | triglyceride lipase activity, phosphatidylcholine 1-acylhydrolase activity, hydrolase activity, galactolipase activity, |
| | **At3g22370** | **alternative oxidase 1A(AOX1A)** |
| | GOTERM_BP_DIRECT | response to cold, mitochondria-nucleus signaling pathway, cellular respiration, oxidation-reduction process, |
| | GOTERM_CC_DIRECT | mitochondrion, mitochondrial inner membrane, integral component of membrane, respiratory chain, |
| | GOTERM_MF_DIRECT | alternative oxidase activity, metal ion binding, |
| | **At1g32350** | **alternative oxidase 1D(AOX1D)** |
| | GOTERM_BP_DIRECT | oxidation-reduction process, |
| | GOTERM_CC_DIRECT | mitochondrion, mitochondrial inner membrane, integral component of membrane, respiratory chain, |
| | GOTERM_MF_DIRECT | alternative oxidase activity, metal ion binding, |
| | **At5g19140** | **aluminum induced protein with YGL and LRDR motifs(AILP1)** |
| | GOTERM_BP_DIRECT | asparagine biosynthetic process, glutamine metabolic process, response to auxin, response to aluminum ion, |
| | GOTERM_CC_DIRECT | cytoplasm, cytosol, plasma membrane, |
| | GOTERM_MF_DIRECT | asparagine synthase (glutamine-hydrolyzing) activity, protein homodimerization activity, |
| | **At3g13950** | **ankyrin(AT3G13950)** |
| | GOTERM_CC_DIRECT | integral component of membrane, |
| | **At3g44720** | **arogenate dehydratase 4(ADT4)** |
| | GOTERM_BP_DIRECT | L-phenylalanine biosynthetic process, response to karrikin, |

| | | |
|---|---|---|
| | GOTERM_CC_DIRECT | chloroplast, chloroplast stroma, |
| | GOTERM_MF_DIRECT | prephenate dehydratase activity, amino acid binding, arogenate dehydratase activity, |
| | **At2g44480** | **beta glucosidase 17(BGLU17)** |
| | GOTERM_BP_DIRECT | carbohydrate metabolic process, response to salt stress, response to hormone, glucosinolate catabolic process, |
| | GOTERM_CC_DIRECT | extracellular region, |
| | GOTERM_MF_DIRECT | hydrolase activity, hydrolyzing O-glycosyl compounds, beta-glucosidase activity, scopolin beta-glucosidase activity, |
| | **At3g60120** | **beta glucosidase 27(BGLU27)** |
| | GOTERM_BP_DIRECT | carbohydrate metabolic process, response to salt stress, response to hormone, glucosinolate catabolic process, |
| | GOTERM_CC_DIRECT | nucleus, |
| | GOTERM_MF_DIRECT | hydrolase activity, hydrolyzing O-glycosyl compounds, beta-glucosidase activity, scopolin beta-glucosidase activity, |
| | **At5g24540** | **beta glucosidase 31(BGLU31)** |
| | GOTERM_BP_DIRECT | carbohydrate metabolic process, response to salt stress, response to hormone, glucosinolate catabolic process, response to other organism, |
| | GOTERM_CC_DIRECT | extracellular region, |
| | GOTERM_MF_DIRECT | hydrolase activity, hydrolyzing O-glycosyl compounds, beta-glucosidase activity, scopolin beta-glucosidase activity, |
| | **At5g24550** | beta glucosidase 32(BGLU32) |
| | GOTERM_BP_DIRECT | carbohydrate metabolic process, response to salt stress, response to hormone, glucosinolate catabolic process, response to other organism, |
| | GOTERM_CC_DIRECT | extracellular region, |
| | GOTERM_MF_DIRECT | hydrolase activity, hydrolyzing O-glycosyl compounds, beta-glucosidase activity, scopolin beta-glucosidase activity, |
| | **At1g61820** | **beta glucosidase 46(BGLU46)** |
| | GOTERM_BP_DIRECT | carbohydrate metabolic process, lignin biosynthetic process, glycosyl compound metabolic process, |
| | GOTERM_CC_DIRECT | extracellular region, |
| | GOTERM_MF_DIRECT | hydrolase activity, hydrolyzing O-glycosyl compounds, beta-glucosidase activity, coniferin beta-glucosidase activity, scopolin beta-glucosidase activity, |
| | **At3g57260** | **beta-1,3-glucanase 2(BGL2)** |
| | GOTERM_BP_DIRECT | carbohydrate metabolic process, response to cold, systemic acquired resistance, |
| | GOTERM_CC_DIRECT | cell wall, vacuole, endoplasmic reticulum, chloroplast, anchored component of plasma membrane, apoplast, |
| | GOTERM_MF_DIRECT | glucan exo-1,3-beta-glucosidase activity, hydrolase activity, hydrolyzing O-glycosyl compounds, protein binding, cellulase activity, polysaccharide binding, glucan endo-1,3-beta-D-glucosidase activity, |
| | **At5g20230** | **blue-copper-binding protein(BCB)** |
| | GOTERM_BP_DIRECT | response to oxidative stress, response to wounding, response to absence of light, aluminum cation transport, defense response to fungus, oxidation-reduction process, cellular response to cold, regulation of lignin biosynthetic process, |
| | GOTERM_CC_DIRECT | vacuole, plasma membrane, anchored component of membrane, anchored component of plasma membrane, |
| | GOTERM_MF_DIRECT | copper ion binding, protein binding, electron carrier activity, metal ion binding, |

| | | |
|---|---|---|
| | **At1g67980** | **caffeoyl-CoA 3-O-methyltransferase(CCOAMT)** |
| | GOTERM_BP_DIRECT | lignin biosynthetic process, methylation, |
| | GOTERM_CC_DIRECT | cytoplasm, cytosol, |
| | GOTERM_MF_DIRECT | O-methyltransferase activity, caffeoyl-CoA O-methyltransferase activity, metal ion binding, |
| | **At2g41730** | **calcium-binding site protein(AT2G41730)** |
| | GOTERM_BP_DIRECT | anaerobic respiration, |
| | GOTERM_CC_DIRECT | nucleus, cytoplasm, |
| | **At5g42380** | **calmodulin like 37(CML37)** |
| | GOTERM_BP_DIRECT | response to ozone, |
| | GOTERM_CC_DIRECT | nucleus, cytoplasm, |
| | GOTERM_MF_DIRECT | calcium ion binding, |
| | **At1g76650** | **calmodulin-like 38(CML38)** |
| | GOTERM_BP_DIRECT | response to wounding, |
| | GOTERM_CC_DIRECT | nucleus, plasma membrane, |
| | GOTERM_MF_DIRECT | calcium ion binding, |
| | **At3g50770** | **calmodulin-like 41(CML41)** |
| | GOTERM_MF_DIRECT | calcium ion binding, |
| | **At4g23700** | **cation/H+ exchanger 17(CHX17)** |
| | GOTERM_BP_DIRECT | protein targeting to vacuole, cation transport, potassium ion transport, regulation of pH, |
| | GOTERM_CC_DIRECT | nucleus, late endosome, integral component of membrane, |
| | GOTERM_MF_DIRECT | monovalent cation:proton antiporter activity, sodium:proton antiporter activity, |
| | **At5g41610** | **cation/H+ exchanger 18(CHX18)** |
| | GOTERM_BP_DIRECT | cation transport, potassium ion transport, regulation of pH, |
| | GOTERM_CC_DIRECT | late endosome, chloroplast, integral component of membrane, |
| | GOTERM_MF_DIRECT | monovalent cation:proton antiporter activity, sodium:proton antiporter activity, |
| | **At1g21250** | **cell wall-associated kinase(WAK1)** |
| | GOTERM_BP_DIRECT | protein phosphorylation, cell surface receptor signaling pathway, response to virus, response to salicylic acid, defense response to fungus, |
| | GOTERM_CC_DIRECT | extracellular region, vacuole, plasma membrane, plant-type cell wall, integral component of membrane, |
| | GOTERM_MF_DIRECT | protein serine/threonine kinase activity, calcium ion binding, protein binding, ATP binding, kinase activity, polysaccharide binding, |
| | **At1g55850** | **cellulose synthase like E1(CSLE1)** |
| | GOTERM_BP_DIRECT | polysaccharide biosynthetic process, plant-type cell wall biogenesis, plant-type primary cell wall biogenesis, cellulose biosynthetic process, cell wall organization, |

| | |
|---|---|
| GOTERM_CC_DIRECT | Golgi membrane, endoplasmic reticulum, endoplasmic reticulum membrane, Golgi apparatus, plasma membrane, integral component of membrane, |
| GOTERM_MF_DIRECT | RNA polymerase II regulatory region sequence-specific DNA binding, transferase activity, transferring glycosyl groups, cellulose synthase activity, cellulose synthase (UDP-forming) activity, |
| **At2g43570** | **chitinase(CHI)** |
| GOTERM_BP_DIRECT | polysaccharide catabolic process, carbohydrate metabolic process, chitin catabolic process, amino sugar metabolic process, response to virus, systemic acquired resistance, leaf senescence, response to silver ion, cell wall macromolecule catabolic process, response to amitrole, |
| GOTERM_CC_DIRECT | extracellular region, intracellular, plant-type cell wall, apoplast, |
| GOTERM_MF_DIRECT | chitinase activity, chitin binding, |
| **At1g80820** | **cinnamoyl coa reductase(CCR2)** |
| GOTERM_BP_DIRECT | defense response, circadian rhythm, response to cold, phenylpropanoid biosynthetic process, lignin biosynthetic process, negative regulation of circadian rhythm, oxidation-reduction process, |
| GOTERM_MF_DIRECT | cinnamoyl-CoA reductase activity, coenzyme binding, |
| **At4g12280** | **copper amine oxidase family protein(AT4G12280)** |
| GOTERM_BP_DIRECT | amine metabolic process, oxidation-reduction process, |
| GOTERM_CC_DIRECT | nucleus, |
| GOTERM_MF_DIRECT | copper ion binding, primary amine oxidase activity, quinone binding, |
| **At5g57510** | **cotton fiber protein(AT5G57510)** |
| GOTERM_BP_DIRECT | cellular response to hypoxia, |
| GOTERM_CC_DIRECT | nucleus, |
| **At3g17690** | **cyclic nucleotide gated channel 19(CNGC19)** |
| GOTERM_BP_DIRECT | transcription, DNA-templated, regulation of transcription, DNA-templated, red, far-red light phototransduction, regulation of membrane potential, |
| GOTERM_CC_DIRECT | nucleus, integral component of plasma membrane, chloroplast, |
| GOTERM_MF_DIRECT | transcription factor activity, sequence-specific DNA binding, ion channel activity, voltage-gated potassium channel activity, protein binding, calmodulin binding, cyclic nucleotide binding, cAMP binding, cGMP binding, sequence-specific DNA binding, |
| **At4g23180** | **cysteine-rich RLK (RECEPTOR-like protein kinase) 10(CRK10)** |
| GOTERM_BP_DIRECT | protein phosphorylation, defense response to bacterium, |
| GOTERM_CC_DIRECT | plasma membrane, plasmodesma, integral component of membrane, |
| GOTERM_MF_DIRECT | protein serine/threonine kinase activity, ATP binding, kinase activity, |
| **At4g23190** | **cysteine-rich RLK (RECEPTOR-like protein kinase) 11(CRK11)** |
| GOTERM_BP_DIRECT | protein phosphorylation, response to oxidative stress, defense response to bacterium, incompatible interaction, |
| GOTERM_CC_DIRECT | extracellular region, plasma membrane, plasmodesma, membrane, integral component of membrane, |
| GOTERM_MF_DIRECT | protein kinase activity, protein serine/threonine kinase activity, ATP binding, kinase activity, |
| **At4g11470** | **cysteine-rich RLK (RECEPTOR-like protein kinase) 31(CRK31)** |
| GOTERM_BP_DIRECT | protein phosphorylation, defense response to bacterium, |

| | |
|---|---|
| GOTERM_CC_DIRECT | extracellular region, plasma membrane, plasmodesma, integral component of membrane, |
| GOTERM_MF_DIRECT | protein kinase activity, protein serine/threonine kinase activity, ATP binding, |
| **At4g04540** | **cysteine-rich RLK (RECEPTOR-like protein kinase) 39(CRK39)** |
| GOTERM_BP_DIRECT | protein phosphorylation, defense response to bacterium, |
| GOTERM_CC_DIRECT | extracellular region, plasma membrane, plasmodesma, integral component of membrane, |
| GOTERM_MF_DIRECT | protein serine/threonine kinase activity, ATP binding, kinase activity, |
| **At2g32190** | **cysteine-rich/transmembrane domain A-like protein(AT2G32190)** |
| GOTERM_BP_DIRECT | developmental process involved in reproduction, |
| GOTERM_CC_DIRECT | plasma membrane, |
| **At3g50930** | **cytochrome BC1 synthesi(BCS1)** |
| GOTERM_BP_DIRECT | response to molecule of bacterial origin, cell death, response to UV, response to bacterium, plant-type hypersensitive response, salicylic acid mediated signaling pathway, |
| GOTERM_CC_DIRECT | mitochondrion, mitochondrial envelope, mitochondrial outer membrane, plasma membrane, integral component of membrane, |
| GOTERM_MF_DIRECT | ATP binding, ATPase activity, identical protein binding, |
| **At2g30750** | **cytochrome P450 family 71 polypeptide(CYP71A12)** |
| GOTERM_BP_DIRECT | response to bacterium, induced systemic resistance, secondary metabolite biosynthetic process, |
| GOTERM_CC_DIRECT | membrane, integral component of membrane, |
| GOTERM_MF_DIRECT | monooxygenase activity, iron ion binding, oxidoreductase activity, acting on paired donors, with incorporation or reduction of molecular oxygen, oxidoreductase activity, acting on paired donors, with incorporation or reduction of molecular oxygen, NAD(P)H as one donor, and incorporation of one atom of oxygen, oxygen binding, heme binding, |
| **At1g55940** | **cytochrome P450 family protein(CYP708A1)** |
| GOTERM_BP_DIRECT | multicellular organism development, brassinosteroid homeostasis, sterol metabolic process, brassinosteroid biosynthetic process, oxidation-reduction process, |
| GOTERM_CC_DIRECT | nucleus, endoplasmic reticulum, integral component of membrane, |
| GOTERM_MF_DIRECT | monooxygenase activity, iron ion binding, oxidoreductase activity, acting on paired donors, with incorporation or reduction of molecular oxygen, oxygen binding, heme binding, |
| **At4g22690** | **cytochrome P450, family 706, subfamily A, polypeptide 1(CYP706A1)** |
| GOTERM_BP_DIRECT | secondary metabolite biosynthetic process, oxidation-reduction process, |
| GOTERM_CC_DIRECT | cell wall, mitochondrion, vacuolar membrane, Golgi apparatus, plasma membrane, chloroplast, |
| GOTERM_MF_DIRECT | iron ion binding, oxidoreductase activity, acting on paired donors, with incorporation or reduction of molecular oxygen, NAD(P)H as one donor, and incorporation of one atom of oxygen, oxygen binding, heme binding, |
| **At5g36110** | **cytochrome P450, family 716, subfamily A, polypeptide 1(CYP716A1)** |
| GOTERM_BP_DIRECT | multicellular organism development, brassinosteroid homeostasis, sterol metabolic process, brassinosteroid biosynthetic process, oxidation-reduction process, |
| GOTERM_CC_DIRECT | extracellular region, |

| | | |
|---|---|---|
| | GOTERM_MF_DIRECT | monooxygenase activity, iron ion binding, oxidoreductase activity, acting on paired donors, with incorporation or reduction of molecular oxygen, oxygen binding, heme binding, |
| | **At3g14660** | **cytochrome P450, family 72, subfamily A, polypeptide 13(CYP72A13)** |
| | GOTERM_BP_DIRECT | oxidation-reduction process, |
| | GOTERM_CC_DIRECT | integral component of membrane, |
| | GOTERM_MF_DIRECT | monooxygenase activity, iron ion binding, oxidoreductase activity, acting on paired donors, with incorporation or reduction of molecular oxygen, oxygen binding, heme binding, |
| | **At4g37370** | **cytochrome P450, family 81, subfamily D, polypeptide 8(CYP81D8)** |
| | GOTERM_BP_DIRECT | indole glucosinolate metabolic process, secondary metabolite biosynthetic process, oxidation-reduction process, response to karrikin, defense response to other organism, |
| | GOTERM_CC_DIRECT | extracellular region, endoplasmic reticulum, plasma membrane, integral component of membrane, |
| | GOTERM_MF_DIRECT | iron ion binding, oxidoreductase activity, acting on paired donors, with incorporation or reduction of molecular oxygen, NAD(P)H as one donor, and incorporation of one atom of oxygen, oxygen binding, heme binding, |
| | **At5g57220** | **cytochrome P450, family 81, subfamily F, polypeptide 2(CYP81F2)** |
| | GOTERM_BP_DIRECT | defense response to insect, response to bacterium, induced systemic resistance, indole glucosinolate biosynthetic process, glucosinolate metabolic process, indole glucosinolate metabolic process, defense response to bacterium, defense response to fungus, defense response by callose deposition in cell wall, oxidation-reduction process, cellular response to hypoxia, |
| | GOTERM_CC_DIRECT | membrane, integral component of membrane, |
| | GOTERM_MF_DIRECT | monooxygenase activity, iron ion binding, oxidoreductase activity, acting on paired donors, with incorporation or reduction of molecular oxygen, oxidoreductase activity, acting on paired donors, with incorporation or reduction of molecular oxygen, NAD(P)H as one donor, and incorporation of one atom of oxygen, oxygen binding, heme binding, |
| | **At4g31970** | **cytochrome P450, family 82, subfamily C, polypeptide 2(CYP82C2)** |
| | GOTERM_BP_DIRECT | secondary metabolite biosynthetic process, oxidation-reduction process, cellular response to hypoxia, |
| | GOTERM_CC_DIRECT | chloroplast, membrane, integral component of membrane, |
| | GOTERM_MF_DIRECT | monooxygenase activity, iron ion binding, oxidoreductase activity, acting on paired donors, with incorporation or reduction of molecular oxygen, oxidoreductase activity, acting on paired donors, with incorporation or reduction of molecular oxygen, NAD(P)H as one donor, and incorporation of one atom of oxygen, oxygen binding, heme binding, |
| | **At2g24180** | **cytochrome p450 71b6(CYP71B6)** |
| | GOTERM_BP_DIRECT | indole-containing compound metabolic process, secondary metabolite biosynthetic process, oxidation-reduction process, |
| | GOTERM_CC_DIRECT | mitochondrion, endoplasmic reticulum, Golgi apparatus, plasma membrane, membrane, integral component of membrane, |
| | GOTERM_MF_DIRECT | monooxygenase activity, iron ion binding, oxidoreductase activity, acting on paired donors, with incorporation or reduction of molecular oxygen, NAD(P)H as one donor, and incorporation of one atom of oxygen, oxygen binding, heme binding, |
| | **At5g64890** | **elicitor peptide 2 precursor(PROPEP2)** |
| | GOTERM_BP_DIRECT | defense response, incompatible interaction, |
| | GOTERM_CC_DIRECT | cytoplasm, |
| | **At5g64905** | **elicitor peptide 3 precursor(PROPEP3)** |

| | |
|---|---|
| GOTERM_BP_DIRECT | defense response, cellular response to hypoxia, |
| GOTERM_CC_DIRECT | nucleus, |
| **At5g47220** | **ethylene responsive element binding factor 2(ERF2)** |
| GOTERM_BP_DIRECT | vasculature development, transcription, DNA-templated, regulation of transcription, DNA-templated, induced systemic resistance, jasmonic acid mediated signaling pathway, ethylene-activated signaling pathway, response to chitin, positive regulation of transcription, DNA-templated, cell division, |
| GOTERM_CC_DIRECT | intracellular, nucleus, |
| GOTERM_MF_DIRECT | DNA binding, transcription factor activity, sequence-specific DNA binding, |
| **At5g22500** | **fatty acid reductase 1(FAR1)** |
| GOTERM_BP_DIRECT | lipid metabolic process, microsporogenesis, response to wounding, response to salt stress, suberin biosynthetic process, oxidation-reduction process, |
| GOTERM_CC_DIRECT | plasma membrane, chloroplast, |
| GOTERM_MF_DIRECT | oxidoreductase activity, acting on the CH-CH group of donors, long-chain-fatty-acyl-CoA reductase activity, fatty-acyl-CoA reductase (alcohol-forming) activity, |
| **At3g44540** | **fatty acid reductase 4(FAR4)** |
| GOTERM_BP_DIRECT | lipid metabolic process, microsporogenesis, response to wounding, response to salt stress, suberin biosynthetic process, long-chain fatty-acyl-CoA metabolic process, oxidation-reduction process, |
| GOTERM_CC_DIRECT | chloroplast, intracellular membrane-bounded organelle, |
| GOTERM_MF_DIRECT | oxidoreductase activity, acting on the CH-CH group of donors, long-chain-fatty-acyl-CoA reductase activity, fatty-acyl-CoA reductase (alcohol-forming) activity, |
| **At3g44550** | **fatty acid reductase 5(FAR5)** |
| GOTERM_BP_DIRECT | lipid metabolic process, microsporogenesis, response to wounding, response to salt stress, suberin biosynthetic process, long-chain fatty-acyl-CoA metabolic process, oxidation-reduction process, |
| GOTERM_CC_DIRECT | intracellular membrane-bounded organelle, |
| GOTERM_MF_DIRECT | oxidoreductase activity, acting on the CH-CH group of donors, long-chain-fatty-acyl-CoA reductase activity, fatty-acyl-CoA reductase (alcohol-forming) activity, |
| **At1g55790** | **ferredoxin-fold anticodon-binding domain protein(AT1G55790)** |
| GOTERM_BP_DIRECT | metal ion transport, rRNA base methylation, |
| GOTERM_CC_DIRECT | nucleus, cytoplasm, |
| GOTERM_MF_DIRECT | metal ion binding, rRNA (uridine-N3-)-methyltransferase activity, |
| **At1g79160** | **filamentous hemagglutinin transporter(AT1G79160)** |
| **At1g19250** | **flavin-dependent monooxygenase 1(FMO1)** |
| GOTERM_BP_DIRECT | plant-type hypersensitive response, systemic acquired resistance, defense response signaling pathway, resistance gene-dependent, defense response signaling pathway, resistance gene-independent, defense response to bacterium, defense response to fungus, response to other organism, oxidation-reduction process, cellular response to hypoxia, |
| GOTERM_CC_DIRECT | chloroplast, |
| GOTERM_MF_DIRECT | monooxygenase activity, N,N-dimethylaniline monooxygenase activity, flavin adenine dinucleotide binding, NADP binding, |
| **At1g56600** | **galactinol synthase 2(GolS2)** |

| | | |
|---|---|---|
| | GOTERM_BP_DIRECT | galactose metabolic process, response to oxidative stress, response to cold, response to water deprivation, response to salt stress, response to abscisic acid, carbohydrate biosynthetic process, |
| | GOTERM_CC_DIRECT | nucleus, cytoplasm, |
| | GOTERM_MF_DIRECT | transferase activity, transferring glycosyl groups, transferase activity, transferring hexosyl groups, metal ion binding, inositol 3-alpha-galactosyltransferase activity, |
| | **At1g60470** | **galactinol synthase 4(GolS4)** |
| | GOTERM_BP_DIRECT | galactose metabolic process, response to oxidative stress, carbohydrate biosynthetic process, |
| | GOTERM_CC_DIRECT | nucleus, cytoplasm, |
| | GOTERM_MF_DIRECT | transferase activity, transferring glycosyl groups, transferase activity, transferring hexosyl groups, metal ion binding, inositol 3-alpha-galactosyltransferase activity, |
| | **At1g18970** | **germin-like protein 4(GLP4)** |
| | GOTERM_BP_DIRECT | oxalate metabolic process, |
| | GOTERM_CC_DIRECT | extracellular region, cell wall, apoplast, |
| | GOTERM_MF_DIRECT | manganese ion binding, nutrient reservoir activity, oxalate decarboxylase activity, |
| | **At5g39100** | **germin-like protein 6(GLP6)** |
| | GOTERM_BP_DIRECT | oxalate metabolic process, |
| | GOTERM_CC_DIRECT | extracellular region, cell wall, apoplast, |
| | GOTERM_MF_DIRECT | manganese ion binding, nutrient reservoir activity, oxalate decarboxylase activity, |
| | **At2g02000** | **glutamate decarboxylase 3(GAD3)** |
| | GOTERM_BP_DIRECT | glutamate metabolic process, |
| | GOTERM_MF_DIRECT | glutamate decarboxylase activity, calmodulin binding, pyridoxal phosphate binding, |
| | **At3g03910** | **glutamate dehydrogenase 3(GDH3)** |
| | GOTERM_BP_DIRECT | cellular amino acid metabolic process, regulation of nitrogen compound metabolic process, oxidation-reduction process, |
| | GOTERM_CC_DIRECT | mitochondrion, |
| | GOTERM_MF_DIRECT | glutamate dehydrogenase (NAD+) activity, glutamate dehydrogenase [NAD(P)+] activity, oxidoreductase activity, |
| | **At5g11210** | **glutamate receptor 2.5(GLR2.5)** |
| | GOTERM_BP_DIRECT | calcium ion transport, cellular calcium ion homeostasis, response to light stimulus, calcium-mediated signaling, cellular response to amino acid stimulus, |
| | GOTERM_CC_DIRECT | intracellular, plasma membrane, integral component of membrane, |
| | GOTERM_MF_DIRECT | G-protein coupled receptor activity, ionotropic glutamate receptor activity, intracellular ligand-gated ion channel activity, calcium channel activity, glutamate receptor activity, |
| | **At4g25760** | **glutamine dumper 2(GDU2)** |
| | GOTERM_BP_DIRECT | amino acid transport, regulation of amino acid export, |
| | GOTERM_CC_DIRECT | nucleus, integral component of membrane, |
| | **At5g16570** | **glutamine synthetase 1;4(GLN1;4)** |

| | |
|---|---|
| GOTERM_BP_DIRECT | glutamine biosynthetic process, nitrogen fixation, nitrate assimilation, |
| GOTERM_CC_DIRECT | cytoplasm, cytosol, |
| GOTERM_MF_DIRECT | glutamate-ammonia ligase activity, protein binding, ATP binding, |
| **At1g74590** | **glutathione S-transferase TAU 10(GSTU10)** |
| GOTERM_BP_DIRECT | glutathione metabolic process, toxin catabolic process, response to toxic substance, |
| GOTERM_CC_DIRECT | cytoplasm, cytosol, |
| GOTERM_MF_DIRECT | glutathione transferase activity, |
| **At1g69930** | **glutathione S-transferase TAU 11(GSTU11)** |
| GOTERM_BP_DIRECT | glutathione metabolic process, toxin catabolic process, response to toxic substance, |
| GOTERM_CC_DIRECT | cytoplasm, cytosol, |
| GOTERM_MF_DIRECT | glutathione transferase activity, |
| **At1g69920** | **glutathione S-transferase TAU 12(GSTU12)** |
| GOTERM_BP_DIRECT | glutathione metabolic process, toxin catabolic process, response to toxic substance, |
| GOTERM_CC_DIRECT | nucleus, cytoplasm, |
| GOTERM_MF_DIRECT | glutathione transferase activity, |
| **At1g78340** | **glutathione S-transferase TAU 22(GSTU22)** |
| GOTERM_BP_DIRECT | glutathione metabolic process, toxin catabolic process, response to toxic substance, |
| GOTERM_CC_DIRECT | nucleus, cytoplasm, cytosol, |
| GOTERM_MF_DIRECT | glutathione transferase activity, |
| **At1g17170** | **glutathione S-transferase TAU 24(GSTU24)** |
| GOTERM_BP_DIRECT | glutathione metabolic process, toxin catabolic process, response to toxic substance, 2,4,6-trinitrotoluene catabolic process, |
| GOTERM_CC_DIRECT | cytoplasm, cytosol, |
| GOTERM_MF_DIRECT | glutathione transferase activity, glutathione binding, |
| **At3g09270** | **glutathione S-transferase TAU 8(GSTU8)** |
| GOTERM_BP_DIRECT | glutathione metabolic process, toxin catabolic process, response to toxic substance, response to cadmium ion, |
| GOTERM_CC_DIRECT | cytoplasm, cytosol, |
| GOTERM_MF_DIRECT | glutathione transferase activity, |
| **At2g29480** | **glutathione S-transferase tau 2(GSTU2)** |
| GOTERM_BP_DIRECT | glutathione metabolic process, toxin catabolic process, response to toxic substance, |
| GOTERM_CC_DIRECT | cytoplasm, cytosol, |
| GOTERM_MF_DIRECT | glutathione transferase activity, |

| | | |
|---|---|---|
| | **At5g02780** | **glutathione transferase lambda 1(GSTL1)** |
| | GOTERM_BP_DIRECT | glutathione metabolic process, response to toxic substance, protein glutathionylation, |
| | GOTERM_CC_DIRECT | cytoplasm, cytosol, |
| | GOTERM_MF_DIRECT | glutathione transferase activity, |
| | **At2g41690** | **heat shock transcription factor B3(HSFB3)** |
| | GOTERM_BP_DIRECT | transcription, DNA-templated, regulation of transcription, DNA-templated, response to cadmium ion, |
| | GOTERM_CC_DIRECT | nucleus, cytoplasm, |
| | GOTERM_MF_DIRECT | DNA binding, transcription factor activity, sequence-specific DNA binding, sequence-specific DNA binding, |
| | **At4g13420** | **high affinity K+ transporter 5(HAK5)** |
| | GOTERM_BP_DIRECT | potassium ion transport, potassium ion transmembrane transport, |
| | GOTERM_CC_DIRECT | plasma membrane, membrane, integral component of membrane, |
| | GOTERM_MF_DIRECT | potassium:sodium symporter activity, potassium ion transmembrane transporter activity, |
| | **At2g18050** | **histone H1-3(HIS1-3)** |
| | GOTERM_BP_DIRECT | nucleosome assembly, response to water deprivation, |
| | GOTERM_CC_DIRECT | nucleosome, nucleus, |
| | GOTERM_MF_DIRECT | DNA binding, nucleosomal DNA binding, |
| | **At5g03390** | **hypothetical protein (DUF295)(AT5G03390)** |
| | GOTERM_CC_DIRECT | chloroplast, |
| | **At1g28190** | **hypothetical protein(AT1G28190)** |
| | GOTERM_CC_DIRECT | nucleus, |
| | **At1g32920** | **hypothetical protein(AT1G32920)** |
| | GOTERM_BP_DIRECT | response to wounding, |
| | **At1g35210** | **hypothetical protein(AT1G35210)** |
| | GOTERM_BP_DIRECT | response to UV-B, |
| | GOTERM_CC_DIRECT | mitochondrion, |
| | **At1g56320** | **hypothetical protein(AT1G56320)** |
| | GOTERM_CC_DIRECT | anchored component of membrane, |
| | **At1g63530** | **hypothetical protein(AT1G63530)** |
| | GOTERM_BP_DIRECT | pollen tube growth, |
| | GOTERM_CC_DIRECT | nucleus, |
| | **At2g36440** | **hypothetical protein(AT2G36440)** |

| | | |
|---|---|---|
| | GOTERM_CC_DIRECT | mitochondrion, |
| | **At2g47200** | **hypothetical protein(AT2G47200)** |
| | GOTERM_CC_DIRECT | mitochondrion, |
| | **At2g48090** | **hypothetical protein(AT2G48090)** |
| | GOTERM_CC_DIRECT | nucleus, |
| | **At3g23170** | **hypothetical protein(AT3G23170)** |
| | GOTERM_CC_DIRECT | chloroplast, |
| | **At4g33666** | **hypothetical protein(AT4G33666)** |
| | **At5g05300** | **hypothetical protein(AT5G05300)** |
| | GOTERM_BP_DIRECT | regulation of innate immune response, |
| | GOTERM_CC_DIRECT | chloroplast, |
| | **At5g17350** | **hypothetical protein(AT5G17350)** |
| | GOTERM_CC_DIRECT | chloroplast, |
| | **At5g22270** | **hypothetical protein(AT5G22270)** |
| | GOTERM_CC_DIRECT | nucleus, |
| | **At5g22530** | **hypothetical protein(AT5G22530)** |
| | **At5g60350** | **hypothetical protein(AT5G60350)** |
| | GOTERM_CC_DIRECT | mitochondrion, |
| | **At4g14365** | **hypothetical protein(XBAT34)** |
| | GOTERM_BP_DIRECT | protein ubiquitination, |
| | GOTERM_CC_DIRECT | nucleus, |
| | GOTERM_MF_DIRECT | zinc ion binding, ligase activity, |
| | **At1g33790** | **jacalin lectin family protein(AT1G33790)** |
| | GOTERM_CC_DIRECT | cytoplasm, chloroplast, |
| | GOTERM_MF_DIRECT | carbohydrate binding, |
| | **At1g18140** | **laccase 1(LAC1)** |
| | GOTERM_BP_DIRECT | lignin biosynthetic process, lignin catabolic process, oxidation-reduction process, |
| | GOTERM_CC_DIRECT | extracellular region, apoplast, |
| | GOTERM_MF_DIRECT | copper ion binding, oxidoreductase activity, oxidizing metal ions, hydroquinone:oxygen oxidoreductase activity, |
| | **At5g01540** | **lectin receptor kinase a4.1(LECRKA4.1)** |
| | GOTERM_BP_DIRECT | protein phosphorylation, response to abscisic acid, abscisic acid-activated signaling pathway, seed germination, defense response to bacterium, pathogen-associated molecular pattern dependent induction by symbiont of host innate immune response, |

| | |
|---|---|
| GOTERM_CC_DIRECT | plasma membrane, integral component of membrane, |
| GOTERM_MF_DIRECT | protein serine/threonine kinase activity, protein binding, ATP binding, kinase activity, carbohydrate binding, |
| **At5g01550** | **lectin receptor kinase a4.1(LECRKA4.2)** |
| GOTERM_BP_DIRECT | protein phosphorylation, abscisic acid-activated signaling pathway, seed germination, |
| GOTERM_CC_DIRECT | plasma membrane, integral component of membrane, |
| GOTERM_MF_DIRECT | protein serine/threonine kinase activity, ATP binding, kinase activity, carbohydrate binding, |
| **At1g55020** | **lipoxygenase 1(LOX1)** |
| GOTERM_BP_DIRECT | fatty acid biosynthetic process, defense response, response to wounding, jasmonic acid biosynthetic process, response to abscisic acid, response to jasmonic acid, defense response to bacterium, incompatible interaction, lateral root formation, membrane disassembly, oxylipin biosynthetic process, lipid oxidation, growth, root development, |
| GOTERM_CC_DIRECT | cytoplasm, plastid, |
| GOTERM_MF_DIRECT | metal ion binding, linoleate 9S-lipoxygenase activity, |
| **At1g15010** | **mediator of RNA polymerase II transcription subunit(AT1G15010)** |
| GOTERM_BP_DIRECT | defense response to fungus, |
| GOTERM_CC_DIRECT | mitochondrion, integral component of membrane, |
| **At2g01300** | **mediator of RNA polymerase II transcription subunit(AT2G01300)** |
| GOTERM_CC_DIRECT | chloroplast, integral component of membrane, |
| **At3g13100** | **multidrug resistance-associated protein 7(ABCC7)** |
| GOTERM_BP_DIRECT | response to other organism, |
| GOTERM_CC_DIRECT | vacuolar membrane, integral component of membrane, |
| GOTERM_MF_DIRECT | ATP binding, xenobiotic-transporting ATPase activity, ATPase activity, coupled to transmembrane movement of substances, |
| **At3g13090** | **multidrug resistance-associated protein 8(ABCC6)** |
| GOTERM_CC_DIRECT | vacuolar membrane, plasmodesma, integral component of membrane, |
| GOTERM_MF_DIRECT | ATP binding, xenobiotic-transporting ATPase activity, ATPase activity, coupled to transmembrane movement of substances, |
| **At3g23250** | **myb domain protein 15(MYB15)** |
| GOTERM_BP_DIRECT | regulation of transcription, DNA-templated, regulation of transcription from RNA polymerase II promoter, response to salt stress, response to ethylene, response to auxin, response to jasmonic acid, response to chitin, cell differentiation, response to cadmium ion, |
| GOTERM_CC_DIRECT | nucleus, |
| GOTERM_MF_DIRECT | RNA polymerase II transcription factor activity, sequence-specific DNA binding, transcription factor activity, RNA polymerase II transcription factor recruiting, DNA binding, transcription factor activity, sequence-specific DNA binding, sequence-specific DNA binding, transcription regulatory region DNA binding, |
| **At4g17785** | **myb domain protein 39(MYB39)** |
| GOTERM_BP_DIRECT | regulation of transcription, DNA-templated, |
| GOTERM_CC_DIRECT | nucleus, |

| | | |
|---|---|---|
| | GOTERM_MF_DIRECT | DNA binding, transcription factor activity, sequence-specific DNA binding, |
| | **At1g17950** | **myb domain protein 52(MYB52)** |
| | GOTERM_BP_DIRECT | regulation of transcription, DNA-templated, regulation of transcription from RNA polymerase II promoter, response to abscisic acid, cell differentiation, regulation of secondary cell wall biogenesis, |
| | GOTERM_CC_DIRECT | nucleus, |
| | GOTERM_MF_DIRECT | RNA polymerase II transcription factor activity, sequence-specific DNA binding, transcription factor activity, RNA polymerase II transcription factor recruiting, DNA binding, transcription factor activity, sequence-specific DNA binding, sequence-specific DNA binding, transcription regulatory region DNA binding, |
| | **At3g12720** | **myb domain protein 67(MYB67)** |
| | GOTERM_BP_DIRECT | regulation of transcription, DNA-templated, regulation of transcription from RNA polymerase II promoter, cell differentiation, |
| | GOTERM_CC_DIRECT | nucleus, |
| | GOTERM_MF_DIRECT | RNA polymerase II transcription factor activity, sequence-specific DNA binding, transcription factor activity, RNA polymerase II transcription factor recruiting, DNA binding, transcription factor activity, sequence-specific DNA binding, sequence-specific DNA binding, transcription regulatory region DNA binding, |
| | **At2g47950** | **myelin transcription factor-like protein(AT2G47950)** |
| | GOTERM_CC_DIRECT | nucleus, |
| | **At4g26260** | **myo-inositol oxygenase 4(MIOX4)** |
| | GOTERM_BP_DIRECT | syncytium formation, inositol catabolic process, L-ascorbic acid biosynthetic process, oxidation-reduction process, |
| | GOTERM_CC_DIRECT | cytoplasm, |
| | GOTERM_MF_DIRECT | iron ion binding, inositol oxygenase activity, |
| | **At1g08100** | **nitrate transporter 2.2(NRT2.2)** |
| | GOTERM_BP_DIRECT | nitrate transport, nitrate assimilation, lateral root development, transmembrane transport, |
| | GOTERM_CC_DIRECT | plasma membrane, integral component of membrane, |
| | GOTERM_MF_DIRECT | nitrate transmembrane transporter activity, |
| | **At5g60770** | **nitrate transporter 2.4(NRT2.4)** |
| | GOTERM_BP_DIRECT | nitrate transport, nitrate assimilation, transmembrane transport, cellular response to nitrate, |
| | GOTERM_CC_DIRECT | plasma membrane, integral component of membrane, |
| | GOTERM_MF_DIRECT | nitrate transmembrane transporter activity, |
| | **At1g12940** | **nitrate transporter2.5(NRT2.5)** |
| | GOTERM_BP_DIRECT | nitrate transport, nitrate assimilation, transmembrane transport, |
| | GOTERM_CC_DIRECT | plasma membrane, integral component of membrane, |
| | GOTERM_MF_DIRECT | nitrate transmembrane transporter activity, |
| | **At3g03530** | **non-specific phospholipase C4(NPC4)** |
| | GOTERM_BP_DIRECT | phospholipid catabolic process, |
| | GOTERM_CC_DIRECT | plasma membrane, |

| | |
|---|---|
| GOTERM_MF_DIRECT | acid phosphatase activity, phospholipase C activity, hydrolase activity, acting on ester bonds, phosphatidylcholine phospholipase C activity, |
| **At2g04450** | **nudix hydrolase homolog 6(NUDT6)** |
| GOTERM_BP_DIRECT | response to other organism, positive regulation of salicylic acid mediated signaling pathway, |
| GOTERM_CC_DIRECT | cytosol, |
| GOTERM_MF_DIRECT | NAD+ diphosphatase activity, hydrolase activity, NADH pyrophosphatase activity, metal ion binding, ADP-ribose diphosphatase activity, NAD binding, |
| **At1g73220** | **organic cation/carnitine transporter1(OCT1)** |
| GOTERM_BP_DIRECT | ion transport, leaf senescence, cadaverine transport, |
| GOTERM_CC_DIRECT | plasma membrane, membrane, integral component of membrane, |
| GOTERM_MF_DIRECT | transporter activity, ATP binding, carbohydrate transmembrane transporter activity, carnitine transmembrane transporter activity, transmembrane transporter activity, |
| **At4g11650** | **osmotin 34(OSM34)** |
| GOTERM_BP_DIRECT | response to salt stress, defense response to bacterium, incompatible interaction, defense response to fungus, incompatible interaction, response to other organism, |
| GOTERM_CC_DIRECT | extracellular region, |
| **At1g75040** | **pathogenesis-related protein 5(PR5)** |
| GOTERM_BP_DIRECT | response to virus, systemic acquired resistance, response to UV-B, regulation of anthocyanin biosynthetic process, response to cadmium ion, response to other organism, |
| GOTERM_CC_DIRECT | extracellular region, cell wall, vacuole, apoplast, |
| **At5g66670** | **pectinesterase, putative (DUF677)(AT5G66670)** |
| GOTERM_CC_DIRECT | integral component of membrane, |
| **At3g49110** | **peroxidase CA(PRXCA)** |
| GOTERM_BP_DIRECT | defense response, response to oxidative stress, response to light stimulus, unidimensional cell growth, defense response to bacterium, hydrogen peroxide catabolic process, defense response to fungus, pathogen-associated molecular pattern dependent induction by symbiont of host innate immune response, oxidation-reduction process, reactive oxygen species metabolic process, |
| GOTERM_CC_DIRECT | extracellular region, vacuole, plant-type cell wall, |
| GOTERM_MF_DIRECT | peroxidase activity, heme binding, metal ion binding, |
| **At2g40180** | **phosphatase 2C5(PP2C5)** |
| GOTERM_BP_DIRECT | protein dephosphorylation, abscisic acid-activated signaling pathway, stomatal lineage progression, |
| GOTERM_CC_DIRECT | nucleus, |
| GOTERM_MF_DIRECT | protein serine/threonine phosphatase activity, metal ion binding, |
| **At3g48850** | **phosphate transporter 3;2(PHT3;2)** |
| GOTERM_BP_DIRECT | translation, transport, response to salt stress, |
| GOTERM_CC_DIRECT | mitochondrion, mitochondrial inner membrane, integral component of membrane, |
| GOTERM_MF_DIRECT | structural constituent of ribosome, |

| | |
|---|---|
| **At4g26270** | **phosphofructokinase 3(PFK3)** |
| GOTERM_BP_DIRECT | fructose 6-phosphate metabolic process, glycolytic process, root epidermal cell differentiation, |
| GOTERM_CC_DIRECT | cytoplasm, cytosol, 6-phosphofructokinase complex, |
| GOTERM_MF_DIRECT | 6-phosphofructokinase activity, ATP binding, metal ion binding, |
| **At3g60420** | **phosphoglycerate mutase family protein(AT3G60420)** |
| **At2g26560** | **phospholipase A 2A(PLA2A)** |
| GOTERM_BP_DIRECT | lipid metabolic process, cell death, plant-type hypersensitive response, lipid catabolic process, oxylipin biosynthetic process, response to cadmium ion, defense response to virus, cellular response to hypoxia, |
| GOTERM_CC_DIRECT | cytoplasm, chloroplast, membrane, |
| GOTERM_MF_DIRECT | phospholipase activity, lipase activity, nutrient reservoir activity, acylglycerol lipase activity, |
| **At5g54490** | **pinoid-binding protein 1(PBP1)** |
| GOTERM_BP_DIRECT | response to auxin, |
| GOTERM_CC_DIRECT | nucleus, |
| GOTERM_MF_DIRECT | calcium ion binding, protein binding, |
| **At2g18660** | **plant natriuretic peptide A(PNP-A)** |
| GOTERM_BP_DIRECT | amino sugar metabolic process, systemic acquired resistance, alternative respiration, |
| GOTERM_CC_DIRECT | extracellular region, cell wall, intracellular, apoplast, |
| GOTERM_MF_DIRECT | chitinase activity, |
| **At2g02850** | **plantacyanin(ARPN)** |
| GOTERM_BP_DIRECT | pollination, anther development, oxidation-reduction process, |
| GOTERM_CC_DIRECT | proteinaceous extracellular matrix, plasma membrane, extracellular matrix, anchored component of plasma membrane, apoplast, |
| GOTERM_MF_DIRECT | copper ion binding, electron carrier activity, metal ion binding, |
| **At5g37840** | **plastid movement impaired protein(AT5G37840)** |
| GOTERM_CC_DIRECT | nucleus, |
| **At1g76600** | **poly polymerase(AT1G76600)** |
| GOTERM_CC_DIRECT | nucleus, nucleolus, |
| **At5g06860** | **polygalacturonase inhibiting protein 1(PGIP1)** |
| GOTERM_BP_DIRECT | defense response, signal transduction, |
| GOTERM_CC_DIRECT | extracellular region, cell wall, Golgi apparatus, cytosol, plant-type cell wall, plasmodesma, membrane, |
| **At3g47420** | **putative glycerol-3-phosphate transporter 1(G3Pp1)** |
| GOTERM_BP_DIRECT | anion transport, carbohydrate transport, phosphate ion homeostasis, transmembrane transport, |
| GOTERM_CC_DIRECT | integral component of membrane, |

| | |
|---|---|
| GOTERM_MF_DIRECT | sugar:proton symporter activity, transmembrane transporter activity, |
| **At4g18250** | **receptor Serine/Threonine kinase-like protein(AT4G18250)** |
| GOTERM_BP_DIRECT | protein phosphorylation, |
| GOTERM_CC_DIRECT | integral component of membrane, |
| GOTERM_MF_DIRECT | protein kinase activity, transmembrane receptor protein serine/threonine kinase activity, ATP binding, kinase activity, |
| **At4g21380** | **receptor kinase 3(RK3)** |
| GOTERM_BP_DIRECT | protein phosphorylation, defense response, recognition of pollen, |
| GOTERM_CC_DIRECT | vacuole, plasma membrane, plasmodesma, integral component of membrane, |
| GOTERM_MF_DIRECT | protein serine/threonine kinase activity, transmembrane receptor protein serine/threonine kinase activity, ATP binding, kinase activity, carbohydrate binding, ubiquitin protein ligase binding, |
| **At3g23110** | **receptor like protein 37(RLP37)** |
| GOTERM_BP_DIRECT | defense response, signal transduction, |
| GOTERM_CC_DIRECT | plasma membrane, integral component of membrane, |
| GOTERM_MF_DIRECT | kinase activity, |
| **At1g47890** | **receptor like protein 7(RLP7)** |
| GOTERM_BP_DIRECT | defense response, signal transduction, |
| GOTERM_CC_DIRECT | integral component of membrane, |
| GOTERM_MF_DIRECT | kinase activity, |
| **At5g48540** | **receptor-like protein kinase-related family protein(AT5G48540)** |
| GOTERM_BP_DIRECT | response to karrikin, |
| GOTERM_CC_DIRECT | extracellular region, |
| **At5g13330** | **related to AP2 6l(Rap2.6L)** |
| GOTERM_BP_DIRECT | transcription, DNA-templated, regulation of transcription, DNA-templated, response to water deprivation, response to salt stress, response to ethylene, response to abscisic acid, response to salicylic acid, response to jasmonic acid, ethylene-activated signaling pathway, glucosinolate metabolic process, positive regulation of transcription, DNA-templated, cellular response to freezing, |
| GOTERM_CC_DIRECT | nucleus, |
| GOTERM_MF_DIRECT | DNA binding, transcription factor activity, sequence-specific DNA binding, sequence-specific DNA binding, |
| **At4g17670** | **senescence-associated family protein (DUF581)(AT4G17670)** |
| GOTERM_CC_DIRECT | nucleus, |
| **At3g56710** | **sigma factor binding protein 1(SIB1)** |
| GOTERM_BP_DIRECT | defense response to bacterium, incompatible interaction, positive regulation of sequence-specific DNA binding transcription factor activity, cellular response to light stimulus, |
| GOTERM_CC_DIRECT | nucleus, chloroplast, |

| | | |
|---|---|---|
| | GOTERM_MF_DIRECT | protein binding, |
| | **At5g10180** | **slufate transporter 2;1(SULTR2;1)** |
| | GOTERM_BP_DIRECT | sulfate transport, sulfate transmembrane transport, |
| | GOTERM_CC_DIRECT | integral component of plasma membrane, |
| | GOTERM_MF_DIRECT | secondary active sulfate transmembrane transporter activity, sulfate transmembrane transporter activity, symporter activity, |
| | **At1g51420** | **sucrose-phosphatase 1(SPP1)** |
| | GOTERM_BP_DIRECT | sucrose biosynthetic process, |
| | GOTERM_CC_DIRECT | vacuolar membrane, |
| | GOTERM_MF_DIRECT | magnesium ion binding, sucrose-phosphate phosphatase activity, |
| | **At1g22150** | **sulfate transporter 1;3(SULTR1;3)** |
| | GOTERM_CC_DIRECT | integral component of plasma membrane, |
| | GOTERM_MF_DIRECT | secondary active sulfate transmembrane transporter activity, sulfate transmembrane transporter activity, symporter activity, |
| | **At2g03760** | **sulfotransferase 12(SOT12)** |
| | GOTERM_BP_DIRECT | defense response, response to salt stress, response to salicylic acid, brassinosteroid metabolic process, |
| | GOTERM_CC_DIRECT | cytoplasm, Golgi apparatus, |
| | GOTERM_MF_DIRECT | sulfotransferase activity, brassinosteroid sulfotransferase activity, flavonoid sulfotransferase activity, |
| | **At1g67810** | **sulfur E2(SUFE2)** |
| | GOTERM_BP_DIRECT | iron-sulfur cluster assembly, positive regulation of sulfur metabolic process, |
| | GOTERM_CC_DIRECT | chloroplast, |
| | GOTERM_MF_DIRECT | enzyme activator activity, |
| | **At5g66170** | **sulfurtransferase 18(STR18)** |
| | GOTERM_CC_DIRECT | cytoplasm, chloroplast, |
| | GOTERM_MF_DIRECT | thiosulfate sulfurtransferase activity, transferase activity, |
| | **At3g25820** | **terpene synthase-like sequence-1,8-cineole(TPS-CIN)** |
| | GOTERM_BP_DIRECT | metabolic process, monoterpenoid biosynthetic process, terpenoid biosynthetic process, root development, defense response to fungus, |
| | GOTERM_CC_DIRECT | chloroplast, |
| | GOTERM_MF_DIRECT | magnesium ion binding, terpene synthase activity, (E)-beta-ocimene synthase activity, myrcene synthase activity, 1,8-cineole synthase activity, |
| | **At1g69880** | **thioredoxin H-type 8(TH8)** |
| | GOTERM_BP_DIRECT | sulfate assimilation, protein folding, glycerol ether metabolic process, cellular response to oxidative stress, cell redox homeostasis, oxidation-reduction process, |
| | GOTERM_CC_DIRECT | cytoplasm, cytosol, plasma membrane, |
| | GOTERM_MF_DIRECT | thioredoxin-disulfide reductase activity, protein disulfide oxidoreductase activity, oxidoreductase activity, acting on a sulfur group of donors, disulfide as acceptor, protein-disulfide reductase activity, |

| | | |
|---|---|---|
| | **At1g19960** | **transcription factor(AT1G19960)** |
| | **At3g27070** | **translocase outer membrane 20-1(TOM20-1)** |
| | GOTERM_BP_DIRECT | protein targeting to mitochondrion, protein import into mitochondrial outer membrane, protein transmembrane transport, |
| | GOTERM_CC_DIRECT | mitochondrion, mitochondrial outer membrane translocase complex, mitochondrial inner membrane presequence translocase complex, integral component of membrane, |
| | GOTERM_MF_DIRECT | P-P-bond-hydrolysis-driven protein transmembrane transporter activity, |
| | **At1g20180** | **transmembrane protein (DUF677)(AT1G20180)** |
| | GOTERM_CC_DIRECT | mitochondrion, integral component of membrane, |
| | **At1g25400** | **transmembrane protein(AT1G25400)** |
| | **At1g51920** | **transmembrane protein(AT1G51920)** |
| | **At1g65500** | **transmembrane protein(AT1G65500)** |
| | **At2g18690** | **transmembrane protein(AT2G18690)** |
| | GOTERM_CC_DIRECT | chloroplast, membrane, integral component of membrane, |
| | **At2g25510** | **transmembrane protein(AT2G25510)** |
| | GOTERM_CC_DIRECT | mitochondrion, integral component of membrane, |
| | **At2g31945** | **transmembrane protein(AT2G31945)** |
| | **At3g55790** | **transmembrane protein(AT3G55790)** |
| | GOTERM_BP_DIRECT | cellular response to hypoxia, |
| | GOTERM_CC_DIRECT | extracellular region, anchored component of membrane, |
| | **At4g28460** | **transmembrane protein(AT4G28460)** |
| | GOTERM_CC_DIRECT | extracellular region, apoplast, |
| | **At4g37290** | **transmembrane protein(AT4G37290)** |
| | GOTERM_BP_DIRECT | response to karrikin, |
| | GOTERM_CC_DIRECT | mitochondrion, apoplast, |
| | **At5g10040** | **transmembrane protein(AT5G10040)** |
| | GOTERM_BP_DIRECT | anaerobic respiration, |
| | GOTERM_CC_DIRECT | integral component of membrane, |
| | **At5g44580** | **transmembrane protein(AT5G44580)** |
| | GOTERM_CC_DIRECT | mitochondrion, |
| | **At5g48175** | **transmembrane protein(AT5G48175)** |
| | GOTERM_CC_DIRECT | integral component of membrane, |
| | **At3g60470** | **transmembrane protein, putative (DUF247)(AT3G60470)** |

| | | |
|---|---|---|
| | GOTERM_CC_DIRECT | plasma membrane, integral component of membrane, |
| | **At3g21520** | **transmembrane protein, putative (DUF679 domain membrane protein 1)(DMP1)** |
| | GOTERM_BP_DIRECT | endomembrane system organization, |
| | GOTERM_CC_DIRECT | endoplasmic reticulum, chloroplast, plant-type vacuole membrane, integral component of membrane, |
| | **At2g32140** | **transmembrane receptor(AT2G32140)** |
| | GOTERM_BP_DIRECT | defense response, signal transduction, |
| | GOTERM_CC_DIRECT | chloroplast, integral component of membrane, |
| | **At5g39040** | **transporter associated with antigen processing protein 2(ABCB27)** |
| | GOTERM_BP_DIRECT | response to aluminum ion, oligopeptide transmembrane transport, transmembrane transport, |
| | GOTERM_CC_DIRECT | plant-type vacuole, vacuole, vacuolar membrane, plasma membrane, integral component of membrane, |
| | GOTERM_MF_DIRECT | transporter activity, ATP binding, oligopeptide-transporting ATPase activity, ATPase activity, coupled to transmembrane movement of substances, |
| | **At2g43510** | **trypsin inhibitor protein 1(TI1)** |
| | GOTERM_BP_DIRECT | defense response, killing of cells of other organism, defense response to fungus, |
| | GOTERM_CC_DIRECT | extracellular region, |
| | GOTERM_MF_DIRECT | serine-type endopeptidase inhibitor activity, ion channel inhibitor activity, |
| | **At5g45380** | **urea-proton symporter DEGRADATION OF UREA 3 (DUR3)(DUR3)** |
| | GOTERM_BP_DIRECT | cellular response to nitrogen starvation, urea transmembrane transport, |
| | GOTERM_CC_DIRECT | plasma membrane, integral component of plasma membrane, integral component of membrane, |
| | GOTERM_MF_DIRECT | urea transmembrane transporter activity, symporter activity, solute:sodium symporter activity, |
| | **At1g21270** | **wall-associated kinase 2(WAK2)** |
| | GOTERM_BP_DIRECT | protein phosphorylation, cell surface receptor signaling pathway, oligosaccharide metabolic process, response to salicylic acid, unidimensional cell growth, cellular water homeostasis, |
| | GOTERM_CC_DIRECT | cell, plasma membrane, integral component of membrane, |
| | GOTERM_MF_DIRECT | protein serine/threonine kinase activity, calcium ion binding, ATP binding, polysaccharide binding, |
| | **At3g28210** | **zinc finger (AN1-like) family protein(PMZ)** |
| | GOTERM_BP_DIRECT | response to abscisic acid, response to chitin, |
| | GOTERM_CC_DIRECT | nucleus, |
| | GOTERM_MF_DIRECT | zinc ion binding, |
