## Supplementary material for "Evidences for a nutritional role of iodine in plants": Table S7

| AGI code | Name |
|---|---|
| **At3g11150** | **2-oxoglutarate (2OG) and Fe(II)-dependent oxygenase superfamily protein(AT3G11150)** |
| GOTERM_CC_DIRECT | chloroplast, |
| **At3g07490** | **ARF-GAP domain 11(AGD11)** |
| GOTERM_CC_DIRECT | peroxisome, plasma membrane, |
| GOTERM_MF_DIRECT | calcium ion binding, |
| **At5g43590** | **Acyl transferase/acyl hydrolase/lysophospholipase superfamily protein(AT5G43590)** |
| GOTERM_BP_DIRECT | defense response, lipid catabolic process, |
| GOTERM_CC_DIRECT | nucleus, cytoplasm, membrane, |
| GOTERM_MF_DIRECT | phospholipase activity, transferase activity, nutrient reservoir activity, acylglycerol lipase activity, |
| **At2g03720** | **Adenine nucleotide alpha hydrolases-like superfamily protein(MRH6)** |
| GOTERM_BP_DIRECT | response to stress, root hair cell differentiation, |
| **At3g14415** | **Aldolase-type TIM barrel family protein(GOX2)** |
| GOTERM_BP_DIRECT | oxidative photosynthetic carbon pathway, defense response to bacterium, hydrogen peroxide biosynthetic process, oxidation-reduction process, |
| GOTERM_CC_DIRECT | nucleus, vacuole, peroxisome, chloroplast, chloroplast stroma, apoplast, |
| GOTERM_MF_DIRECT | glycolate oxidase activity, FMN binding, oxidoreductase activity, very-long-chain-(S)-2-hydroxy-acid oxidase activity, long-chain-(S)-2-hydroxy-long-chain-acid oxidase activity, medium-chain-(S)-2-hydroxy-acid oxidase activity, |
| **At5g04730** | **Ankyrin-repeat containing protein(AT5G04730)** |
| GOTERM_BP_DIRECT | signal transduction, |
| GOTERM_CC_DIRECT | membrane, integral component of membrane, |
| **At4g12550** | **Auxin-Induced in Root cultures 1(AIR1)** |
| GOTERM_BP_DIRECT | lipid transport, response to auxin, lateral root morphogenesis, |
| GOTERM_CC_DIRECT | extracellular region, |
| GOTERM_MF_DIRECT | lipid binding, |
| **At1g23160** | **Auxin-responsive GH3 family protein(AT1G23160)** |
| GOTERM_BP_DIRECT | response to auxin, |
| GOTERM_CC_DIRECT | cytoplasm, |
| **At1g62510** | **Bifunctional inhibitor/lipid-transfer protein/seed storage 2S albumin superfamily protein(AT1G62510)** |
| GOTERM_BP_DIRECT | lipid transport, |
| GOTERM_MF_DIRECT | lipid binding, |
| **At3g22570** | **Bifunctional inhibitor/lipid-transfer protein/seed storage 2S albumin superfamily protein(AT3G22570)** |
| GOTERM_BP_DIRECT | proteolysis, lipid transport, |

| | | |
|---|---|---|
| | GOTERM_CC_DIRECT | extracellular region, |
| | GOTERM_MF_DIRECT | peptidase activity, lipid binding, |
| | **At4g12360** | **Bifunctional inhibitor/lipid-transfer protein/seed storage 2S albumin superfamily protein(AT4G12360)** |
| | GOTERM_BP_DIRECT | lipid transport, |
| | GOTERM_CC_DIRECT | chloroplast, integral component of membrane, anchored component of membrane, |
| | GOTERM_MF_DIRECT | lipid binding, |
| | **At4g25790** | **CAP (Cysteine-rich secretory proteins, Antigen 5, and Pathogenesis-related 1 protein) superfamily protein(AT4G25790)** |
| | GOTERM_CC_DIRECT | extracellular region, |
| | **At4g30320** | **CAP (Cysteine-rich secretory proteins, Antigen 5, and Pathogenesis-related 1 protein) superfamily protein(AT4G30320)** |
| | GOTERM_CC_DIRECT | extracellular region, |
| | **At4g33730** | **CAP (Cysteine-rich secretory proteins, Antigen 5, and Pathogenesis-related 1 protein) superfamily protein(AT4G33730)** |
| | GOTERM_BP_DIRECT | negative regulation of response to salt stress, |
| | GOTERM_CC_DIRECT | extracellular region, |
| | **At5g26130** | **CAP (Cysteine-rich secretory proteins, Antigen 5, and Pathogenesis-related 1 protein) superfamily protein(AT5G26130)** |
| | GOTERM_CC_DIRECT | extracellular region, |
| | **At2g34180** | **CBL-interacting protein kinase 13(CIPK13)** |
| | GOTERM_BP_DIRECT | protein phosphorylation, signal transduction, |
| | GOTERM_CC_DIRECT | plasma membrane, |
| | GOTERM_MF_DIRECT | protein serine/threonine kinase activity, protein binding, ATP binding, kinase activity, |
| | **At2g25090** | **CBL-interacting protein kinase 16(CIPK16)** |
| | GOTERM_BP_DIRECT | protein phosphorylation, sodium ion transport, signal transduction, hyperosmotic salinity response, |
| | GOTERM_CC_DIRECT | plasma membrane, |
| | GOTERM_MF_DIRECT | protein serine/threonine kinase activity, ATP binding, kinase activity, |
| | **At2g26980** | **CBL-interacting protein kinase 3(CIPK3)** |
| | GOTERM_BP_DIRECT | protein phosphorylation, signal transduction, response to cytokinin, response to abscisic acid, abscisic acid-activated signaling pathway, intracellular signal transduction, |
| | GOTERM_CC_DIRECT | intracellular, nucleus, cytoplasm, plasma membrane, |
| | GOTERM_MF_DIRECT | protein kinase activity, protein serine/threonine kinase activity, ATP binding, kinase activity, |
| | **At4g18510** | **CLAVATA3/ESR-related 2(CLE2)** |
| | GOTERM_BP_DIRECT | multicellular organism development, cell-cell signaling involved in cell fate commitment, |
| | GOTERM_CC_DIRECT | extracellular region, extracellular space, apoplast, |
| | GOTERM_MF_DIRECT | receptor serine/threonine kinase binding, |

| | | |
|---|---|---|
| | **At3g29810** | **COBRA-like protein 2 precursor(COBL2)** |
| | GOTERM_BP_DIRECT | plant-type cell wall organization, plant-type cell wall biogenesis, seed coat development, cellulose microfibril organization, cell growth, seed development, mucilage biosynthetic process involved in seed coat development, plant-type cell wall cellulose biosynthetic process, |
| | GOTERM_CC_DIRECT | Golgi apparatus, plasma membrane, anchored component of membrane, anchored component of plasma membrane, |
| | **At5g54790** | **CTD small phosphatase-like protein(AT5G54790)** |
| | GOTERM_CC_DIRECT | nucleus, |
| | **At5g46370** | **Ca2+ activated outward rectifying K+ channel 2(KCO2)** |
| | GOTERM_BP_DIRECT | thiamine diphosphate biosynthetic process, phosphorylation, potassium ion transmembrane transport, |
| | GOTERM_CC_DIRECT | nucleus, plant-type vacuole membrane, integral component of membrane, |
| | GOTERM_MF_DIRECT | potassium channel activity, calcium ion binding, ATP binding, outward rectifier potassium channel activity, kinase activity, thiamine binding, metal ion binding, |
| | **At1g66860** | **Class I glutamine amidotransferase-like superfamily protein(AT1G66860)** |
| | GOTERM_BP_DIRECT | glutamine metabolic process, |
| | GOTERM_CC_DIRECT | nucleus, cytosol, |
| | GOTERM_MF_DIRECT | transferase activity, hydrolase activity, |
| | **At4g37220** | **Cold acclimation protein WCOR413 family(AT4G37220)** |
| | GOTERM_BP_DIRECT | response to sucrose, response to glucose, response to fructose, |
| | GOTERM_CC_DIRECT | mitochondrion, plasma membrane, integral component of membrane, |
| | **At1g55440** | **Cysteine/Histidine-rich C1 domain family protein(AT1G55440)** |
| | GOTERM_CC_DIRECT | nucleus, |
| | GOTERM_MF_DIRECT | zinc ion binding, |
| | **At2g01900** | **DNAse I-like superfamily protein(AT2G01900)** |
| | GOTERM_BP_DIRECT | phosphatidylinositol dephosphorylation, cellular response to salt stress, regulation of clathrin-mediated endocytosis, regulation of reactive oxygen species metabolic process, |
| | GOTERM_CC_DIRECT | nucleus, |
| | GOTERM_MF_DIRECT | phosphatidylinositol-4,5-bisphosphate 5-phosphatase activity, hydrolase activity, phosphatidylinositol-3,4,5-trisphosphate 5-phosphatase activity, phosphatidylinositol phosphate 5-phosphatase activity, |
| | **At2g46220** | **DUF2358 family protein (DUF2358)(AT2G46220)** |
| | GOTERM_CC_DIRECT | chloroplast, |
| | **At4g11190** | **Disease resistance-responsive (dirigent-like protein) family protein(AT4G11190)** |
| | GOTERM_BP_DIRECT | defense response, phenylpropanoid biosynthetic process, lignan biosynthetic process, |
| | GOTERM_CC_DIRECT | extracellular region, apoplast, |
| | GOTERM_MF_DIRECT | guiding stereospecific synthesis activity, |
| | **At3g46240** | **ER protein carbohydrate-binding protein(AT3G46240)** |

| | | |
|---|---|---|
| | GOTERM_CC_DIRECT | integral component of membrane, |
| | **At2g20520** | **FASCICLIN-like arabinogalactan 6(FLA6)** |
| | GOTERM_CC_DIRECT | plasma membrane, integral component of membrane, anchored component of membrane, |
| | **At5g19970** | **GRAS family transcription factor family protein(AT5G19970)** |
| | GOTERM_CC_DIRECT | mitochondrion, |
| | **At3g47170** | **HXXXD-type acyl-transferase family protein(AT3G47170)** |
| | GOTERM_CC_DIRECT | cytoplasm, |
| | GOTERM_MF_DIRECT | transferase activity, transferase activity, transferring acyl groups other than amino-acyl groups, |
| | **At5g47950** | **HXXXD-type acyl-transferase family protein(AT5G47950)** |
| | GOTERM_CC_DIRECT | cytoplasm, |
| | GOTERM_MF_DIRECT | transferase activity, transferase activity, transferring acyl groups other than amino-acyl groups, |
| | **At5g47980** | **HXXXD-type acyl-transferase family protein(AT5G47980)** |
| | GOTERM_CC_DIRECT | cytoplasm, |
| | GOTERM_MF_DIRECT | transferase activity, transferase activity, transferring acyl groups, transferase activity, transferring acyl groups other than amino-acyl groups, |
| | **At2g22190** | **Haloacid dehalogenase-like hydrolase (HAD) superfamily protein(TPPE)** |
| | GOTERM_BP_DIRECT | trehalose biosynthetic process, trehalose metabolism in response to stress, |
| | GOTERM_CC_DIRECT | cytoplasm, chloroplast, |
| | GOTERM_MF_DIRECT | trehalose-phosphatase activity, |
| | **At4g39770** | **Haloacid dehalogenase-like hydrolase (HAD) superfamily protein(TPPH)** |
| | GOTERM_BP_DIRECT | trehalose biosynthetic process, trehalose metabolism in response to stress, |
| | GOTERM_CC_DIRECT | nucleus, cytoplasm, cytosol, |
| | GOTERM_MF_DIRECT | trehalose-phosphatase activity, |
| | **At5g61590** | **Integrase-type DNA-binding superfamily protein(AT5G61590)** |
| | GOTERM_BP_DIRECT | transcription, DNA-templated, regulation of transcription, DNA-templated, response to water deprivation, ethylene-activated signaling pathway, glucosinolate metabolic process, positive regulation of transcription, DNA-templated, negative regulation of wax biosynthetic process, |
| | GOTERM_CC_DIRECT | nucleus, |
| | GOTERM_MF_DIRECT | transcription regulatory region sequence-specific DNA binding, DNA binding, transcription factor activity, sequence-specific DNA binding, sequence-specific DNA binding, transcription regulatory region DNA binding, |
| | **At4g37540** | **LOB domain-containing protein 39(LBD39)** |
| | **At2g34330** | **LOW protein: protein BOBBER-like protein(AT2G34330)** |
| | GOTERM_CC_DIRECT | mitochondrion, |
| | **At1g64210** | **Leucine-rich repeat protein kinase family protein(AT1G64210)** |

| | | |
|---|---|---|
| | GOTERM_BP_DIRECT | protein phosphorylation, transmembrane receptor protein tyrosine kinase signaling pathway, |
| | GOTERM_CC_DIRECT | mitochondrion, plasma membrane, integral component of membrane, |
| | GOTERM_MF_DIRECT | protein kinase activity, protein serine/threonine kinase activity, ATP binding, kinase activity, |
| | **At5g59650** | **Leucine-rich repeat protein kinase family protein(AT5G59650)** |
| | GOTERM_CC_DIRECT | plasma membrane, integral component of membrane, |
| | GOTERM_MF_DIRECT | protein serine/threonine kinase activity, ATP binding, kinase activity, |
| | **At2g19210** | **Leucine-rich repeat transmembrane protein kinase protein(AT2G19210)** |
| | GOTERM_BP_DIRECT | protein phosphorylation, |
| | GOTERM_CC_DIRECT | plasma membrane, integral component of membrane, |
| | GOTERM_MF_DIRECT | protein serine/threonine kinase activity, ATP binding, kinase activity, |
| | **At1g34490** | **MBOAT (membrane bound O-acyl transferase) family protein(AT1G34490)** |
| | GOTERM_BP_DIRECT | lipid metabolic process, |
| | GOTERM_CC_DIRECT | integral component of membrane, |
| | GOTERM_MF_DIRECT | transferase activity, transferring acyl groups, long-chain-alcohol O-fatty-acyltransferase activity, wax ester synthase activity, |
| | **At5g23840** | **MD-2-related lipid recognition domain-containing protein(AT5G23840)** |
| | GOTERM_CC_DIRECT | extracellular region, |
| | **At2g01530** | **MLP-like protein 329(MLP329)** |
| | GOTERM_BP_DIRECT | defense response, response to biotic stimulus, response to cytokinin, |
| | GOTERM_CC_DIRECT | nucleus, |
| | GOTERM_MF_DIRECT | copper ion binding, |
| | **At1g70890** | **MLP-like protein 43(MLP43)** |
| | GOTERM_BP_DIRECT | defense response, response to biotic stimulus, |
| | GOTERM_CC_DIRECT | chloroplast, |
| | **At3g20460** | **Major facilitator superfamily protein(AT3G20460)** |
| | GOTERM_BP_DIRECT | hexose transmembrane transport, glucose import, |
| | GOTERM_CC_DIRECT | plasma membrane, integral component of plasma membrane, membrane, |
| | GOTERM_MF_DIRECT | sugar:proton symporter activity, glucose transmembrane transporter activity, carbohydrate transmembrane transporter activity, |
| | **At3g45700** | **Major facilitator superfamily protein(AT3G45700)** |
| | GOTERM_BP_DIRECT | oligopeptide transport, nitrate assimilation, chloride transmembrane transport, |
| | GOTERM_CC_DIRECT | plasma membrane, membrane, integral component of membrane, |
| | GOTERM_MF_DIRECT | transporter activity, chloride transmembrane transporter activity, |

| | | |
|---|---|---|
| | **At3g45710** | **Major facilitator superfamily protein(AT3G45710)** |
| | GOTERM_BP_DIRECT | oligopeptide transport, nitrate assimilation, |
| | GOTERM_CC_DIRECT | plasma membrane, membrane, integral component of membrane, |
| | GOTERM_MF_DIRECT | transporter activity, |
| | **At5g14120** | **Major facilitator superfamily protein(AT5G14120)** |
| | GOTERM_CC_DIRECT | vacuole, vacuolar membrane, integral component of membrane, |
| | **At3g29035** | **NAC domain containing protein 3(NAC3)** |
| | GOTERM_BP_DIRECT | transcription, DNA-templated, multicellular organism development, response to wounding, leaf senescence, response to hydrogen peroxide, positive regulation of sequence-specific DNA binding transcription factor activity, positive regulation of leaf senescence, |
| | GOTERM_CC_DIRECT | nucleus, |
| | GOTERM_MF_DIRECT | transcription factor activity, sequence-specific DNA binding, sequence-specific DNA binding, protein heterodimerization activity, |
| | **At5g07680** | **NAC domain containing protein 80(NAC080)** |
| | GOTERM_BP_DIRECT | transcription, DNA-templated, regulation of transcription, DNA-templated, multicellular organism development, |
| | GOTERM_CC_DIRECT | nucleus, |
| | GOTERM_MF_DIRECT | DNA binding, transcription factor activity, sequence-specific DNA binding, |
| | **At1g66800** | **NAD(P)-binding Rossmann-fold superfamily protein(AT1G66800)** |
| | GOTERM_BP_DIRECT | lignin biosynthetic process, |
| | GOTERM_MF_DIRECT | alcohol dehydrogenase (NAD) activity, cinnamyl-alcohol dehydrogenase activity, coenzyme binding, |
| | **At5g01740** | **Nuclear transport factor 2 (NTF2) family protein(AT5G01740)** |
| | **At1g33930** | **P-loop containing nucleoside triphosphate hydrolases superfamily protein(AT1G33930)** |
| | GOTERM_BP_DIRECT | response to bacterium, |
| | GOTERM_CC_DIRECT | mitochondrion, |
| | GOTERM_MF_DIRECT | GTP binding, |
| | **At3g18460** | **PLAC8 family protein(AT3G18460)** |
| | GOTERM_CC_DIRECT | plasma membrane, integral component of membrane, |
| | **At1g11920** | **Pectin lyase-like superfamily protein(AT1G11920)** |
| | GOTERM_BP_DIRECT | pectin catabolic process, |
| | GOTERM_CC_DIRECT | extracellular region, |
| | GOTERM_MF_DIRECT | lyase activity, pectate lyase activity, metal ion binding, |
| | **At1g34510** | **Peroxidase superfamily protein(AT1G34510)** |
| | GOTERM_BP_DIRECT | response to oxidative stress, plant-type cell wall organization, hydrogen peroxide catabolic process, oxidation-reduction process, |

| | | |
|---|---|---|
| | GOTERM_CC_DIRECT | extracellular region, plant-type cell wall, |
| | GOTERM_MF_DIRECT | peroxidase activity, heme binding, metal ion binding, |
| | **At5g62360** | **Plant invertase/pectin methylesterase inhibitor superfamily protein(AT5G62360)** |
| | GOTERM_BP_DIRECT | negative regulation of catalytic activity, |
| | GOTERM_MF_DIRECT | enzyme inhibitor activity, pectinesterase inhibitor activity, |
| | **At1g70880** | **Polyketide cyclase/dehydrase and lipid transport superfamily protein(AT1G70880)** |
| | GOTERM_BP_DIRECT | defense response, response to biotic stimulus, |
| | GOTERM_CC_DIRECT | nucleus, |
| | **At5g28010** | **Polyketide cyclase/dehydrase and lipid transport superfamily protein(AT5G28010)** |
| | GOTERM_BP_DIRECT | defense response, response to biotic stimulus, |
| | GOTERM_CC_DIRECT | cytoplasm, |
| | **At3g54580** | **Proline-rich extensin-like family protein(AT3G54580)** |
| | GOTERM_BP_DIRECT | plant-type cell wall organization, |
| | GOTERM_MF_DIRECT | structural constituent of cell wall, |
| | **At4g08410** | **Proline-rich extensin-like family protein(AT4G08410)** |
| | GOTERM_BP_DIRECT | plant-type cell wall organization, |
| | GOTERM_MF_DIRECT | structural constituent of cell wall, |
| | **At5g19810** | **Proline-rich extensin-like family protein(AT5G19810)** |
| | **At3g07070** | **Protein kinase superfamily protein(AT3G07070)** |
| | GOTERM_BP_DIRECT | protein phosphorylation, |
| | GOTERM_CC_DIRECT | plasma membrane, |
| | GOTERM_MF_DIRECT | protein serine/threonine kinase activity, ATP binding, kinase activity, |
| | **At1g35625** | **RING/U-box superfamily protein(AT1G35625)** |
| | GOTERM_BP_DIRECT | protein transport, |
| | GOTERM_CC_DIRECT | integral component of membrane, late endosome membrane, protein storage vacuole membrane, |
| | GOTERM_MF_DIRECT | peptidase activity, zinc ion binding, |
| | **At1g53820** | **RING/U-box superfamily protein(AT1G53820)** |
| | GOTERM_BP_DIRECT | protein ubiquitination, proteasome-mediated ubiquitin-dependent protein catabolic process, |
| | GOTERM_CC_DIRECT | nucleus, integral component of membrane, |
| | GOTERM_MF_DIRECT | zinc ion binding, ubiquitin protein ligase activity, |
| | **At4g09130** | **RING/U-box superfamily protein(AT4G09130)** |

| | | |
|---|---|---|
| | GOTERM_BP_DIRECT | protein ubiquitination, |
| | GOTERM_CC_DIRECT | extracellular region, integral component of membrane, |
| | GOTERM_MF_DIRECT | zinc ion binding, |
| | **At1g22500** | **RING/U-box superfamily protein(ATL15)** |
| | GOTERM_BP_DIRECT | response to light stimulus, protein ubiquitination, response to L-ascorbic acid, |
| | GOTERM_CC_DIRECT | extracellular region, integral component of membrane, |
| | GOTERM_MF_DIRECT | ubiquitin-protein transferase activity, zinc ion binding, ligase activity, |
| | **At3g48940** | **Remorin family protein(AT3G48940)** |
| | GOTERM_CC_DIRECT | plasma membrane, |
| | GOTERM_MF_DIRECT | DNA binding, |
| | **At4g33110** | **S-adenosyl-L-methionine-dependent methyltransferases superfamily protein(AT4G33110)** |
| | GOTERM_BP_DIRECT | methylation, |
| | GOTERM_CC_DIRECT | cytoplasm, endosome, vacuolar membrane, trans-Golgi network, plasma membrane, |
| | GOTERM_MF_DIRECT | S-adenosylmethionine-dependent methyltransferase activity, (S)-coclaurine-N-methyltransferase activity, |
| | **At2g21210** | **SAUR-like auxin-responsive protein family(AT2G21210)** |
| | GOTERM_BP_DIRECT | response to auxin, response to chitin, |
| | GOTERM_CC_DIRECT | chloroplast, |
| | **At5g15600** | **SPIRAL1-like4(SP1L4)** |
| | GOTERM_CC_DIRECT | microtubule, |
| | **At4g34580** | **Sec14p-like phosphatidylinositol transfer family protein(COW1)** |
| | GOTERM_BP_DIRECT | transport, cell tip growth, root epidermal cell differentiation, protein transport, root hair elongation, root hair cell tip growth, |
| | GOTERM_CC_DIRECT | Golgi membrane, intracellular, mitochondrion, plasma membrane, root hair tip, |
| | GOTERM_MF_DIRECT | transporter activity, phosphatidylinositol transporter activity, |
| | **At4g28410** | **Tyrosine transaminase family protein(AT4G28410)** |
| | GOTERM_BP_DIRECT | cellular amino acid metabolic process, biosynthetic process, |
| | GOTERM_CC_DIRECT | cytoplasm, |
| | GOTERM_MF_DIRECT | transaminase activity, pyridoxal phosphate binding, |
| | **At3g58000** | **VQ motif-containing protein(AT3G58000)** |
| | GOTERM_BP_DIRECT | defense response, |
| | GOTERM_CC_DIRECT | nucleus, |
| | **At5g39630** | **Vesicle transport v-SNARE family protein(AT5G39630)** |

| | | |
|---|---|---|
| | GOTERM_BP_DIRECT | protein targeting to vacuole, ER to Golgi vesicle-mediated transport, intra-Golgi vesicle-mediated transport, Golgi to vacuole transport, retrograde transport, endosome to Golgi, vesicle fusion with Golgi apparatus, |
| | GOTERM_CC_DIRECT | endoplasmic reticulum membrane, Golgi apparatus, cytosol, ER to Golgi transport vesicle membrane, SNARE complex, late endosome membrane, |
| | GOTERM_MF_DIRECT | SNARE binding, receptor activity, SNAP receptor activity, |
| | **At2g25900** | **Zinc finger C-x8-C-x5-C-x3-H type family protein(ATCTH)** |
| | GOTERM_BP_DIRECT | regulation of transcription, DNA-templated, mRNA destabilization, |
| | GOTERM_CC_DIRECT | nucleus, cytoplasm, |
| | GOTERM_MF_DIRECT | DNA binding, transcription factor activity, sequence-specific DNA binding, RNA binding, single-stranded RNA binding, metal ion binding, |
| | **At2g13360** | **alanine:glyoxylate aminotransferase(AGT)** |
| | GOTERM_BP_DIRECT | photorespiration, |
| | GOTERM_CC_DIRECT | peroxisome, plasma membrane, chloroplast, chloroplast stroma, membrane, apoplast, |
| | GOTERM_MF_DIRECT | catalytic activity, serine-pyruvate transaminase activity, alanine-glyoxylate transaminase activity, serine-glyoxylate transaminase activity, |
| | **At5g45310** | **coiled-coil protein(AT5G45310)** |
| | GOTERM_CC_DIRECT | Golgi apparatus, integral component of membrane, |
| | **At3g46900** | **copper transporter 2(COPT2)** |
| | GOTERM_BP_DIRECT | copper ion transport, high-affinity copper ion transport, copper ion transmembrane transport, |
| | GOTERM_CC_DIRECT | integral component of membrane, anchored component of plasma membrane, |
| | GOTERM_MF_DIRECT | copper ion transmembrane transporter activity, high-affinity copper ion transmembrane transporter activity, |
| | **At4g11490** | **cysteine-rich RLK (RECEPTOR-like protein kinase) 33(CRK33)** |
| | GOTERM_BP_DIRECT | protein phosphorylation, defense response to bacterium, |
| | GOTERM_CC_DIRECT | extracellular region, plasma membrane, plasmodesma, integral component of membrane, |
| | GOTERM_MF_DIRECT | protein serine/threonine kinase activity, ATP binding, kinase activity, |
| | **At5g42580** | **cytochrome P450, family 705, subfamily A, polypeptide 12(CYP705A12)** |
| | GOTERM_BP_DIRECT | secondary metabolite biosynthetic process, oxidation-reduction process, |
| | GOTERM_CC_DIRECT | extracellular region, plasmodesma, membrane, integral component of membrane, |
| | GOTERM_MF_DIRECT | iron ion binding, oxidoreductase activity, acting on paired donors, with incorporation or reduction of molecular oxygen, NAD(P)H as one donor, and incorporation of one atom of oxygen, oxygen binding, heme binding, |
| | **At4g12330** | **cytochrome P450, family 706, subfamily A, polypeptide 7(CYP706A7)** |
| | GOTERM_BP_DIRECT | secondary metabolite biosynthetic process, oxidation-reduction process, |
| | GOTERM_CC_DIRECT | membrane, integral component of membrane, |
| | GOTERM_MF_DIRECT | iron ion binding, oxidoreductase activity, acting on paired donors, with incorporation or reduction of molecular oxygen, NAD(P)H as one donor, and incorporation of one atom of oxygen, oxygen binding, heme binding, |
| | **At4g27710** | **cytochrome P450, family 709, subfamily B, polypeptide 3(CYP709B3)** |

| | |
|---|---|
| GOTERM_BP_DIRECT | response to salt stress, response to abscisic acid, oxidation-reduction process, |
| GOTERM_CC_DIRECT | integral component of membrane, |
| GOTERM_MF_DIRECT | monooxygenase activity, iron ion binding, oxidoreductase activity, acting on paired donors, with incorporation or reduction of molecular oxygen, oxygen binding, heme binding, |
| **At5g42590** | **cytochrome P450, family 71, subfamily A, polypeptide 16(CYP71A16)** |
| GOTERM_BP_DIRECT | secondary metabolite biosynthetic process, oxidation-reduction process, |
| GOTERM_CC_DIRECT | extracellular region, endoplasmic reticulum, chloroplast, membrane, integral component of membrane, |
| GOTERM_MF_DIRECT | iron ion binding, oxidoreductase activity, acting on paired donors, with incorporation or reduction of molecular oxygen, NAD(P)H as one donor, and incorporation of one atom of oxygen, oxygen binding, heme binding, |
| **At1g13080** | **cytochrome P450, family 71, subfamily B, polypeptide 2(CYP71B2)** |
| GOTERM_BP_DIRECT | heat acclimation, secondary metabolite biosynthetic process, oxidation-reduction process, defense response to other organism, |
| GOTERM_CC_DIRECT | extracellular region, membrane, integral component of membrane, |
| GOTERM_MF_DIRECT | iron ion binding, oxidoreductase activity, acting on paired donors, with incorporation or reduction of molecular oxygen, NAD(P)H as one donor, and incorporation of one atom of oxygen, oxygen binding, heme binding, |
| **At3g26290** | **cytochrome P450, family 71, subfamily B, polypeptide 26(CYP71B26)** |
| GOTERM_BP_DIRECT | secondary metabolite biosynthetic process, oxidation-reduction process, |
| GOTERM_CC_DIRECT | membrane, integral component of membrane, |
| GOTERM_MF_DIRECT | iron ion binding, oxidoreductase activity, acting on paired donors, with incorporation or reduction of molecular oxygen, NAD(P)H as one donor, and incorporation of one atom of oxygen, oxygen binding, heme binding, |
| **At2g42850** | **cytochrome P450, family 718(CYP718)** |
| GOTERM_BP_DIRECT | multicellular organism development, brassinosteroid homeostasis, sterol metabolic process, brassinosteroid biosynthetic process, oxidation-reduction process, |
| GOTERM_CC_DIRECT | chloroplast, integral component of membrane, |
| GOTERM_MF_DIRECT | monooxygenase activity, iron ion binding, oxidoreductase activity, acting on paired donors, with incorporation or reduction of molecular oxygen, oxygen binding, heme binding, |
| **At1g13710** | **cytochrome P450, family 78, subfamily A, polypeptide 5(CYP78A5)** |
| GOTERM_BP_DIRECT | positive regulation of cell proliferation, regulation of meristem growth, leaf formation, organ growth, regulation of growth rate, secondary metabolite biosynthetic process, positive regulation of organ growth, floral organ development, oxidation-reduction process, |
| GOTERM_CC_DIRECT | endoplasmic reticulum, membrane, integral component of membrane, |
| GOTERM_MF_DIRECT | monooxygenase activity, iron ion binding, oxidoreductase activity, acting on paired donors, with incorporation or reduction of molecular oxygen, oxidoreductase activity, acting on paired donors, with incorporation or reduction of molecular oxygen, NAD(P)H as one donor, and incorporation of one atom of oxygen, oxygen binding, heme binding, |
| **At2g46660** | **cytochrome P450, family 78, subfamily A, polypeptide 6(CYP78A6)** |
| GOTERM_BP_DIRECT | regulation of growth, secondary metabolite biosynthetic process, seed development, oxidation-reduction process, |
| GOTERM_CC_DIRECT | chloroplast, membrane, integral component of membrane, |
| GOTERM_MF_DIRECT | iron ion binding, oxidoreductase activity, acting on paired donors, with incorporation or reduction of molecular oxygen, NAD(P)H as one donor, and incorporation of one atom of oxygen, oxygen binding, heme binding, |

| | | |
|---|---|---|
| | **At1g34540** | **cytochrome P450, family 94, subfamily D, polypeptide 1(CYP94D1)** |
| | GOTERM_BP_DIRECT | oxidation-reduction process, |
| | GOTERM_CC_DIRECT | chloroplast, |
| | GOTERM_MF_DIRECT | monooxygenase activity, iron ion binding, oxidoreductase activity, acting on paired donors, with incorporation or reduction of molecular oxygen, oxygen binding, heme binding, |
| | **At4g36220** | **ferulic acid 5-hydroxylase 1(FAH1)** |
| | GOTERM_BP_DIRECT | phenylpropanoid biosynthetic process, lignin biosynthetic process, response to UV-B, oxidation-reduction process, |
| | GOTERM_CC_DIRECT | endoplasmic reticulum, membrane, integral component of membrane, |
| | GOTERM_MF_DIRECT | monooxygenase activity, iron ion binding, protein binding, oxidoreductase activity, acting on paired donors, with incorporation or reduction of molecular oxygen, oxidoreductase activity, acting on paired donors, with incorporation or reduction of molecular oxygen, NAD(P)H as one donor, and incorporation of one atom of oxygen, heme binding, ferulate 5-hydroxylase activity, |
| | **At1g63930** | **from the Czech 'roh' meaning 'corner'(ROH1)** |
| | GOTERM_BP_DIRECT | seed coat development, mucilage biosynthetic process involved in seed coat development, |
| | GOTERM_CC_DIRECT | nucleus, |
| | **At1g30260** | **galactosyltransferase family protein(AT1G30260)** |
| | GOTERM_BP_DIRECT | response to cytokinin, |
| | GOTERM_CC_DIRECT | mitochondrion, |
| | **At1g34760** | **general regulatory factor 11(GRF11)** |
| | GOTERM_CC_DIRECT | cytoplasm, |
| | GOTERM_MF_DIRECT | amino acid binding, protein domain specific binding, protein phosphorylated amino acid binding, ATPase binding, |
| | **At5g56100** | **glycine-rich protein / oleosin(AT5G56100)** |
| | GOTERM_BP_DIRECT | lipid storage, |
| | GOTERM_CC_DIRECT | nucleus, membrane, integral component of membrane, |
| | **At4g39000** | **glycosyl hydrolase 9B17(GH9B17)** |
| | GOTERM_BP_DIRECT | cellulose catabolic process, cell wall organization, |
| | GOTERM_CC_DIRECT | extracellular region, |
| | GOTERM_MF_DIRECT | hydrolase activity, hydrolyzing O-glycosyl compounds, cellulase activity, |
| | **At4g10310** | **high-affinity K+ transporter 1(HKT1)** |
| | GOTERM_BP_DIRECT | potassium ion transport, sodium ion transport, response to osmotic stress, response to salt stress, potassium ion transmembrane transport, |
| | GOTERM_CC_DIRECT | plasma membrane, integral component of membrane, |
| | GOTERM_MF_DIRECT | potassium ion transmembrane transporter activity, sodium ion transmembrane transporter activity, |
| | **At5g53980** | **homeobox protein 52(HB52)** |
| | GOTERM_BP_DIRECT | transcription, DNA-templated, regulation of transcription, DNA-templated, response to blue light, response to absence of light, |

| | |
|---|---|
| GOTERM_CC_DIRECT | nucleus, |
| GOTERM_MF_DIRECT | transcription factor activity, sequence-specific DNA binding, sequence-specific DNA binding, |
| **At3g60700** | **hypothetical protein (DUF1163)(AT3G60700)** |
| GOTERM_CC_DIRECT | nucleus, |
| **At5g44660** | **hypothetical protein(AT5G44660)** |
| GOTERM_CC_DIRECT | nucleus, |
| **At3g17600** | **indole-3-acetic acid inducible 31(IAA31)** |
| GOTERM_BP_DIRECT | transcription, DNA-templated, regulation of transcription, DNA-templated, gravitropism, response to auxin, auxin-activated signaling pathway, root development, shoot system development, |
| GOTERM_CC_DIRECT | nucleus, |
| GOTERM_MF_DIRECT | transcription factor activity, sequence-specific DNA binding, |
| **At5g19040** | **isopentenyltransferase 5(IPT5)** |
| GOTERM_BP_DIRECT | tRNA modification, cytokinin biosynthetic process, |
| GOTERM_CC_DIRECT | mitochondrion, chloroplast, plastid, |
| GOTERM_MF_DIRECT | ATP binding, AMP dimethylallyltransferase activity, transferase activity, transferring alkyl or aryl (other than methyl) groups, tRNA dimethylallyltransferase activity, ATP dimethylallyltransferase activity, ADP dimethylallyltransferase activity, |
| **At2g23620** | **methyl esterase 1(MES1)** |
| GOTERM_BP_DIRECT | systemic acquired resistance, salicylic acid metabolic process, defense response to fungus, incompatible interaction, |
| GOTERM_CC_DIRECT | cytoplasm, |
| GOTERM_MF_DIRECT | hydrolase activity, acting on ester bonds, methyl indole-3-acetate esterase activity, methyl salicylate esterase activity, methyl jasmonate esterase activity, |
| **At2g25680** | **molybdate transporter 1(MOT1)** |
| GOTERM_BP_DIRECT | molybdate ion transport, |
| GOTERM_CC_DIRECT | mitochondrion, vacuole, plasma membrane, chloroplast, endomembrane system, integral component of membrane, mitochondrial membrane, |
| GOTERM_MF_DIRECT | molybdate ion transmembrane transporter activity, sulfate transmembrane transporter activity, |
| **At4g08290** | **nodulin MtN21 /EamA-like transporter family protein(UMAMIT20)** |
| GOTERM_BP_DIRECT | amino acid export, amino acid import, amino acid homeostasis, |
| GOTERM_CC_DIRECT | extracellular region, plasma membrane, integral component of membrane, |
| GOTERM_MF_DIRECT | amino acid transmembrane transporter activity, |
| **At1g16370** | **organic cation/carnitine transporter 6(OCT6)** |
| GOTERM_BP_DIRECT | ion transport, cellular response to water deprivation, transmembrane transport, cellular response to salt stress, |
| GOTERM_CC_DIRECT | plasma membrane, plant-type vacuole membrane, membrane, integral component of membrane, |
| GOTERM_MF_DIRECT | sugar:proton symporter activity, ATP binding, carbohydrate transmembrane transporter activity, substrate-specific transmembrane transporter activity, |

| | |
|---|---|
| **At5g58360** | **ovate family protein 3(OFP3)** |
| GOTERM_BP_DIRECT | transcription, DNA-templated, negative regulation of transcription, DNA-templated, |
| GOTERM_CC_DIRECT | nucleus, |
| GOTERM_MF_DIRECT | DNA binding, |
| **At4g15340** | **pentacyclic triterpene synthase 1(PEN1)** |
| GOTERM_BP_DIRECT | tricyclic triterpenoid biosynthetic process, triterpenoid biosynthetic process, |
| GOTERM_MF_DIRECT | lyase activity, arabidiol synthase activity, |
| **At4g26050** | **plant intracellular ras group-related LRR 8(PIRL8)** |
| **At3g02610** | **plant stearoyl-acyl-carrier desaturase family protein(AT3G02610)** |
| GOTERM_BP_DIRECT | fatty acid metabolic process, fatty acid biosynthetic process, unsaturated fatty acid biosynthetic process, endosperm development, oxidation-reduction process, |
| GOTERM_CC_DIRECT | chloroplast, |
| GOTERM_MF_DIRECT | acyl-[acyl-carrier-protein] desaturase activity, metal ion binding, |
| **At5g60660** | **plasma membrane intrinsic protein 2;4(PIP2;4)** |
| GOTERM_BP_DIRECT | transport, response to abscisic acid, cellular water homeostasis, ion transmembrane transport, root hair elongation, hydrogen peroxide transmembrane transport, |
| GOTERM_CC_DIRECT | vacuole, plasma membrane, integral component of plasma membrane, plasmodesma, membrane, integral component of membrane, |
| GOTERM_MF_DIRECT | transporter activity, water channel activity, glycerol channel activity, |
| **At3g20850** | **proline-rich family protein(AT3G20850)** |
| **At5g26080** | **proline-rich family protein(AT5G26080)** |
| GOTERM_CC_DIRECT | plasma membrane, |
| **At3g62680** | **proline-rich protein 3(PRP3)** |
| GOTERM_BP_DIRECT | trichoblast differentiation, cellular response to auxin stimulus, cellular response to ethylene stimulus, cellular response to calcium ion starvation, |
| GOTERM_CC_DIRECT | extracellular region, cell wall, plasmodesma, |
| **At5g19790** | **related to AP2 11(RAP2.11)** |
| GOTERM_BP_DIRECT | response to reactive oxygen species, transcription, DNA-templated, regulation of transcription, DNA-templated, response to ethylene, ethylene-activated signaling pathway, cellular response to potassium ion, post-embryonic root development, |
| GOTERM_CC_DIRECT | nucleus, |
| GOTERM_MF_DIRECT | DNA binding, transcription factor activity, sequence-specific DNA binding, sequence-specific DNA binding, |
| **At1g43160** | **related to AP2 6(RAP2.6)** |
| GOTERM_BP_DIRECT | transcription, DNA-templated, regulation of transcription, DNA-templated, response to osmotic stress, response to cold, response to water deprivation, response to wounding, response to salt stress, chloroplast organization, response to abscisic acid, response to salicylic acid, response to jasmonic acid, ethylene-activated signaling pathway, cellular response to heat, positive regulation of transcription, DNA-templated, |
| GOTERM_CC_DIRECT | nucleus, |
| GOTERM_MF_DIRECT | DNA binding, transcription factor activity, sequence-specific DNA binding, |

| | | |
|---|---|---|
| | **At1g70460** | **root hair specific 10(PERK13)** |
| | GOTERM_BP_DIRECT | protein phosphorylation, |
| | GOTERM_CC_DIRECT | plasma membrane, membrane, integral component of membrane, |
| | GOTERM_MF_DIRECT | protein kinase activity, protein serine/threonine kinase activity, ATP binding, protein kinase binding, |
| | **At4g25220** | **root hair specific 15(G3Pp2)** |
| | GOTERM_BP_DIRECT | transport, anion transport, carbohydrate transport, phosphate ion homeostasis, transmembrane transport, |
| | GOTERM_CC_DIRECT | cytoplasm, integral component of membrane, |
| | GOTERM_MF_DIRECT | transporter activity, transmembrane transporter activity, |
| | **At4g38390** | **root hair specific 17(RHS17)** |
| | GOTERM_CC_DIRECT | Golgi apparatus, integral component of membrane, |
| | GOTERM_MF_DIRECT | transferase activity, transferring glycosyl groups, |
| | **At3g23800** | **selenium-binding protein 3(SBP3)** |
| | GOTERM_CC_DIRECT | nucleus, |
| | GOTERM_MF_DIRECT | selenium binding, |
| | **At2g33810** | **squamosa promoter binding protein-like 3(SPL3)** |
| | GOTERM_BP_DIRECT | transcription, DNA-templated, regulation of transcription, DNA-templated, flower development, positive regulation of flower development, vegetative to reproductive phase transition of meristem, inflorescence development, regulation of vegetative phase change, cell differentiation, |
| | GOTERM_CC_DIRECT | nucleus, cytoplasm, |
| | GOTERM_MF_DIRECT | DNA binding, transcription factor activity, sequence-specific DNA binding, metal ion binding, |
| | **At5g25240** | **stress induced protein(AT5G25240)** |
| | **At1g49500** | **transcription initiation factor TFIID subunit 1b-like protein(AT1G49500)** |
| | **At3g25805** | **transmembrane protein(AT3G25805)** |
| | GOTERM_CC_DIRECT | chloroplast, integral component of membrane, |
| | **At5g24313** | **transmembrane protein(AT5G24313)** |
| | GOTERM_CC_DIRECT | mitochondrion, |
| | **At5g26270** | **transmembrane protein(AT5G26270)** |
| | GOTERM_CC_DIRECT | nucleus, integral component of membrane, |
| | **At5g57530** | **xyloglucan endotransglucosylase/hydrolase 12(XTH12)** |
| | GOTERM_BP_DIRECT | xyloglucan metabolic process, cell wall biogenesis, cell wall organization, |
| | GOTERM_CC_DIRECT | extracellular region, cell wall, cytoplasm, apoplast, |
| | GOTERM_MF_DIRECT | hydrolase activity, hydrolyzing O-glycosyl compounds, xyloglucan:xyloglucosyl transferase activity, hydrolase activity, acting on glycosyl bonds, xyloglucan-specific endo-beta-1,4-glucanase activity, xyloglucan endotransglucosylase activity, |
