## Supplementary material for "Evidences for a nutritional role of iodine in plants": Table S8

| GO term | Description | P-value | FDR q-value | Enrichment (N, B, n, b) | Genes |
|---|---|---|---|---|---|
| GO:0050896 | response to stimulus | 2.08E-21 | 8.4E-18 | 2.82 (11869,1306,296,92) | [-] Hide genes<br><br>AT5G22530 - hypothetical protein<br>AT4G20860 - fad-binding berberine family protein<br>AT5G18470 - curculin-like (mannose-binding) lectin family protein<br>AT2G21210 - saur-like auxin-responsive protein<br>AT4G37290 - hypothetical protein<br>AT5G61590 - ethylene-responsive transcription factor erf107<br>AT4G28460 - hypothetical protein<br>AT1G66090 - tir-nbs class of disease resistance protein<br>AT1G68620 - probable carboxylesterase 6<br>AT1G57630 - toll-interleukin-resistance domain-containing protein<br>AT1G15010 - hypothetical protein<br>AT1G68690 - proline-rich receptor-like protein kinase perk9<br>AT1G30260 - hypothetical protein<br>AT5G57510 - hypothetical protein<br>AT4G36430 - peroxidase 49<br>AT2G40460 - putative peptide/nitrate transporter<br>AT1G12200 - putative flavin monooxygenase.<br>AT5G65600 - concanavalin a-like lectin kinase-like protein<br>AT5G02490 - heat shock protein 70<br>AT1G35910 - probable trehalose-phosphate phosphatase d<br>AT5G64120 - peroxidase 71<br>AT1G72520 - lipoxygenase 4<br>AT4G20830 - fad-binding berberine family protein<br>AT3G23170 - hypothetical protein<br>AT3G16530 - legume lectin-like protein<br>AT5G64110 - peroxidase 70<br>AT1G19020 - hypothetical protein |

| | | | | | |
|---|---|---|---|---|---|
| | | | | | AT5G06730 - peroxidase<br>AT1G76600 - hypothetical protein<br>AT1G73805 - protein sar deficient 1<br>AT4G10500 - oxidoreductase, 2og-fe(ii) oxygenase family protein<br>AT5G14130 - peroxidase 55<br>AT5G19880 - peroxidase<br>AT4G37710 - vq motif-containing protein<br>AT5G05340 - peroxidase 52<br>AT2G22880 - vq motif-containing protein<br>AT1G13340 - regulator of vps4 activity in the mvb pathway protein<br>AT1G26380 - fad-binding and bbe domain-containing protein<br>AT1G72910 - toll-interleukin-resistance domain-containing protein<br>AT1G34510 - peroxidase 8<br>AT1G11330 - g-type lectin s-receptor-like serine/threonine-protein kinase<br>AT1G26410 - fad-binding and bbe domain-containing protein<br>AT1G72940 - toll-interleukin-resistance domain-containing protein<br>AT2G37430 - zinc finger protein zat11<br>AT1G76520 - auxin efflux carrier family protein<br>AT3G21670 - nitrate transporter 1.3<br>AT1G14550 - peroxidase 5<br>AT5G39580 - peroxidase 62<br>AT1G14540 - peroxidase 4<br>AT4G08780 - peroxidase 38<br>AT1G35210 - hypothetical protein<br>AT4G23680 - polyketide cyclase/dehydrase and lipid transport superfamily protein<br>AT1G58420 - hypothetical protein<br>AT1G65500 - hypothetical protein<br>AT3G45710 - major facilitator superfamily protein<br>AT5G10760 - aspartyl protease family protein<br>AT4G26120 - regulatory protein npr2 |

| | | | | | |
|---|---|---|---|---|---|
| | | | | | AT3G22550 - hypothetical protein<br>AT3G12700 - aspartyl protease family protein<br>AT3G28580 - aaa-type atpase family protein<br>AT5G44660 - hypothetical protein<br>AT1G22500 - putative c3hc4-type ring zinc finger protein<br>AT2G41090 - calmodulin-like protein 10<br>AT2G38340 - dehydration-responsive element-binding protein 2e<br>AT3G55790 - hypothetical protein<br>AT1G43910 - p-loop containing nucleoside triphosphate hydrolases superfamily protein<br>AT3G26470 - rpw8 domain-containing powdery mildew resistance protein<br>AT1G53070 - legume lectin-like protein<br>AT5G22270 - hypothetical protein<br>AT4G17670 - hypothetical protein<br>AT2G41380 - s-adenosyl-l-methionine-dependent methyltransferase-like protein<br>AT4G11170 - putative disease resistance protein<br>AT5G25250 - flotillin-like protein 1<br>AT1G61550 - g-type lectin s-receptor-like serine/threonine-protein kinase<br>AT5G19240 - gpi-anchored glycoprotein membrane precursor<br>AT5G48540 - receptor-like protein kinase-related family protein<br>AT2G18140 - peroxidase 14<br>AT5G38710 - proline dehydrogenase 2<br>AT1G61500 - g-type lectin s-receptor-like serine/threonine-protein kinase<br>AT2G38870 - pr-6 proteinase inhibitor family protein<br>AT2G29250 - concanavalin a-like lectin protein kinase-like protein<br>AT2G35380 - peroxidase 20<br>AT2G29220 - putative inactive l-type lectin-domain containing receptor kinase iii.1 |

| | | | | | |
|---|---|---|---|---|---|
| | | | | | AT3G53600 - c2h2-type zinc finger protein<br>AT3G02840 - hypothetical protein<br>AT3G14415 - peroxisomal (s)-2-hydroxy-acid oxidase glo2<br>AT2G32020 - acyl-coa n-acyltransferases (nat) superfamily protein<br>AT5G38900 - thioredoxin superfamily protein<br>AT4G37220 - cold acclimation protein wcor413<br>AT1G70880 - srpbcc domain-containing protein<br>AT4G19810 - class v chitinase<br>AT2G01900 - dnase i-like superfamily protein |
| GO:0006950 | response to stress | 9.97E-20 | 2.01E-16 | 3.23 (11869,893,296,72) | [-] Hide genes<br><br>AT4G20860 - fad-binding berberine family protein<br>AT5G22530 - hypothetical protein<br>AT5G61590 - ethylene-responsive transcription factor erf107<br>AT4G28460 - hypothetical protein<br>AT1G66090 - tir-nbs class of disease resistance protein<br>AT1G68620 - probable carboxylesterase 6<br>AT1G57630 - toll-interleukin-resistance domain-containing protein<br>AT1G15010 - hypothetical protein<br>AT1G68690 - proline-rich receptor-like protein kinase perk9<br>AT5G57510 - hypothetical protein<br>AT4G36430 - peroxidase 49<br>AT1G12200 - putative flavin monooxygenase.<br>AT5G65600 - concanavalin a-like lectin kinase-like protein<br>AT5G02490 - heat shock protein 70<br>AT1G35910 - probable trehalose-phosphate phosphatase d<br>AT5G64120 - peroxidase 71<br>AT1G72520 - lipoxygenase 4<br>AT4G20830 - fad-binding berberine family protein<br>AT3G23170 - hypothetical protein<br>AT5G64110 - peroxidase 70<br>AT1G19020 - hypothetical protein |

| | | | | | |
|---|---|---|---|---|---|
| | | | | | AT5G06730 - peroxidase<br>AT1G76600 - hypothetical protein<br>AT1G73805 - protein sar deficient 1<br>AT5G14130 - peroxidase 55<br>AT4G10500 - oxidoreductase, 2og-fe(ii) oxygenase family protein<br>AT5G19880 - peroxidase<br>AT4G37710 - vq motif-containing protein<br>AT5G05340 - peroxidase 52<br>AT1G13340 - regulator of vps4 activity in the mvb pathway protein<br>AT1G26380 - fad-binding and bbe domain-containing protein<br>AT1G72910 - toll-interleukin-resistance domain-containing protein<br>AT1G34510 - peroxidase 8<br>AT1G26410 - fad-binding and bbe domain-containing protein<br>AT1G72940 - toll-interleukin-resistance domain-containing protein<br>AT2G37430 - zinc finger protein zat11<br>AT1G14550 - peroxidase 5<br>AT1G14540 - peroxidase 4<br>AT5G39580 - peroxidase 62<br>AT4G08780 - peroxidase 38<br>AT4G23680 - polyketide cyclase/dehydrase and lipid transport superfamily protein<br>AT1G58420 - hypothetical protein<br>AT1G65500 - hypothetical protein<br>AT5G10760 - aspartyl protease family protein<br>AT4G26120 - regulatory protein npr2<br>AT3G22550 - hypothetical protein<br>AT3G28580 - aaa-type atpase family protein<br>AT5G44660 - hypothetical protein<br>AT2G41090 - calmodulin-like protein 10<br>AT2G38340 - dehydration-responsive element-binding protein 2e<br>AT3G55790 - hypothetical protein<br>AT3G26470 - rpw8 domain-containing powdery mildew resistance |

| | | | | | |
|---|---|---|---|---|---|
| | | | | | protein<br>AT5G22270 - hypothetical protein<br>AT4G17670 - hypothetical protein<br>AT4G11170 - putative disease resistance protein<br>AT5G25250 - flotillin-like protein 1<br>AT1G61550 - g-type lectin s-receptor-like serine/threonine-protein kinase<br>AT5G19240 - gpi-anchored glycoprotein membrane precursor<br>AT2G18140 - peroxidase 14<br>AT1G61500 - g-type lectin s-receptor-like serine/threonine-protein kinase<br>AT5G38710 - proline dehydrogenase 2<br>AT2G38870 - pr-6 proteinase inhibitor family protein<br>AT2G29250 - concanavalin a-like lectin protein kinase-like protein<br>AT2G35380 - peroxidase 20<br>AT2G29220 - putative inactive l-type lectin-domain containing receptor kinase iii.1<br>AT3G53600 - c2h2-type zinc finger protein<br>AT3G02840 - hypothetical protein<br>AT3G14415 - peroxisomal (s)-2-hydroxy-acid oxidase glo2<br>AT5G38900 - thioredoxin superfamily protein<br>AT1G70880 - srpbcc domain-containing protein<br>AT4G19810 - class v chitinase<br>AT2G01900 - dnase i-like superfamily protein |
| GO:0042221 | response to chemical | 1.53E-15 | 2.06E-12 | 3.54 (11869,578,296,51) | [-] Hide genes<br><br>AT5G22530 - hypothetical protein<br>AT4G20860 - fad-binding berberine family protein<br>AT2G21210 - saur-like auxin-responsive protein<br>AT5G61590 - ethylene-responsive transcription factor erf107<br>AT1G66090 - tir-nbs class of disease resistance protein<br>AT1G68620 - probable carboxylesterase 6 |

| | | | | | |
|---|---|---|---|---|---|
| | | | | | AT1G57630 - toll-interleukin-resistance domain-containing protein<br>AT1G68690 - proline-rich receptor-like protein kinase perk9<br>AT1G15010 - hypothetical protein<br>AT1G30260 - hypothetical protein<br>AT5G57510 - hypothetical protein<br>AT5G02490 - heat shock protein 70<br>AT1G72520 - lipoxygenase 4<br>AT3G23170 - hypothetical protein<br>AT3G16530 - legume lectin-like protein<br>AT1G19020 - hypothetical protein<br>AT1G76600 - hypothetical protein<br>AT1G73805 - protein sar deficient 1<br>AT4G10500 - oxidoreductase, 2og-fe(ii) oxygenase family protein<br>AT4G37710 - vq motif-containing protein<br>AT1G26380 - fad-binding and bbe domain-containing protein<br>AT1G72910 - toll-interleukin-resistance domain-containing protein<br>AT1G26410 - fad-binding and bbe domain-containing protein<br>AT1G72940 - toll-interleukin-resistance domain-containing protein<br>AT2G37430 - zinc finger protein zat11<br>AT1G76520 - auxin efflux carrier family protein<br>AT3G21670 - nitrate transporter 1.3<br>AT1G14550 - peroxidase 5<br>AT1G14540 - peroxidase 4<br>AT1G65500 - hypothetical protein<br>AT3G45710 - major facilitator superfamily protein<br>AT4G26120 - regulatory protein npr2<br>AT3G22550 - hypothetical protein<br>AT3G12700 - aspartyl protease family protein<br>AT3G28580 - aaa-type atpase family protein<br>AT1G22500 - putative c3hc4-type ring zinc finger protein<br>AT2G41090 - calmodulin-like protein 10<br>AT2G38340 - dehydration-responsive element-binding protein 2e |

| | | | | | |
|---|---|---|---|---|---|
| | | | | | AT3G55790 - hypothetical protein<br>AT1G43910 - p-loop containing nucleoside triphosphate hydrolases superfamily protein<br>AT4G17670 - hypothetical protein<br>AT2G41380 - s-adenosyl-l-methionine-dependent methyltransferase-like protein<br>AT4G11170 - putative disease resistance protein<br>AT5G25250 - flotillin-like protein 1<br>AT5G19240 - gpi-anchored glycoprotein membrane precursor<br>AT5G38710 - proline dehydrogenase 2<br>AT3G53600 - c2h2-type zinc finger protein<br>AT3G02840 - hypothetical protein<br>AT2G32020 - acyl-coa n-acyltransferases (nat) superfamily protein<br>AT4G37220 - cold acclimation protein wcor413<br>AT4G19810 - class v chitinase |
| GO:0071453 | cellular response to oxygen levels | 2.62E-14 | 2.64E-11 | 8.02 (11869,110,296,22) | [-] Hide genes<br><br>AT1G19020 - hypothetical protein<br>AT1G76600 - hypothetical protein<br>AT4G20860 - fad-binding berberine family protein<br>AT5G25250 - flotillin-like protein 1<br>AT1G66090 - tir-nbs class of disease resistance protein<br>AT4G37710 - vq motif-containing protein<br>AT1G68620 - probable carboxylesterase 6<br>AT5G19240 - gpi-anchored glycoprotein membrane precursor<br>AT1G68690 - proline-rich receptor-like protein kinase perk9<br>AT1G57630 - toll-interleukin-resistance domain-containing protein<br>AT1G15010 - hypothetical protein<br>AT1G26380 - fad-binding and bbe domain-containing protein<br>AT3G22550 - hypothetical protein<br>AT5G57510 - hypothetical protein<br>AT1G72910 - toll-interleukin-resistance domain-containing protein |

| | | | | | |
|---|---|---|---|---|---|
| | | | | | AT1G26410 - fad-binding and bbe domain-containing protein<br>AT1G72940 - toll-interleukin-resistance domain-containing protein<br>AT2G41090 - calmodulin-like protein 10<br>AT1G14550 - peroxidase 5<br>AT1G14540 - peroxidase 4<br>AT3G55790 - hypothetical protein<br>AT3G23170 - hypothetical protein |
| GO:0071456 | cellular response to hypoxia | 2.62E-14 | 2.11E-11 | 8.02 (11869,110,296,22) | [-] Hide genes<br><br>AT1G19020 - hypothetical protein<br>AT1G76600 - hypothetical protein<br>AT4G20860 - fad-binding berberine family protein<br>AT5G25250 - flotillin-like protein 1<br>AT1G66090 - tir-nbs class of disease resistance protein<br>AT4G37710 - vq motif-containing protein<br>AT1G68620 - probable carboxylesterase 6<br>AT5G19240 - gpi-anchored glycoprotein membrane precursor<br>AT1G68690 - proline-rich receptor-like protein kinase perk9<br>AT1G57630 - toll-interleukin-resistance domain-containing protein<br>AT1G15010 - hypothetical protein<br>AT1G26380 - fad-binding and bbe domain-containing protein<br>AT3G22550 - hypothetical protein<br>AT5G57510 - hypothetical protein<br>AT1G72910 - toll-interleukin-resistance domain-containing protein<br>AT1G26410 - fad-binding and bbe domain-containing protein<br>AT1G72940 - toll-interleukin-resistance domain-containing protein<br>AT2G41090 - calmodulin-like protein 10<br>AT1G14550 - peroxidase 5<br>AT1G14540 - peroxidase 4<br>AT3G55790 - hypothetical protein<br>AT3G23170 - hypothetical protein |

| | | | | | |
|---|---|---|---|---|---|
| GO:0036294 | cellular response to decreased oxygen levels | 2.62E-14 | 1.76E-11 | 8.02 (11869,110,296,22) | [-] Hide genes<br><br>AT1G19020 - hypothetical protein<br>AT1G76600 - hypothetical protein<br>AT4G20860 - fad-binding berberine family protein<br>AT5G25250 - flotillin-like protein 1<br>AT1G66090 - tir-nbs class of disease resistance protein<br>AT4G37710 - vq motif-containing protein<br>AT1G68620 - probable carboxylesterase 6<br>AT5G19240 - gpi-anchored glycoprotein membrane precursor<br>AT1G57630 - toll-interleukin-resistance domain-containing protein<br>AT1G68690 - proline-rich receptor-like protein kinase perk9<br>AT1G15010 - hypothetical protein<br>AT1G26380 - fad-binding and bbe domain-containing protein<br>AT3G22550 - hypothetical protein<br>AT5G57510 - hypothetical protein<br>AT1G72910 - toll-interleukin-resistance domain-containing protein<br>AT1G26410 - fad-binding and bbe domain-containing protein<br>AT1G72940 - toll-interleukin-resistance domain-containing protein<br>AT2G41090 - calmodulin-like protein 10<br>AT1G14550 - peroxidase 5<br>AT1G14540 - peroxidase 4<br>AT3G55790 - hypothetical protein<br>AT3G23170 - hypothetical protein |
| GO:0001666 | response to hypoxia | 3.88E-14 | 2.24E-11 | 7.88 (11869,112,296,22) | [-] Hide genes<br><br>AT1G19020 - hypothetical protein<br>AT1G76600 - hypothetical protein<br>AT4G20860 - fad-binding berberine family protein<br>AT5G25250 - flotillin-like protein 1<br>AT1G66090 - tir-nbs class of disease resistance protein<br>AT4G37710 - vq motif-containing protein |

| | | | | | |
|---|---|---|---|---|---|
| | | | | | AT1G68620 - probable carboxylesterase 6<br>AT5G19240 - gpi-anchored glycoprotein membrane precursor<br>AT1G57630 - toll-interleukin-resistance domain-containing protein<br>AT1G68690 - proline-rich receptor-like protein kinase perk9<br>AT1G15010 - hypothetical protein<br>AT1G26380 - fad-binding and bbe domain-containing protein<br>AT3G22550 - hypothetical protein<br>AT5G57510 - hypothetical protein<br>AT1G72910 - toll-interleukin-resistance domain-containing protein<br>AT1G26410 - fad-binding and bbe domain-containing protein<br>AT1G72940 - toll-interleukin-resistance domain-containing protein<br>AT2G41090 - calmodulin-like protein 10<br>AT1G14550 - peroxidase 5<br>AT1G14540 - peroxidase 4<br>AT3G55790 - hypothetical protein<br>AT3G23170 - hypothetical protein |
| GO:0036293 | response to decreased oxygen levels | 4.72E-14 | 2.38E-11 | 7.81 (11869,113,296,22) | [-] Hide genes<br><br>AT1G19020 - hypothetical protein<br>AT1G76600 - hypothetical protein<br>AT4G20860 - fad-binding berberine family protein<br>AT5G25250 - flotillin-like protein 1<br>AT1G66090 - tir-nbs class of disease resistance protein<br>AT4G37710 - vq motif-containing protein<br>AT1G68620 - probable carboxylesterase 6<br>AT5G19240 - gpi-anchored glycoprotein membrane precursor<br>AT1G68690 - proline-rich receptor-like protein kinase perk9<br>AT1G57630 - toll-interleukin-resistance domain-containing protein<br>AT1G15010 - hypothetical protein<br>AT1G26380 - fad-binding and bbe domain-containing protein<br>AT3G22550 - hypothetical protein<br>AT5G57510 - hypothetical protein |

| | | | | | |
|---|---|---|---|---|---|
| | | | | | AT1G72910 - toll-interleukin-resistance domain-containing protein<br>AT1G26410 - fad-binding and bbe domain-containing protein<br>AT1G72940 - toll-interleukin-resistance domain-containing protein<br>AT2G41090 - calmodulin-like protein 10<br>AT1G14550 - peroxidase 5<br>AT1G14540 - peroxidase 4<br>AT3G55790 - hypothetical protein<br>AT3G23170 - hypothetical protein |
| GO:0070482 | response to oxygen levels | 5.71E-14 | 2.56E-11 | 7.74 (11869,114,296,22) | [-] Hide genes<br><br>AT1G19020 - hypothetical protein<br>AT1G76600 - hypothetical protein<br>AT4G20860 - fad-binding berberine family protein<br>AT5G25250 - flotillin-like protein 1<br>AT1G66090 - tir-nbs class of disease resistance protein<br>AT4G37710 - vq motif-containing protein<br>AT1G68620 - probable carboxylesterase 6<br>AT5G19240 - gpi-anchored glycoprotein membrane precursor<br>AT1G57630 - toll-interleukin-resistance domain-containing protein<br>AT1G68690 - proline-rich receptor-like protein kinase perk9<br>AT1G15010 - hypothetical protein<br>AT1G26380 - fad-binding and bbe domain-containing protein<br>AT3G22550 - hypothetical protein<br>AT5G57510 - hypothetical protein<br>AT1G72910 - toll-interleukin-resistance domain-containing protein<br>AT1G26410 - fad-binding and bbe domain-containing protein<br>AT1G72940 - toll-interleukin-resistance domain-containing protein<br>AT2G41090 - calmodulin-like protein 10<br>AT1G14550 - peroxidase 5<br>AT1G14540 - peroxidase 4 |

| | | | | | |
|---|---|---|---|---|---|
| | | | | | AT3G55790 - hypothetical protein<br>AT3G23170 - hypothetical protein |
| GO:0070887 | cellular response to chemical stimulus | 1.2E-13 | 4.86E-11 | 6.06 (11869,172,296,26) | [-] Hide genes<br><br>AT4G20860 - fad-binding berberine family protein<br>AT3G45710 - major facilitator superfamily protein<br>AT1G66090 - tir-nbs class of disease resistance protein<br>AT1G68620 - probable carboxylesterase 6<br>AT1G68690 - proline-rich receptor-like protein kinase perk9<br>AT1G57630 - toll-interleukin-resistance domain-containing protein<br>AT1G15010 - hypothetical protein<br>AT5G57510 - hypothetical protein<br>AT3G22550 - hypothetical protein<br>AT5G02490 - heat shock protein 70<br>AT2G41090 - calmodulin-like protein 10<br>AT3G55790 - hypothetical protein<br>AT3G23170 - hypothetical protein<br>AT1G19020 - hypothetical protein<br>AT1G76600 - hypothetical protein<br>AT5G25250 - flotillin-like protein 1<br>AT4G37710 - vq motif-containing protein<br>AT5G19240 - gpi-anchored glycoprotein membrane precursor<br>AT1G26380 - fad-binding and bbe domain-containing protein<br>AT1G72910 - toll-interleukin-resistance domain-containing protein<br>AT1G26410 - fad-binding and bbe domain-containing protein<br>AT1G72940 - toll-interleukin-resistance domain-containing protein<br>AT3G53600 - c2h2-type zinc finger protein<br>AT2G37430 - zinc finger protein zat11<br>AT1G14550 - peroxidase 5<br>AT1G14540 - peroxidase 4 |

| | | | | | |
|---|---|---|---|---|---|
| GO:0006979 | response to oxidative stress | 8.55E-13 | 3.14E-10 | 6.13 (11869,157,296,24) | [-] Hide genes<br><br>AT1G19020 - hypothetical protein<br>AT4G08780 - peroxidase 38<br>AT5G06730 - peroxidase<br>AT4G11170 - putative disease resistance protein<br>AT5G14130 - peroxidase 55<br>AT5G05340 - peroxidase 52<br>AT5G19880 - peroxidase<br>AT1G13340 - regulator of vps4 activity in the mvb pathway protein<br>AT2G18140 - peroxidase 14<br>AT1G34510 - peroxidase 8<br>AT3G28580 - aaa-type atpase family protein<br>AT5G44660 - hypothetical protein<br>AT2G35380 - peroxidase 20<br>AT4G36430 - peroxidase 49<br>AT1G35910 - probable trehalose-phosphate phosphatase d<br>AT3G02840 - hypothetical protein<br>AT2G41090 - calmodulin-like protein 10<br>AT1G14550 - peroxidase 5<br>AT5G64120 - peroxidase 71<br>AT1G14540 - peroxidase 4<br>AT4G20830 - fad-binding berberine family protein<br>AT1G72520 - lipoxygenase 4<br>AT5G39580 - peroxidase 62<br>AT5G64110 - peroxidase 70 |
| GO:0009628 | response to abiotic stimulus | 7.2E-11 | 2.42E-8 | 3.18 (11869,505,296,40) | [-] Hide genes<br><br>AT4G20860 - fad-binding berberine family protein<br>AT1G35210 - hypothetical protein<br>AT5G18470 - curculin-like (mannose-binding) lectin family protein<br>AT4G37290 - hypothetical protein |

| | | | | | |
|---|---|---|---|---|---|
| | | | | | AT5G61590 - ethylene-responsive transcription factor erf107<br>AT1G65500 - hypothetical protein<br>AT1G66090 - tir-nbs class of disease resistance protein<br>AT1G68620 - probable carboxylesterase 6<br>AT1G15010 - hypothetical protein<br>AT1G57630 - toll-interleukin-resistance domain-containing protein<br>AT1G68690 - proline-rich receptor-like protein kinase perk9<br>AT3G22550 - hypothetical protein<br>AT5G57510 - hypothetical protein<br>AT1G22500 - putative c3hc4-type ring zinc finger protein<br>AT5G02490 - heat shock protein 70<br>AT1G35910 - probable trehalose-phosphate phosphatase d<br>AT2G41090 - calmodulin-like protein 10<br>AT2G38340 - dehydration-responsive element-binding protein 2e<br>AT3G55790 - hypothetical protein<br>AT3G23170 - hypothetical protein<br>AT1G53070 - legume lectin-like protein<br>AT1G19020 - hypothetical protein<br>AT1G76600 - hypothetical protein<br>AT5G22270 - hypothetical protein<br>AT1G73805 - protein sar deficient 1<br>AT5G25250 - flotillin-like protein 1<br>AT4G37710 - vq motif-containing protein<br>AT2G22880 - vq motif-containing protein<br>AT5G19240 - gpi-anchored glycoprotein membrane precursor<br>AT5G48540 - receptor-like protein kinase-related family protein<br>AT5G38710 - proline dehydrogenase 2<br>AT1G26380 - fad-binding and bbe domain-containing protein<br>AT1G72910 - toll-interleukin-resistance domain-containing protein<br>AT1G26410 - fad-binding and bbe domain-containing protein<br>AT1G72940 - toll-interleukin-resistance domain-containing protein<br>AT3G53600 - c2h2-type zinc finger protein |

| | | | | | |
|---|---|---|---|---|---|
| | | | | | AT1G14550 - peroxidase 5<br>AT1G14540 - peroxidase 4<br>AT4G19810 - class v chitinase<br>AT2G01900 - dnase i-like superfamily protein |
| GO:0051704 | multi-organism process | 3.19E-9 | 9.9E-7 | 4.17 (11869,231,296,24) | [-] Hide genes<br><br>AT1G73805 - protein sar deficient 1<br>AT4G28460 - hypothetical protein<br>AT4G10500 - oxidoreductase, 2og-fe(ii) oxygenase family protein<br>AT5G10760 - aspartyl protease family protein<br>AT1G15010 - hypothetical protein<br>AT4G26120 - regulatory protein npr2<br>AT2G38870 - pr-6 proteinase inhibitor family protein<br>AT1G23160 - auxin-responsive gh3 family protein<br>AT2G29250 - concanavalin a-like lectin protein kinase-like protein<br>AT2G29220 - putative inactive l-type lectin-domain containing receptor kinase iii.1<br>AT4G36430 - peroxidase 49<br>AT1G12200 - putative flavin monooxygenase.<br>AT2G40460 - putative peptide/nitrate transporter<br>AT5G65600 - concanavalin a-like lectin kinase-like protein<br>AT5G02490 - heat shock protein 70<br>AT3G14415 - peroxisomal (s)-2-hydroxy-acid oxidase glo2<br>AT5G64120 - peroxidase 71<br>AT1G72520 - lipoxygenase 4<br>AT5G38900 - thioredoxin superfamily protein<br>AT5G39580 - peroxidase 62<br>AT4G20830 - fad-binding berberine family protein<br>AT3G23170 - hypothetical protein<br>AT3G16530 - legume lectin-like protein |

| | | | | | |
|---|---|---|---|---|---|
| | | | | | AT3G26470 - rpw8 domain-containing powdery mildew resistance protein |
| GO:0006952 | defense response | 4.18E-9 | 1.21E-6 | 3.96 (11869,253,296,25) | [-] Hide genes<br><br>AT4G23680 - polyketide cyclase/dehydrase and lipid transport superfamily protein<br>AT1G58420 - hypothetical protein<br>AT4G28460 - hypothetical protein<br>AT5G10760 - aspartyl protease family protein<br>AT1G66090 - tir-nbs class of disease resistance protein<br>AT1G15010 - hypothetical protein<br>AT4G26120 - regulatory protein npr2<br>AT1G12200 - putative flavin monooxygenase.<br>AT5G65600 - concanavalin a-like lectin kinase-like protein<br>AT5G64120 - peroxidase 71<br>AT4G20830 - fad-binding berberine family protein<br>AT1G72520 - lipoxygenase 4<br>AT3G23170 - hypothetical protein<br>AT3G26470 - rpw8 domain-containing powdery mildew resistance protein<br>AT1G73805 - protein sar deficient 1<br>AT4G10500 - oxidoreductase, 2og-fe(ii) oxygenase family protein<br>AT1G61550 - g-type lectin s-receptor-like serine/threonine-protein kinase<br>AT1G61500 - g-type lectin s-receptor-like serine/threonine-protein kinase<br>AT2G38870 - pr-6 proteinase inhibitor family protein<br>AT2G29250 - concanavalin a-like lectin protein kinase-like protein<br>AT2G29220 - putative inactive l-type lectin-domain containing receptor kinase iii.1<br>AT3G14415 - peroxisomal (s)-2-hydroxy-acid oxidase glo2<br>AT5G39580 - peroxidase 62 |

| | | | | | |
|---|---|---|---|---|---|
| | | | | | AT5G38900 - thioredoxin superfamily protein<br>AT1G70880 - srpbcc domain-containing protein |
| GO:0009605 | response to external stimulus | 5.52E-9 | 1.49E-6 | 3.55 (11869,316,296,28) | [-] Hide genes<br><br>AT4G28460 - hypothetical protein<br>AT5G10760 - aspartyl protease family protein<br>AT1G15010 - hypothetical protein<br>AT4G26120 - regulatory protein npr2<br>AT3G22550 - hypothetical protein<br>AT3G28580 - aaa-type atpase family protein<br>AT4G36430 - peroxidase 49<br>AT1G22500 - putative c3hc4-type ring zinc finger protein<br>AT2G40460 - putative peptide/nitrate transporter<br>AT1G12200 - putative flavin monooxygenase.<br>AT5G02490 - heat shock protein 70<br>AT5G65600 - concanavalin a-like lectin kinase-like protein<br>AT5G64120 - peroxidase 71<br>AT4G20830 - fad-binding berberine family protein<br>AT1G72520 - lipoxygenase 4<br>AT3G23170 - hypothetical protein<br>AT3G16530 - legume lectin-like protein<br>AT3G26470 - rpw8 domain-containing powdery mildew resistance protein<br>AT4G17670 - hypothetical protein<br>AT1G73805 - protein sar deficient 1<br>AT4G10500 - oxidoreductase, 2og-fe(ii) oxygenase family protein<br>AT2G38870 - pr-6 proteinase inhibitor family protein<br>AT2G29250 - concanavalin a-like lectin protein kinase-like protein<br>AT2G29220 - putative inactive l-type lectin-domain containing receptor kinase iii.1<br>AT3G53600 - c2h2-type zinc finger protein<br>AT3G14415 - peroxisomal (s)-2-hydroxy-acid oxidase glo2 |

| | | | | | |
|---|---|---|---|---|---|
| | | | | | AT5G39580 - peroxidase 62<br>AT5G38900 - thioredoxin superfamily protein |
| GO:0051707 | response to other organism | 7.91E-9 | 2E-6 | 4.14 (11869,223,296,23) | [-] Hide genes<br><br>AT1G73805 - protein sar deficient 1<br>AT4G28460 - hypothetical protein<br>AT4G10500 - oxidoreductase, 2og-fe(ii) oxygenase family protein<br>AT5G10760 - aspartyl protease family protein<br>AT1G15010 - hypothetical protein<br>AT4G26120 - regulatory protein npr2<br>AT2G38870 - pr-6 proteinase inhibitor family protein<br>AT2G29250 - concanavalin a-like lectin protein kinase-like protein<br>AT2G29220 - putative inactive l-type lectin-domain containing receptor kinase iii.1<br>AT4G36430 - peroxidase 49<br>AT1G12200 - putative flavin monooxygenase.<br>AT2G40460 - putative peptide/nitrate transporter<br>AT5G65600 - concanavalin a-like lectin kinase-like protein<br>AT5G02490 - heat shock protein 70<br>AT3G14415 - peroxisomal (s)-2-hydroxy-acid oxidase glo2<br>AT5G64120 - peroxidase 71<br>AT1G72520 - lipoxygenase 4<br>AT5G38900 - thioredoxin superfamily protein<br>AT5G39580 - peroxidase 62<br>AT4G20830 - fad-binding berberine family protein<br>AT3G23170 - hypothetical protein<br>AT3G16530 - legume lectin-like protein<br>AT3G26470 - rpw8 domain-containing powdery mildew resistance protein |

| | | | | | |
|---|---|---|---|---|---|
| GO:0043207 | response to external biotic stimulus | 1.21E-8 | 2.87E-6 | 4.04 (11869,228,296,23) | [-] Hide genes<br><br>AT1G73805 - protein sar deficient 1<br>AT4G28460 - hypothetical protein<br>AT4G10500 - oxidoreductase, 2og-fe(ii) oxygenase family protein<br>AT5G10760 - aspartyl protease family protein<br>AT1G15010 - hypothetical protein<br>AT4G26120 - regulatory protein npr2<br>AT2G38870 - pr-6 proteinase inhibitor family protein<br>AT2G29250 - concanavalin a-like lectin protein kinase-like protein<br>AT2G29220 - putative inactive l-type lectin-domain containing receptor kinase iii.1<br>AT4G36430 - peroxidase 49<br>AT1G12200 - putative flavin monooxygenase.<br>AT2G40460 - putative peptide/nitrate transporter<br>AT5G65600 - concanavalin a-like lectin kinase-like protein<br>AT5G02490 - heat shock protein 70<br>AT3G14415 - peroxisomal (s)-2-hydroxy-acid oxidase glo2<br>AT5G64120 - peroxidase 71<br>AT1G72520 - lipoxygenase 4<br>AT5G38900 - thioredoxin superfamily protein<br>AT5G39580 - peroxidase 62<br>AT4G20830 - fad-binding berberine family protein<br>AT3G23170 - hypothetical protein<br>AT3G16530 - legume lectin-like protein<br>AT3G26470 - rpw8 domain-containing powdery mildew resistance protein |
| GO:0009607 | response to biotic stimulus | 1.21E-8 | 2.71E-6 | 4.04 (11869,228,296,23) | [-] Hide genes<br><br>AT1G73805 - protein sar deficient 1<br>AT4G28460 - hypothetical protein<br>AT4G10500 - oxidoreductase, 2og-fe(ii) oxygenase family protein |

| | | | | | |
|---|---|---|---|---|---|
| | | | | | AT5G10760 - aspartyl protease family protein<br>AT1G15010 - hypothetical protein<br>AT4G26120 - regulatory protein npr2<br>AT2G38870 - pr-6 proteinase inhibitor family protein<br>AT2G29250 - concanavalin a-like lectin protein kinase-like protein<br>AT2G29220 - putative inactive l-type lectin-domain containing receptor kinase iii.1<br>AT4G36430 - peroxidase 49<br>AT1G12200 - putative flavin monooxygenase.<br>AT2G40460 - putative peptide/nitrate transporter<br>AT5G65600 - concanavalin a-like lectin kinase-like protein<br>AT5G02490 - heat shock protein 70<br>AT3G14415 - peroxisomal (s)-2-hydroxy-acid oxidase glo2<br>AT5G64120 - peroxidase 71<br>AT1G72520 - lipoxygenase 4<br>AT5G38900 - thioredoxin superfamily protein<br>AT5G39580 - peroxidase 62<br>AT4G20830 - fad-binding berberine family protein<br>AT3G23170 - hypothetical protein<br>AT3G16530 - legume lectin-like protein<br>AT3G26470 - rpw8 domain-containing powdery mildew resistance protein |
| GO:0051716 | cellular response to stimulus | 2.64E-8 | 5.62E-6 | 3.22 (11869,361,296,29) | [-] Hide genes<br><br>AT4G20860 - fad-binding berberine family protein<br>AT3G45710 - major facilitator superfamily protein<br>AT1G66090 - tir-nbs class of disease resistance protein<br>AT1G68620 - probable carboxylesterase 6<br>AT1G15010 - hypothetical protein<br>AT1G68690 - proline-rich receptor-like protein kinase perk9<br>AT1G57630 - toll-interleukin-resistance domain-containing protein<br>AT3G22550 - hypothetical protein |

| | | | | | |
|---|---|---|---|---|---|
| | | | | | AT5G57510 - hypothetical protein<br>AT5G02490 - heat shock protein 70<br>AT2G41090 - calmodulin-like protein 10<br>AT2G38340 - dehydration-responsive element-binding protein 2e<br>AT3G55790 - hypothetical protein<br>AT3G23170 - hypothetical protein<br>AT1G19020 - hypothetical protein<br>AT1G76600 - hypothetical protein<br>AT1G73805 - protein sar deficient 1<br>AT5G25250 - flotillin-like protein 1<br>AT4G37710 - vq motif-containing protein<br>AT5G19240 - gpi-anchored glycoprotein membrane precursor<br>AT1G26380 - fad-binding and bbe domain-containing protein<br>AT1G72910 - toll-interleukin-resistance domain-containing protein<br>AT1G26410 - fad-binding and bbe domain-containing protein<br>AT1G72940 - toll-interleukin-resistance domain-containing protein<br>AT3G53600 - c2h2-type zinc finger protein<br>AT2G37430 - zinc finger protein zat11<br>AT1G14550 - peroxidase 5<br>AT1G14540 - peroxidase 4<br>AT2G01900 - dnase i-like superfamily protein |
| GO:1901700 | response to oxygen-containing compound | 1.09E-7 | 2.21E-5 | 3.47 (11869,277,296,24) | [-] Hide genes<br><br>AT4G20860 - fad-binding berberine family protein<br>AT4G17670 - hypothetical protein<br>AT2G21210 - saur-like auxin-responsive protein<br>AT5G61590 - ethylene-responsive transcription factor erf107<br>AT1G65500 - hypothetical protein<br>AT4G11170 - putative disease resistance protein<br>AT4G10500 - oxidoreductase, 2og-fe(ii) oxygenase family protein<br>AT4G26120 - regulatory protein npr2<br>AT5G38710 - proline dehydrogenase 2 |

| | | | | | |
|---|---|---|---|---|---|
| | | | | | AT3G22550 - hypothetical protein<br>AT3G28580 - aaa-type atpase family protein<br>AT3G12700 - aspartyl protease family protein<br>AT1G22500 - putative c3hc4-type ring zinc finger protein<br>AT3G53600 - c2h2-type zinc finger protein<br>AT2G37430 - zinc finger protein zat11<br>AT3G02840 - hypothetical protein<br>AT2G32020 - acyl-coa n-acyltransferases (nat) superfamily protein<br>AT3G21670 - nitrate transporter 1.3<br>AT2G38340 - dehydration-responsive element-binding protein 2e<br>AT4G37220 - cold acclimation protein wcor413<br>AT1G72520 - lipoxygenase 4<br>AT4G19810 - class v chitinase<br>AT3G16530 - legume lectin-like protein<br>AT1G43910 - p-loop containing nucleoside triphosphate hydrolases superfamily protein |
| GO:0033554 | cellular response to stress | 1.23E-7 | 2.36E-5 | 3.25 (11869,321,296,26) | [-] Hide genes<br><br>AT4G20860 - fad-binding berberine family protein<br>AT1G66090 - tir-nbs class of disease resistance protein<br>AT1G68620 - probable carboxylesterase 6<br>AT1G15010 - hypothetical protein<br>AT1G68690 - proline-rich receptor-like protein kinase perk9<br>AT1G57630 - toll-interleukin-resistance domain-containing protein<br>AT5G57510 - hypothetical protein<br>AT3G22550 - hypothetical protein<br>AT5G02490 - heat shock protein 70<br>AT2G41090 - calmodulin-like protein 10<br>AT2G38340 - dehydration-responsive element-binding protein 2e<br>AT3G55790 - hypothetical protein<br>AT3G23170 - hypothetical protein<br>AT1G19020 - hypothetical protein |

| | | | | | |
|---|---|---|---|---|---|
| | | | | | AT1G76600 - hypothetical protein<br>AT5G25250 - flotillin-like protein 1<br>AT4G37710 - vq motif-containing protein<br>AT5G19240 - gpi-anchored glycoprotein membrane precursor<br>AT1G26380 - fad-binding and bbe domain-containing protein<br>AT1G72910 - toll-interleukin-resistance domain-containing protein<br>AT1G26410 - fad-binding and bbe domain-containing protein<br>AT1G72940 - toll-interleukin-resistance domain-containing protein<br>AT3G53600 - c2h2-type zinc finger protein<br>AT1G14550 - peroxidase 5<br>AT1G14540 - peroxidase 4<br>AT2G01900 - dnase i-like superfamily protein |
| GO:0098542 | defense response to other organism | 3.98E-7 | 7.3E-5 | 4.10 (11869,176,296,18) | [-] Hide genes<br><br>AT1G73805 - protein sar deficient 1<br>AT4G28460 - hypothetical protein<br>AT4G10500 - oxidoreductase, 2og-fe(ii) oxygenase family protein<br>AT5G10760 - aspartyl protease family protein<br>AT1G15010 - hypothetical protein<br>AT4G26120 - regulatory protein npr2<br>AT2G38870 - pr-6 proteinase inhibitor family protein<br>AT2G29250 - concanavalin a-like lectin protein kinase-like protein<br>AT2G29220 - putative inactive l-type lectin-domain containing receptor kinase iii.1<br>AT1G12200 - putative flavin monooxygenase.<br>AT5G65600 - concanavalin a-like lectin kinase-like protein<br>AT3G14415 - peroxisomal (s)-2-hydroxy-acid oxidase glo2<br>AT5G64120 - peroxidase 71<br>AT5G38900 - thioredoxin superfamily protein<br>AT5G39580 - peroxidase 62<br>AT4G20830 - fad-binding berberine family protein<br>AT3G23170 - hypothetical protein |

| | | | | | |
|---|---|---|---|---|---|
| | | | | | AT3G26470 - rpw8 domain-containing powdery mildew resistance protein |
| GO:0009620 | response to fungus | 8.45E-6 | 1.48E-3 | 5.73 (11869,70,296,10) | [-] Hide genes<br><br>AT2G38870 - pr-6 proteinase inhibitor family protein<br>AT1G12200 - putative flavin monooxygenase.<br>AT4G10500 - oxidoreductase, 2og-fe(ii) oxygenase family protein<br>AT5G64120 - peroxidase 71<br>AT1G15010 - hypothetical protein<br>AT5G38900 - thioredoxin superfamily protein<br>AT5G39580 - peroxidase 62<br>AT4G20830 - fad-binding berberine family protein<br>AT4G26120 - regulatory protein npr2<br>AT3G26470 - rpw8 domain-containing powdery mildew resistance protein |
| GO:0050832 | defense response to fungus | 1.2E-5 | 2.02E-3 | 6.22 (11869,58,296,9) | [-] Hide genes<br><br>AT2G38870 - pr-6 proteinase inhibitor family protein<br>AT1G12200 - putative flavin monooxygenase.<br>AT5G64120 - peroxidase 71<br>AT1G15010 - hypothetical protein<br>AT5G38900 - thioredoxin superfamily protein<br>AT5G39580 - peroxidase 62<br>AT4G20830 - fad-binding berberine family protein<br>AT4G26120 - regulatory protein npr2<br>AT3G26470 - rpw8 domain-containing powdery mildew resistance protein |
| GO:0001101 | response to acid chemical | 1.55E-5 | 2.51E-3 | 3.47 (11869,185,296,16) | [-] Hide genes<br><br>AT4G20860 - fad-binding berberine family protein<br>AT5G61590 - ethylene-responsive transcription factor erf107 |

| | | | | | |
|---|---|---|---|---|---|
| | | | | | AT1G65500 - hypothetical protein<br>AT4G11170 - putative disease resistance protein<br>AT4G10500 - oxidoreductase, 2og-fe(ii) oxygenase family protein<br>AT5G38710 - proline dehydrogenase 2<br>AT3G28580 - aaa-type atpase family protein<br>AT1G22500 - putative c3hc4-type ring zinc finger protein<br>AT3G53600 - c2h2-type zinc finger protein<br>AT3G02840 - hypothetical protein<br>AT2G32020 - acyl-coa n-acyltransferases (nat) superfamily protein<br>AT2G38340 - dehydration-responsive element-binding protein 2e<br>AT3G21670 - nitrate transporter 1.3<br>AT1G72520 - lipoxygenase 4<br>AT4G19810 - class v chitinase<br>AT1G43910 - p-loop containing nucleoside triphosphate hydrolases superfamily protein |
| GO:0042493 | response to drug | 5.86E-5 | 9.11E-3 | 4.61 (11869,87,296,10) | [-] Hide genes<br><br>AT3G28580 - aaa-type atpase family protein<br>AT2G21210 - saur-like auxin-responsive protein<br>AT3G53600 - c2h2-type zinc finger protein<br>AT4G11170 - putative disease resistance protein<br>AT2G37430 - zinc finger protein zat11<br>AT4G10500 - oxidoreductase, 2og-fe(ii) oxygenase family protein<br>AT3G02840 - hypothetical protein<br>AT1G72520 - lipoxygenase 4<br>AT4G26120 - regulatory protein npr2<br>AT3G16530 - legume lectin-like protein |
| GO:0010033 | response to organic substance | 7.25E-5 | 1.08E-2 | 2.65 (11869,303,296,20) | [-] Hide genes<br><br>AT4G20860 - fad-binding berberine family protein<br>AT4G17670 - hypothetical protein<br>AT1G73805 - protein sar deficient 1 |

| | | | | | |
|---|---|---|---|---|---|
| | | | | | AT2G21210 - saur-like auxin-responsive protein<br>AT4G10500 - oxidoreductase, 2og-fe(ii) oxygenase family protein<br>AT4G26120 - regulatory protein npr2<br>AT1G30260 - hypothetical protein<br>AT3G22550 - hypothetical protein<br>AT3G12700 - aspartyl protease family protein<br>AT3G28580 - aaa-type atpase family protein<br>AT1G22500 - putative c3hc4-type ring zinc finger protein<br>AT3G53600 - c2h2-type zinc finger protein<br>AT2G37430 - zinc finger protein zat11<br>AT1G76520 - auxin efflux carrier family protein<br>AT5G02490 - heat shock protein 70<br>AT2G32020 - acyl-coa n-acyltransferases (nat) superfamily protein<br>AT4G37220 - cold acclimation protein wcor413<br>AT4G19810 - class v chitinase<br>AT3G16530 - legume lectin-like protein<br>AT1G43910 - p-loop containing nucleoside triphosphate hydrolases superfamily protein |
| GO:0009617 | response to bacterium | 1.04E-4 | 1.5E-2 | 4.31 (11869,93,296,10) | [-] Hide genes<br><br>AT2G29250 - concanavalin a-like lectin protein kinase-like protein<br>AT1G73805 - protein sar deficient 1<br>AT2G29220 - putative inactive l-type lectin-domain containing receptor kinase iii.1<br>AT4G10500 - oxidoreductase, 2og-fe(ii) oxygenase family protein<br>AT5G65600 - concanavalin a-like lectin kinase-like protein<br>AT5G02490 - heat shock protein 70<br>AT4G28460 - hypothetical protein<br>AT3G14415 - peroxisomal (s)-2-hydroxy-acid oxidase glo2<br>AT1G72520 - lipoxygenase 4<br>AT4G26120 - regulatory protein npr2 |

| | | | | | |
|---|---|---|---|---|---|
| GO:0010193 | response to ozone | 1.48E-4 | 2.06E-2 | 24.06 (11869,5,296,3) | [-] Hide genes<br>AT4G11170 - putative disease resistance protein<br>AT3G02840 - hypothetical protein<br>AT1G72520 - lipoxygenase 4 |
| GO:0010224 | response to UV-B | 2.27E-4 | 3.05E-2 | 12.34 (11869,13,296,4) | [-] Hide genes<br>AT1G35210 - hypothetical protein<br>AT1G73805 - protein sar deficient 1<br>AT3G53600 - c2h2-type zinc finger protein<br>AT2G22880 - vq motif-containing protein |
| GO:2000377 | regulation of reactive oxygen species metabolic process | 2.27E-4 | 2.95E-2 | 12.34 (11869,13,296,4) | [-] Hide genes<br>AT4G20860 - fad-binding berberine family protein<br>AT5G22270 - hypothetical protein<br>AT5G65600 - concanavalin a-like lectin kinase-like protein<br>AT2G01900 - dnase i-like superfamily protein |
| GO:0018958 | phenol-containing compound metabolic process | 2.9E-4 | 3.66E-2 | 20.05 (11869,6,296,3) | [-] Hide genes<br>AT1G64160 - dirigent-like protein dir5<br>AT4G10500 - oxidoreductase, 2og-fe(ii) oxygenase family protein<br>AT5G63560 - hxxxd-type acyl-transferase-like protein |
| GO:0005991 | trehalose metabolic process | 2.9E-4 | 3.55E-2 | 20.05 (11869,6,296,3) | [-] Hide genes<br>AT4G39770 - probable trehalose-phosphate phosphatase h<br>AT2G22190 - probable trehalose-phosphate phosphatase e<br>AT1G35910 - probable trehalose-phosphate phosphatase d |

| | | | | | |
|---|---|---|---|---|---|
| GO:0005992 | trehalose biosynthetic process | 2.9E-4 | 3.45E-2 | 20.05 (11869,6,296,3) | [-] Hide genes<br>AT4G39770 - probable trehalose-phosphate phosphatase h<br>AT2G22190 - probable trehalose-phosphate phosphatase e<br>AT1G35910 - probable trehalose-phosphate phosphatase d |
| GO:0034285 | response to disaccharide | 3.11E-4 | 3.59E-2 | 11.46 (11869,14,296,4) | [-] Hide genes<br>AT3G22550 - hypothetical protein<br>AT3G12700 - aspartyl protease family protein<br>AT4G17670 - hypothetical protein<br>AT4G37220 - cold acclimation protein wcor413 |
| GO:0009744 | response to sucrose | 3.11E-4 | 3.49E-2 | 11.46 (11869,14,296,4) | [-] Hide genes<br>AT3G22550 - hypothetical protein<br>AT3G12700 - aspartyl protease family protein<br>AT4G17670 - hypothetical protein<br>AT4G37220 - cold acclimation protein wcor413 |
| GO:0009698 | phenylpropanoid metabolic process | 3.29E-4 | 3.59E-2 | 8.02 (11869,25,296,5) | [-] Hide genes<br>AT1G64160 - dirigent-like protein dir5<br>AT5G05340 - peroxidase 52<br>AT5G63560 - hxxxd-type acyl-transferase-like protein<br>AT2G37360 - abc transporter g family member 2<br>AT5G64120 - peroxidase 71 |
| GO:0010035 | response to inorganic substance | 3.79E-4 | 4.03E-2 | 2.88 (11869,195,296,14) | [-] Hide genes<br>AT5G22530 - hypothetical protein<br>AT2G41380 - s-adenosyl-l-methionine-dependent methyltransferase-like protein<br>AT5G61590 - ethylene-responsive transcription factor erf107<br>AT1G65500 - hypothetical protein |

| | | | | | |
|---|---|---|---|---|---|
| | | | | | AT4G11170 - putative disease resistance protein<br>AT5G38710 - proline dehydrogenase 2<br>AT3G28580 - aaa-type atpase family protein<br>AT3G53600 - c2h2-type zinc finger protein<br>AT2G37430 - zinc finger protein zat11<br>AT5G02490 - heat shock protein 70<br>AT3G02840 - hypothetical protein<br>AT2G38340 - dehydration-responsive element-binding protein 2e<br>AT3G21670 - nitrate transporter 1.3<br>AT1G72520 - lipoxygenase 4 |
| GO:0009743 | response to carbohydrate | 7.96E-4 | 8.25E-2 | 6.68 (11869,30,296,5) | [-] Hide genes<br><br>AT3G12700 - aspartyl protease family protein<br>AT3G22550 - hypothetical protein<br>AT4G17670 - hypothetical protein<br>AT1G22500 - putative c3hc4-type ring zinc finger protein<br>AT4G37220 - cold acclimation protein wcor413 |
| GO:0019748 | secondary metabolic process | 9.25E-4 | 9.34E-2 | 5.23 (11869,46,296,6) | [-] Hide genes<br><br>AT5G61590 - ethylene-responsive transcription factor erf107<br>AT1G64160 - dirigent-like protein dir5<br>AT5G05340 - peroxidase 52<br>AT5G63560 - hxxxd-type acyl-transferase-like protein<br>AT2G37360 - abc transporter g family member 2<br>AT5G64120 - peroxidase 71 |
