## Supplementary material for "Evidences for a nutritional role of iodine in plants": Table S9

| GO term | Description | P-value | FDR q-value | Enrichment (N, B, n, b) | Genes |
|---|---|---|---|---|---|
| GO:0048037 | cofactor binding | 9.08E-12 | 1.58E-8 | 4.68 (11869,240,296,28) | [-] Hide genes<br><br>AT1G30700 - fad-binding berberine family protein<br>AT4G08780 - peroxidase 38<br>AT4G20860 - fad-binding berberine family protein<br>AT4G36430 - peroxidase 49<br>AT1G12200 - putative flavin monooxygenase.<br>AT4G12280 - copper amine oxidase family protein<br>AT5G64120 - peroxidase 71<br>AT4G20830 - fad-binding berberine family protein<br>AT5G64110 - peroxidase 70<br>AT5G06730 - peroxidase<br>AT4G15610 - hypothetical protein<br>AT5G14130 - peroxidase 55<br>AT5G19880 - peroxidase<br>AT5G05340 - peroxidase 52<br>AT2G46750 - d-arabinono-1,4-lactone oxidase-like protein<br>AT2G18140 - peroxidase 14<br>AT5G38710 - proline dehydrogenase 2<br>AT1G26380 - fad-binding and bbe domain-containing protein<br>AT1G34510 - peroxidase 8<br>AT5G44400 - fad-binding and bbe domain-containing protein<br>AT1G26390 - fad-binding berberine family protein<br>AT2G35380 - peroxidase 20<br>AT1G26420 - fad-binding and bbe domain-containing protein<br>AT1G26410 - fad-binding and bbe domain-containing protein<br>AT3G14415 - peroxisomal (s)-2-hydroxy-acid oxidase glo2<br>AT1G14550 - peroxidase 5<br>AT5G39580 - peroxidase 62<br>AT1G14540 - peroxidase 4 |

| | | | | | |
|---|---|---|---|---|---|
| GO:0004601 | peroxidase activity | 2.63E-10 | 2.28E-7 | 9.05 (11869,62,296,14) | [-] Hide genes<br><br>AT4G08780 - peroxidase 38<br>AT5G06730 - peroxidase<br>AT5G14130 - peroxidase 55<br>AT5G05340 - peroxidase 52<br>AT5G19880 - peroxidase<br>AT2G18140 - peroxidase 14<br>AT1G34510 - peroxidase 8<br>AT2G35380 - peroxidase 20<br>AT4G36430 - peroxidase 49<br>AT5G64120 - peroxidase 71<br>AT1G14550 - peroxidase 5<br>AT5G39580 - peroxidase 62<br>AT1G14540 - peroxidase 4<br>AT5G64110 - peroxidase 70 |
| GO:0016684 | oxidoreductase activity, acting on peroxide as acceptor | 3.31E-10 | 1.91E-7 | 8.91 (11869,63,296,14) | [-] Hide genes<br><br>AT4G08780 - peroxidase 38<br>AT5G06730 - peroxidase<br>AT5G14130 - peroxidase 55<br>AT5G19880 - peroxidase<br>AT5G05340 - peroxidase 52<br>AT2G18140 - peroxidase 14<br>AT1G34510 - peroxidase 8<br>AT2G35380 - peroxidase 20<br>AT4G36430 - peroxidase 49<br>AT5G64120 - peroxidase 71<br>AT1G14550 - peroxidase 5<br>AT5G39580 - peroxidase 62<br>AT1G14540 - peroxidase 4<br>AT5G64110 - peroxidase 70 |

| | | | | | |
|---|---|---|---|---|---|
| GO:0016209 | antioxidant activity | 1.19E-9 | 5.19E-7 | 8.14 (11869,69,296,14) | [-] Hide genes<br><br>AT4G08780 - peroxidase 38<br>AT5G06730 - peroxidase<br>AT5G14130 - peroxidase 55<br>AT5G05340 - peroxidase 52<br>AT5G19880 - peroxidase<br>AT2G18140 - peroxidase 14<br>AT1G34510 - peroxidase 8<br>AT2G35380 - peroxidase 20<br>AT4G36430 - peroxidase 49<br>AT5G64120 - peroxidase 71<br>AT1G14550 - peroxidase 5<br>AT5G39580 - peroxidase 62<br>AT1G14540 - peroxidase 4<br>AT5G64110 - peroxidase 70 |
| GO:0020037 | heme binding | 1.78E-9 | 6.18E-7 | 7.91 (11869,71,296,14) | [-] Hide genes<br><br>AT4G08780 - peroxidase 38<br>AT5G06730 - peroxidase<br>AT5G14130 - peroxidase 55<br>AT5G05340 - peroxidase 52<br>AT5G19880 - peroxidase<br>AT2G18140 - peroxidase 14<br>AT1G34510 - peroxidase 8<br>AT2G35380 - peroxidase 20<br>AT4G36430 - peroxidase 49<br>AT5G64120 - peroxidase 71<br>AT1G14550 - peroxidase 5<br>AT1G14540 - peroxidase 4<br>AT5G39580 - peroxidase 62<br>AT5G64110 - peroxidase 70 |

| | | | | | |
|---|---|---|---|---|---|
| GO:0046906 | tetrapyrrole binding | 2.16E-9 | 6.25E-7 | 7.80 (11869,72,296,14) | [-] Hide genes<br><br>AT4G08780 - peroxidase 38<br>AT5G06730 - peroxidase<br>AT5G14130 - peroxidase 55<br>AT5G05340 - peroxidase 52<br>AT5G19880 - peroxidase<br>AT2G18140 - peroxidase 14<br>AT1G34510 - peroxidase 8<br>AT2G35380 - peroxidase 20<br>AT4G36430 - peroxidase 49<br>AT5G64120 - peroxidase 71<br>AT1G14550 - peroxidase 5<br>AT1G14540 - peroxidase 4<br>AT5G39580 - peroxidase 62<br>AT5G64110 - peroxidase 70 |
| GO:0071949 | FAD binding | 4.46E-9 | 1.11E-6 | 12.15 (11869,33,296,10) | [-] Hide genes<br><br>AT1G26380 - fad-binding and bbe domain-containing protein<br>AT1G30700 - fad-binding berberine family protein<br>AT5G44400 - fad-binding and bbe domain-containing protein<br>AT4G20860 - fad-binding berberine family protein<br>AT1G26390 - fad-binding berberine family protein<br>AT1G26420 - fad-binding and bbe domain-containing protein<br>AT1G26410 - fad-binding and bbe domain-containing protein<br>AT2G46750 - d-arabinono-1,4-lactone oxidase-like protein<br>AT4G20830 - fad-binding berberine family protein<br>AT5G38710 - proline dehydrogenase 2 |
| GO:0050660 | flavin adenine dinucleotide binding | 1.86E-7 | 4.04E-5 | 7.48 (11869,59,296,11) | [-] Hide genes<br><br>AT1G30700 - fad-binding berberine family protein<br>AT1G26380 - fad-binding and bbe domain-containing protein |

| | | | | | |
|---|---|---|---|---|---|
| | | | | | AT5G44400 - fad-binding and bbe domain-containing protein<br>AT4G20860 - fad-binding berberine family protein<br>AT1G26390 - fad-binding berberine family protein<br>AT1G26420 - fad-binding and bbe domain-containing protein<br>AT1G26410 - fad-binding and bbe domain-containing protein<br>AT1G12200 - putative flavin monooxygenase.<br>AT2G46750 - d-arabinono-1,4-lactone oxidase-like protein<br>AT4G20830 - fad-binding berberine family protein<br>AT5G38710 - proline dehydrogenase 2 |
| GO:0050662 | coenzyme binding | 7.14E-5 | 1.38E-2 | 3.82 (11869,126,296,12) | [-] Hide genes<br><br>AT1G30700 - fad-binding berberine family protein<br>AT1G26380 - fad-binding and bbe domain-containing protein<br>AT4G20860 - fad-binding berberine family protein<br>AT5G44400 - fad-binding and bbe domain-containing protein<br>AT1G26390 - fad-binding berberine family protein<br>AT1G26420 - fad-binding and bbe domain-containing protein<br>AT1G26410 - fad-binding and bbe domain-containing protein<br>AT1G12200 - putative flavin monooxygenase.<br>AT3G14415 - peroxisomal (s)-2-hydroxy-acid oxidase glo2<br>AT2G46750 - d-arabinono-1,4-lactone oxidase-like protein<br>AT4G20830 - fad-binding berberine family protein<br>AT5G38710 - proline dehydrogenase 2 |
| GO:0016491 | oxidoreductase activity | 8.36E-5 | 1.45E-2 | 2.32 (11869,433,296,25) | [-] Hide genes<br><br>AT4G08780 - peroxidase 38<br>AT1G66800 - alcohol dehydrogenase-like protein<br>AT4G36430 - peroxidase 49<br>AT3G02610 - acyl-[acyl-carrier-protein] desaturase<br>AT1G12200 - putative flavin monooxygenase.<br>AT4G12280 - copper amine oxidase family protein<br>AT5G64120 - peroxidase 71 |

| | | | | | |
|---|---|---|---|---|---|
| | | | | | AT4G20830 - fad-binding berberine family protein<br>AT1G72520 - lipoxygenase 4<br>AT5G64110 - peroxidase 70<br>AT5G06730 - peroxidase<br>AT5G14130 - peroxidase 55<br>AT5G05340 - peroxidase 52<br>AT4G39830 - putative l-ascorbate oxidase<br>AT5G19880 - peroxidase<br>AT2G46750 - d-arabinono-1,4-lactone oxidase-like protein<br>AT2G18140 - peroxidase 14<br>AT5G38710 - proline dehydrogenase 2<br>AT1G34510 - peroxidase 8<br>AT2G35380 - peroxidase 20<br>AT3G14415 - peroxisomal (s)-2-hydroxy-acid oxidase glo2<br>AT1G14550 - peroxidase 5<br>AT5G38900 - thioredoxin superfamily protein<br>AT5G39580 - peroxidase 62<br>AT1G14540 - peroxidase 4 |
| GO:0004805 | trehalose-phosphatase activity | 2.9E-4 | 4.59E-2 | 20.05 (11869,6,296,3) | [-] Hide genes<br>AT4G39770 - probable trehalose-phosphate phosphatase h<br>AT2G22190 - probable trehalose-phosphate phosphatase e<br>AT1G35910 - probable trehalose-phosphate phosphatase d |
