## Supplementary material for "Evidences for a nutritional role of iodine in plants": Table S10

| Accession | Description | Sum PEP Score | Coverage [%] | # Peptides | # PSMs | # Unique Peptides | # AAs | MW [kDa] | calc. pI | Score Mascot |
|---|---|---|---|---|---|---|---|---|---|---|
| ATCG00480.1 | ATP synthase subunit beta | 1118.679 | 94 | 55 | 2681 | 53 | 498 | 53.9 | 5.5 | 89814 |
| AT4G10340.1 | light harvesting complex of photosystem II 5 | 481.351 | 80 | 26 | 1930 | 26 | 280 | 30.1 | 6.37 | 53733 |
| AT1G29930.1 | chlorophyll A/B binding protein 1 | 285.348 | 90 | 16 | 1841 | 1 | 267 | 28.2 | 5.66 | 50302 |
| AT1G29910.1 | chlorophyll A/B binding protein 3 | 285.194 | 91 | 16 | 1836 | 1 | 267 | 28.2 | 5.43 | 50031 |
| ATCG00120.1 | ATP synthase subunit alpha | 812.99 | 66 | 35 | 1669 | 34 | 507 | 55.3 | 5.25 | 55652 |
| AT2G34420.1 | photosystem II light harvesting complex gene B1B2 | 224.566 | 80 | 15 | 1489 | 1 | 265 | 28 | 5.44 | 41172 |
| ATCG00680.1 | photosystem II reaction center protein B | 481.679 | 44 | 26 | 1458 | 26 | 508 | 56 | 6.89 | 51434 |
| AT2G05070.1 | photosystem II light harvesting complex gene 2.2 | 238.416 | 67 | 12 | 1407 | 1 | 265 | 28.6 | 5.43 | 36589 |
| AT2G34430.1 | light-harvesting chlorophyll-protein complex II subunit B1 | 230.469 | 79 | 14 | 1402 | 2 | 266 | 28.2 | 5.27 | 39385 |
| AT3G27690.1 | photosystem II light harvesting complex gene 2.3 | 233.11 | 67 | 12 | 1373 | 1 | 266 | 28.8 | 5.9 | 35845 |
| AT2G05100.1 | photosystem II light harvesting complex gene 2.1 | 229.129 | 67 | 12 | 1366 | 1 | 265 | 28.6 | 5.43 | 35569 |
| AT1G15820.1 | light harvesting complex photosystem II subunit 6 | 275.792 | 59 | 16 | 1202 | 16 | 258 | 27.5 | 7.4 | 38076 |
| AT5G01530.1 | light harvesting complex photosystem II | 231.779 | 51 | 14 | 1018 | 10 | 290 | 31.1 | 6.14 | 32354 |
| ATCG00280.1 | photosystem II reaction center protein C | 297.223 | 39 | 22 | 998 | 22 | 473 | 51.8 | 7.2 | 31312 |
| ATCG00540.1 | photosynthetic electron transfer A | 381.171 | 61 | 23 | 919 | 23 | 320 | 35.3 | 8.29 | 29544 |
| AT5G54270.1 | light-harvesting chlorophyll B-binding protein 3 | 222.252 | 72 | 13 | 900 | 11 | 265 | 28.7 | 5.12 | 30414 |
| AT5G42270.1 | FtsH extracellular protease family | 538.389 | 58 | 37 | 889 | 13 | 704 | 75.2 | 5.49 | 31084 |
| AT4G21280.1 | photosystem II subunit QA | 289.351 | 59 | 19 | 858 | 17 | 223 | 23.8 | 9.64 | 21265 |
| AT1G50250.1 | FTSH protease 1 | 502.229 | 56 | 37 | 748 | 13 | 716 | 76.7 | 5.83 | 26247 |
| AT1G06680.1 | photosystem II subunit P-1 | 316.397 | 59 | 13 | 704 | 11 | 263 | 28.1 | 7.39 | 20344 |
| AT3G08940.2 | light harvesting complex photosystem II | 179.896 | 68 | 14 | 666 | 10 | 287 | 31.2 | 6.24 | 18601 |
| AT3G50820.1 | photosystem II subunit O-2 | 374.438 | 64 | 23 | 652 | 11 | 331 | 35 | 6.16 | 18509 |
| ATCG00130.1 | ATPase, F0 complex, subunit B/B', bacterial/chloroplast | 238.426 | 64 | 15 | 647 | 15 | 184 | 21 | 8.13 | 16277 |
| AT4G05180.1 | photosystem II subunit Q-2 | 248.942 | 50 | 13 | 638 | 11 | 230 | 24.6 | 9.72 | 19274 |
| ATCG00270.1 | photosystem II reaction center protein D | 237.544 | 31 | 12 | 636 | 12 | 353 | 39.5 | 5.74 | 19072 |
| AT2G39730.1 | rubisco activase | 353.503 | 72 | 26 | 623 | 26 | 474 | 51.9 | 6.15 | 23207 |
| AT5G66570.1 | PS II oxygen-evolving complex 1 | 346.584 | 61 | 22 | 579 | 10 | 332 | 35.1 | 5.66 | 15283 |
| AT5G66190.1 | ferredoxin-NADP(+)-oxidoreductase 1 | 230.644 | 61 | 22 | 505 | 20 | 360 | 40.3 | 8.13 | 15297 |
| AT1G20020.1 | ferredoxin-NADP(+)-oxidoreductase 2 | 315.147 | 59 | 21 | 491 | 19 | 369 | 41.1 | 8.31 | 14846 |
| AT4G04640.1 | ATPase, F1 complex, gamma subunit protein | 311.259 | 72 | 24 | 475 | 24 | 373 | 40.9 | 8 | 15368 |
| ATCG00020.1 | photosystem II reaction center protein A | 128.76 | 28 | 10 | 460 | 10 | 353 | 38.9 | 5.25 | 11818 |
| AT1G31330.1 | photosystem I subunit F | 156.248 | 47 | 12 | 455 | 12 | 221 | 24.2 | 9.54 | 11771 |
| AT1G74470.1 | Pyridine nucleotide-disulphide oxidoreductase family protein | 311.65 | 58 | 26 | 454 | 26 | 467 | 51.8 | 8.91 | 16041 |
| ATCG00490.1 | ribulose-bisphosphate carboxylases | 424.17 | 64 | 34 | 427 | 34 | 479 | 52.9 | 6.29 | 10286 |
| AT4G27440.1 | protochlorophyllide oxidoreductase B | 298.869 | 63 | 20 | 425 | 12 | 401 | 43.3 | 9.16 | 17218 |
| AT1G79040.1 | photosystem II subunit R | 150.365 | 38 | 5 | 424 | 5 | 140 | 14.6 | 9.63 | 11962 |
| AT1G61520.1 | photosystem I light harvesting complex gene 3 | 127.049 | 59 | 12 | 404 | 12 | 273 | 29.2 | 8.68 | 12823 |
| AT3G47470.1 | light-harvesting chlorophyll-protein complex I subunit A4 | 205.004 | 67 | 13 | 359 | 13 | 251 | 27.7 | 6.7 | 10201 |
| AT1G54780.1 | thylakoid lumen 18.3 kDa protein | 160.846 | 44 | 12 | 328 | 12 | 285 | 31.1 | 8.84 | 8800 |

| | | | | | | | | | | |
|---|---|---|---|---|---|---|---|---|---|---|
| AT4G22890.1 | PGR5-LIKE A | 244.415 | 59 | 17 | 285 | 2 | 324 | 35.7 | 5.29 | 9484 |
| AT1G71500.1 | Rieske (2Fe-2S) domain-containing protein | 192.004 | 56 | 17 | 285 | 17 | 287 | 31.7 | 8.73 | 11268 |
| AT4G22890.5 | PGR5-LIKE A | 222.462 | 58 | 16 | 267 | 1 | 322 | 35.5 | 5.29 | 9040 |
| AT4G03280.1 | photosynthetic electron transfer C | 128.878 | 62 | 9 | 255 | 9 | 229 | 24.4 | 8.54 | 8202 |
| ATCG00340.1 | Photosystem I, PsaA/PsaB protein | 206 | 25 | 15 | 233 | 15 | 734 | 82.4 | 7.4 | 6655 |
| AT4G38970.1 | fructose-bisphosphate aldolase 2 | 257.65 | 58 | 21 | 225 | 12 | 398 | 43 | 7.24 | 6845 |
| AT4G02770.1 | photosystem I subunit D-1 | 177.844 | 72 | 18 | 207 | 18 | 208 | 22.6 | 9.77 | 5935 |
| AT3G23400.1 | Plastid-lipid associated protein PAP / fibrillin family protein | 154.937 | 54 | 11 | 206 | 11 | 284 | 30.4 | 6.13 | 5744 |
| AT5G54190.1 | protochlorophyllide oxidoreductase A | 133.889 | 27 | 10 | 201 | 2 | 405 | 43.8 | 9.36 | 8478 |
| AT2G20890.1 | photosystem II reaction center PSB29 protein | 121.071 | 48 | 14 | 180 | 14 | 300 | 33.8 | 9.13 | 5049 |
| AT2G30790.1 | photosystem II subunit P-2 | 98.991 | 43 | 4 | 126 | 2 | 125 | 13.4 | 6.07 | 3883 |
| AT4G28750.1 | Photosystem I reaction centre subunit IV / PsaE protein | 84.314 | 57 | 6 | 125 | 5 | 143 | 15 | 9.92 | 4659 |
| AT1G03600.1 | photosystem II family protein | 70.229 | 33 | 4 | 123 | 4 | 174 | 18.8 | 9.88 | 3909 |
| AT1G52230.1 | photosystem I subunit H2 | 88.019 | 57 | 6 | 111 | 2 | 145 | 15.3 | 9.91 | 3952 |
| AT2G20260.1 | photosystem I subunit E-2 | 123.144 | 46 | 5 | 110 | 4 | 145 | 15.2 | 9.95 | 4815 |
| AT3G16140.1 | photosystem I subunit H-1 | 76.57 | 57 | 6 | 109 | 2 | 145 | 15.2 | 9.95 | 4043 |
| AT1G67090.1 | ribulose bisphosphate carboxylase small chain 1A | 123.098 | 57 | 14 | 108 | 11 | 180 | 20.2 | 7.71 | 2838 |
| AT5G07020.1 | proline-rich family protein | 69.27 | 23 | 5 | 84 | 5 | 235 | 24.4 | 4.75 | 3304 |
| ATCG01060.1 | iron-sulfur cluster binding;electron carriers;4 iron, 4 sulfur clust | 79.419 | 70 | 6 | 79 | 6 | 81 | 9 | 7.08 | 2687 |
| ATCG00710.1 | photosystem II reaction center protein H | 37.726 | 37 | 3 | 77 | 3 | 73 | 7.7 | 6.55 | 1137 |

| Accession | Description | Sum PEP Score | Coverage [%] | # Peptides | # PSMs | # Unique Peptides | # AAs | MW [kDa] | calc. pI | Score Mascot |
|---|---|---|---|---|---|---|---|---|---|---|
| ATCG00490.1 | ribulose-bisphosphate carboxylases | 609.949 | 60 | 30 | 725 | 30 | 479 | 52.9 | 6.29 | 24124 |

| Accession | Description | Sum PEP Score | Coverage [%] | # Peptides | # PSMs | # Unique Peptid | # AAs | MW [kDa] | calc. pI | Score Mascot |
|---|---|---|---|---|---|---|---|---|---|---|
| ATCG00490.1 | ribulose-bisphosphate carboxylases | 990.276 | 66 | 38 | 6603 | 38 | 479 | 52.9 | 6.29 | 232891 |
| AT1G67090.1 | ribulose bisphosphate carboxylase small chain 1A | 312.759 | 64 | 17 | 1946 | 11 | 180 | 20.2 | 7.71 | 69414 |
| AT5G38410.1 | Ribulose bisphosphate carboxylase (small chain) family protein | 320.98 | 65 | 14 | 1820 | 6 | 181 | 20.3 | 8.05 | 69681 |
| AT5G38430.1 | Ribulose bisphosphate carboxylase (small chain) family protein | 295.171 | 65 | 14 | 1695 | 6 | 181 | 20.3 | 7.71 | 63557 |
| AT1G06680.1 | photosystem II subunit P-1 | 321.279 | 55 | 10 | 486 | 8 | 263 | 28.1 | 7.39 | 22113 |
| AT2G34430.1 | light-harvesting chlorophyll-protein complex II subunit B1 | 157.356 | 47 | 8 | 316 | 4 | 266 | 28.2 | 5.27 | 14347 |

| Accession | Description | Sum PEP Score | Coverage [%] | # Peptides | # PSMs | # Unique Peptides | # AAs | MW [kDa] | calc. pI | Score Mascot |
|---|---|---|---|---|---|---|---|---|---|---|
| ATCG00480.1 | ATP synthase subunit beta | 1030.446 | 91 | 36 | 863 | 35 | 498 | 53.9 | 5.5 | 29226 |
| AT3G50820.1 | photosystem II subunit O-2 | 379.107 | 59 | 20 | 348 | 9 | 331 | 35 | 6.16 | 11742 |
| AT5G66570.1 | PS II oxygen-evolving complex 1 | 351.442 | 57 | 19 | 288 | 8 | 332 | 35.1 | 5.66 | 9014 |

| Accession | Description | Sum PEP Score | Coverage [%] | # Peptides | # PSMs | # Unique Peptides | # AAs | MW [kDa] | calc. pI | Score Mascot |
|---|---|---|---|---|---|---|---|---|---|---|
| AT1G29930.1 | chlorophyll A/B binding protein 1 | 294.004 | 90 | 17 | 922 | 6 | 267 | 28.2 | 5.66 | 19926 |
| AT2G34420.1 | photosystem II light harvesting complex gene B1B2 | 227.193 | 66 | 14 | 810 | 1 | 265 | 28 | 5.44 | 17876 |
| ATCG00680.1 | photosystem II reaction center protein B | 367.635 | 45 | 22 | 546 | 22 | 508 | 56 | 6.89 | 16666 |
| ATCG00020.1 | photosystem II reaction center protein A | 109.972 | 24 | 9 | 179 | 9 | 353 | 38.9 | 5.25 | 4314 |

| Accession | Description | Sum PEP Score | Coverage [%] | # Peptides | # PSMs | # Unique Peptides | # AAs | MW [kDa] | calc. pI | Score Mascot |
|---|---|---|---|---|---|---|---|---|---|---|
| ATCG00490.1 | ribulose-bisphosphate carboxylases | 1463.516 | 72 | 55 | 11402 | 55 | 479 | 52.9 | 6.29 | 334482 |
| AT1G67090.1 | ribulose bisphosphate carboxylase small chain 1A | 531.753 | 78 | 26 | 5620 | 15 | 180 | 20.2 | 7.71 | 150482 |
| AT2G39730.1 | rubisco activase | 1757.686 | 76 | 38 | 4073 | 1 | 474 | 51.9 | 6.15 | 209005 |
| AT5G38410.1 | Ribulose bisphosphate carboxylase (small chain) family protein | 501.611 | 80 | 24 | 4043 | 9 | 181 | 20.3 | 8.05 | 115188 |
| AT2G39730.2 | rubisco activase | 1714.37 | 78 | 38 | 4024 | 1 | 446 | 49.1 | 7.68 | 207178 |
| AT5G38430.1 | Ribulose bisphosphate carboxylase (small chain) family protein | 476.656 | 77 | 23 | 3616 | 6 | 181 | 20.3 | 7.71 | 99885 |
| AT3G50820.1 | photosystem II subunit O-2 | 569.48 | 69 | 28 | 3225 | 13 | 331 | 35 | 6.16 | 95694 |
| AT5G66570.1 | PS II oxygen-evolving complex 1 | 516.715 | 66 | 26 | 3168 | 11 | 332 | 35.1 | 5.66 | 91634 |
| AT1G29930.1 | chlorophyll A/B binding protein 1 | 255.455 | 63 | 12 | 2321 | 1 | 267 | 28.2 | 5.66 | 78499 |
| AT1G29910.1 | chlorophyll A/B binding protein 3 | 256.525 | 64 | 12 | 2315 | 1 | 267 | 28.2 | 5.43 | 78180 |
| AT2G34420.1 | photosystem II light harvesting complex gene B1B2 | 170.117 | 53 | 11 | 1965 | 1 | 265 | 28 | 5.44 | 66535 |
| ATCG00680.1 | photosystem II reaction center protein B | 656.431 | 41 | 22 | 1932 | 22 | 508 | 56 | 6.89 | 86949 |
| AT3G01500.1 | carbonic anhydrase 1 | 633.975 | 90 | 31 | 1907 | 1 | 270 | 29.5 | 5.73 | 74496 |
| AT3G01500.2 | carbonic anhydrase 1 | 615.417 | 76 | 31 | 1871 | 1 | 347 | 37.4 | 5.99 | 73033 |
| ATCG00280.1 | photosystem II reaction center protein C | 533.022 | 35 | 18 | 1859 | 18 | 473 | 51.8 | 7.2 | 82036 |
| AT5G26000.1 | thioglucoside glucohydrolase 1 | 801.177 | 61 | 38 | 1846 | 36 | 541 | 61.1 | 5.92 | 63080 |
| AT4G21280.1 | photosystem II subunit QA | 487.348 | 63 | 24 | 1699 | 3 | 223 | 23.8 | 9.64 | 60416 |
| AT5G14740.2 | carbonic anhydrase 2 | 619.456 | 83 | 31 | 1665 | 3 | 259 | 28.3 | 5.53 | 64596 |
| AT4G05180.1 | photosystem II subunit Q-2 | 426.381 | 59 | 17 | 1660 | 15 | 230 | 24.6 | 9.72 | 54621 |
| AT4G21280.2 | photosystem II subunit QA | 415.687 | 63 | 22 | 1484 | 1 | 224 | 23.9 | 9.64 | 51415 |
| AT5G14740.4 | carbonic anhydrase 2 | 523.244 | 60 | 28 | 1425 | 1 | 330 | 36.5 | 7.64 | 52999 |
| AT4G33010.1 | glycine decarboxylase P-protein 1 | 1329.66 | 66 | 54 | 1211 | 28 | 1037 | 112.9 | 6.98 | 44490 |
| AT2G26080.1 | glycine decarboxylase P-protein 2 | 1101.214 | 54 | 50 | 845 | 24 | 1044 | 113.7 | 6.65 | 31957 |
| AT1G20340.1 | Cupredoxin superfamily protein | 262.787 | 41 | 3 | 689 | 3 | 167 | 17 | 5.2 | 26446 |
| AT1G11860.1 | Glycine cleavage T-protein family | 757.317 | 70 | 29 | 686 | 29 | 408 | 44.4 | 8.37 | 24544 |
| AT1G20020.1 | ferredoxin-NADP(+)-oxidoreductase 2 | 566.35 | 62 | 26 | 597 | 21 | 369 | 41.1 | 8.31 | 21228 |
| ATCG00540.1 | photosynthetic electron transfer A | 361.258 | 52 | 20 | 578 | 20 | 320 | 35.3 | 8.29 | 20649 |
| AT3G27830.1 | ribosomal protein L12-A | 114.02 | 35 | 12 | 473 | 12 | 191 | 20.1 | 5.64 | 15650 |
| AT2G05520.2 | glycine-rich protein 3 | 20.323 | 56 | 4 | 13 | 1 | 138 | 13.6 | 8.66 | 661 |

| Accession | Description | Sum PEP Score | Coverage [%] | # Peptides | # PSMs | # Unique Peptides | # AAs | MW [kDa] | calc. pI | Score Mascot |
|---|---|---|---|---|---|---|---|---|---|---|
| ATCG00490.1 | ribulose-bisphosphate carboxylases | 1057.6 | 70 | 45 | 7864 | 40 | 479 | 52.9 | 6.29 | 252479 |
| ATMG00280.1 | Ribulose bisphosphate carboxylase large chain, catalytic domain | 240.11 | 50 | 7 | 1732 | 2 | 110 | 12.7 | 8.95 | 70001 |
| AT4G38740.1 | rotamase CYP 1 | 263.15 | 84 | 13 | 166 | 6 | 172 | 18.4 | 7.81 | 6123 |
| AT2G21130.1 | Cyclophilin-like peptidyl-prolyl cis-trans isomerase family protein | 206.1 | 62 | 10 | 142 | 3 | 174 | 18.5 | 8.16 | 5185 |

| Accession | Description | Sum PEP Score | Coverage [%] | # Peptides | # PSMs | # Unique Peptid | # AAs | MW [kDa] | calc. pI | Score Mascot |
|---|---|---|---|---|---|---|---|---|---|---|
| ATCG00490.1 | ribulose-bisphosphate carboxylases | 1423.708 | 74 | 54 | 10165 | 54 | 479 | 52.9 | 6.29 | 297807 |
| AT1G67090.1 | ribulose bisphosphate carboxylase small chain 1A | 497.813 | 69 | 24 | 4699 | 16 | 180 | 20.2 | 7.71 | 129482 |
| AT2G39730.1 | rubisco activase | 1824.046 | 83 | 44 | 4101 | 2 | 474 | 51.9 | 6.15 | 199771 |
| AT2G39730.2 | rubisco activase | 1781.982 | 84 | 43 | 4045 | 1 | 446 | 49.1 | 7.68 | 197592 |
| AT5G38410.1 | Ribulose bisphosphate carboxylase (small chain) family protein | 475.511 | 75 | 22 | 3630 | 8 | 181 | 20.3 | 8.05 | 108373 |
| AT5G38430.1 | Ribulose bisphosphate carboxylase (small chain) family protein | 433.294 | 69 | 20 | 3228 | 6 | 181 | 20.3 | 7.71 | 93742 |
| AT3G50820.1 | photosystem II subunit O-2 | 503.292 | 69 | 28 | 2674 | 13 | 331 | 35 | 6.16 | 79844 |
| AT5G66570.1 | PS II oxygen-evolving complex 1 | 491.313 | 67 | 26 | 2526 | 11 | 332 | 35.1 | 5.66 | 72902 |
| ATCG00680.1 | photosystem II reaction center protein B | 685.334 | 41 | 24 | 2124 | 24 | 508 | 56 | 6.89 | 94154 |
| AT1G29930.1 | chlorophyll A/B binding protein 1 | 278.52 | 70 | 13 | 2005 | 1 | 267 | 28.2 | 5.66 | 71213 |
| AT1G29910.1 | chlorophyll A/B binding protein 3 | 278.33 | 71 | 13 | 2000 | 1 | 267 | 28.2 | 5.43 | 70969 |
| ATCG00280.1 | photosystem II reaction center protein C | 563.275 | 37 | 18 | 1743 | 18 | 473 | 51.8 | 7.2 | 77507 |
| AT3G01500.1 | carbonic anhydrase 1 | 621.835 | 89 | 33 | 1706 | 22 | 270 | 29.5 | 5.73 | 66675 |
| AT4G21280.1 | photosystem II subunit QA | 447.595 | 60 | 23 | 1507 | 21 | 223 | 23.8 | 9.64 | 59744 |
| AT5G14740.2 | carbonic anhydrase 2 | 614.793 | 87 | 32 | 1424 | 21 | 259 | 28.3 | 5.53 | 56337 |
| AT4G05180.1 | photosystem II subunit Q-2 | 404.572 | 60 | 18 | 1355 | 16 | 230 | 24.6 | 9.72 | 46333 |
| AT4G33010.1 | glycine decarboxylase P-protein 1 | 1155.235 | 64 | 49 | 1133 | 28 | 1037 | 112.9 | 6.98 | 41829 |
| AT1G31330.1 | photosystem I subunit F | 225.524 | 43 | 10 | 853 | 10 | 221 | 24.2 | 9.54 | 28333 |
| AT2G26080.1 | glycine decarboxylase P-protein 2 | 870.727 | 52 | 42 | 789 | 21 | 1044 | 113.7 | 6.65 | 29491 |
| AT1G11860.1 | Glycine cleavage T-protein family | 734.918 | 70 | 30 | 718 | 30 | 408 | 44.4 | 8.37 | 27002 |
| ATCG00540.1 | photosynthetic electron transfer A | 374.624 | 73 | 23 | 570 | 23 | 320 | 35.3 | 8.29 | 20305 |
| AT1G20340.1 | Cupredoxin superfamily protein | 219.526 | 47 | 4 | 537 | 4 | 167 | 17 | 5.2 | 20150 |

| Accession | Description | Sum PEP Score | Coverage [%] | # Peptides | # PSMs | # Unique Peptides | # AAs | MW [kDa] | calc. pI | Score Mascot |
|---|---|---|---|---|---|---|---|---|---|---|
| ATMG00280.1 | Ribulose bisphosphate carboxylase large chain, catalytic domain | 259.63 | 55 | 8 | 1515 | 2 | 110 | 12.7 | 8.95 | 57215 |
| AT2G34420.1 | photosystem II light harvesting complex gene B1B2 | 72.68 | 51 | 8 | 398 | 1 | 265 | 28 | 5.44 | 12434 |
| ATCG00280.1 | photosystem II reaction center protein C | 301.59 | 35 | 17 | 393 | 17 | 473 | 51.8 | 7.2 | 17155 |
| AT2G34430.1 | light-harvesting chlorophyll-protein complex II subunit B1 | 87.876 | 52 | 9 | 355 | 3 | 266 | 28.2 | 5.27 | 10814 |
| AT1G65270.2 | unknown protein; FUNCTIONS IN: molecular_function unknown; INVOl | 45.061 | 34 | 9 | 10 | 9 | 292 | 32.5 | 6.43 | 355 |

| Accession | Description | Sum PEP Score | Coverage [%] | # Peptides | # PSMs | # Unique Peptides | # AAs | MW [kDa] | calc. pI | Score Mascot |
|---|---|---|---|---|---|---|---|---|---|---|
| ATCG00280.1 | photosystem II reaction center protein C | 290.184 | 36 | 17 | 616 | 17 | 473 | 51.8 | 7.2 | 26297 |
| AT2G07732.1 | Ribulose bisphosphate carboxylase large chain, catalytic domain | 41.505 | 38 | 5 | 352 | 2 | 116 | 13.1 | 8.47 | 11488 |
| AT1G65270.2 | unknown protein; FUNCTIONS IN: molecular_function unknown; INVOLVED I | 37.064 | 27 | 9 | 14 | 9 | 292 | 32.5 | 6.43 | 333 |

| Accession | Description | Sum PEP Score | Coverage [%] | # Peptides | # PSMs | # Unique Peptid | # AAs | MW [kDa] | calc. pI | Score Mascot |
|---|---|---|---|---|---|---|---|---|---|---|
| ATMG00280.1 | Ribulose bisphosphate carboxylase large chain, catalytic domain | 254.6 | 48 | 6 | 1404 | 2 | 110 | 12.7 | 8.95 | 53054 |
| AT3G09440.1 | Heat shock protein 70 (Hsp 70) family protein | 663.9 | 62 | 45 | 463 | 19 | 649 | 71.1 | 5.07 | 15552 |
| AT5G02490.1 | Heat shock protein 70 (Hsp 70) family protein | 487.8 | 53 | 34 | 381 | 10 | 653 | 71.3 | 5.12 | 13416 |
| AT3G12580.1 | heat shock protein 70 | 414.8 | 50 | 32 | 367 | 9 | 650 | 71.1 | 5.25 | 12346 |
| AT2G05520.4 | glycine-rich protein 3 | 21.84 | 51 | 3 | 12 | 1 | 124 | 12.3 | 8.47 | 499 |
| AT2G14835.1 | RING/U-box superfamily protein | 51.26 | 26 | 5 | 10 | 5 | 343 | 37.4 | 8.1 | 289 |

| Accession | Description | Sum PEP Score | Coverage [%] | # Peptides | # PSMs | # Unique Peptides | # AAs | MW [kDa] | calc. pI | Score Mascot |
|---|---|---|---|---|---|---|---|---|---|---|
| AT3G09260.1 | Glycosyl hydrolase superfamily protein | 777.704 | 60 | 27 | 3523 | 21 | 524 | 59.7 | 6.92 | 115469 |
| AT2G01520.1 | MLP-like protein 328 | 482.855 | 96 | 18 | 2221 | 10 | 151 | 17.5 | 5.73 | 65206 |
| AT4G23670.1 | Polyketide cyclase/dehydrase and lipid transport superfamily protein | 383.13 | 96 | 22 | 2117 | 16 | 151 | 17.5 | 6.37 | 57815 |
| AT5G17920.1 | Cobalamin-independent synthase family protein | 826.832 | 64 | 50 | 1655 | 27 | 765 | 84.3 | 6.51 | 63353 |
| AT1G66270.1 | Glycosyl hydrolase superfamily protein | 442.306 | 60 | 28 | 1597 | 16 | 524 | 59.6 | 7.02 | 43178 |
| AT1G07920.1 | GTP binding Elongation factor Tu family protein | 571.247 | 58 | 29 | 1587 | 29 | 449 | 49.5 | 9.11 | 53003 |
| AT5G02500.1 | heat shock cognate protein 70-1 | 1030.06 | 69 | 47 | 1514 | 15 | 651 | 71.3 | 5.12 | 61914 |
| AT5G08680.1 | ATP synthase alpha/beta family protein | 777.404 | 76 | 36 | 1411 | 34 | 559 | 59.8 | 6.49 | 63727 |
| AT1G78900.1 | vacuolar ATP synthase subunit A | 845.436 | 74 | 46 | 1248 | 46 | 623 | 68.8 | 5.24 | 52984 |
| AT3G52930.1 | Aldolase superfamily protein | 608.079 | 84 | 35 | 1245 | 21 | 358 | 38.5 | 6.46 | 49620 |
| AT1G66280.1 | Glycosyl hydrolase superfamily protein | 340.812 | 53 | 27 | 1185 | 16 | 524 | 59.7 | 7.21 | 26728 |
| AT5G28540.1 | heat shock protein 70 (Hsp 70) family protein | 804.721 | 63 | 53 | 1168 | 4 | 669 | 73.6 | 5.17 | 47114 |
| AT2G01530.1 | MLP-like protein 329 | 296.231 | 73 | 16 | 1161 | 9 | 151 | 17.6 | 5.55 | 28818 |
| AT5G42020.1 | Heat shock protein 70 (Hsp 70) family protein | 782.649 | 63 | 53 | 1143 | 5 | 668 | 73.5 | 5.21 | 45899 |
| AT3G09440.1 | Heat shock protein 70 (Hsp 70) family protein | 848.754 | 60 | 41 | 1137 | 18 | 649 | 71.1 | 5.07 | 44328 |
| AT5G02490.1 | Heat shock protein 70 (Hsp 70) family protein | 610.956 | 54 | 31 | 1084 | 9 | 653 | 71.3 | 5.12 | 46143 |
| AT3G03780.1 | methionine synthase 2 | 611.27 | 50 | 40 | 1065 | 18 | 765 | 84.5 | 6.51 | 42466 |
| AT3G32980.1 | Peroxidase superfamily protein | 349.946 | 50 | 20 | 1004 | 13 | 352 | 38.8 | 6.67 | 35476 |
| AT2G36530.1 | Enolase | 835.147 | 85 | 38 | 1003 | 37 | 444 | 47.7 | 5.77 | 43938 |
| AT3G12580.1 | heat shock protein 70 | 605.379 | 56 | 35 | 965 | 11 | 650 | 71.1 | 5.25 | 39772 |
| AT3G16460.1 | Mannose-binding lectin superfamily protein | 867.958 | 56 | 45 | 953 | 45 | 705 | 72.4 | 5.5 | 46447 |
| AT1G78850.1 | D-mannose binding lectin protein with Apple-like carbohydrate-bindir | 567.052 | 70 | 34 | 949 | 19 | 441 | 49 | 7.65 | 41327 |
| AT4G23680.1 | Polyketide cyclase/dehydrase and lipid transport superfamily protein | 278.189 | 74 | 16 | 931 | 11 | 151 | 17.5 | 6.34 | 27683 |
| AT1G04410.1 | Lactate/malate dehydrogenase family protein | 610.25 | 81 | 29 | 884 | 16 | 332 | 35.5 | 6.55 | 37406 |
| AT3G16420.1 | PYK10-binding protein 1 | 626.35 | 82 | 23 | 829 | 18 | 298 | 32.1 | 5.77 | 35063 |
| AT4G14060.1 | Polyketide cyclase/dehydrase and lipid transport superfamily protein | 256.608 | 73 | 13 | 806 | 8 | 151 | 17.3 | 5.94 | 23901 |
| AT4G30170.1 | Peroxidase family protein | 547.834 | 78 | 23 | 788 | 18 | 325 | 35.8 | 9.1 | 34691 |
| AT2G36460.1 | Aldolase superfamily protein | 504.503 | 74 | 28 | 753 | 14 | 358 | 38.4 | 7.39 | 31990 |
| AT4G16260.1 | Glycosyl hydrolase superfamily protein | 486.653 | 65 | 19 | 656 | 19 | 344 | 37.7 | 6.84 | 27313 |
| AT2G47730.1 | glutathione S-transferase phi 8 | 450.464 | 62 | 18 | 652 | 17 | 263 | 29.2 | 8.5 | 32056 |
| AT3G49120.1 | peroxidase CB | 383.58 | 57 | 19 | 596 | 13 | 353 | 38.8 | 7.64 | 22453 |
| AT1G49240.1 | actin 8 | 537.042 | 63 | 29 | 571 | 13 | 377 | 41.8 | 5.58 | 21875 |
| AT5G11670.1 | NADP-malic enzyme 2 | 456.934 | 65 | 26 | 562 | 16 | 588 | 64.4 | 6.42 | 24239 |
| AT4G37910.1 | mitochondrial heat shock protein 70-1 | 807.38 | 61 | 51 | 551 | 42 | 682 | 73 | 5.62 | 24729 |
| AT1G65930.1 | cytosolic NADP+-dependent isocitrate dehydrogenase | 544.257 | 75 | 36 | 545 | 29 | 410 | 45.7 | 6.57 | 24515 |
| AT3G55440.1 | triosephosphate isomerase | 399.943 | 79 | 19 | 535 | 18 | 254 | 27.2 | 5.5 | 22220 |
| AT5G09810.1 | actin 7 | 540.181 | 73 | 29 | 523 | 11 | 377 | 41.7 | 5.49 | 19531 |
| AT4G02520.1 | glutathione S-transferase PHI 2 | 289.459 | 87 | 16 | 505 | 8 | 212 | 24.1 | 6.35 | 19191 |
| AT1G07890.3 | ascorbate peroxidase 1 | 440.647 | 64 | 17 | 480 | 16 | 250 | 27.5 | 6.13 | 16767 |

| | | | | | | | | | | |
|---|---|---|---|---|---|---|---|---|---|---|
| AT5G43330.1 | Lactate/malate dehydrogenase family protein | 341.66 | 51 | 21 | 440 | 8 | 332 | 35.7 | 6.79 | 17506 |
| AT4G20260.4 | plasma-membrane associated cation-binding protein 1 | 190.187 | 72 | 17 | 425 | 17 | 226 | 24.7 | 4.96 | 16466 |
| AT5G17820.1 | Peroxidase superfamily protein | 437.404 | 73 | 20 | 415 | 20 | 313 | 34.1 | 10.05 | 16131 |
| AT5G03690.2 | Aldolase superfamily protein | 159.506 | 21 | 9 | 389 | 2 | 359 | 38.6 | 7.74 | 20172 |
| AT1G22840.1 | CYTOCHROME C-1 | 156.186 | 73 | 12 | 383 | 9 | 114 | 12.4 | 9.31 | 13874 |
| AT4G09000.1 | general regulatory factor 1 | 329.558 | 76 | 22 | 381 | 10 | 267 | 29.9 | 4.81 | 10714 |
| AT2G39310.1 | jacalin-related lectin 22 | 556.929 | 62 | 32 | 370 | 29 | 458 | 50.4 | 5.3 | 14522 |
| AT4G08770.1 | Peroxidase superfamily protein | 186.023 | 51 | 17 | 368 | 7 | 346 | 38.2 | 7.64 | 10218 |
| AT4G10040.1 | cytochrome c-2 | 95.964 | 71 | 9 | 342 | 6 | 112 | 12.2 | 9.32 | 11415 |
| AT3G01190.1 | Peroxidase superfamily protein | 408.113 | 67 | 24 | 333 | 23 | 321 | 34.9 | 8.95 | 13753 |
| AT1G74030.1 | enolase 1 | 413.006 | 53 | 25 | 326 | 24 | 477 | 51.4 | 6.13 | 15396 |
| AT5G38480.1 | general regulatory factor 3 | 265.733 | 75 | 19 | 326 | 10 | 255 | 28.6 | 4.79 | 8980 |
| AT1G09640.1 | Translation elongation factor EF1B, gamma chain | 386.939 | 49 | 22 | 324 | 12 | 414 | 46.6 | 5.48 | 13035 |
| AT3G16430.1 | jacalin-related lectin 31 | 205.441 | 41 | 14 | 319 | 9 | 296 | 32.2 | 6.29 | 13892 |
| AT5G16050.1 | general regulatory factor 5 | 267.933 | 72 | 20 | 308 | 8 | 268 | 30.2 | 4.81 | 8591 |
| AT3G16470.1 | Mannose-binding lectin superfamily protein | 434.127 | 67 | 26 | 302 | 26 | 451 | 48.5 | 5.26 | 15061 |
| AT3G12110.1 | actin-11 | 358.926 | 64 | 23 | 301 | 3 | 377 | 41.6 | 5.39 | 8974 |
| AT2G38390.1 | Peroxidase superfamily protein | 184.441 | 50 | 12 | 295 | 7 | 349 | 38.1 | 8.02 | 10005 |
| AT1G35160.1 | GF14 protein phi chain | 265.77 | 77 | 22 | 289 | 10 | 267 | 30.2 | 4.87 | 7634 |
| AT1G55020.1 | lipoxygenase 1 | 536.744 | 42 | 34 | 279 | 33 | 859 | 98 | 5.52 | 11240 |
| AT2G24940.1 | membrane-associated progesterone binding protein 2 | 296.128 | 93 | 11 | 271 | 11 | 100 | 11 | 4.88 | 14000 |
| AT1G78300.1 | general regulatory factor 2 | 238.277 | 71 | 20 | 268 | 11 | 259 | 29.1 | 4.79 | 7478 |
| AT3G02520.1 | general regulatory factor 7 | 240.997 | 67 | 18 | 266 | 5 | 265 | 29.8 | 4.82 | 7056 |
| AT1G58270.1 | TRAF-like family protein | 216.653 | 44 | 19 | 265 | 19 | 396 | 45 | 5.82 | 11821 |
| AT5G65430.3 | general regulatory factor 8 | 213.598 | 60 | 18 | 264 | 8 | 260 | 29.5 | 5.39 | 7494 |
| AT2G22170.1 | Lipase/lipooxygenase, PLAT/LH2 family protein | 133.119 | 44 | 7 | 264 | 6 | 183 | 20.1 | 5.31 | 11514 |
| AT5G19510.1 | Translation elongation factor EF1B/ribosomal protein S6 family prote | 257.189 | 65 | 11 | 259 | 8 | 224 | 24.2 | 4.56 | 9513 |
| AT1G80240.1 | Protein of unknown function, DUF642 | 291.843 | 52 | 19 | 253 | 19 | 370 | 40.2 | 8.37 | 9826 |
| AT1G45201.1 | triacylglycerol lipase-like 1 | 201.581 | 39 | 18 | 249 | 18 | 479 | 54.7 | 8.98 | 9954 |
| AT4G34670.1 | Ribosomal protein S3Ae | 249.798 | 68 | 20 | 230 | 11 | 262 | 29.8 | 9.74 | 7851 |
| AT2G37620.1 | actin 1 | 314.41 | 57 | 21 | 226 | 6 | 377 | 41.8 | 5.49 | 7033 |
| AT5G12940.1 | Leucine-rich repeat (LRR) family protein | 192.203 | 52 | 19 | 225 | 19 | 371 | 39.9 | 9.41 | 7955 |
| AT4G08780.1 | Peroxidase superfamily protein | 288.768 | 58 | 22 | 223 | 13 | 346 | 38.1 | 7.64 | 8131 |
| AT5G58420.1 | Ribosomal protein S4 (RPS4A) family protein | 176.193 | 58 | 17 | 220 | 1 | 262 | 29.8 | 10.21 | 7264 |
| AT5G07090.1 | Ribosomal protein S4 (RPS4A) family protein | 185.728 | 58 | 17 | 220 | 1 | 262 | 29.9 | 10.17 | 7318 |
| AT3G04840.1 | Ribosomal protein S3Ae | 201.771 | 57 | 16 | 216 | 7 | 262 | 29.8 | 9.76 | 7574 |
| AT5G10450.4 | G-box regulating factor 6 | 184.013 | 51 | 15 | 212 | 6 | 273 | 31 | 5.33 | 5402 |
| AT1G47128.1 | Granulin repeat cysteine protease family protein | 181.804 | 25 | 14 | 204 | 13 | 462 | 50.9 | 5.41 | 5823 |
| AT1G70840.1 | MLP-like protein 31 | 87.428 | 42 | 9 | 198 | 2 | 171 | 19.1 | 6.37 | 4756 |
| AT3G04400.2 | Ribosomal protein L14p/L23e family protein | 278.765 | 78 | 14 | 196 | 14 | 125 | 13.4 | 10.05 | 7126 |
| AT3G06860.1 | multifunctional protein 2 | 360.831 | 51 | 36 | 191 | 36 | 725 | 78.8 | 9.17 | 5740 |
| AT2G26080.1 | glycine decarboxylase P-protein 2 | 357.938 | 35 | 31 | 189 | 15 | 1044 | 113.7 | 6.65 | 6183 |
| AT2G02930.1 | glutathione S-transferase F3 | 106.331 | 39 | 9 | 187 | 1 | 212 | 24.1 | 6.74 | 6095 |

| | | | | | | | | | | |
|---|---|---|---|---|---|---|---|---|---|---|
| AT5G20010.1 | RAS-related nuclear protein-1 | 139.075 | 48 | 10 | 184 | 10 | 221 | 25.3 | 6.86 | 5470 |
| AT2G47510.1 | fumarase 1 | 331.204 | 49 | 21 | 179 | 10 | 492 | 53 | 7.88 | 7446 |
| AT4G33010.1 | glycine decarboxylase P-protein 1 | 340.729 | 34 | 27 | 178 | 11 | 1037 | 112.9 | 6.98 | 5901 |
| AT3G19390.1 | Granulin repeat cysteine protease family protein | 312.325 | 54 | 16 | 177 | 16 | 452 | 49.3 | 5.74 | 8452 |
| AT2G43610.1 | Chitinase family protein | 264.551 | 54 | 15 | 164 | 14 | 281 | 30 | 9.39 | 6372 |
| AT2G29560.1 | cytosolic enolase | 192.472 | 45 | 16 | 107 | 15 | 475 | 51.6 | 5.52 | 3660 |
| AT2G43535.1 | Scorpion toxin-like knottin superfamily protein | 16.655 | 31 | 3 | 107 | 3 | 97 | 10.7 | 5.38 | 2122 |
| AT5G20080.1 | FAD/NAD(P)-binding oxidoreductase | 198.945 | 61 | 16 | 103 | 16 | 328 | 36 | 8.69 | 3438 |
| AT4G26010.1 | Peroxidase superfamily protein | 141.001 | 43 | 17 | 103 | 17 | 310 | 33.8 | 9.94 | 3648 |
| AT5G52240.1 | membrane steroid binding protein 1 | 123.649 | 51 | 7 | 102 | 5 | 220 | 24.4 | 4.74 | 4903 |
| AT5G14590.1 | Isocitrate/isopropylmalate dehydrogenase family protein | 206.667 | 52 | 23 | 99 | 19 | 485 | 54.2 | 7.97 | 4260 |
| AT3G48890.1 | membrane-associated progesterone binding protein 3 | 106.645 | 45 | 7 | 98 | 5 | 233 | 25.4 | 4.63 | 5277 |
| AT5G50950.1 | FUMARASE 2 | 190.319 | 28 | 13 | 97 | 2 | 510 | 55.7 | 7.36 | 4604 |
| AT5G27770.1 | Ribosomal L22e protein family | 70.684 | 47 | 9 | 84 | 3 | 124 | 14 | 9.55 | 2096 |
| AT3G44110.1 | DNAJ homologue 3 | 211.202 | 45 | 13 | 84 | 7 | 420 | 46.4 | 6.11 | 2707 |
| AT3G05560.1 | Ribosomal L22e protein family | 69.472 | 47 | 9 | 75 | 3 | 124 | 14 | 9.55 | 1745 |
| AT1G54340.1 | isocitrate dehydrogenase | 111.54 | 43 | 15 | 73 | 9 | 416 | 47.2 | 7.71 | 2445 |
| AT2G41475.1 | Embryo-specific protein 3, (ATS3) | 23.449 | 25 | 5 | 72 | 5 | 179 | 19.9 | 8.82 | 1627 |
| AT4G02270.1 | root hair specific 13 | 79.958 | 52 | 9 | 37 | 9 | 165 | 18.1 | 9.11 | 1693 |
| AT2G47540.1 | Pollen Ole e 1 allergen and extensin family protein | 78.553 | 48 | 6 | 25 | 6 | 173 | 19 | 5.66 | 1230 |
| AT5G39570.1 | FUNCTIONS IN: molecular_function unknown; INVOLVED IN: biologica | 78.72 | 47 | 10 | 24 | 10 | 381 | 43.5 | 4.78 | 862 |
| AT1G75750.1 | GAST1 protein homolog 1 | 15.139 | 33 | 3 | 16 | 3 | 98 | 10.7 | 9.11 | 528 |

| Accession | Description | Sum PEP Score | Coverag e [%] | # Peptides | # PSMs | # Unique Peptides | # AAs | MW [kDa] | calc. pI | Score Mascot |
|---|---|---|---|---|---|---|---|---|---|---|
| AT3G04120.1 | glyceraldehyde-3-phosphate dehydrogenase C subui | 371.849 | 84 | 33 | 981 | 4 | 338 | 36.9 | 7.12 | 39930 |
| AT1G79550.1 | phosphoglycerate kinase | 410.954 | 78 | 30 | 451 | 25 | 401 | 42.1 | 5.68 | 17577 |
| AT4G30170.1 | Peroxidase family protein | 256.761 | 78 | 22 | 200 | 16 | 325 | 35.8 | 9.1 | 8245 |
| AT1G78850.1 | D-mannose binding lectin protein with Apple-like ca | 201.483 | 51 | 20 | 196 | 8 | 441 | 49 | 7.65 | 9143 |
| AT3G01190.1 | Peroxidase superfamily protein | 236.27 | 73 | 23 | 193 | 21 | 321 | 34.9 | 8.95 | 6370 |
| AT1G68560.1 | alpha-xylosidase 1 | 282.377 | 44 | 31 | 176 | 30 | 915 | 102.3 | 6.77 | 5208 |
| AT1G28290.1 | arabinogalactan protein 31 | 111.678 | 23 | 7 | 155 | 7 | 359 | 38.5 | 10.17 | 5823 |
| AT2G37130.1 | Peroxidase superfamily protein | 174.167 | 50 | 15 | 120 | 15 | 327 | 36.7 | 7.4 | 3919 |
| AT3G13790.1 | Glycosyl hydrolases family 32 protein | 221.88 | 48 | 28 | 118 | 25 | 584 | 66.2 | 9.07 | 3911 |
| AT1G05240.1 | Peroxidase superfamily protein | 165.468 | 59 | 16 | 113 | 15 | 325 | 35.6 | 9.25 | 3981 |
| AT5G17820.1 | Peroxidase superfamily protein | 166.886 | 61 | 14 | 107 | 14 | 313 | 34.1 | 10.05 | 3599 |
| AT4G20830.1 | FAD-binding Berberine family protein | 185.783 | 42 | 25 | 107 | 23 | 570 | 63.5 | 9.58 | 3653 |
| AT1G78860.1 | D-mannose binding lectin protein with Apple-like ca | 134.867 | 46 | 18 | 93 | 6 | 443 | 49.2 | 6.28 | 3182 |
| AT4G20860.1 | FAD-binding Berberine family protein | 167.07 | 43 | 25 | 91 | 25 | 530 | 60.4 | 9.04 | 2819 |
| AT4G19410.2 | Pectinacetylesterase family protein | 150.492 | 52 | 18 | 91 | 2 | 517 | 55.8 | 9.04 | 3065 |
| AT5G44380.1 | FAD-binding Berberine family protein | 111.359 | 34 | 20 | 90 | 17 | 541 | 60.6 | 9.48 | 3016 |
| AT4G19410.1 | Pectinacetylesterase family protein | 144.814 | 66 | 17 | 89 | 1 | 391 | 42.1 | 9.11 | 3004 |
| AT1G80240.1 | Protein of unknown function, DUF642 | 151.119 | 56 | 17 | 78 | 17 | 370 | 40.2 | 8.37 | 2572 |
| AT5G12940.1 | Leucine-rich repeat (LRR) family protein | 125.986 | 55 | 16 | 73 | 16 | 371 | 39.9 | 9.41 | 2138 |
| AT4G30280.1 | xyloglucan endotransglucosylase/hydrolase 18 | 96.468 | 50 | 12 | 64 | 5 | 282 | 32.1 | 8.56 | 2351 |
| AT1G30730.1 | FAD-binding Berberine family protein | 119.106 | 34 | 19 | 63 | 13 | 526 | 58.9 | 9.17 | 1903 |
| AT4G30270.1 | xyloglucan endotransglucosylase/hydrolase 24 | 101.435 | 39 | 8 | 59 | 7 | 269 | 30.7 | 8.31 | 2335 |
| AT5G64260.1 | EXORDIUM like 2 | 71.091 | 28 | 9 | 58 | 7 | 305 | 32.7 | 9.38 | 1528 |
| AT4G30290.1 | xyloglucan endotransglucosylase/hydrolase 19 | 76.089 | 40 | 10 | 58 | 5 | 277 | 31.5 | 8.78 | 2056 |
| AT2G45220.1 | Plant invertase/pectin methylesterase inhibitor supe | 127.257 | 40 | 16 | 58 | 15 | 511 | 55.9 | 9.04 | 2463 |
| AT1G65310.1 | xyloglucan endotransglucosylase/hydrolase 17 | 72.178 | 49 | 12 | 56 | 5 | 282 | 32 | 8.56 | 1929 |
| AT5G56870.1 | beta-galactosidase 4 | 145.503 | 40 | 22 | 49 | 17 | 724 | 80.5 | 8.68 | 1547 |
| AT1G70840.1 | MLP-like protein 31 | 41.574 | 40 | 8 | 43 | 2 | 171 | 19.1 | 6.37 | 971 |
| AT1G12140.1 | flavin-monooxygenase glucosinolate S-oxygenase 5 | 127.37 | 41 | 16 | 42 | 16 | 459 | 52.1 | 6.19 | 1220 |
| AT1G45130.1 | beta-galactosidase 5 | 114.369 | 29 | 17 | 38 | 16 | 732 | 81.4 | 8.25 | 1311 |
| AT3G49960.1 | Peroxidase superfamily protein | 108.359 | 50 | 13 | 37 | 11 | 329 | 35.7 | 9.06 | 1422 |
| AT2G37170.1 | plasma membrane intrinsic protein 2 | 33.523 | 18 | 5 | 34 | 4 | 285 | 30.4 | 7.81 | 1028 |
| AT5G09440.1 | EXORDIUM like 4 | 52.822 | 32 | 9 | 34 | 7 | 278 | 29.5 | 9.48 | 983 |
| AT1G30870.1 | Peroxidase superfamily protein | 68.433 | 36 | 12 | 32 | 12 | 349 | 38.3 | 6.89 | 955 |
| AT4G25260.1 | Plant invertase/pectin methylesterase inhibitor supe | 74.366 | 54 | 8 | 25 | 8 | 201 | 22.2 | 5.1 | 1093 |
| AT3G43670.1 | Copper amine oxidase family protein | 84.669 | 28 | 13 | 21 | 11 | 687 | 77.5 | 6.04 | 711 |
| AT2G42570.1 | TRICHOME BIREFRINGENCE-LIKE 39 | 37.665 | 24 | 8 | 20 | 8 | 367 | 43.1 | 9.44 | 452 |
| AT3G09925.1 | Pollen Ole e 1 allergen and extensin family protein | 38.437 | 39 | 6 | 19 | 6 | 171 | 19.2 | 7.64 | 561 |
| AT5G05500.1 | Pollen Ole e 1 allergen and extensin family protein | 45.126 | 34 | 4 | 19 | 4 | 183 | 20.2 | 8.24 | 674 |

| | | | | | | | | | | |
|---|---|---|---|---|---|---|---|---|---|---|
| AT2G33790.1 | arabinogalactan protein 30 | 45.505 | 18 | 4 | 16 | 4 | 239 | 25.7 | 9.95 | 527 |
| AT5G51490.1 | Plant invertase/pectin methylesterase inhibitor supe | 50.948 | 21 | 8 | 11 | 6 | 536 | 58.8 | 9.48 | 450 |
| AT3G47400.1 | Plant invertase/pectin methylesterase inhibitor supe | 35.517 | 13 | 6 | 8 | 5 | 594 | 65.4 | 9.82 | 220 |
| AT5G51500.1 | Plant invertase/pectin methylesterase inhibitor supe | 15.716 | 10 | 5 | 5 | 3 | 540 | 59.6 | 9.6 | 131 |
| AT5G53370.1 | pectin methylesterase PCR fragment F | 9.842 | 4 | 3 | 3 | 2 | 587 | 64.2 | 7.64 | 108 |

| Accession | Description | Sum PEP Score | Coverage [%] | # Peptides | # PSMs | # Unique Peptides | # AAs | MW [kDa] | calc. pI | Score Mascot |
|---|---|---|---|---|---|---|---|---|---|---|
| AT3G04120.1 | glyceraldehyde-3-phosphate dehydrogenase C subunit 1 | 465.82 | 86 | 32 | 1546 | 4 | 338 | 36.9 | 7.12 | 65217 |
| AT3G01190.1 | Peroxidase superfamily protein | 396.49 | 74 | 25 | 339 | 23 | 321 | 34.9 | 8.95 | 12263 |
| AT1G78850.1 | D-mannose binding lectin protein with Apple-like carbohydrate-binding dor | 259.48 | 54 | 21 | 210 | 9 | 441 | 49 | 7.65 | 9551 |
| AT1G05240.1 | Peroxidase superfamily protein | 214.08 | 57 | 16 | 204 | 15 | 325 | 35.6 | 9.25 | 8181 |
| AT2G37130.1 | Peroxidase superfamily protein | 236.35 | 54 | 19 | 176 | 19 | 327 | 36.7 | 7.4 | 6258 |
| AT4G19410.2 | Pectinacetylesterase family protein | 222.46 | 50 | 17 | 146 | 2 | 517 | 55.8 | 9.04 | 5774 |
| AT4G19410.1 | Pectinacetylesterase family protein | 215.13 | 63 | 16 | 145 | 1 | 391 | 42.1 | 9.11 | 5734 |
| AT4G20830.1 | FAD-binding Berberine family protein | 235.7 | 42 | 25 | 136 | 23 | 570 | 63.5 | 9.58 | 4850 |
| AT4G30280.1 | xyloglucan endotransglucosylase/hydrolase 18 | 121.22 | 46 | 12 | 85 | 6 | 282 | 32.1 | 8.56 | 3358 |
| AT1G65310.1 | xyloglucan endotransglucosylase/hydrolase 17 | 102.51 | 45 | 12 | 78 | 6 | 282 | 32 | 8.56 | 2939 |
| AT4G30290.1 | xyloglucan endotransglucosylase/hydrolase 19 | 82.421 | 35 | 9 | 74 | 4 | 277 | 31.5 | 8.78 | 2933 |
| AT1G45130.1 | beta-galactosidase 5 | 212.39 | 41 | 23 | 63 | 23 | 732 | 81.4 | 8.25 | 2207 |
| AT1G13950.1 | eukaryotic elongation factor 5A-1 | 126.38 | 73 | 11 | 52 | 7 | 158 | 17.4 | 5.76 | 1684 |
| AT3G46280.1 | protein kinase-related | 123.73 | 30 | 11 | 49 | 10 | 471 | 50.5 | 5.15 | 1674 |
| AT3G62680.1 | proline-rich protein 3 | 87.886 | 31 | 9 | 39 | 9 | 313 | 34.4 | 9.41 | 1517 |
| AT4G02270.1 | root hair specific 13 | 32.531 | 39 | 5 | 34 | 5 | 165 | 18.1 | 9.11 | 1127 |
| AT4G30270.1 | xyloglucan endotransglucosylase/hydrolase 24 | 77.918 | 34 | 7 | 32 | 6 | 269 | 30.7 | 8.31 | 1487 |
| AT3G09925.1 | Pollen Ole e 1 allergen and extensin family protein | 63.285 | 54 | 8 | 31 | 8 | 171 | 19.2 | 7.64 | 1002 |
| AT5G05500.1 | Pollen Ole e 1 allergen and extensin family protein | 92.716 | 45 | 6 | 30 | 6 | 183 | 20.2 | 8.24 | 1317 |
| AT5G48070.1 | xyloglucan endotransglucosylase/hydrolase 20 | 64.213 | 41 | 9 | 27 | 6 | 282 | 32.4 | 8.56 | 1406 |
| AT4G26220.1 | S-adenosyl-L-methionine-dependent methyltransferases superfamily protei | 57.765 | 50 | 9 | 20 | 9 | 232 | 25.9 | 5.53 | 549 |
| AT2G18800.1 | xyloglucan endotransglucosylase/hydrolase 21 | 29.355 | 15 | 4 | 17 | 3 | 305 | 34.1 | 8.1 | 929 |
| AT3G46270.1 | receptor protein kinase-related | 47.55 | 24 | 7 | 13 | 6 | 470 | 50.4 | 5.16 | 630 |
| AT5G62130.2 | Per1-like family protein | 15.452 | 14 | 3 | 3 | 3 | 345 | 39.9 | 7.9 | 123 |

| Accession | Description | Sum PEP Score | Coverage [%] | # Peptides | # PSMs | # Unique Peptides | # AAs | MW [kDa] | calc. pI | Score Mascot |
|---|---|---|---|---|---|---|---|---|---|---|
| AT3G04120.1 | glyceraldehyde-3-phosphate dehydrogenase C subunit 1 | 340.304 | 83 | 29 | 1114 | 4 | 338 | 36.9 | 7.12 | 46360 |
| AT4G30170.1 | Peroxidase family protein | 230.362 | 78 | 21 | 190 | 16 | 325 | 35.8 | 9.1 | 7658 |
| AT3G01190.1 | Peroxidase superfamily protein | 278.793 | 73 | 22 | 188 | 20 | 321 | 34.9 | 8.95 | 6820 |
| AT1G05240.1 | Peroxidase superfamily protein | 180.791 | 56 | 16 | 170 | 15 | 325 | 35.6 | 9.25 | 6623 |
| AT3G13790.1 | Glycosyl hydrolases family 32 protein | 226.874 | 47 | 27 | 154 | 25 | 584 | 66.2 | 9.07 | 5210 |
| AT2G37130.1 | Peroxidase superfamily protein | 148.783 | 50 | 14 | 106 | 14 | 327 | 36.7 | 7.4 | 3728 |
| AT5G44380.1 | FAD-binding Berberine family protein | 130.679 | 36 | 21 | 96 | 17 | 541 | 60.6 | 9.48 | 3225 |
| AT4G19410.1 | Pectinacetylesterase family protein | 138.923 | 52 | 13 | 80 | 1 | 391 | 42.1 | 9.11 | 2419 |
| AT4G19410.2 | Pectinacetylesterase family protein | 137.73 | 38 | 13 | 80 | 1 | 517 | 55.8 | 9.04 | 2421 |
| AT1G45130.1 | beta-galactosidase 5 | 140.715 | 36 | 21 | 44 | 21 | 732 | 81.4 | 8.25 | 1397 |
| AT1G12140.1 | flavin-monooxygenase glucosinolate S-oxygenase 5 | 135.816 | 44 | 16 | 42 | 16 | 459 | 52.1 | 6.19 | 1141 |
| AT1G55120.1 | beta-fructofuranosidase 5 | 69.549 | 21 | 10 | 36 | 8 | 594 | 67 | 5.78 | 1011 |
| AT1G70840.1 | MLP-like protein 31 | 30.531 | 40 | 7 | 29 | 2 | 171 | 19.1 | 6.37 | 720 |
| AT4G02270.1 | root hair specific 13 | 29.507 | 39 | 5 | 26 | 5 | 165 | 18.1 | 9.11 | 750 |
| AT1G48930.1 | glycosyl hydrolase 9C1 | 84.023 | 33 | 15 | 25 | 14 | 627 | 69 | 8.91 | 860 |
| AT3G43670.1 | Copper amine oxidase family protein | 68.704 | 24 | 12 | 19 | 10 | 687 | 77.5 | 6.04 | 599 |
| AT3G62680.1 | proline-rich protein 3 | 45.397 | 26 | 7 | 17 | 7 | 313 | 34.4 | 9.41 | 690 |
| AT2G17760.1 | Eukaryotic aspartyl protease family protein | 39.934 | 15 | 6 | 15 | 6 | 513 | 56 | 5.27 | 396 |
| AT2G14835.1 | RING/U-box superfamily protein | 41.071 | 24 | 5 | 13 | 5 | 343 | 37.4 | 8.1 | 442 |
| AT5G05500.1 | Pollen Ole e 1 allergen and extensin family protein | 45.788 | 35 | 4 | 12 | 4 | 183 | 20.2 | 8.24 | 406 |

| Accession | Description | Sum PEP Score | Coverage [%] | # Peptides | # PSMs | # Unique Peptides | # AAs | MW [kDa] | calc. pI | Score Mascot |
|---|---|---|---|---|---|---|---|---|---|---|
| AT2G19760.1 | profilin 1 | 132.218 | 55 | 7 | 80 | 7 | 131 | 14.3 | 4.82 | 2938 |
| AT2G24180.1 | cytochrome p450 71b6 | 59.113 | 28 | 11 | 26 | 11 | 503 | 57 | 7.85 | 597 |
