## Supplementary material for "Evidences for a nutritional role of iodine in plants": Table S11

| Master Protein Accession | Description | Iodinated Sequence | Iodinated site | DATASET |
|---|---|---|---|---|
| ATCG00120.1 | ATP synthase subunit alpha | EAYPGDVFYLHSR | [Y3] | ChlorBN |
| | | SVYEPLQTGLIAIDSMIPIGR | [Y3] | ChlorBN |
| ATCG00480.1 | ATP synthase subunit beta | GIYPAVDPLDSTSTMLQPR | [Y3] | ChlorBN |
| | | GSITSIQAVYVPADDLTDPAPATTFAHLDATTVLSR | [Y10] | ChlorBN |
| | | IVGEEHYETAQQVK | [Y7] | ChlorBN, Chlo_3516 |
| | | VALVYGQMNEPPGAR | [Y5] | ChlorBN |
| | | VGLTALTMAEYFR | [Y11] | ChlorBN |
| AT4G04640.1 | ATPase, F1 complex, gamma subunit protein | GLGLEYTVISVGK | [Y6] | ChlorBN |
| AT1G29910.1 | chlorophyll A/B binding protein 3 | YLGPFSGESPSYLTGEFPGDYGWDTAGLSADPETFAR | [Y5]/[Y12]/[Y21] | ChlorBN |
| AT1G20340.1 | Cupredoxin superfamily protein | NNAGYPHNVVFDEDEIPSGVDVAK | [Y5]/[H7] | CAU, Ros |
| AT5G66190.1 | ferredoxin-NADP(+)-oxidoreductase 1 | LVYTNDGGEIVK | [Y3] | ChlorBN |
| AT4G38970.1 | fructose-bisphosphate aldolase 2 | ATPEQVAAYTLK | [Y9] | ChlorBN |
| AT5G42270.1 | FtsH extracellular protease family | DYSMATADVVDAEVR | [Y2] | ChlorBN |
| AT3G09260.1 | Glycosyl hydrolase superfamily protein | CSSYVNAK | [Y4] | ChlorBN |
| | | GPALWDIYCR | [Y8] | ChlorBN |
| | | FGLYYVDFK | [Y4]/[Y5] | ChlorBN |
| AT4G10340.1 | light harvesting complex of photosystem II 5 | SEIPEYLNGEVAGDYGYDPFGLGK | [Y6]/[Y15]/Y17] | ChlorBN |
| | | TGALLLDGNTLNYFGK | [Y13] | ChlorBN |
| AT5G54270.1 | light-harvesting chlorophyll B-binding protein 3 | YLGPFSVQTPSYLTGEFPGDYGWDTAGLSADPEAFAK | [Y12]/[Y21] | ChlorBN |
| AT2G34430.1 | light-harvesting chlorophyll-protein complex II subunit B1 | YLGPFSGEPPSYLTGEFPGDYGWDTAGLSADPETFAR | [Y12]/[Y21] | ChlorBN, PXD010730, LFD |
| AT2G24940.1 | membrane-associated progesterone binding protein 2 | SFYGSGGDYSMFAGK | [Y3] | Ros |
| AT4G22890.5 | PGR5-LIKE A | FLEASMAYVSGNPILNDEEYDKLK | [Y20] | ChlorBN |
| ATCG00540.1 | photosynthetic electron transfer A | GGYEITIVDASNGR | [Y3] | ChlorBN |
| AT4G03280.1 | photosynthetic electron transfer C | GDPTYLVVENDK | [Y5] | ChlorBN |
| AT4G28750.1 | Photosystem I reaction centre subunit IV / PsaE protein | VNYANISTNNYALDEVEEVAA | [Y3] | ChlorBN |
| AT2G20260.1 | photosystem I subunit E-2 | VNYANISTNNYALDEVEEVK | [Y3] | ChlorBN |
| AT1G31330.1 | photosystem I subunit F | LYAPESAPALALNAQIEK | [Y2] | ChlorBN, Ros |
| AT1G52230.1 | photosystem I subunit H2 | SVYFDLEDLGNTTGQWDVYGSDAPSPYNPLQSK | [Y19] | ChlorBN |
| ATCG00340.1 | Photosystem I, PsaA/PsaB protein | TSYGFDVLLSSTSGPAFNAGR | [Y3] | ChlorBN |
| AT1G03600.1 | photosystem II family protein | DIYSALNAVSGHYVSFGPTAPIPAK | [Y3] | ChlorBN |
| AT1G06680.1 | photosystem II subunit P-1 | SITDYGSPEEFLSQVNYLLGK | [Y5/Y17] | ChlorBN, PXD010730 |
| AT1G79040.1 | photosystem II subunit R | YGANVDGYSPIYNENEWSASGDVYK | [Y1]/[Y8]/[Y12]/[Y24] | ChlorBN |

| | | | | |
|---|---|---|---|---|
| AT2G05100.1 | photosystem II light harvesting complex gene 2.1 | STPQSIWYGPDRPK | [Y8] | ChlorBN |
| | | YLGPFSENTPSYLTGEYPGDYGWDTAGLSADPETFAK | [Y1//Y12/Y17/Y21] | ChlorBN |
| AT2G20890.1 | photosystem II reaction center PSB29 protein | AYIEALNEDPK | [Y2] | ChlorBN |
| AT2G30790.1 | photosystem II subunit P-2 | SITDYGSPEQFLSQVNYLLGK | [Y5]/[Y17] | ChlorBN |
| AT3G50820.1 | photosystem II subunit O-2 | GGSTGYDNAVALPAGGR | [Y6] | ChlorBN, Chlo_3516 |
| AT4G05180.1 | photosystem II subunit Q-2 | YYSETVSSLNNVLAK | [Y2] | ChlorBN |
| AT4G21280.1 | photosystem II subunit QA | LFDTIDNLDYAAK | [Y10] | ChlorBN |
| | | YYAETVSALNEVLAK | [Y1]/[Y2] | ChlorBN |
| ATCG00020.1 | photosystem II reaction center protein A | ETTENESANEGYR | [Y12] | ChlorBN, TMAR |
| | | FGQEEETYNIVAAHGYFGR | [Y8] | ChlorBN |
| ATCG00270.1 | photosystem II reaction center protein D | AAEDPEFETFYTK | [Y3] | ChlorBN |
| ATCG00280.1 | photosystem II reaction center protein C | DIQPWQERRSAEYMTHAPLGSLNSVGGVATEINAVNYVSPR | [H16] | LFD |
| | | RSAEYMTHAPLGSLNSVGGVATEINAVNYVSPR | [H8] | CAU, Ros |
| | | SAEYMTHAPLGSLNSVGGVATEINAVNYVSPR | [H7]/[Y28] | ChlorBN, LFP |
| ATCG00680.1 | photosystem II reaction center protein B | LAFYDYIGNNPAK | [Y4]/[Y6] | ChlorBN, TMAR, CAU, Ros |
| | | YQWDQGYFQQEIYR | [Y1]/[Y7] | ChlorBN |
| AT5G07020.1 | proline-rich family protein | AVDYSGPSLSYYINK | [Y12] | ChlorBN |
| AT1G74470.1 | Pyridine nucleotide-disulphide oxidoreductase family protein | SIDAGDYDYAIAFQER | [Y7] | ChlorBN |
| ATMG00280.1 | Ribulose bisphosphate carboxylase large chain, catalytic domain | GGLYFTKDDENVNSQPFMR | [Y4] | CLLF, LFD, LFP, LFPT |
| AT1G67090.1 | ribulose bisphosphate carboxylase small chain 1A | EYPNAFIR | [Y2] | ChlorBN, CAU, Ros |
| ATCG00490.1 | ribulose-bisphosphate carboxylases | GHYLNATAGTCEEMIKR | [H2]/[Y3] | ChlorBN, Chlo_10545, CAU |
| | | LTYYTPEYETKDTDILAAFR | [Y3]/[Y4]/[Y8] | CAU, Ros |
| AT1G71500.1 | Rieske (2Fe-2S) domain-containing protein | SPAEGAYSEGLLNAR | [Y7] | ChlorBN |
| AT2G39730.1 | rubisco activase | GLAYDTSDDQQDITR | [Y4] | ChlorBN |
| AT1G54780.1 | thylakoid lumen 18.3 kDa protein | ADAFEYADQVLEK | [Y6] | ChlorBN |
| | | ETYVVDDAGVLSR | [Y3] | ChlorBN |
| AT5G09810.1 | actin 7 | NYELPDGQVITIGAER | [Y2] | Root |
| AT1G28290.1 | arabinogalactan protein 31 | NGYFLLLAPK | [Y3] | RT |
| AT5G08680.1 | ATP synthase alpha/beta family protein | VGLTGLTVAEYFR | [Y11] | Root |
| AT1G45130.1 | beta-galactosidase 5 | YDEDIATYGNR | [Y1] | RT, RTTP, RTUZ |
| AT2G43610.1 | Chitinase family protein | YCSPSTTYPCQPGK | [Y1] | Root |
| AT3G43670.1 | Copper amine oxidase family protein | GTAYENVEDLGEK | [Y4] | RT, RTUZ |
| AT1G78850.1 | D-mannose binding lectin protein | TGDSSLVAYVK | [Y9] | Root, RT, RTTP |
| AT5G20080.1 | FAD/NAD(P)-binding oxidoreductase | IFYTVDNPTK | [Y3] | Root |
| AT4G20830.1 | FAD-binding Berberine family protein | DVDIGVNDHGANSYK | [Y14] | RTTP |

| | | | | |
|---|---|---|---|---|
| AT5G44380.1 | FAD-binding Berberine family protein | YGLAGDNVLDVK | [Y1] | RTUZ |
| AT5G50950.1 | FUMARASE 2 | IGYDNAAAVAK | [Y3] | Root |
| AT3G04120.1 | glyceraldehyde-3-phosphate dehydrogenase C sub 1 | LKGILGYTEDDVVSTDFVGDNR | [Y7] | RT, RTTP, RTUZ |
| AT4G16260.1 | Glycosyl hydrolase superfamily protein | AFYTNLASR<br>LYDPNQAALNALR | [Y3]<br>[Y2] | Root<br>Root |
| AT3G13790.1 | Glycosyl hydrolases family 32 protein | HDYYTIGTYDR | [Y3]/[Y4]/[Y9] | RT, RTUZ |
| AT3G19390.1 | Granulin repeat cysteine protease family protein | VVTIDGYEDVPQNDEK | [Y7] | Root |
| AT3G12580.1 | heat shock protein 70 | NALENYAYNMR<br>TTPSYVAFTDSER | [Y6]<br>[Y5] | Root<br>Root |
| AT5G42020.1 | Heat shock protein 70 (Hsp 70) family protein | NALETYVYNMK | [Y6]/[Y8] | Root |
| AT3G16430.1 | jacalin-related lectin 31 | VYVGQAQDGISAVK | [Y2] | Root |
| AT2G22170.1 | Lipase/lipooxygenase, PLAT/LH2 family protein | VYDKYGDYIGIR | [Y5]/[Y8] | Root |
| AT3G16460.1 | Mannose-binding lectin superfamily protein | IYASYGGEGIQYVK | [Y5]/[Y12] | Root |
| AT3G48890.1 | membrane-associated progesterone binding protein 3 | MFYGPGGPYALFAGK | [Y3] | Root |
| AT4G19410.1 | Pectinacetylesterase family protein | DITGGSYIQSYYSK | [Y7] | RT |
| AT4G30170.1 | Peroxidase family protein | EVVVLTGGPSYPVELGR<br>IYNFSPTTR<br>TGFYQNSCPNVETIVR | [Y11]<br>[Y2]<br>[Y4] | Root, RT, RTUZ<br>Root<br>Root, RT |
| AT1G05240.1 | Peroxidase superfamily protein | GDSDPSMNPSYVR | [Y11] | RT, RTTP, RTUZ |
| AT2G37130.1 | Peroxidase superfamily protein | CPSPTPDPNAVLYSR<br>QQVETLYYK | [Y13]<br>[Y8] | RT, RTUZ<br>RT, RTTP, RTUZ |
| AT3G01190.1 | Peroxidase superfamily protein | TFDLSYFTLVAK | [Y6] | RT, RTUZ |
| AT5G17820.1 | Peroxidase superfamily protein | DSVALAGGPSYSIPTGR<br>VGFYSQSCPQAETIVR | [Y11]<br>[Y4] | Root, RT<br>RT |
| AT1G79550.1 | phosphoglycerate kinase | YSLKPLVPRLSELLGVEVVMANDSIGEEVQK | [Y1] | RT |
| AT4G20260.4 | plasma-membrane associated cation-binding protein 1 | VVETYEATSAEVK | [Y5] | Root |
| AT2G47540.1 | Pollen Ole e 1 allergen and extensin family protein | GISGAILQNYR | [Y10] | Root |
| AT5G05500.1 | Pollen Ole e 1 allergen and extensin family protein | TDSYGHFYGELK | [Y4] | RT, RTTP, RTUZ |
| AT4G14060.1 | Polyketide cyclase/dehydrase and lipid transport superfamily protein | VYDTILQFIQK<br>ATSGTYVTEVPLKGSAEK | [Y2]<br>[Y6] | Root<br>Root |
| AT4G23680.1 | Polyketide cyclase/dehydrase and lipid transport superfamily protein | VYDVVYQFIPK | [Y2] | Root |
| AT2G19760.1 | profilin 1 | TNQALVFGFYDEPMTGGQCNLVVER | [Y10] | RTTOF |
| AT4G02270.1 | root hair specific 13 | VDAYGNELVPISILSSK | [Y4] | Root, RTTP, RTUZ |
| AT4G26220.1 | S-adenosyl-L-methionine-dependent methyltransferases | GLLKSEELYKYILETSVYPR | [Y9] | RTTP |
| AT1G58270.1 | TRAF-like family protein | FLDSYTSDSFSSGGR | [Y5] | Root |

| | | | | |
|---|---|---|---|---|
| AT1G45201.1 | triacylglycerol lipase-like 1 | FVYNNDVVPR | [Y3] | Root |
| AT3G55440.1 | triosephosphate isomerase | IIYGGSVNGGNCK | [Y3] | Root |
| AT4G30270.1 | xyloglucan endotransglucosylase/hydrolase 24 | NYESLGVLFPK | [Y2] | RT |
| | | LVPGNSAGTVTTFYLK | [Y14] | RTTP |
