## Supplementary material for "Evidences for a nutritional role of iodine in plants": Table S12

| *A thaliana* leaves | | |
|---|---|---|
| **Accession code** | **STRING symbol** | **Protein description** |
| ATCG00120 | ATPA | Encodes the ATPase alpha subunit, which is a subunit of ATP synthase |
| ATCG00480 | PB | ATP synthase subunit beta, chloroplastic |
| AT4G04640 | ATPC1 | ATP synthase gamma chain 1, chloroplastic |
| AT1G29910 | CAB3 | Chlorophyll A/B binding protein 3 |
| AT1G20340 | DRT112 | Plastocyanin major isoform, chloroplastic |
| AT5G66190 | FNR1 | Ferredoxin-NADP reductase, leaf isozyme 1, chloroplastic |
| AT4G38970 | FBA2 | Fructose-bisphosphate aldolase 2, chloroplastic |
| AT5G42270 | VAR1 | ATP-dependent zinc metalloprotease FTSH 5, chloroplastic |
| AT3G09260 | PYK10 | Glycosyl hydrolase superfamily protein |
| AT4G10340 | LHCB5 | Chlorophyll a-b binding protein CP26, chloroplastic |
| AT5G54270 | LHCB3 | Chlorophyll a-b binding protein 3, chloroplastic |
| AT2G34430 | LHB1B1 | Light-harvesting chlorophyll-protein complex II subunit B1 |
| AT2G24940 | MAPR2 | Membrane-associated progesterone binding protein 2 (MAPR2) |
| AT4G22890 | PGRL1A | Encodes PGRL1A, a transmembrane protein present in thylakoids |
| ATCG00540 | PETA | Photosynthetic electron transfer A |
| AT4G03280 | PETC | Cytochrome b6-f complex iron-sulfur subunit, chloroplastic |
| AT4G28750 | PSAE-1 | Photosystem I reaction center subunit IV A, chloroplastic |
| AT2G20260 | PSAE-2 | Photosystem I reaction center subunit IV B, chloroplastic |
| AT1G31330 | PSAF | Photosystem I reaction center subunit III, chloroplastic |
| AT1G52230 | PSAH2 | Photosystem I reaction center subunit VI-2, chloroplastic |
| ATCG00340 | PSAB | Photosystem I P700 chlorophyll a apoprotein A2 |
| AT1G03600 | PSB27 | Photosystem II repair protein PSB27-H1, chloroplastic |
| AT1G06680 | PSBP-1 | Oxygen-evolving enhancer protein 2-1, chloroplastic |
| AT1G79040 | PSBR | Encodes for the 10 kDa PsbR subunit of photosystem II (PSII) |
| AT2G05100 | LHCB2.1 | Chlorophyll a-b binding protein 2.1, chloroplastic |
| AT2G20890 | PSB29 | Protein THYLAKOID FORMATION 1, chloroplastic |
| AT2G30790 | PSBP-2 | Putative oxygen-evolving enhancer protein 2-2 |
| AT3G50820 | PSBO2 | Oxygen-evolving enhancer protein 1-2, chloroplastic |
| AT4G05180 | PSBQ-2 | Oxygen-evolving enhancer protein 3-2, chloroplastic |
| AT4G21280 | PSBQA | |
| ATCG00020 | PSBA | Photosystem II reaction center protein A |
| ATCG00270 | PSBD | Photosystem II reaction center protein D |
| ATCG00280 | PSBC | Photosystem II CP43 reaction center protein |

| | | |
|---|---|---|
| ATCG00680 | PSBB | Photosystem II CP47 reaction center protein |
| AT5G07020 | AT5G07020 | Protein MAINTENANCE OF PSII UNDER HIGH LIGHT 1 |
| AT1G74470 | AT1G74470 | Pyridine nucleotide-disulphide oxidoreductase family protein |
| ATMG00280 | ORF110A | Ribulose bisphosphate carboxylase large chain |
| AT1G67090 | RBCS1A | Ribulose bisphosphate carboxylase small chain 1A, chloroplastic |
| ATCG00490 | RBCL | Ribulose bisphosphate carboxylase large chain |
| AT1G71500 | AT1G71500 | Uncharacterized protein At1g71500/F26A9_12 |
| AT2G39730 | RCA | Ribulose bisphosphate carboxylase/oxygenase activase, chloroplastic |
| AT1G54780 | TLP18.3 | UPF0603 protein At1g54780, chloroplastic |

| ***A thaliana roots*** | | |
|---|---|---|
| **Accession code** | **STRING symbol** | **Protein description** |
| AT5G09810 | ACT7 | Actin-7 |
| AT1G28290 | AGP31 | Atypical plasma membrange arabinogalactan protein |
| AT5G08680 | AT5G08680 | ATP synthase subunit beta-3, mitochondrial |
| AT1G45130 | BGAL5 | Beta-galactosidase 5 (BGAL5) |
| AT2G43610 | AT2G43610 | Chitinase family protein |
| AT3G43670 | AT3G43670 | Copper amine oxidase family protein |
| AT1G78850 | AT1G78850 | D-mannose binding lectin protein with Apple-like carbohydrate-binding domain |
| AT5G20080 | AT5G20080 | NADH-cytochrome b5 reductase-like protein |
| AT4G20830 | AT4G20830 | FAD-binding Berberine family protein |
| AT5G44380 | AT5G44380 | FAD-binding Berberine family protein |
| AT5G50950 | FUM2 | Fumarate hydratase 2, chloroplastic |
| AT3G04120 | GAPC1 | Glyceraldehyde-3-phosphate dehydrogenase GAPC1, cytosolic |
| AT4G16260 | AT4G16260 | Probable glucan endo-1,3-beta-glucosidase At4g16260 |
| AT3G13790 | ATBFRUCT1 | Beta-fructofuranosidase, insoluble isoenzyme CWINV1 |
| AT3G19390 | AT3G19390 | Granulin repeat cysteine protease family protein |
| AT3G12580 | HSP70 | Probable mediator of RNA polymerase II transcription subunit 37c |
| AT5G42020 | BIP2 | Mediator of RNA polymerase II transcription subunit 37f |
| AT3G16430 | JAL31 | Jacalin-related lectin 31 |
| AT2G22170 | PLAT2 | Lipase/lipooxygenase, PLAT/LH2 family protein |
| AT3G16460 | JAL34 | Mannose-binding lectin superfamily protein |
| AT3G48890 | MAPR3 | Putative progesterone-binding protein homolog (Atmp2) mRNA |
| AT4G19410 | AT4G19410 | Pectinacetylesterase family protein |
| AT4G30170 | AT4G30170 | Peroxidase family protein |
| AT1G05240 | AT1G05240 | Peroxidase superfamily protein |
| AT2G37130 | AT2G37130 | Peroxidase superfamily protein |

| | | |
|---|---|---|
| AT3G01190 | AT3G01190 | Peroxidase superfamily protein |
| AT5G17820 | AT5G17820 | Peroxidase superfamily protein |
| AT1G79550 | PGK | Encodes cytosolic phosphoglycerate kinase (PGK) |
| AT4G20260 | PCAP1 | Plasma membrane-associated cation-binding protein 1 |
| AT2G47540 | AT2G47540 | Pollen Ole e 1 allergen and extensin family protein |
| AT5G05500 | MOP10 | Pollen Ole e 1 allergen and extensin family protein |
| AT4G14060 | AT4G14060 | Polyketide cyclase/dehydrase and lipid transport superfamily protein |
| AT4G23680 | AT4G23680 | Polyketide cyclase/dehydrase and lipid transport superfamily protein |
| AT2G19760 | PRF1 | Profilin-1 |
| AT4G02270 | RHS13 | Uncharacterized protein At4g02270 |
| AT4G26220 | AT4G26220 | S-adenosyl-L-methionine-dependent methyltransferases superfamily protein |
| AT1G58270 | ZW9 | Uncharacterized protein At1g58270 |
| AT1G45201 | TLL1 | Triacylglycerol lipase-like 1 |
| AT3G55440 | TPI | Triosephosphate isomerase, cytosolic |
| AT4G30270 | XTH24 | Xyloglucan endotransglucosylase/hydrolase protein 24 |
