## Supplementary material for "Evidences for a nutritional role of iodine in plants": Table S13

| LEAVES – Biological processes (GO terms enrichment)<br>*OGC: observed gene count; **BGC: background gene count | | | | | |
|---|---|---|---|---|---|
| #term ID | term description | OGC * | BGC** | FDR | matching proteins in your network (labels) |
| GO:0015979 | photosynthesis | 31 | 215 | 7.82e-53 | AT1G74470,ATPC1,CAB3,FNR1,LHCB2.1,LHCB3,LHCB5,PETA,PETC,PGRL1A,PSAB,PSAE-1,PSAE-2,PSAF,PSAH2,PSB27,PSB29,PSBA,PSBB,PSBC,PSBD,PSBO2,PSBP-1,PSBP-2,PSBQ-2,PSBQA,PSBR,RBCL,RBCS1A,TLP18.3,VAR1 |
| GO:0019684 | photosynthesis, light reaction | 20 | 98 | 8.52e-35 | ATPC1,CAB3,FNR1,LHCB2.1,LHCB3,LHCB5,PETA,PETC,PGRL1A,PSB27,PSB29,PSBA,PSBB,PSBC,PSBD,PSBO2,PSBP-1,PSBR,TLP18.3,VAR1 |
| GO:0006091 | generation of precursor metabolites and energy | 22 | 360 | 4.38e-28 | ATPC1,CAB3,DRT112,FBA2,FNR1,LHCB2.1,LHCB3,LHCB5,PETA,PETC,PGRL1A,PSB27,PSB29,PSBA,PSBB,PSBC,PSBD,PSBO2,PSBP-1,PSBR,TLP18.3,VAR1 |
| GO:0009767 | photosynthetic electron transport chain | 9 | 40 | 3.80e-15 | ATPC1,FNR1,PETA,PETC,PGRL1A,PSBA,PSBB,PSBC,PSBD |
| GO:0018298 | protein-chromophore linkage | 9 | 43 | 5.43e-15 | CAB3,LHCB2.1,LHCB3,LHCB5,PSAB,PSBA,PSBB,PSBC,PSBD |
| GO:0022900 | electron transport chain | 10 | 160 | 5.55e-12 | ATPC1,DRT112,FNR1,PETA,PETC,PGRL1A,PSBA,PSBB,PSBC,PSBD |
| GO:0044237 | cellular metabolic process | 36 | 8432 | 1.45e-11 | AT1G74470,ATPA,ATPC1,CAB3,DRT112,FBA2,FNR1,LHCB2.1,LHCB3,LHCB5,PB,PETA,PETC,PGRL1A,PSAB,PSAE-1,PSAE-2,PSAF,PSAH2,PSB27,PSB29,PSBA,PSBB,PSBC,PSBD,PSBO2,PSBP-1,PSBP-2,PSBQ-2,PSBQA,PSBR,PYK10,RBCL,RBCS1A,TLP18.3,VAR1 |
| GO:0009765 | photosynthesis, light harvesting | 7 | 35 | 1.59e-11 | CAB3,LHCB2.1,LHCB3,LHCB5,PSB27,TLP18.3,VAR1 |
| GO:0010207 | photosystem II assembly | 6 | 15 | 2.21e-11 | LHCB5,PSB27,PSB29,PSBB,PSBO2,PSBR |
| GO:0009772 | photosynthetic electron transport in photosystem II | 5 | 8 | 3.53e-10 | ATPC1,PSBA,PSBB,PSBC,PSBD |
| GO:0055114 | oxidation-reduction process | 15 | 1348 | 2.30e-08 | AT1G74470,ATPC1,DRT112,FNR1,PETA,PETC,PGRL1A,PSAB,PSBA,PSBB,PSBC,PSBD,PSBO2,RBCL,RBCS1A |
| GO:0050896 | response to stimulus | 25 | 5064 | 1.34e-07 | AT5G07020,ATPA,CAB3,DRT112,FBA2,FNR1,LHCB2.1,LHCB3,LHCB5,PB,PETC,PSAE-1,PSAE-2,PSAH2,PSB27,PSB29,PSBA,PSBO2,PSBP-1,PSBR,PYK10,RBCL,RBCS1A,RCA,VAR1 |
| GO:0009416 | response to light stimulus | 10 | 585 | 4.71e-07 | CAB3,LHCB2.1,LHCB3,LHCB5,PETC,PSB27,PSBO2,RBCS1A,RCA,VAR1 |
| GO:0009735 | response to cytokinin | 7 | 212 | 9.89e-07 | AT5G07020,PSAE-1,PSAE-2,PSAH2,PSBR,PYK10,VAR1 |
| GO:0009768 | photosynthesis, light harvesting in photosystem I | 4 | 19 | 9.89e-07 | CAB3,LHCB2.1,LHCB3,LHCB5 |
| GO:0034622 | cellular protein-containing complex assembly | 8 | 332 | 9.89e-07 | LHCB5,PSB27,PSB29,PSBB,PSBO2,PSBR,RBCS1A,VAR1 |
| GO:0010033 | response to organic substance | 14 | 1786 | 4.79e-06 | AT5G07020,CAB3,FBA2,LHCB2.1,LHCB3,PSAE-1,PSAE-2,PSAH2,PSB29,PSBR,PYK10,RBCL,RCA,VAR1 |
| GO:0042221 | response to chemical | 16 | 2654 | 1.59e-05 | AT5G07020,CAB3,DRT112,FBA2,LHCB2.1,LHCB3,PSAE-1,PSAE-2,PSAH2,PSB29,PSBA,PSBR,PYK10,RBCL,RCA,VAR1 |
| GO:0009628 | response to abiotic stimulus | 13 | 1699 | 1.68e-05 | ATPA,CAB3,LHCB2.1,LHCB3,LHCB5,PB,PETC,PSB27,PSBO2,PYK10,RBCS1A,RCA,VAR1 |

| | | | | | |
|---|---|---|---|---|---|
| GO:0009725 | response to hormone | 12 | 1502 | 3.02e-05 | AT5G07020,FBA2,LHCB2.1,LHCB3,PSAE-1,PSAE-2,PSAH2,PSBR,PYK10,RBCL,RCA,VAR1 |
| GO:0010218 | response to far red light | 4 | 57 | 3.50e-05 | CAB3,LHCB2.1,LHCB5,RBCS1A |
| GO:0010206 | photosystem II repair | 3 | 15 | 3.90e-05 | PSB27,TLP18.3,VAR1 |
| GO:0009642 | response to light intensity | 5 | 142 | 4.20e-05 | LHCB2.1,LHCB3,PSB27,PSBO2,VAR1 |
| GO:1901564 | organonitrogen compound metabolic process | 19 | 4116 | 4.20e-05 | AT1G74470,ATPA,ATPC1,CAB3,FBA2,LHCB2.1,LHCB3,LHCB5,PB,PSAB,PSB27,PSB29,PSBA,PSBB,PSBC,PSBD,PYK10,TLP18.3,VAR1 |
| GO:0010114 | response to red light | 4 | 64 | 4.74e-05 | CAB3,LHCB2.1,LHCB5,RBCS1A |
| GO:0009644 | response to high light intensity | 4 | 76 | 8.81e-05 | LHCB2.1,LHCB3,PSBO2,VAR1 |
| GO:0009637 | response to blue light | 4 | 80 | 0.00010 | CAB3,LHCB2.1,LHCB5,RBCS1A |
| GO:0006754 | ATP biosynthetic process | 4 | 89 | 0.00015 | ATPA,ATPC1,FBA2,PB |
| GO:0009409 | response to cold | 6 | 347 | 0.00018 | ATPA,LHCB2.1,PB,PYK10,RBCS1A,RCA |
| GO:0015986 | ATP synthesis coupled proton transport | 3 | 31 | 0.00020 | ATPA,ATPC1,PB |
| GO:0009168 | purine ribonucleoside monophosphate biosynthetic process | 4 | 110 | 0.00027 | ATPA,ATPC1,FBA2,PB |
| GO:0009769 | photosynthesis, light harvesting in photosystem II | 2 | 4 | 0.00031 | LHCB2.1,LHCB3 |
| GO:0010196 | nonphotochemical quenching | 2 | 6 | 0.00052 | LHCB5,PETC |
| GO:1903426 | regulation of reactive oxygen species biosynthetic process | 2 | 7 | 0.00059 | LHCB2.1,PSB29 |
| GO:0042742 | defense response to bacterium | 5 | 315 | 0.00080 | ATPA,FNR1,PETC,PSBP-1,RCA |
| GO:0010119 | regulation of stomatal movement | 3 | 70 | 0.0012 | LHCB2.1,LHCB3,PSB29 |
| GO:0009635 | response to herbicide | 2 | 13 | 0.0014 | LHCB3,PSBA |
| GO:0010205 | photoinhibition | 2 | 14 | 0.0015 | PSBO2,VAR1 |
| GO:0090662 | ATP hydrolysis coupled transmembrane transport | 3 | 77 | 0.0015 | ATPA,ATPC1,PB |

| | | | | | |
|---|---|---|---|---|---|
| GO:0099131 | ATP hydrolysis coupled ion transmembrane transport | 3 | 77 | 0.0015 | ATPA,ATPC1,PB |
| GO:0099132 | ATP hydrolysis coupled cation transmembrane transport | 3 | 78 | 0.0015 | ATPA,ATPC1,PB |
| GO:0019253 | reductive pentose-phosphate cycle | 2 | 20 | 0.0026 | RBCL,RBCS1A |
| GO:0044267 | cellular protein metabolic process | 12 | 2826 | 0.0039 | CAB3,LHCB2.1,LHCB3,LHCB5,PSAB,PSB27,PSBA,PSBB,PSBC,PSBD,TLP18.3,VAR1 |
| GO:0051707 | response to other organism | 7 | 1079 | 0.0055 | ATPA,FNR1,PB,PETC,PSBP-1,PYK10,RCA |
| GO:0098542 | defense response to other organism | 6 | 847 | 0.0078 | ATPA,FNR1,PB,PETC,PSBP-1,RCA |
| GO:0009853 | photorespiration | 2 | 49 | 0.0115 | RBCL,RBCS1A |
| GO:0015994 | chlorophyll metabolic process | 2 | 56 | 0.0145 | AT1G74470,PSB29 |
| GO:0006950 | response to stress | 11 | 2932 | 0.0154 | ATPA,FNR1,LHCB2.1,LHCB3,PB,PETC,PSBA,PSBP-1,PYK10,RBCS1A,RCA |
| GO:1901135 | carbohydrate derivative metabolic process | 5 | 701 | 0.0170 | ATPA,ATPC1,FBA2,PB,PYK10 |
| GO:0044270 | cellular nitrogen compound catabolic process | 3 | 214 | 0.0174 | FBA2,PSB29,PYK10 |
| GO:0046700 | heterocycle catabolic process | 3 | 214 | 0.0174 | FBA2,PSB29,PYK10 |
| GO:1901700 | response to oxygen-containing compound | 7 | 1398 | 0.0191 | CAB3,FBA2,LHCB2.1,LHCB3,PSB29,RBCL,RCA |
| GO:0019439 | aromatic compound catabolic process | 3 | 238 | 0.0214 | FBA2,PSB29,PYK10 |
| GO:0016043 | cellular component organization | 9 | 2271 | 0.0237 | LHCB5,PSB27,PSB29,PSBB,PSBO2,PSBR,PYK10,RBCS1A,VAR1 |
| GO:0051704 | multi-organism process | 7 | 1475 | 0.0237 | ATPA,FNR1,PB,PETC,PSBP-1,PYK10,RCA |
| GO:1901361 | organic cyclic compound catabolic process | 3 | 249 | 0.0237 | FBA2,PSB29,PYK10 |
| GO:0009737 | response to abscisic acid | 4 | 511 | 0.0268 | FBA2,LHCB2.1,LHCB3,RBCL |
| GO:0071704 | organic substance metabolic process | 21 | 8632 | 0.0319 | AT1G74470,ATPA,ATPC1,CAB3,FBA2,LHCB2.1,LHCB3,LHCB5,PB,PSAB,PSB27,PSB29,PSBA,PSBB,PSBC,PSBD,PYK10,RBCL,RBCS1A,TLP18.3,VAR1 |
| GO:0016051 | carbohydrate biosynthetic process | 3 | 295 | 0.0351 | FBA2,RBCL,RBCS1A |

| | | | | | |
|---|---|---|---|---|---|
| GO:0017144 | drug metabolic process | 4 | 626 | 0.0498 | ATPA,ATPC1,FBA2,PB |

### LEAVES – Molecular functions (GO terms enrichment)

*OGC: observed gene count; **BGC: background gene count

| #term ID | term description | OGC * | BGC** | FDR | matching proteins in your network (labels) |
|---|---|---|---|---|---|
| GO:0016168 | chlorophyll binding | 9 | 30 | 4.35e-16 | CAB3,LHCB2.1,LHCB3,LHCB5,PSAB,PSBA,PSBB,PSBC,PSBD |
| GO:0019904 | protein domain specific binding | 10 | 71 | 2.22e-15 | DRT112,FNR1,LHCB2.1,LHCB3,LHCB5,PGRL1A,PSAE-1,PSAE-2,PSAF,PSBP-1 |
| GO:0046906 | tetrapyrrole binding | 11 | 299 | 3.86e-11 | CAB3,LHCB2.1,LHCB3,LHCB5,MAPR2,PETA,PSAB,PSBA,PSBB,PSBC,PSBD |
| GO:0009055 | electron transfer activity | 8 | 105 | 1.95e-10 | DRT112,FNR1,PETA,PETC,PSBA,PSBB,PSBC,PSBD |
| GO:0045156 | electron transporter, transferring electrons within the cyclic electron transport pathway of photosynthesis activity | 5 | 9 | 3.01e-10 | FNR1,PSBA,PSBB,PSBC,PSBD |
| GO:0046872 | metal ion binding | 22 | 2940 | 5.68e-10 | ATPA,CAB3,DRT112,LHCB2.1,LHCB3,LHCB5,ORF110A,PB,PETA,PETC,PSAB,PSBA,PSBC,PSBD,PSBP-1,PSBP-2,PSBQ-2,PSBQA,PYK10,RBCL,RBCS1A,VAR1 |
| GO:0016491 | oxidoreductase activity | 14 | 1201 | 2.46e-08 | AT1G74470,DRT112,FNR1,PETA,PETC,PGRL1A,PSAB,PSBA,PSBB,PSBC,PSBD,PSBO2,RBCL,RBCS1A |
| GO:0048037 | cofactor binding | 12 | 860 | 5.80e-08 | CAB3,LHCB2.1,LHCB3,LHCB5,MAPR2,PETA,PETC,PSAB,PSBA,PSBB,PSBC,PSBD |
| GO:0005488 | binding | 31 | 8611 | 2.45e-07 | ATPA,CAB3,DRT112,FNR1,LHCB2.1,LHCB3,LHCB5,MAPR2,ORF110A,PB,PETA,PETC,PGRL1A,PSAB,PSAE-1,PSAE-2,PSAF,PSBA,PSBB,PSBC,PSBD,PSBO2,PSBP-1,PSBP-2,PSBQ-2,PSBQA,PYK10,RBCL,RBCS1A,RCA,VAR1 |
| GO:0043167 | ion binding | 24 | 5070 | 2.97e-07 | ATPA,CAB3,DRT112,LHCB2.1,LHCB3,LHCB5,ORF110A,PB,PETA,PETC,PSAB,PSBA,PSBB,PSBC,PSBD,PSBP-1,PSBP-2,PSBQ-2,PSBQA,PYK10,RBCL,RBCS1A,RCA,VAR1 |
| GO:0031409 | pigment binding | 4 | 19 | 3.96e-07 | CAB3,LHCB2.1,LHCB3,LHCB5 |
| GO:0008266 | poly(U) RNA binding | 3 | 16 | 2.83e-05 | FNR1,PSBO2,PSBP-1 |
| GO:0046933 | proton-transporting ATP synthase activity, rotational mechanism | 3 | 19 | 3.88e-05 | ATPA,ATPC1,PB |
| GO:0046028 | electron transporter, transferring electrons from cytochrome b6/f complex of | 2 | 2 | 9.92e-05 | DRT112,PETC |

| | | | | | |
|---|---|---|---|---|---|
| | photosystem II activity | | | | |
| GO:0044769 | ATPase activity, coupled to transmembrane movement of ions, rotational mechanism | 3 | 37 | 0.00022 | ATPA,ATPC1,PB |
| GO:0016984 | ribulose-bisphosphate carboxylase activity | 2 | 5 | 0.00031 | RBCL,RBCS1A |
| GO:0016730 | oxidoreductase activity, acting on iron-sulfur proteins as donors | 2 | 8 | 0.00059 | FNR1,PGRL1A |
| GO:0043168 | anion binding | 13 | 2629 | 0.00059 | ATPA,CAB3,LHCB2.1,LHCB3,LHCB5,PB,PSAB,PSBA,PSBB,PSBC,PSBD,RCA,VAR1 |
| GO:0005515 | protein binding | 11 | 1988 | 0.00081 | DRT112,FNR1,LHCB2.1,LHCB3,LHCB5,PGRL1A,PSAE-1,PSAE-2,PSAF,PSBP-1,PYK10 |
| GO:0019829 | cation-transporting ATPase activity | 3 | 74 | 0.0011 | ATPA,ATPC1,PB |
| GO:0005509 | calcium ion binding | 4 | 226 | 0.0019 | PSBP-1,PSBP-2,PSBQ-2,PSBQA |
| GO:0016830 | carbon-carbon lyase activity | 3 | 96 | 0.0020 | FBA2,RBCL,RBCS1A |
| GO:0003824 | catalytic activity | 21 | 7239 | 0.0039 | AT1G74470,ATPA,ATPC1,DRT112,FBA2,FNR1,PB,PETA,PETC,PGRL1A,PSAB,PSBA,PSBB,PSBC,PSBD,PSBO2,PYK10,RBCL,RBCS1A,TLP18.3,VAR1 |
| GO:0097159 | organic cyclic compound binding | 18 | 5841 | 0.0055 | ATPA,CAB3,FNR1,LHCB2.1,LHCB3,LHCB5,MAPR2,PB,PETA,PSAB,PSBA,PSBB,PSBC,PSBD,PSBO2,PSBP-1,RCA,VAR1 |
| GO:1901363 | heterocyclic compound binding | 18 | 5835 | 0.0055 | ATPA,CAB3,FNR1,LHCB2.1,LHCB3,LHCB5,MAPR2,PB,PETA,PSAB,PSBA,PSBB,PSBC,PSBD,PSBO2,PSBP-1,RCA,VAR1 |
| GO:0005507 | copper ion binding | 3 | 157 | 0.0071 | DRT112,PYK10,RBCS1A |
| GO:0046914 | transition metal ion binding | 6 | 933 | 0.0101 | ATPA,DRT112,PB,PETA,PYK10,RBCS1A |
| GO:0042623 | ATPase activity, coupled | 4 | 391 | 0.0105 | ATPA,ATPC1,PB,VAR1 |
| GO:0043492 | ATPase activity, coupled to movement of substances | 3 | 199 | 0.0118 | ATPA,ATPC1,PB |
| GO:0015405 | P-P-bond-hydrolysis-driven transmembrane transporter activity | 3 | 207 | 0.0127 | ATPA,ATPC1,PB |

| | | | | | |
|---|---|---|---|---|---|
| GO:0016887 | ATPase activity | 4 | 498 | 0.0202 | ATPA,ATPC1,PB,VAR1 |
| GO:0000287 | magnesium ion binding | 2 | 115 | 0.0374 | ORF110A,RBCL |
