## Supplementary material for "Evidences for a nutritional role of iodine in plants": Table S14

| Mercator analysis | | |
|---|---|---|
| **IDENTIFIER** | **BINCODE** | **NAME** |
| atcg00120.1 | '1.1.9.2.1' | 'Photosynthesis.photophosphorylation.ATP synthase complex.peripheral CF1 subcomplex.subunit alpha' |
| atcg00480.1 | '1.1.9.2.2' | 'Photosynthesis.photophosphorylation.ATP synthase complex.peripheral CF1 subcomplex.subunit beta' |
| at4g04640.1 | '1.1.9.2.3' | 'Photosynthesis.photophosphorylation.ATP synthase complex.peripheral CF1 subcomplex.subunit gamma' |
| at1g29910.1 | '1.1.1.1.1' | 'Photosynthesis.photophosphorylation.photosystem II.LHC-II complex.component LHCb1/2/3' |
| at1g20340.1 | '1.1.3.1' | 'Photosynthesis.photophosphorylation.Cytb6/f to PS-I electron carriers.plastocyanin' |
| at5g66190.1 | '1.1.5.2.1' | 'Photosynthesis.photophosphorylation.linear electron flow.ferredoxin-NADP reductase (FNR) activity.ferredoxin-NADP oxidoreductase' |
| at4g38970.1 | '1.2.5' | 'Photosynthesis.calvin cycle.fructose 1,6-bisphosphate aldolase' |
| at4g38970.1 | '3.12.2' | 'Carbohydrate metabolism.plastidial glycolysis.fructose-1,6-bisphosphate aldolase' |
| at5g42270.1 | '19.4.5.8.2.1' | 'Protein homeostasis.proteolysis.metallopeptidase activities.FtsH endopeptidase activities.FtsH plastidial protease complexes.component FtsH1\|2\|5\|6\|8' |
| at3g09260.1 | '50.3.2' | 'Enzyme classification.EC_3 hydrolases.EC_3.2 glycosylase' |
| at4g10340.1 | '1.1.1.1.3' | 'Photosynthesis.photophosphorylation.photosystem II.LHC-II complex.component LHCb5' |
| at5g54270.1 | '1.1.1.1.1' | 'Photosynthesis.photophosphorylation.photosystem II.LHC-II complex.component LHCb1/2/3' |
| at2g34430.1 | '1.1.1.1.1' | 'Photosynthesis.photophosphorylation.photosystem II.LHC-II complex.component LHCb1/2/3' |
| at2g24940.1 | '11.3.2.1.4' | 'Phytohormone action.brassinosteroid.perception and signal transduction.receptor complex.receptor kinase regulator protein (MSBP)' |
| at4g22890.1 | '1.1.6.1.2' | 'Photosynthesis.photophosphorylation.cyclic electron flow.PGR5/PGRL1 complex.component PGRL1-like' |
| atcg00540.1 | '1.1.2.1' | 'Photosynthesis.photophosphorylation.cytochrome b6/f complex.apocytochrome f component PetA' |
| at4g03280.1 | '1.1.2.3' | 'Photosynthesis.photophosphorylation.cytochrome b6/f complex.Rieske iron-sulfur component PetC' |
| at4g28750.1 | '1.1.4.2.5' | 'Photosynthesis.photophosphorylation.photosystem I.PS-I complex.component PsaE' |
| at2g20260.1 | '1.1.4.2.5' | 'Photosynthesis.photophosphorylation.photosystem I.PS-I complex.component PsaE' |
| at1g31330.1 | '1.1.4.2.6' | 'Photosynthesis.photophosphorylation.photosystem I.PS-I complex.component PsaF' |
| at1g52230.1 | '1.1.4.2.8' | 'Photosynthesis.photophosphorylation.photosystem I.PS-I complex.component PsaH' |
| atcg00340.1 | '1.1.4.2.2' | 'Photosynthesis.photophosphorylation.photosystem I.PS-I complex.apoprotein component PsaB' |
| at1g03600.1 | '1.1.1.3.8' | 'Photosynthesis.photophosphorylation.photosystem II.assembly and maintenance.regulatory protein (Psb27)' |
| at1g06680.1 | '1.1.1.2.2.2.1' | 'Photosynthesis.photophosphorylation.photosystem II.PS-II complex.oxygen-evolving center (OEC) extrinsic proteins.Viridiplantae-specific components.component OEC23/PsbP' |
| at1g79040.1 | '1.1.1.2.9' | 'Photosynthesis.photophosphorylation.photosystem II.PS-II complex.component PsbR' |

| | | |
|---|---|---|
| at2g05100.1 | '1.1.1.1.1' | 'Photosynthesis.photophosphorylation.photosystem II.LHC-II complex.component LHCb1/2/3' |
| at2g20890.1 | '1.1.1.3.10' | 'Photosynthesis.photophosphorylation.photosystem II.assembly and maintenance.assembly factor (Psb29)' |
| at2g30790.1 | '1.1.1.2.2.2.1' | 'Photosynthesis.photophosphorylation.photosystem II.PS-II complex.oxygen-evolving center (OEC) extrinsic proteins.Viridiplantae-specific components.component OEC23/PsbP' |
| at3g50820.1 | '1.1.1.2.2.1' | 'Photosynthesis.photophosphorylation.photosystem II.PS-II complex.oxygen-evolving center (OEC) extrinsic proteins.component OEC33/PsbO' |
| at4g05180.1 | '1.1.1.2.2.2.2' | 'Photosynthesis.photophosphorylation.photosystem II.PS-II complex.oxygen-evolving center (OEC) extrinsic proteins.Viridiplantae-specific components.component OEC16/PsbQ' |
| at4g21280.2 | '1.1.1.2.2.2.2' | 'Photosynthesis.photophosphorylation.photosystem II.PS-II complex.oxygen-evolving center (OEC) extrinsic proteins.Viridiplantae-specific components.component OEC16/PsbQ' |
| atcg00020.1 | '1.1.1.2.1.1' | 'Photosynthesis.photophosphorylation.photosystem II.PS-II complex.reaction center complex.component D1/PsbA' |
| atcg00270.1 | '1.1.1.2.1.2' | 'Photosynthesis.photophosphorylation.photosystem II.PS-II complex.reaction center complex.component D2/PsbD' |
| atcg00280.1 | '1.1.1.2.1.4' | 'Photosynthesis.photophosphorylation.photosystem II.PS-II complex.reaction center complex.component CP43/PsbC' |
| atcg00680.1 | '1.1.1.2.1.3' | 'Photosynthesis.photophosphorylation.photosystem II.PS-II complex.reaction center complex.component CP47/PsbB' |
| at5g07020.1 | '1.1.1.5.2' | 'Photosynthesis.photophosphorylation.photosystem II.photoprotection.protein (MPH1)' |
| at1g74470.1 | '7.12.6.6.3' | 'Coenzyme metabolism.tetrapyrrol biosynthesis.chlorophyll metabolism.chlorophyll(ide) interconversions.geranylgeranyl reductase (ChlP)' |
| atmg00280.1 | '35.1' | 'not assigned.annotated' |
| at1g67090.1 | '1.2.1.1.2' | 'Photosynthesis.calvin cycle.ribulose-1,5-bisphosphat carboxylase/oxygenase (RuBisCo) activity.RuBisCo heterodimer.small subunit' |
| atcg00490.1 | '1.2.1.1.1' | 'Photosynthesis.calvin cycle.ribulose-1,5-bisphosphat carboxylase/oxygenase (RuBisCo) activity.RuBisCo heterodimer.large subunit' |
| at1g71500.1 | '1.1.1.3.13' | 'Photosynthesis.photophosphorylation.photosystem II.assembly and maintenance.stabilizing factor (Psb33)' |
| at2g39730.1 | '1.2.1.3.2' | 'Photosynthesis.calvin cycle.ribulose-1,5-bisphosphat carboxylase/oxygenase (RuBisCo) activity.RuBisCo regulation.ATP-dependent activase (RCA)' |
| at1g54780.1 | '1.1.1.3.12' | 'Photosynthesis.photophosphorylation.photosystem II.assembly and maintenance.assembly factor (Psb32)' |
| at5g09810.1 | '20.2.1' | 'Cytoskeleton organisation.microfilament network.actin filament protein' |
| at1g28290.1 | '35.1' | 'not assigned.annotated' |
| at5g08680.1 | '2.4.6.2.2' | 'Cellular respiration.oxidative phosphorylation.ATP synthase complex.peripheral MF1 subcomplex.subunit beta' |
| at1g45130.1 | '21.3.2.2.2' | 'Cell wall organisation.pectin.rhamnogalacturonan I.modification and degradation.beta-galactosidase (BGAL)' |
| at2g43610.1 | '35.1' | 'not assigned.annotated' |
| at3g43670.1 | '8.5.1' | 'Polyamine metabolism.polyamine degradation.copper-containing amine oxidase (CuAO)' |
| at1g78850.1 | '35.1' | 'not assigned.annotated' |

| | | |
|---|---|---|
| at5g20080.1 | '50.1.6' | 'Enzyme classification.EC_1 oxidoreductases.EC_1.6 oxidoreductase acting on NADH or NADPH' |
| at4g20830.1 | '50.1.1' | 'Enzyme classification.EC_1 oxidoreductases.EC_1.1 oxidoreductase acting on CH-OH group of donor' |
| at5g44380.1 | '50.1.1' | 'Enzyme classification.EC_1 oxidoreductases.EC_1.1 oxidoreductase acting on CH-OH group of donor' |
| at5g50950.1 | '2.3.7' | 'Cellular respiration.tricarboxylic acid cycle.fumarase' |
| at3g04120.1 | '2.1.1.4.1' | 'Cellular respiration.glycolysis.cytosolic glycolysis.glyceraldehyde 3-phosphate dehydrogenase activities.NAD-dependent glyceraldehyde 3-phosphate dehydrogenase' |
| at4g16260.1 | '50.3.2' | 'Enzyme classification.EC_3 hydrolases.EC_3.2 glycosylase' |
| at3g13790.1 | '3.1.4.1.1' | 'Carbohydrate metabolism.sucrose metabolism.degradation.invertase activities.acid beta-fructofuranosidase (CWIN)' |
| at3g19390.1 | '50.3.4' | 'Enzyme classification.EC_3 hydrolases.EC_3.4 hydrolase acting on peptide bond (peptidase)' |
| at3g12580.1 | '19.1.5.1' | 'Protein homeostasis.protein quality control.cytosolic Hsp70 chaperone system.chaperone (Hsp70)' |
| at5g42020.1 | '35.1' | 'not assigned.annotated' |
| at3g16430.1 | '35.1' | 'not assigned.annotated' |
| at2g22170.1 | '35.1' | 'not assigned.annotated' |
| at3g16460.1 | '35.1' | 'not assigned.annotated' |
| at3g48890.1 | '11.3.2.1.4' | 'Phytohormone action.brassinosteroid.perception and signal transduction.receptor complex.receptor kinase regulator protein (MSBP)' |
| at4g19410.1 | '21.3.5.4' | 'Cell wall organisation.pectin.modification and degradation.pectin acetylesterase' |
| at4g30170.1 | '35.1' | 'not assigned.annotated' |
| at1g05240.1 | '35.1' | 'not assigned.annotated' |
| at2g37130.1 | '35.1' | 'not assigned.annotated' |
| at3g01190.1 | '35.1' | 'not assigned.annotated' |
| at5g17820.1 | '35.1' | 'not assigned.annotated' |
| at1g79550.1 | '2.1.1.5' | 'Cellular respiration.glycolysis.cytosolic glycolysis.phosphoglycerate kinase' |
| at4g20260.4 | '35.1' | 'not assigned.annotated' |
| at2g47540.1 | '21.4.1.2.1' | 'Cell wall organisation.cell wall proteins.hydroxyproline-rich glycoprotein activities.proline-rich protein activities.glycoprotein' |
| at5g05500.1 | '35.2' | 'not assigned.not annotated' |
| at4g14060.1 | '35.1' | 'not assigned.annotated' |
| at4g23680.1 | '35.1' | 'not assigned.annotated' |
| at2g19760.1 | '20.2.2.3' | 'Cytoskeleton organisation.microfilament network.actin polymerisation.profilin actin nucleation protein' |
| at4g02270.1 | '21.4.1.2.1' | 'Cell wall organisation.cell wall proteins.hydroxyproline-rich glycoprotein activities.proline-rich protein activities.glycoprotein' |
| at4g26220.1 | '21.6.1.4' | 'Cell wall organisation.lignin.monolignol biosynthesis.caffeoyl-CoA 3-O-methyltransferase (CCoA-OMT)' |
| at1g58270.1 | '35.2' | 'not assigned.not annotated' |

| | | |
|---|---|---|
| at1g45201.1 | '5.7.1.2.3' | 'Lipid metabolism.lipid degradation.triacylglycerol lipase activities.diacyl-/triacylglycerol lipase activities.lipase (OBL)' |
| at3g55440.1 | '2.1.1.3' | 'Cellular respiration.glycolysis.cytosolic glycolysis.triosephosphate isomerase' |
| at3g55440.1 | '3.1.2.1' | 'Carbohydrate metabolism.sucrose metabolism.biosynthesis.cytosolic triose-phosphate isomerase' |
| at4g30270.1 | '50.2.4' | 'Enzyme classification.EC_2 transferases.EC_2.4 glycosyltransferase' |

| Mercator analysis | | |
|---|---|---|
| IDENTIFIER | BINCODE | DESCRIPTION |
| atcg00120.1 | '1.1.9.2.1' | subunit Symbols:ATPA \| ATP synthase subunit alpha \| ChrC:9938-11461 REVERSE LENGTH=507 ) &' |
| atcg00480.1 | '1.1.9.2.2' | subunit Symbols:PB,ATPB \| ATP synthase subunit beta \| ChrC:52660-54156 REVERSE LENGTH=498 ) &' |
| at4g04640.1 | '1.1.9.2.3' | subunit Symbols:ATPC1 \| \| Chr4:2350761-2351882 REVERSE LENGTH=373 ) &' |
| at1g29910.1 | '1.1.1.1.1' | component Symbols:CAB3,LHCB1.2,AB180 \| LIGHT HARVESTING CHLOROPHYLL A/B BINDING PROTEIN 1.2,chlorophyll A/B binding protein 3 \| Chr1:10472443-10473246 REVERSE LENGTH=267 ) &' |
| at1g20340.1 | '1.1.3.1' | plastocyanin Symbols:PETE2,DRT112 \| DNA-DAMAGE-REPAIR/TOLERATION PROTEIN 112,PLASTOCYANIN 2 \| Chr1:7042770-7043273 REVERSE LENGTH=167 ) &' |
| at5g66190.1 | '1.1.5.2.1' | ferredoxin-NADP Symbols:LFNR1,ATLFNR1,FNR1 \| leaf-type chloroplast-targeted FNR 1,LEAF FNR 1,ferredoxin-NADP(+)-oxidoreductase 1 \| Chr5:26451203-26453012 REVERSE LENGTH=360 ) &' |
| at4g38970.1 | '1.2.5' | fructose Symbols:AtFBA2,FBA2 \| fructose-bisphosphate aldolase 2 \| Chr4:18163714-18165659 REVERSE LENGTH=398 ) &' |
| at4g38970.1 | '3.12.2' | fructose-1,6-bisphosphate Symbols:AtFBA2,FBA2 \| fructose-bisphosphate aldolase 2 \| Chr4:18163714-18165659 REVERSE LENGTH=398 ) &' |
| at5g42270.1 | '19.4.5.8.2.1' | component Symbols:FTSH5,VAR1 \| VARIEGATED 1 \| Chr5:16902659-16905102 FORWARD LENGTH=704 ) &' |
| at3g09260.1 | '50.3.2' | Beta-glucosidase Symbols:LEB,BGLU23,PYK10,PSR3.1 \| LONG ER BODY \| Chr3:2840657-2843730 REVERSE LENGTH=524 ) &' |
| at4g10340.1 | '1.1.1.1.3' | component Symbols:LHCB5 \| light harvesting complex of photosystem II 5 \| Chr4:6408200-6409496 FORWARD LENGTH=280 ) &' |
| at5g54270.1 | '1.1.1.1.1' | component Symbols:LHCB3,LHCB3*1 \| light-harvesting chlorophyll B-binding protein 3 \| Chr5:22038424-22039383 FORWARD LENGTH=265 ) &' |
| at2g34430.1 | '1.1.1.1.1' | component Symbols:LHCB1.4,LHB1B1 \| light-harvesting chlorophyll-protein complex II subunit B1,LIGHT-HARVESTING CHLOROPHYLL-PROTEIN COMPLEX II SUBUNIT B1 \| Chr2:14524818-14525618 FORWARD LENGTH=266 ) &' |
| at2g24940.1 | '11.3.2.1.4' | brassinosteroid Symbols:MAPR2,AtMAPR2 \| membrane-associated progesterone binding protein 2 \| Chr2:10609447-10609749 FORWARD LENGTH=100 ) &' |
| at4g22890.1 | '1.1.6.1.2' | component Symbols:PGR5-LIKE A \| \| Chr4:12007157-12009175 FORWARD LENGTH=324 ) &' |
| atcg00540.1 | '1.1.2.1' | apocytochrome Symbols:PETA \| photosynthetic electron transfer A \| ChrC:61657-62619 FORWARD LENGTH=320 ) &' |
| at4g03280.1 | '1.1.2.3' | Rieske Symbols:PGR1,PETC \| PROTON GRADIENT REGULATION 1,photosynthetic electron transfer C \| Chr4:1440314-1441717 FORWARD LENGTH=229 ) &' |

| | | |
|---|---|---|
| at4g28750.1 | '1.1.4.2.5' | component Symbols:PSAE-1 \| PSA E1 KNOCKOUT \| Chr4:14202951-14203888 REVERSE LENGTH=143 ) &' |
| at2g20260.1 | '1.1.4.2.5' | component Symbols:PSAE-2 \| photosystem I subunit E-2 \| Chr2:8736780-8737644 FORWARD LENGTH=145 ) &' |
| at1g31330.1 | '1.1.4.2.6' | component Symbols:PSAF \| photosystem I subunit F \| Chr1:11215011-11215939 REVERSE LENGTH=221 ) &' |
| at1g52230.1 | '1.1.4.2.8' | component Symbols:PSI-H,PSAH-2,PSAH2 \| photosystem I subunit H2,PHOTOSYSTEM I SUBUNIT H-2 \| Chr1:19454902-19455508 FORWARD LENGTH=145 ) &' |
| atcg00340.1 | '1.1.4.2.2' | apoprotein Symbols:PSAB \| \| ChrC:37375-39579 REVERSE LENGTH=734 ) &' |
| at1g03600.1 | '1.1.1.3.8' | Psb27 Symbols:PSB27 \| \| Chr1:898916-899440 FORWARD LENGTH=174 ) &' |
| at1g06680.1 | '1.1.1.2.2.2.1' | component Symbols:OE23,PSII-P,PSBP-1,OEE2 \| OXYGEN-EVOLVING ENHANCER PROTEIN 2,PHOTOSYSTEM II SUBUNIT P,photosystem II subunit P-1,OXYGEN EVOLVING COMPLEX SUBUNIT 23 KDA \| Chr1:2047940-2049186 FORWARD LENGTH=263 ) &' |
| at1g79040.1 | '1.1.1.2.9' | component Symbols:PSBR \| photosystem II subunit R \| Chr1:29736085-29736781 FORWARD LENGTH=140 ) &' |
| at2g05100.1 | '1.1.1.1.1' | component Symbols:LHCB2,LHCB2.1 \| LIGHT-HARVESTING CHLOROPHYLL B-BINDING 2,photosystem II light harvesting complex gene 2.1 \| Chr2:1823449-1824331 REVERSE LENGTH=265 ) &' |
| at2g20890.1 | '1.1.1.3.10' | Psb29 Symbols:THF1,PSB29 \| THYLAKOID FORMATION1,photosystem II reaction center PSB29 protein \| Chr2:8987783-8989185 FORWARD LENGTH=300 ) &' |
| at2g30790.1 | '1.1.1.2.2.2.1' | component Symbols:PSBP-2 \| photosystem II subunit P-2 \| Chr2:13119047-13119596 REVERSE LENGTH=125 ) &' |
| at3g50820.1 | '1.1.1.2.2.1' | component Symbols:OEC33,PSBO-2,PSBO2 \| photosystem II subunit O-2,OXYGEN EVOLVING COMPLEX SUBUNIT 33 KDA,PHOTOSYSTEM II SUBUNIT O-2 \| Chr3:18891008-18892311 REVERSE LENGTH=331 ) &' |
| at4g05180.1 | '1.1.1.2.2.2.2' | component Symbols:PSBQ-2,PSBQ,PSII-Q \| photosystem II subunit Q-2,PHOTOSYSTEM II SUBUNIT Q \| Chr4:2672093-2673170 REVERSE LENGTH=230 ) &' |
| at4g21280.2 | '1.1.1.2.2.2.2' | component Symbols:PSBQ-1,PSBQA,PSBQ \| photosystem II subunit QA,PHOTOSYSTEM II SUBUNIT Q-1,PHOTOSYSTEM II SUBUNIT Q \| Chr4:11334446-11335587 FORWARD LENGTH=224 ) &' |
| atcg00020.1 | '1.1.1.2.1.1' | component Symbols:PSBA \| photosystem II reaction center protein A \| ChrC:383-1444 REVERSE LENGTH=353 ) &' |
| atcg00270.1 | '1.1.1.2.1.2' | component Symbols:PSBD \| photosystem II reaction center protein D \| ChrC:32711-33772 FORWARD LENGTH=353 ) &' |
| atcg00280.1 | '1.1.1.2.1.4' | component Symbols:PSBC \| photosystem II reaction center protein C \| ChrC:33720-35141 FORWARD LENGTH=473 ) &' |
| atcg00680.1 | '1.1.1.2.1.3' | component Symbols:PSBB \| photosystem II reaction center protein B \| ChrC:72371-73897 FORWARD LENGTH=508 ) &' |

| | | |
|---|---|---|
| at5g07020.1 | '1.1.1.5.2' | MPH1 Symbols:MPH1 \| Maintenance of PSII under High light 1 \| Chr5:2180669-2182284 REVERSE LENGTH=235 ) &' |
| at1g74470.1 | '7.12.6.6.3' | geranylgeranyl Symbols:no symbol available \| no full name available \| Chr1:27991248-27992845 FORWARD LENGTH=467 ) &' |
| atmg00280.1 | '35.1' | Symbols:ORF110A \| \| ChrM:77819-78151 REVERSE LENGTH=110 ) & Putative uncharacterized mitochondrial protein AtMg00280 OS=Arabidopsis thaliana (sp\|p93292\|m280_arath : 225.0)' |
| at1g67090.1 | '1.2.1.1.2' | small Symbols:RBCS1A \| ribulose bisphosphate carboxylase small chain 1A \| Chr1:25048465-25049249 REVERSE LENGTH=180 ) &' |
| atcg00490.1 | '1.2.1.1.1' | large Symbols:RBCL \| \| ChrC:54958-56397 FORWARD LENGTH=479 ) &' |
| at1g71500.1 | '1.1.1.3.13' | Psb33 Symbols:PSB33 \| PhotoSystem B protein 33 \| Chr1:26936084-26937331 FORWARD LENGTH=287 ) &' |
| at2g39730.1 | '1.2.1.3.2' | ATP-dependent Symbols:RCA \| rubisco activase \| Chr2:16570951-16573345 REVERSE LENGTH=474 ) &' |
| at1g54780.1 | '1.1.1.3.12' | Psb32 Symbols:TLP18.3,AtTLP18.3 \| thylakoid lumen protein 18.3 \| Chr1:20439533-20440953 FORWARD LENGTH=285 ) &' |
| at5g09810.1 | '20.2.1' | actin Symbols:AtACT7,ACT7 \| actin 7 \| Chr5:3052809-3054220 FORWARD LENGTH=377 ) &' |
| at1g28290.1 | '35.1' | Symbols:AGP31 \| arabinogalactan protein 31 \| Chr1:9889331-9890843 REVERSE LENGTH=359 ) & Non-classical arabinogalactan protein 31 OS=Arabidopsis thaliana (sp\|q9fza2\|agp31_arath : 249.0)' |
| at5g08680.1 | '2.4.6.2.2' | subunit Symbols:no symbol available \| no full name available \| Chr5:2821992-2824683 FORWARD LENGTH=559 ) &' |
| at1g45130.1 | '21.3.2.2.2' | beta-galactosidase Symbols:AtBGAL5,BGAL5 \| beta-galactosidase 5 \| Chr1:17065447-17069110 FORWARD LENGTH=732 ) &' |
| at2g43610.1 | '35.1' | Symbols:no symbol available \| no full name available \| Chr2:18088058-18089184 REVERSE LENGTH=281 ) & Endochitinase At2g43610 OS=Arabidopsis thaliana (sp\|o22842\|chi61_arath : 485.0)' |
| at3g43670.1 | '8.5.1' | copper-containing Symbols:no symbol available \| no full name available \| Chr3:15567144-15569734 FORWARD LENGTH=687 ) &' |
| at1g78850.1 | '35.1' | Symbols:no symbol available \| no full name available \| Chr1:29642072-29643397 REVERSE LENGTH=441 ) & EP1-like glycoprotein 3 OS=Arabidopsis thaliana (sp\|q9zva4\|ep1l3_arath : 893.0)' |
| at5g20080.1 | '50.1.6' | NADH-cytochrome Symbols:no symbol available \| no full name available \| Chr5:6782708-6786360 FORWARD LENGTH=328 ) &' |
| at4g20830.1 | '50.1.1' | Berberine Symbols:AtBBE20,AtBBE19 \| \| Chr4:11155486-11157577 FORWARD LENGTH=570 ) &' |
| at5g44380.1 | '50.1.1' | Berberine Symbols:AtBBE28 \| \| Chr5:17878873-17881369 REVERSE LENGTH=541 ) &' |
| at5g50950.1 | '2.3.7' | fumarase Symbols:FUM2 \| FUMARASE 2 \| Chr5:20729687-20733476 FORWARD LENGTH=510 ) &' |

| | | |
|---|---|---|
| at3g04120.1 | '2.1.1.4.1' | NAD-dependent Symbols:GAPC1,GAPC,GAPC-1 \| GLYCERALDEHYDE-3-PHOSPHATE DEHYDROGENASE C SUBUNIT,glyceraldehyde-3-phosphate dehydrogenase C subunit 1 \| Chr3:1081077-1083131 FORWARD LENGTH=338 ) &' |
| at4g16260.1 | '50.3.2' | Probable Symbols:no symbol available \| no full name available \| Chr4:9200180-9201441 REVERSE LENGTH=344 ) &' |
| at3g13790.1 | '3.1.4.1.1' | acid Symbols:ATBFRUCT1,CWI1,AtCWI1,ATCWINV1 \| ARABIDOPSIS THALIANA CELL WALL INVERTASE 1 \| Chr3:4533084-4535831 REVERSE LENGTH=584 ) &' |
| at3g19390.1 | '50.3.4' | Probable Symbols:no symbol available \| no full name available \| Chr3:6723024-6724768 FORWARD LENGTH=452 ) &' |
| at3g12580.1 | '19.1.5.1' | chaperone Symbols:HSP70,ATHSP70 \| ARABIDOPSIS HEAT SHOCK PROTEIN 70,heat shock protein 70 \| Chr3:3991487-3993689 REVERSE LENGTH=650 ) &' |
| at5g42020.1 | '35.1' | Symbols:BIP2,BIP \| luminal binding protein \| Chr5:16807697-16810480 REVERSE LENGTH=668 ) & Mediator of RNA polymerase II transcription subunit 37f OS=Arabidopsis thaliana (sp\|q39043\|md37f_arath : 1157.0)' |
| at3g16430.1 | '35.1' | Symbols:JAL31,PBPII \| jacalin-related lectin 31 \| Chr3:5581830-5582959 FORWARD LENGTH=296 ) & PYK10-binding protein 2 OS=Arabidopsis thaliana (sp\|o04313\|jal31_arath : 554.0)' |
| at2g22170.1 | '35.1' | Symbols:PLAT2 \| PLAT domain protein 2 \| Chr2:9427010-9427742 REVERSE LENGTH=183 ) & PLAT domain-containing protein 2 OS=Arabidopsis thaliana (sp\|q9sie7\|plat2_arath : 376.0)' |
| at3g16460.1 | '35.1' | Symbols:JAL34 \| jacalin-related lectin 34 \| Chr3:5593029-5595522 FORWARD LENGTH=705 ) & Jacalin-related lectin 34 OS=Arabidopsis thaliana (sp\|o04310\|jal34_arath : 816.0)' |
| at3g48890.1 | '11.3.2.1.4' | brassinosteroid Symbols:MSBP2,MAPR3,ATMAPR3,ATMP2 \| ARABIDOPSIS THALIANA MEMBRANE-ASSOCIATED PROGESTERONE BINDING PROTEIN 3,membrane-associated progesterone binding protein 3,MEMBRANE STEROID BINDING PROTEIN 2 \| Chr3:18129669-18131353 FORWARD LENGTH=233 ) &' |
| at4g19410.1 | '21.3.5.4' | pectin Symbols:PAE7 \| pectin acetylesterase 7 \| Chr4:10582188-10584766 REVERSE LENGTH=391 ) &' |
| at4g30170.1 | '35.1' | Symbols:no symbol available \| no full name available \| Chr4:14762922-14764482 FORWARD LENGTH=325 ) & Peroxidase 45 OS=Arabidopsis thaliana (sp\|q96522\|per45_arath : 619.0)' |
| at1g05240.1 | '35.1' | Symbols:no symbol available \| no full name available \| Chr1:1521202-1522447 FORWARD LENGTH=325 ) & Peroxidase 2 OS=Arabidopsis thaliana (sp\|q67z07\|per2_arath : 598.0)' |
| at2g37130.1 | '35.1' | Symbols:no symbol available \| no full name available \| Chr2:15598225-15600004 REVERSE LENGTH=327 ) & Peroxidase 21 OS=Arabidopsis thaliana (sp\|q42580\|per21_arath : 641.0)' |
| at3g01190.1 | '35.1' | Symbols:no symbol available \| no full name available \| Chr3:67236-68477 REVERSE LENGTH=321 ) & Peroxidase 27 OS=Arabidopsis thaliana (sp\|q43735\|per27_arath : 623.0)' |
| at5g17820.1 | '35.1' | Symbols:no symbol available \| no full name available \| Chr5:5888195-5890101 REVERSE LENGTH=313 ) & Peroxidase 57 OS=Arabidopsis thaliana (sp\|q43729\|per57_arath : 606.0)' |
| at1g79550.1 | '2.1.1.5' | phosphoglycerate Symbols:PGK \| phosphoglycerate kinase \| Chr1:29924347-29926295 REVERSE LENGTH=401 ) &' |

| | | |
|---|---|---|
| at4g20260.4 | '35.1' | Symbols:ATPCAP1,PCAP1,MDP25 \| ARABIDOPSIS THALIANA PLASMA-MEMBRANE ASSOCIATED CATION-BINDING PROTEIN 1,microtubule-destabilizing protein 25,plasma-membrane associated cation-binding protein 1 \| Chr4:10941593-10943227 FORWARD LENGTH=226 ) & Plasma membrane-associated cation-binding protein 1 OS=Arabidopsis thaliana (sp\|q96262\|pcap1_arath : 172.0)' |
| at2g47540.1 | '21.4.1.2.1' | cell Symbols:no symbol available \| no full name available \| Chr2:19505901-19506504 FORWARD LENGTH=173 ) &' |
| at5g05500.1 | '35.2' | Symbols:PRPL1,AtPRPL1,MOP10 \| proline-rich protein-like 1 \| Chr5:1629716-1630267 FORWARD LENGTH=183 )' |
| at4g14060.1 | '35.1' | Symbols:no symbol available \| no full name available \| Chr4:8107405-8108158 FORWARD LENGTH=151 ) & MLP-like protein 328 OS=Arabidopsis thaliana (sp\|q9zvf3\|ml328_arath : 281.0)' |
| at4g23680.1 | '35.1' | Symbols:no symbol available \| no full name available \| Chr4:12336416-12337417 REVERSE LENGTH=151 ) & MLP-like protein 328 OS=Arabidopsis thaliana (sp\|q9zvf3\|ml328_arath : 238.0)' |
| at2g19760.1 | '20.2.2.3' | profilin Symbols:PFN1,PRF1 \| profilin 1,PROFILIN 1 \| Chr2:8517074-8518067 REVERSE LENGTH=131 ) &' |
| at4g02270.1 | '21.4.1.2.1' | cell Symbols:RHS13 \| root hair specific 13 \| Chr4:992383-992996 REVERSE LENGTH=165 ) &' |
| at4g26220.1 | '21.6.1.4' | caffeoyl-CoA Symbols:CCoAOMT7 \| caffeoyl coenzyme A ester O-methyltransferase 7 \| Chr4:13284179-13285146 FORWARD LENGTH=232 ) &' |
| at1g58270.1 | '35.2' | Symbols:ZW9 \| \| Chr1:21612394-21614089 REVERSE LENGTH=396 )' |
| at1g45201.1 | '5.7.1.2.3' | lipase Symbols:TLL1,ATTLL1 \| ARABIDOPSIS THALIANA TRIACYLGLYCEROL LIPASE-LIKE 1,triacylglycerol lipase-like 1 \| Chr1:17123889-17128462 FORWARD LENGTH=479 ) &' |
| at3g55440.1 | '2.1.1.3' | triosephosphate Symbols:ATCTIMC,CYTOTPI,TPI \| CYTOSOLIC ISOFORM TRIOSE PHOSPHATE ISOMERASE,triosephosphate isomerase,CYTOSOLIC TRIOSE PHOSPHATE ISOMERASE \| Chr3:20553794-20556078 FORWARD LENGTH=254 ) &' |
| at3g55440.1 | '3.1.2.1' | cytosolic Symbols:ATCTIMC,CYTOTPI,TPI \| CYTOSOLIC ISOFORM TRIOSE PHOSPHATE ISOMERASE,triosephosphate isomerase,CYTOSOLIC TRIOSE PHOSPHATE ISOMERASE \| Chr3:20553794-20556078 FORWARD LENGTH=254 ) &' |
| at4g30270.1 | '50.2.4' | Xyloglucan Symbols:MERI5B,XTH24,MERI-5,SEN4 \| xyloglucan endotransglucosylase/hydrolase 24,SENESCENCE 4,MERISTEM 5,meristem-5 \| Chr4:14819445-14820448 REVERSE LENGTH=269 ) &' |

| The Arabidopsis Information Resources (www.arabidopsis.org) | | |
|---|---|---|
| Accession | Gene Model Description | Primary Gene Symbol |
| ATCG00120.1 | Encodes the ATPase alpha subunit, which is a subunit of ATP synthase and part of the | ATP SYNTHASE SUBUNIT ALPHA (ATPA) |
| ATCG00480.1 | chloroplast-encoded gene for beta subunit of ATP synthase | ATP SYNTHASE SUBUNIT BETA (PB) |
| AT4G04640.1 | One of two genes (with ATPC2) encoding the gamma subunit of Arabidopsis chloropl | (ATPC1) |
| AT1G29910.1 | member of Chlorophyll a/b-binding protein family | CHLOROPHYLL A/B BINDING PROTEIN 3 (CAB3) |
| AT1G20340.1 | recombination and DNA-damage resistance protein (DRT112) One of two Arabidops | DNA-DAMAGE-REPAIR/TOLERATION PROTEIN 112 (DRT112) |
| AT5G66190.1 | Encodes a leaf-type ferredoxin:NADP(H) oxidoreductase. It is present in both chloro | FERREDOXIN-NADP(+)-OXIDOREDUCTASE 1 (FNR1) |
| AT4G38970.1 | Protein is tyrosine-phosphorylated and its phosphorylation state is modulated in res | FRUCTOSE-BISPHOSPHATE ALDOLASE 2 (FBA2) |
| AT4G38970.1 | Protein is tyrosine-phosphorylated and its phosphorylation state is modulated in res | FRUCTOSE-BISPHOSPHATE ALDOLASE 2 (FBA2) |
| AT5G42270.1 | VAR1 contains a conserved motif for ATPase and a metalloprotease characteristic to | VARIEGATED 1 (VAR1) |
| AT3G09260.1 | Encodes beta-glucosidase.The major constituent of ER bodies. One of the most abun | (PYK10) |
| AT4G10340.1 | photosystem II encoding the light-harvesting chlorophyll a/b binding protein CP26 of | LIGHT HARVESTING COMPLEX OF PHOTOSYSTEM II 5 (LHCB5) |
| AT5G54270.1 | Lhcb3 protein is a component of the main light harvesting chlorophyll a/b-protein co | LIGHT-HARVESTING CHLOROPHYLL B-BINDING PROTEIN 3 (LHCB3) |
| AT2G34430.1 | Photosystem II type I chlorophyll a/b-binding protein The mRNA is cell-to-cell r | LIGHT-HARVESTING CHLOROPHYLL-PROTEIN COMPLEX II SUBUNIT B1 (LHB1B1) |
| AT2G24940.1 | membrane-associated progesterone binding protein 2;(source:Araport11) | MEMBRANE-ASSOCIATED PROGESTERONE BINDING PROTEIN 2 (MAPR2) |
| AT4G22890.1 | Encodes PGRL1A, a transmembrane protein present in thylakoids. PGRL1A has a higl | (PGR5-LIKE A) |
| ATCG00540.1 | Encodes cytochrome f apoprotein; involved in photosynthetic electron transport cha | PHOTOSYNTHETIC ELECTRON TRANSFER A (PETA) |
| AT4G03280.1 | Encodes the Rieske FeS center of cytochrome b6f complex. Gene is expressed in shoo | PHOTOSYNTHETIC ELECTRON TRANSFER C (PETC) |
| AT4G28750.1 | mutant has Decreased effective quantum yield of photosystem II; Pale green plants; | PSA E1 KNOCKOUT (PSAE-1) |
| AT2G20260.1 | Encodes subunit E of photosystem I. The mRNA is cell-to-cell mobile. | PHOTOSYSTEM I SUBUNIT E-2 (PSAE-2) |
| AT1G31330.1 | Encodes subunit F of photosystem I. | PHOTOSYSTEM I SUBUNIT F (PSAF) |
| AT1G52230.1 | Phosphorylation of this protein is dependent on calcium. The mRNA is cell-to-cell mo | PHOTOSYSTEM I SUBUNIT H2 (PSAH2) |
| ATCG00340.1 | Encodes the D1 subunit of photosystem I reaction center. | (PSAB) |
| AT1G03600.1 | PSB27 is a chloroplast lumen localized protein that is involved in adaptation to chang | (PSB27) |
| AT1G06680.1 | Encodes a 23 kD extrinsic protein that is part of photosystem II and participates in th | PHOTOSYSTEM II SUBUNIT P-1 (PSBP-1) |
| AT1G79040.1 | Encodes for the 10 kDa PsbR subunit of photosystem II (PSII). This subunit appears to | PHOTOSYSTEM II SUBUNIT R (PSBR) |
| AT2G05100.1 | Lhcb2.1 protein encoding a subunit of the light harvesting complex II. Member of a g | PHOTOSYSTEM II LIGHT HARVESTING COMPLEX GENE 2.1 (LHCB2.1) |
| AT2G20890.1 | Chloroplast-localized Thylakoid formation1 gene product involved in vesicle-mediate | PHOTOSYSTEM II REACTION CENTER PSB29 PROTEIN (PSB29) |
| AT2G30790.1 | Encodes a 23 kD extrinsic protein that is part of photosystem II and participates in th | PHOTOSYSTEM II SUBUNIT P-2 (PSBP-2) |
| AT3G50820.1 | Encodes a protein which is an extrinsic subunit of photosystem II and which has beer | PHOTOSYSTEM II SUBUNIT O-2 (PSBO2) |
| AT4G05180.1 | Encodes the PsbQ subunit of the oxygen evolving complex of photosystem II. | PHOTOSYSTEM II SUBUNIT Q-2 (PSBQ-2) |
| AT4G21280.2 | Encodes the PsbQ subunit of the oxygen evolving complex of photosystem II. | PHOTOSYSTEM II SUBUNIT QA (PSBQA) |
| ATCG00020.1 | Encodes chlorophyll binding protein D1, a part of the photosystem II reaction center | PHOTOSYSTEM II REACTION CENTER PROTEIN A (PSBA) |
| ATCG00270.1 | PSII D2 protein | PHOTOSYSTEM II REACTION CENTER PROTEIN D (PSBD) |
| ATCG00280.1 | chloroplast gene encoding a CP43 subunit of the photosystem II reaction center. pro | PHOTOSYSTEM II REACTION CENTER PROTEIN C (PSBC) |
| ATCG00680.1 | encodes for CP47, subunit of the photosystem II reaction center. | PHOTOSYSTEM II REACTION CENTER PROTEIN B (PSBB) |
| AT5G07020.1 | Encodes an integral thylakoid membrane protein that interacts with PSII core comple | MAINTENANCE OF PSII UNDER HIGH LIGHT 1 (MPH1) |
| AT1G74470.1 | Encodes for a multifunctional protein with geranylgeranyl reductase activity shown t | 0 |
| ATMG00280.1 | Ribulose bisphosphate carboxylase large chain, catalytic domain-containing protein;( | (ORF110A) |
| AT1G67090.1 | Encodes a member of the Rubisco small subunit (RBCS) multigene family: RBCS1A (A | RIBULOSE BISPHOSPHATE CARBOXYLASE SMALL CHAIN 1A (RBCS1A) |
| ATCG00490.1 | large subunit of RUBISCO. Protein is tyrosine-phosphorylated and its phosphorylatior | (RBCL) |

| | | |
|---|---|---|
| AT1G71500.1 | Encodes PSB33, a protein conserved in the plastid lineage. PSB33 is associated with t | PHOTOSYSTEM B PROTEIN 33 (PSB33) |
| AT2G39730.1 | Rubisco activase, a nuclear-encoded chloroplast protein that consists of two isoform | RUBISCO ACTIVASE (RCA) |
| AT1G54780.1 | Encodes a thylakoid lumen protein regulating photosystem II repair cycle. Has acid p | THYLAKOID LUMEN PROTEIN 18.3 (TLP18.3) |
| AT5G09810.1 | Member of Actin gene family.Mutants are defective in germination and root growth. | ACTIN 7 (ACT7) |
| AT1G28290.1 | Encodes an atypical arabinogalactan protein that is localized to the plasma membrar | ARABINOGALACTAN PROTEIN 31 (AGP31) |
| AT5G08680.1 | Encodes the mitochondrial ATP synthase beta-subunit. This subunit is encoded by a | 0 |
| AT1G45130.1 | beta-galactosidase 5;(source:Araport11) | BETA-GALACTOSIDASE 5 (BGAL5) |
| AT2G43610.1 | Chitinase family protein;(source:Araport11) | 0 |
| AT3G43670.1 | Copper amine oxidase family protein;(source:Araport11) | COPPER AMINE OXIDASE GAMMA 2 (CUAOγ2) |
| AT1G78850.1 | curculin-like (mannose-binding) lectin family protein, low similarity to ser/thr proteir | MANNOSE BINDING LECTIN1 (MBL1) |
| AT5G20080.1 | FAD/NAD(P)-binding oxidoreductase;(source:Araport11) | 0 |
| AT4G20830.1 | Encodes an oligogalacturonide oxidase that inactivates the elicitor-active oligogalact | (ATBBE19) |
| AT5G44380.1 | FAD-binding Berberine family protein;(source:Araport11) | (ATBBE28) |
| AT5G50950.1 | Encodes a fumarase enzyme initially shown to be in the mitochondria through prote | FUMARASE 2 (FUM2) |
| AT3G04120.1 | encodes cytosolic GADPH (C subunit) involved in the glycolytic pathway but also inte | GLYCERALDEHYDE-3-PHOSPHATE DEHYDROGENASE C SUBUNIT 1 (GAPC1) |
| AT4G16260.1 | Encodes a putative beta-1,3-endoglucanase that interacts with the 30C02 cyst nemat | 0 |
| AT3G13790.1 | Encodes a protein with invertase activity. | (ATBFRUCT1) |
| AT3G19390.1 | Granulin repeat cysteine protease family protein;(source:Araport11) | 0 |
| AT3G12580.1 | heat shock protein 70;(source:Araport11) | HEAT SHOCK PROTEIN 70 (HSP70) |
| AT5G42020.1 | Luminal binding protein (BiP2) involved in polar nuclei fusion during proliferation of | (BIP2) |
| AT3G16430.1 | Encodes a protein that increases the beta-glucosidase activities of three scopolin glu | JACALIN-RELATED LECTIN 31 (JAL31) |
| AT2G22170.1 | Lipase/lipooxygenase, PLAT/LH2 family protein;(source:Araport11) | PLAT DOMAIN PROTEIN 2 (PLAT2) |
| AT3G16460.1 | Mannose-binding protein | JACALIN-RELATED LECTIN 34 (JAL34) |
| AT3G48890.1 | putative progesterone-binding protein homolog (Atmp2) mRNA, The mRNA is cell-to | MEMBRANE-ASSOCIATED PROGESTERONE BINDING PROTEIN 3 (MAPR3) |
| AT4G19410.1 | Pectinacetylesterase family protein;(source:Araport11) | PECTIN ACETYLESTERASE 7 (PAE7) |
| AT4G30170.1 | Peroxidase family protein;(source:Araport11) | 0 |
| AT1G05240.1 | Peroxidase superfamily protein;(source:Araport11) | PATHOGENESIS RELATED 9 (PR9) |
| AT2G37130.1 | Peroxidase superfamily protein;(source:Araport11) | 0 |
| AT3G01190.1 | Peroxidase superfamily protein;(source:Araport11) | 0 |
| AT5G17820.1 | Peroxidase superfamily protein;(source:Araport11) | PEROXIDASE 57 (PER57) |
| AT1G79550.1 | Encodes cytosolic phosphoglycerate kinase (PGK3). Expression studies in PGK mutan | PHOSPHOGLYCERATE KINASE (PGK) |
| AT4G20260.4 | Encodes a Ca2+ and Cu2+ binding protein. N-terminal myristylation on glycine 2 appe | PLASMA-MEMBRANE ASSOCIATED CATION-BINDING PROTEIN 1 (PCAP1) |
| AT2G47540.1 | Pollen Ole e 1 allergen and extensin family protein;(source:Araport11) | 0 |
| AT5G05500.1 | Encodes Proline-rich protein-like PRPL1, controls elongation of root hairs. | (MOP10) |
| AT4G14060.1 | Polyketide cyclase/dehydrase and lipid transport superfamily protein;(source:Arapor | 0 |
| AT4G23680.1 | Polyketide cyclase/dehydrase and lipid transport superfamily protein;(source:Arapor | 0 |
| AT2G19760.1 | first member of the Arabidopsis profilin multigene family, expressed in all organs of A | PROFILIN 1 (PRF1) |
| AT4G02270.1 | root hair specific 13;(source:Araport11) | ROOT HAIR SPECIFIC 13 (RHS13) |
| AT4G26220.1 | Encodes a caffeoyl-coenzyme A O-methyltransferase (CCoAOMT)-like protein with a | CAFFEOYL COENZYME A ESTER O-METHYLTRANSFERASE 7 (CCOAOMT7) |
| AT1G58270.1 | ZW9 mRNA, complete cds The mRNA is cell-to-cell mobile. | (ZW9) |
| AT1G45201.1 | Target of AtGRP7 regulation. | TRIACYLGLYCEROL LIPASE-LIKE 1 (TLL1) |
| AT3G55440.1 | Encodes triosephosphate isomerase. | TRIOSEPHOSPHATE ISOMERASE (TPI) |

| | | |
|---|---|---|
| AT3G55440.1 | Encodes triosephosphate isomerase. | TRIOSEPHOSPHATE ISOMERASE (TPI) |
| AT4G30270.1 | encodes a protein similar to endo xyloglucan transferase in sequence. It is also very s | XYLOGLUCAN ENDOTRANSGLUCOSYLASE/HYDROLASE 24 (XTH24) |

| LEAVES | | | Datasets | | | | | | | | | | |
|---|---|---|---|---|---|---|---|---|---|---|---|---|---|
| gene | Description | Master Protein Accessions | ChlorBN | Chlo_10545 | Chlo3516 | PXD010730 | TMAR | CAU | CLLF | Ros | LFD | LFP | LFPT |
| ATPA | ATP synthase subunit alpha | ATCG00120.1 | x | | | | | | | | | | |
| PB | ATP synthase subunit beta | ATCG00480.1 | x | | | | | | | | | | |
| ATPC1 | ATPase, F1 complex, gamma subunit | AT4G04640.1 | x | | | | | | | | | | |
| CAB3 | chlorophyll A/B binding protein 3 | AT1G29910.1 | x | | | | | | | | | | |
| DRT112 | Cupredoxin superfamily protein | AT1G20340.1 | | | | | | x | | x | | | |
| FNR1 | ferredoxin-NADP(+)-oxidoreductase 1 | AT5G66190.1 | x | | | | | | | | | | |
| FBA2 | fructose-bisphosphate aldolase 2 | AT4G38970.1 | x | | | | | | | | | | |
| VAR1 | FtsH extracellular protease family | AT5G42270.1 | x | | | | | | | | | | |
| PYK10 | Glycosyl hydrolase superfamily protein | AT3G09260.1 | x | | | | | | | | | | |
| LHCB5 | light harvesting complex of | AT4G10340.1 | x | | | | | | | | | | |
| LHCB3 | light-harvesting chlorophyll B-binding | AT5G54270.1 | x | | | | | | | | | | |
| LHB1B1 | light-harvesting chlorophyll-protein | AT2G34430.1 | x | | | x | | | | | x | | |
| MAPR2 | membrane-associated progesterone | AT2G24940.1 | | | | | | | | x | | | |
| PGR5-LIKE A | PGR5-LIKE A | AT4G22890.5 | x | | | | | | | | | | |
| PETA | photosynthetic electron transfer A | ATCG00540.1 | x | | | | | | | | | | |
| PETC | photosynthetic electron transfer C | AT4G03280.1 | x | | | | | | | | | | |
| PSAE-1 | Photosystem I reaction centre subunit | AT4G28750.1 | x | | | | | | | | | | |
| PSAE-2 | photosystem I subunit E-2 | AT2G20260.1 | x | | | | | | | | | | |
| PSAF | photosystem I subunit F | AT1G31330.1 | x | | | | | | | x | | | |
| PSAH2 | photosystem I subunit H2 | AT1G52230.1 | x | | | | | | | | | | |
| PSAB | Photosystem I, PsaA/PsaB protein | ATCG00340.1 | x | | | | | | | | | | |
| PSB27 | photosystem II family protein | AT1G03600.1 | x | | | | | | | | | | |
| PSBP-1 | photosystem II subunit P-1 | AT1G06680.1 | x | | | x | | | | | | | |
| PSBR | photosystem II subunit R | AT1G79040.1 | x | | | | | | | | | | |
| LHCB2.1 | photosystem II light harvesting | AT2G05100.1 | x | | | | | | | | | | |
| PSB29 | photosystem II reaction center PSB29 | AT2G20890.1 | x | | | | | | | | | | |
| PSBP-2 | photosystem II subunit P-2 | AT2G30790.1 | x | | | | | | | | | | |
| PSBO2 | photosystem II subunit O-2 | AT3G50820.1 | x | | x | | | | | | | | |
| PSBQ-2 | photosystem II subunit Q-2 | AT4G05180.1 | x | | | | | | | | | | |
| PSBQA | photosystem II subunit QA | AT4G21280.1 | x | | | | | | | | | | |
| PSBA | photosystem II reaction center | ATCG00020.1 | x | | | | x | | | | | | |
| PSBD | photosystem II reaction center | ATCG00270.1 | x | | | | | | | | | | |
| PSBC | photosystem II reaction center | ATCG00280.1 | | | | | | | | | x | | |
| PSBB | photosystem II reaction center | ATCG00680.1 | x | | | | x | x | | x | | | |
| MPH1 | proline-rich family protein | AT5G07020.1 | x | | | | | | | | | | |
| 0 | Pyridine nucleotide-disulphide | AT1G74470.1 | x | | | | | | | | | | |

| | | | | | | | | | | | | | |
|---|---|---|---|---|---|---|---|---|---|---|---|---|---|
| ORF110A | Ribulose bisphosphate carboxylase large chain, catalytic domain | ATMG00280.1 | | | | | | | x | | x | x | x |
| RBCS1A | ribulose bisphosphate carboxylase | AT1G67090.1 | x | | | | | x | | x | | | |
| RBCL | ribulose-bisphosphate carboxylases | ATCG00490.1 | x | x | | | | x | | | | | |
| PSB33 | Rieske (2Fe-2S) domain-containing | AT1G71500.1 | x | | | | | | | | | | |
| RCA | rubisco activase | AT2G39730.1 | x | | | | | | | | | | |
| TLP18.3 | thylakoid lumen 18.3 kDa protein | AT1G54780.1 | x | | | | | | | | | | |

| ROOT | | | Datasets | | | | |
| --- | --- | --- | --- | --- | --- | --- | --- |
| **gene** | **Description** | **Master Protein Accessions** | **Root** | **RT** | **RTTP** | **RTUZ** | **RTTOF** |
| ACT7 | actin 7 | AT5G09810.1 | x | | | | |
| AGP31 | arabinogalactan protein 31 | AT1G28290.1 | | x | | | |
| 0 | ATP synthase alpha/beta family protein | AT5G08680.1 | x | | | | |
| BGAL5 | beta-galactosidase 5 | AT1G45130.1 | | x | x | x | |
| 0 | Chitinase family protein | AT2G43610.1 | x | | | | |
| CUAOγ2 | Copper amine oxidase family protein | AT3G43670.1 | | x | | x | |
| MBL1 | D-mannose binding lectin protein | AT1G78850.1 | x | x | x | | |
| 0 | FAD/NAD(P)-binding oxidoreductase | AT5G20080.1 | x | | | | |
| ATBBE19 | FAD-binding Berberine family protein | AT4G20830.1 | | | x | | |
| ATBBE28 | FAD-binding Berberine family protein | AT5G44380.1 | | | | x | |
| FUM2 | FUMARASE 2 | AT5G50950.1 | x | | | | |
| GAPC1 | glyceraldehyde-3-phosphate dehydrogenase C sub 1 | AT3G04120.1 | | x | x | x | |
| 0 | Glycosyl hydrolase superfamily protein | AT4G16260.1 | x | | | | |
| ATBFRUCT1 | Glycosyl hydrolases family 32 protein | AT3G13790.1 | | x | | x | |
| 0 | Granulin repeat cysteine protease family protein | AT3G19390.1 | x | | | | |
| HSP70 | heat shock protein 70 | AT3G12580.1 | x | | | | |
| BIP2 | Heat shock protein 70 (Hsp 70) family protein | AT5G42020.1 | x | | | | |
| JAL31 | jacalin-related lectin 31 | AT3G16430.1 | x | | | | |
| PLAT2 | Lipase/lipooxygenase, PLAT/LH2 family protein | AT2G22170.1 | x | | | | |
| JAL34 | Mannose-binding lectin superfamily protein | AT3G16460.1 | x | | | | |
| MAPR3 | membrane-associated progesterone binding protein 3 | AT3G48890.1 | x | | | | |
| PAE7 | Pectinacetylesterase family protein | AT4G19410.1 | | x | | | |
| 0 | Peroxidase family protein | AT4G30170.1 | x | x | | x | |
| PR9 | Peroxidase superfamily protein | AT1G05240.1 | | x | x | x | |
| 0 | Peroxidase superfamily protein | AT2G37130.1 | | x | | x | |
| 0 | Peroxidase superfamily protein | AT3G01190.1 | | x | | x | |
| PER57 | Peroxidase superfamily protein | AT5G17820.1 | x | x | | | |
| PGK | phosphoglycerate kinase | AT1G79550.1 | | x | | | |
| PCAP1 | plasma-membrane associated cation-binding protein 1 | AT4G20260.4 | x | | | | |
| 0 | Pollen Ole e 1 allergen and extensin family protein | AT2G47540.1 | x | | | | |
| MOP10 | Pollen Ole e 1 allergen and extensin family protein | AT5G05500.1 | | x | x | x | |
| 0 | Polyketide cyclase/dehydrase and lipid transport superfamily protein | AT4G14060.1 | x | | | | |
| 0 | Polyketide cyclase/dehydrase and lipid transport superfamily protein | AT4G23680.1 | x | | | | |
| PRF1 | profilin 1 | AT2G19760.1 | | | | | x |
| RHS13 | root hair specific 13 | AT4G02270.1 | x | | x | x | |
| CCOAOMT7 | S-adenosyl-L-methionine-dependent methyltransferases superfamily protein | AT4G26220.1 | | | x | | |
| ZW9 | TRAF-like family protein | AT1G58270.1 | x | | | | |

| | | | | |
|---|---|---|---|---|
| TLL1 | triacylglycerol lipase-like 1 | AT1G45201.1 | x | |
| TPI | triosephosphate isomerase | AT3G55440.1 | x | |
| XTH24 | xyloglucan endotransglucosylase/hydrolase 24 | AT4G30270.1 | | x |
