## Supplementary material for "Evidences for a nutritional role of iodine in plants": Table S15

| ROOTS – Biological processes (GO terms enrichment)<br>*OGC: observed gene count; **BGC: background gene count | | | | | |
|---|---|---|---|---|---|
| #term ID | term description | OGC * | BGC** | FDR | matching proteins in your network (labels) |
| GO:0017144 | drug metabolic process | 11 | 626 | 3.24e-07 | AT2G37130,AT2G43610,AT3G01190,AT4G30170,AT5G08680,AT5G17820,FUM2,GAPC1,P1,PGK,TPI |
| GO:0006950 | response to stress | 20 | 2932 | 3.24e-07 | ACT7,AT2G37130,AT2G43610,AT3G01190,AT4G16260,AT4G20830,AT4G30170,AT5G17820,AT5G20080,AT5G44380,ATBFRUCT1,BIP2,FUM2,GAPC1,HSP70,JAL34,P1,PCAP1,PGK,TPI |
| GO:0006979 | response to oxidative stress | 9 | 389 | 7.87e-07 | AT2G37130,AT3G01190,AT4G20830,AT4G30170,AT5G17820,AT5G44380,GAPC1,HSP70,P1 |
| GO:0009636 | response to toxic substance | 8 | 330 | 3.30e-06 | AT2G37130,AT3G01190,AT4G30170,AT5G17820,GAPC1,HSP70,P1,PLAT2 |
| GO:0050896 | response to stimulus | 23 | 5064 | 4.53e-06 | ACT7,AGP31,AT2G37130,AT2G43610,AT3G01190,AT4G16260,AT4G20830,AT4G30170,AT5G08680,AT5G17820,AT5G20080,AT5G44380,ATBFRUCT1,BIP2,FUM2,GAPC1,HSP70,JAL34,P1,PCAP1,PGK,PLAT2,TPI |
| GO:0016999 | antibiotic metabolic process | 6 | 171 | 1.75e-05 | AT2G37130,AT3G01190,AT4G30170,AT5G17820,FUM2,P1 |
| GO:0042743 | hydrogen peroxide metabolic process | 5 | 95 | 1.75e-05 | AT2G37130,AT3G01190,AT4G30170,AT5G17820,P1 |
| GO:0042744 | hydrogen peroxide catabolic process | 5 | 88 | 1.75e-05 | AT2G37130,AT3G01190,AT4G30170,AT5G17820,P1 |
| GO:0098869 | cellular oxidant detoxification | 6 | 179 | 1.75e-05 | AT2G37130,AT3G01190,AT4G30170,AT5G17820,P1,PLAT2 |
| GO:0051186 | cofactor metabolic process | 8 | 531 | 3.29e-05 | AT2G37130,AT3G01190,AT4G30170,AT5G17820,GAPC1,P1,PGK,TPI |
| GO:0042737 | drug catabolic process | 6 | 226 | 3.41e-05 | AT2G37130,AT2G43610,AT3G01190,AT4G30170,AT5G17820,P1 |
| GO:0009664 | plant-type cell wall organization | 5 | 139 | 5.80e-05 | AT2G37130,AT3G01190,AT5G17820,P1,XTH24 |
| GO:0042221 | response to chemical | 15 | 2654 | 6.25e-05 | ACT7,AGP31,AT2G37130,AT3G01190,AT4G30170,AT5G08680,AT5G17820,BIP2,GAPC1,HSP70,P1,PCAP1,PGK,PLAT2,TPI |
| GO:0055114 | oxidation-reduction process | 11 | 1348 | 6.39e-05 | AT2G37130,AT3G01190,AT4G20830,AT4G30170,AT5G17820,AT5G20080,AT5G44380,FUM2,GAPC1,P1,PLAT2 |
| GO:0006754 | ATP biosynthetic process | 4 | 89 | 0.00024 | AT5G08680,GAPC1,PGK,TPI |
| GO:0009168 | purine ribonucleoside monophosphate biosynthetic process | 4 | 110 | 0.00044 | AT5G08680,GAPC1,PGK,TPI |
| GO:0009628 | response to abiotic stimulus | 11 | 1699 | 0.00044 | ACT7,AT4G16260,AT5G20080,ATBFRUCT1,FUM2,GAPC1,HSP70,JAL34,PCAP1,PGK,TPI |
| GO:0005975 | carbohydrate metabolic process | 8 | 856 | 0.00049 | AT2G43610,AT4G16260,ATBFRUCT1,BGAL5,GAPC1,PGK,TPI,XTH24 |
| GO:0010038 | response to metal ion | 6 | 414 | 0.00049 | AT5G08680,BIP2,GAPC1,HSP70,PCAP1,TPI |
| GO:0046034 | ATP metabolic process | 4 | 125 | 0.00057 | AT5G08680,GAPC1,PGK,TPI |
| GO:0071554 | cell wall organization or biogenesis | 7 | 639 | 0.00057 | AT2G37130,AT2G43610,AT3G01190,AT4G19410,AT5G17820,P1,XTH24 |

| | | | | | |
|---|---|---|---|---|---|
| GO:0046686 | response to cadmium ion | 5 | 286 | 0.00079 | AT5G08680,BIP2,GAPC1,HSP70,TPI |
| GO:0009167 | purine ribonucleoside monophosphate metabolic process | 4 | 149 | 0.00084 | AT5G08680,GAPC1,PGK,TPI |
| GO:0009651 | response to salt stress | 6 | 492 | 0.00085 | AT4G16260,AT5G20080,FUM2,GAPC1,PCAP1,TPI |
| GO:0006096 | glycolytic process | 3 | 57 | 0.00094 | GAPC1,PGK,TPI |
| GO:0006757 | ATP generation from ADP | 3 | 57 | 0.00094 | GAPC1,PGK,TPI |
| GO:0009266 | response to temperature stimulus | 6 | 505 | 0.00094 | FUM2,GAPC1,HSP70,JAL34,PCAP1,PGK |
| GO:0042866 | pyruvate biosynthetic process | 3 | 57 | 0.00094 | GAPC1,PGK,TPI |
| GO:0071555 | cell wall organization | 6 | 503 | 0.00094 | AT2G37130,AT3G01190,AT4G19410,AT5G17820,P1,XTH24 |
| GO:0009166 | nucleotide catabolic process | 3 | 61 | 0.00097 | GAPC1,PGK,TPI |
| GO:0044248 | cellular catabolic process | 9 | 1345 | 0.00097 | AT2G37130,AT2G43610,AT3G01190,AT4G30170,AT5G17820,GAPC1,P1,PGK,TPI |
| GO:0019359 | nicotinamide nucleotide biosynthetic process | 3 | 67 | 0.0011 | GAPC1,PGK,TPI |
| GO:0006090 | pyruvate metabolic process | 3 | 74 | 0.0014 | GAPC1,PGK,TPI |
| GO:0006094 | gluconeogenesis | 2 | 18 | 0.0024 | GAPC1,TPI |
| GO:0032272 | negative regulation of protein polymerization | 2 | 18 | 0.0024 | PCAP1,PRF1 |
| GO:0046496 | nicotinamide nucleotide metabolic process | 3 | 96 | 0.0024 | GAPC1,PGK,TPI |
| GO:0051494 | negative regulation of cytoskeleton organization | 2 | 19 | 0.0025 | PCAP1,PRF1 |
| GO:1902904 | negative regulation of supramolecular fiber organization | 2 | 19 | 0.0025 | PCAP1,PRF1 |
| GO:0016052 | carbohydrate catabolic process | 4 | 244 | 0.0026 | AT2G43610,GAPC1,PGK,TPI |
| GO:0009743 | response to carbohydrate | 3 | 123 | 0.0044 | GAPC1,PCAP1,PGK |
| GO:0051707 | response to other organism | 7 | 1079 | 0.0047 | AT2G37130,AT4G16260,AT4G20830,ATBFRUCT1,HSP70,PCAP1,PGK |

| | | | | | |
|---|---|---|---|---|---|
| GO:0002237 | response to molecule of bacterial origin | 2 | 33 | 0.0061 | PCAP1,PGK |
| GO:0008152 | metabolic process | 24 | 9671 | 0.0063 | AT2G37130,AT2G43610,AT3G01190,AT3G19390,AT4G16260,AT4G20830,AT4G26220,AT4G30170,AT5G08680,AT5G17820,AT5G20080,AT5G44380,ATBFRUCT1,BGAL5,BIP2,FUM2,GAPC1,HSP70,P1,PCAP1,PGK,PLAT2,TPI,XTH24 |
| GO:0006006 | glucose metabolic process | 2 | 38 | 0.0076 | GAPC1,TPI |
| GO:0006091 | generation of precursor metabolites and energy | 4 | 360 | 0.0091 | FUM2,GAPC1,PGK,TPI |
| GO:0070887 | cellular response to chemical stimulus | 7 | 1245 | 0.0094 | AT2G37130,AT3G01190,AT4G30170,AT5G17820,P1,PCAP1,PLAT2 |
| GO:0009408 | response to heat | 3 | 184 | 0.0114 | GAPC1,HSP70,PGK |
| GO:0007010 | cytoskeleton organization | 3 | 192 | 0.0126 | ACT7,PCAP1,PRF1 |
| GO:1901135 | carbohydrate derivative metabolic process | 5 | 701 | 0.0147 | AT2G43610,AT5G08680,GAPC1,PGK,TPI |
| GO:0009808 | lignin metabolic process | 2 | 59 | 0.0151 | AT4G26220,P1 |
| GO:0009735 | response to cytokinin | 3 | 212 | 0.0158 | BIP2,PCAP1,TPI |
| GO:0042542 | response to hydrogen peroxide | 2 | 63 | 0.0164 | GAPC1,HSP70 |
| GO:0016043 | cellular component organization | 9 | 2271 | 0.0189 | ACT7,AT2G37130,AT3G01190,AT4G19410,AT5G17820,P1,PCAP1,PRF1,XTH24 |
| GO:0050832 | defense response to fungus | 4 | 478 | 0.0203 | AT2G37130,AT4G16260,AT4G20830,ATBFRUCT1 |
| GO:0098542 | defense response to other organism | 5 | 847 | 0.0283 | AT2G37130,AT4G16260,AT4G20830,ATBFRUCT1,PCAP1 |
| GO:0006952 | defense response | 6 | 1277 | 0.0365 | AT2G37130,AT2G43610,AT4G16260,AT4G20830,ATBFRUCT1,PCAP1 |
| GO:0097435 | supramolecular fiber organization | 2 | 106 | 0.0388 | PCAP1,PRF1 |
| GO:0010033 | response to organic substance | 7 | 1786 | 0.0485 | ACT7,AGP31,BIP2,GAPC1,PCAP1,PGK,TPI |
| GO:0009409 | response to cold | 3 | 347 | 0.0494 | FUM2,JAL34,PCAP1 |

ROOTS – Molecular functions (GO terms enrichment)

*OGC: observed gene count; **BGC: background gene count

| #term ID | term description | OGC * | BGC** | FDR | matching proteins in your network (labels) |
|---|---|---|---|---|---|
| GO:0005507 | copper ion binding | 7 | 157 | 5.13e-07 | AT5G08680,AT5G20080,GAPC1,JAL31,JAL34,PCAP1,TPI |
| GO:0004601 | peroxidase activity | 6 | 128 | 2.75e-06 | AT2G37130,AT3G01190,AT4G30170,AT5G17820,P1,PLAT2 |

| | | | | | |
|---|---|---|---|---|---|
| GO:0020037 | heme binding | 7 | 267 | 4.37e-06 | AT2G37130,AT3G01190,AT4G30170,AT5G17820,MAPR3,P1,PLAT2 |
| GO:0048037 | cofactor binding | 10 | 860 | 7.22e-06 | AT2G37130,AT3G01190,AT4G20830,AT4G30170,AT5G17820,AT5G44380,GAPC1,MAPR3,P1,PLAT2 |
| GO:0016491 | oxidoreductase activity | 10 | 1201 | 0.00012 | AT2G37130,AT3G01190,AT4G20830,AT4G30170,AT5G17820,AT5G20080,AT5G44380,GAPC1,P1,PLAT2 |
| GO:0043167 | ion binding | 19 | 5070 | 0.00043 | ACT7,AT2G37130,AT3G01190,AT4G20830,AT4G26220,AT4G30170,AT5G08680,AT5G17820,AT5G20080,AT5G44380,BIP2,GAPC1,HSP70,JAL31,JAL34,P1,PCAP1,PGK,TPI |
| GO:0004553 | hydrolase activity, hydrolyzing O-glycosyl compounds | 5 | 258 | 0.00053 | AT2G43610,AT4G16260,ATBFRUCT1,BGAL5,XTH24 |
| GO:0005488 | binding | 25 | 8611 | 0.00063 | ACT7,AT1G78850,AT2G37130,AT2G43610,AT3G01190,AT4G16260,AT4G20830,AT4G26220,AT4G30170,AT5G08680,AT5G17820,AT5G20080,AT5G44380,BIP2,GAPC1,HSP70,JAL31,JAL34,MAPR3,P1,PCAP1,PGK,PLAT2,PRF1,TPI |
| GO:0003824 | catalytic activity | 22 | 7239 | 0.0013 | AT2G37130,AT2G43610,AT3G01190,AT3G19390,AT4G16260,AT4G19410,AT4G20830,AT4G26220,AT4G30170,AT5G08680,AT5G17820,AT5G20080,AT5G44380,ATBFRUCT1,BGAL5,FUM2,GAPC1,P1,PGK,PLAT2,TPI,XTH24 |
| GO:0046872 | metal ion binding | 13 | 2940 | 0.0018 | AT2G37130,AT3G01190,AT4G26220,AT4G30170,AT5G08680,AT5G17820,AT5G20080,GAPC1,JAL31,JAL34,P1,PCAP1,TPI |
| GO:0030246 | carbohydrate binding | 4 | 306 | 0.0085 | AT1G78850,AT4G16260,JAL31,JAL34 |
